# TF-COMB - discovering grammar of transcription factor binding sites

**DOI:** 10.1101/2022.06.21.496953

**Authors:** Mette Bentsen, Vanessa Heger, Hendrik Schultheis, Carsten Kuenne, Mario Looso

**Affiliations:** Bioinformatics Core Unit (BCU), Max Planck Institute for Heart and Lung Research, Bad Nauheim, Germany; Cardio-Pulmonary Institute (CPI), Bad Nauheim, Germany

**Keywords:** Transcription factors, co-occurrence, grammar, networks, gene regulation

## Abstract

Cooperativity between transcription factors is important to regulate target gene expression. In particular, the binding grammar of TFs in relation to each other, as well as in the context of other genomic elements, is crucial for TF functionality. However, tools to easily uncover co-occurrence between DNA-binding proteins, and investigate the regulatory modules of TFs, are limited. Here we present TF-COMB (Transcription Factor Co-Occurrence using Market Basket analysis) - a tool to investigate co-occurring TFs and binding grammar within regulatory regions. We found that TF-COMB can accurately identify known co-occurring TFs from ChIP-seq data, as well as uncover preferential localization to other genomic elements. With the use of ATAC-seq footprinting and TF motif locations, we found that TFs exhibit both preferred orientation and distance in relation to each other, and that these are biologically significant. Finally, we extended the analysis to not only investigate individual TF pairs, but also TF pairs in the context of networks, which enabled the investigation of TF complexes and TF hubs. In conclusion, TF-COMB is a flexible tool to investigate various aspects of TF binding grammar.

**Graphical abstract:** 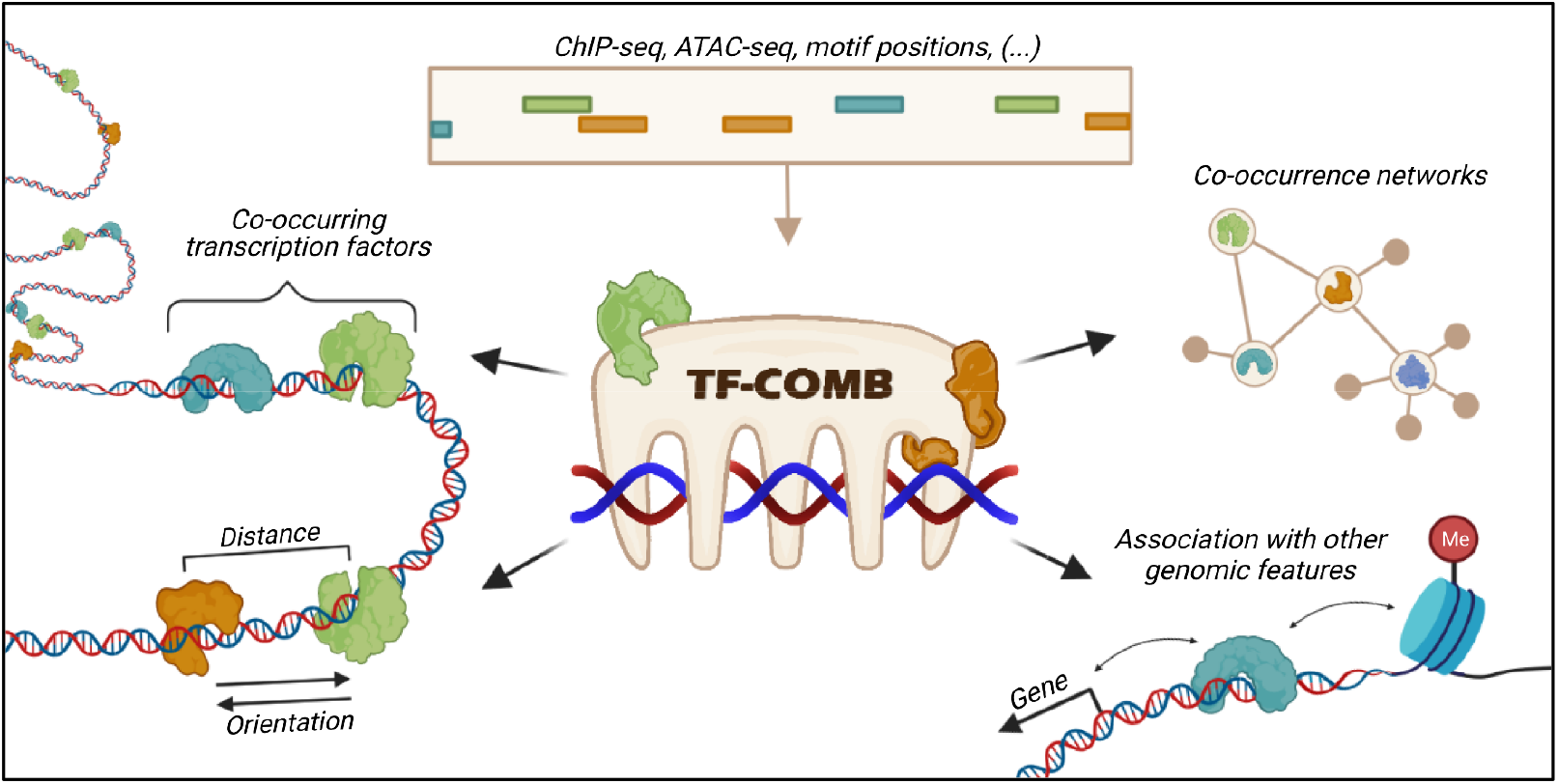

## 1. Introduction

Epigenetic regulation of gene expression is crucial for all processes occurring in cell biology, including cell differentiation and cell development among others. In this context, transcription factors (TFs) are a well-known class of DNA-binding proteins regulating gene expression, operating in direct proximity to target genes, as well as through regulatory enhancer elements further up-or downstream. While some TFs are known to be specific for a certain cell type or a dedicated cellular process, in many cases, combinations of multiple TFs are needed to elicit a specific response - a concept known as TF co-occurrence (Figure 1a). In fact, the amount and types of involved TFs varies, and in higher eukaryotes, it has been shown that 10-15 TF binding sites (TFBS) are required to target a specific gene [1], which can differ in terms of the interaction mode. In the context of enhancers, different models of TF-enhancer interactions have been described [2], which include the enhanceosome, the billboard and the TF collective models (Figure 1b). Of these, the enhanceosome is defined by a certain arrangement, order, distance, affinity and/or relative orientation of the given TFBS. Such rules for binding are known as *TF binding grammar* (please see [3] for a review). In contrast, the billboard and the TF collective models allow for a more flexible grammar where TFBS arrangement and TF binding states are not restricted. In these models, protein-protein interactions (PPI) can enable co-occurrence of TFs through dimerization and binding of additional TF co-factors, but also allow for TFs to co-occur without direct physical interactions. Additionally, the binding of TFs is influenced by other factors including the presence of histone modifications, chromatin accessibility and 3D chromatin organization. A smaller subset of TFs, known as pioneers, are even able to influence chromatin accessibility by opening previously closed chromatin regions, thereby initiating and enabling subsequent binding of other factors [4]. Thus, a higher level of enhancer organization is a prominent factor in gene regulation.

**Figure 1:**
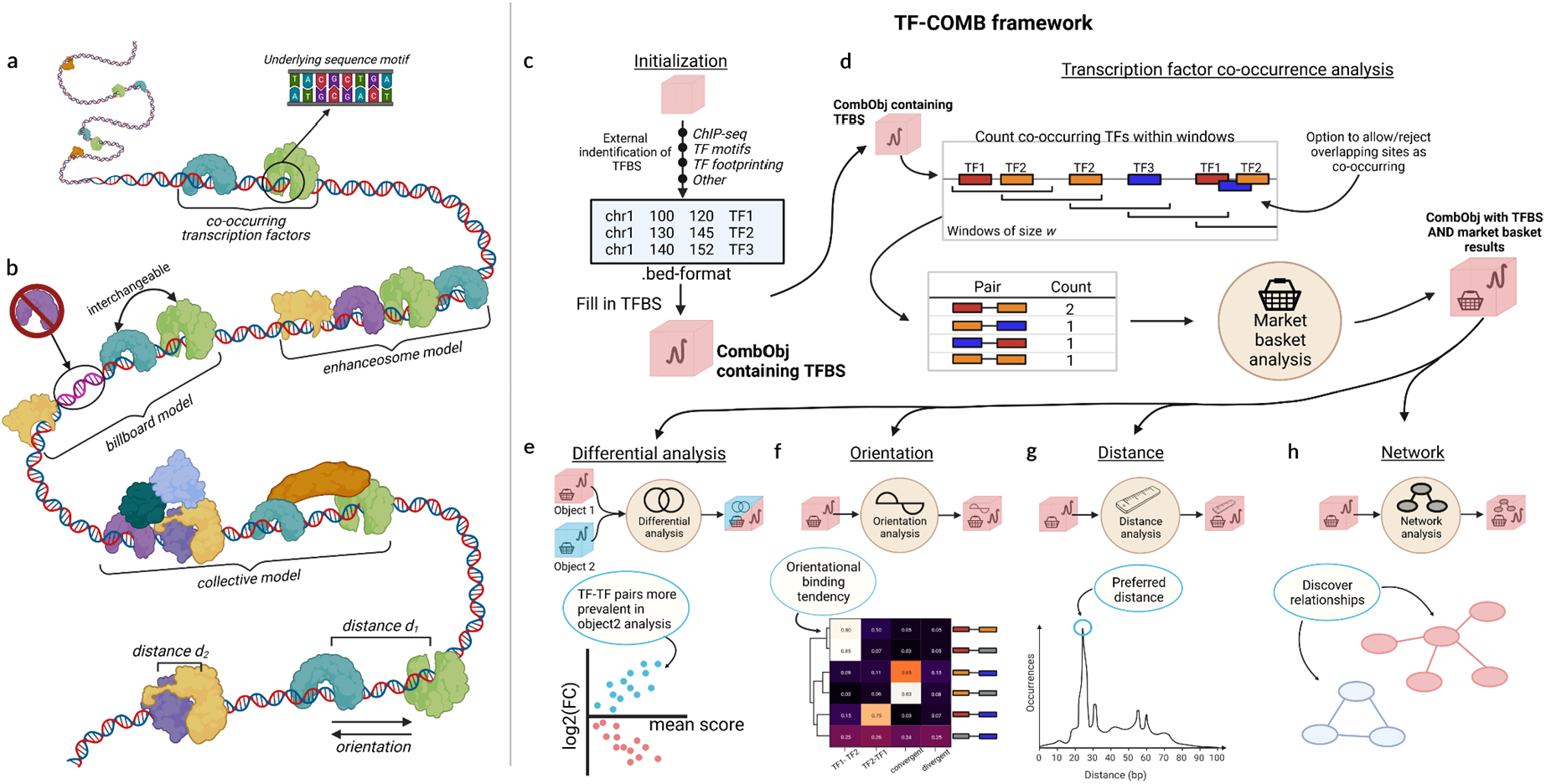
Grammar of regulatory elements and the TF-COMB framework. a) Concept of two co-occurring transcription factors (green + blue) bound in immediate proximity directly to the DNA. b) Models of TF-enhancer interactions and TF binding characteristics. The enhanceosome is defined by strict positioning of TFs, whereas the billboard allows for interchanged positions (green + blue) and absent TF factors (purple). The collective model allows TFs to bind on top of other factors (dark orange, light blue, dark green). TF pairs also exhibit additional characteristics, such as preferred binding distance (d_1_ and d_2_), as well as binding in different orientations on the DNA. Drawn with inspiration from [2]. c-h) The TF-COMB framework: c) Initialization of a TF-COMB object (red square) by providing TFBS and regions of interest from any data origin (e.g. ChIP-Seq, footprinting, ATAC-peaks). d) Co-occurrences are identified, counted and analyzed with an adapted market basket analysis, and are stored in the object for further analyses. e) Differential analysis module allows for the comparison of two independent TF-COMB objects. The module visualizes data to indicate TF pairs more frequent in Object1 (red dots) and Object2 (blue dots) respectively. f) The orientational binding module calculates strand specificity of TF pairs and visualizes preferences via heatmaps. g) TF pairs are analyzed in context of their binding distance, and pairs with prefered binding distance are classified and visualized as histograms. h) Network analysis and visualization module allows to identify higher order relationships between TFs and/or other features. All subfigures were created with BioRender.com.

The detection and analysis of TF binding grammar is complex, as the number of known human TFs is reported to be ∼1600 [5], genes are reported in the range of ∼22000 protein-coding genes [6], and enhancers are reported in the range of millions [7]. Moreover, the collection of active TFs, as well as the chromatin organization, differs for each single cell type. To detect TF co-occurrence, one can broadly distinguish between experimental and computational methods. In the case of experimental methods, the de-facto gold standard in the field are chromatin immunoprecipitation based assays followed by deep sequencing (e.g. ChIP-seq and CUT&RUN-seq). These assays utilize TF specific antibodies and are performed in one cell type at a time. While it is technically possible to run these assays multiple times on different factors of interest, the cost and the dependency on highly specific antibodies generally render these assays unsuitable to investigate global TF co-occurrence. Due to these limitations, the topic of co-occurring TFs have preferably been investigated by *in silico* methods, which utilize a variety of statistical methods [8], linguistic models [9] and enrichment-based algorithms [10]. Most of these methods are based on association analysis of two TFs, and thus simplify the complexity of co-occurring TFs to one specific pair at a time. Additionally, while most of the available tools perform TF motif searches to screen the genome for potential co-occurring events, they are mostly restricted to a single TF anchor point, derived from e.g. ChIP-seq. These limitations motivated us to design a tool that would enable global TF co-occurring analysis independent of data origin. In this context, chromatin accessibility data such as ATAC-seq is of high interest, as we and other groups have recently shown that ATAC-seq can be utilized to find TF binding sites via genome-wide TF footprinting [11, 12]. In addition, co-occurrence of TFs with histone modifications, locations of genes, and other genomic elements are likewise important to accommodate when analyzing TF binding as an epigenetic mechanism. Here we introduce the TF-COMB (Transcription Factor Co-Occurrence using Market Basket analysis) framework, which uses an enhanced market basket analysis to identify and investigate the grammar of co-occurring TFs from a variety of data sources.

## 2. Approach of TF-COMB

In order to detect co-occurring TF binding sites, TF-COMB utilizes an association analysis known as market basket analysis (MBA) [13]. This method has classically been applied to identify shopping habits, also known as association rules, such as “if the customer buys cereal, they are likely to buy milk”. The same approach can also be applied to TF co-occurrence analysis as “if TF1 binds, it is also likely that TF2 binds”, for which the association rule is called a TF co-occurring pair. However, the classical MBA carries a number of shortcomings in the context of TF binding data. For example, the classical MBA reduces items occurring multiple times per transaction to one, and would thus mask the effects of robustness by TFBS replicates in biological networks. In addition, previous applications of MBAs to TF data have likewise excluded all overlapping TFs [14], which influences the discovery of TFs binding in dimers and complexes, where the binding sites might overlap. Moreover, in the context of binding grammar, the order and orientation of TF binding is also of high importance, which has not been taken into account in previous implementations of MBAs. Thus, the current state of publicly available software lacks support for certain aspects needed to characterize the effects of TF co-occurrence. The TF-COMB framework is intended to overcome these limitations.

Firstly, TF-COMB counts all co-occurring TFs in a predefined genomic window. As the source of binding data is flexible, the analysis can be performed for both ChIP-seq peaks, motif positions, and footprints, as well as other input regions. Rather than setting up non-overlapping genomic regions, TF-COMB utilizes sliding windows beginning at each given TFBS, allowing to count all co-occurring TFs relative to that position. A default window size of 100bp was used throughout this paper, however, the parameters of the framework can be changed in order to control the size of the window (minimum and maximum distance), whether to count TFs more than once, as well as basic grammar parameters of TFs such as the inclusion of directionality and strandedness of TF pairs. Thus, the framework is very flexible in terms of investigating a variety of aspects of co-occurring TFs.

Depending on the input data, the total number of TF binding sites and the resulting number of TF combinations can be immense, which gives rise to the need for a measure to rank biologically or statistically important TF combinations. Using the information of pairwise counted TF1/TF2 pairs, different association metrics are supported. The classical MBA utilizes the support, confidence and lift measures to filter and rank interesting associations, however, additional scoring schemes exist, such as the cosine association score [15]. Of note, there is no established method of choosing the correct threshold for association scores of MBA [16]. To overcome this limitation, TF-COMB calculates a Z-score of significance, which helps to reduce false-positives from TFs which are ubiquitously present across the whole genome. This is done through a null-model of random co-occurrences, which is calculated by shuffling the TF labels, rather than randomly shuffling the positions across the genome, as TFBS positions naturally appear in clusters [17]. In summary, the TF-COMB supported metrics allow to rate and select a subset of TF co-occurrences of interest.

## 3. Materials and methods

### 3.1. Availability and implementation

TF-COMB is a Python package intended to be used as a toolbox within Jupyter notebooks or within custom analysis scripts. Due to the high computational needs, it is supported by C-code through Cython [18] integration in Python. Additionally, given functions support multiprocessing when applicable.

TF-COMB is open source and freely available on github at: https://github.com/loosolab/TF-COMB. Details on the individual TF-COMB modules are given at: https://tf-comb.readthedocs.io.

### 3.2. Sequencing data

We obtained TF ChIP-seq peaks, histone ChIP-seq peaks, RNA-seq and ATAC-seq for cell lines HepG2, K562, HEK293, GM12878, MCF-7, H1, A549 and HeLa-S3 from ENCODE [19]. The cell lines were chosen based on a requirement of at least 50 unique TF ChIP-seq experiments available. In case of more than one available experiment per cell line and/or ChIP target, the most recent experiment was used. For ChIP-seq, all peaks were centered at the peak summit and reduced to 1bp regions, while peaks overlapping blacklisted regions were excluded. For ATAC-seq, individual replicates were merged per experimental condition. Accession numbers for all ENCODE datasets used are given in Supplementary Table 1. Additionally, we obtained genomic coordinates for HiC anchor regions for cell line GM12878 (GEO accession GSE63525; file “GSE63525_GM12878_primary+replicate_HiCCUPS_looplist.txt”) [20].

### 3.3. Transcription factor motifs

Motifs were obtained from the JASPAR database (JASPAR 2022 CORE vertebrates) [21]. Annotated dimers (e.g. “Ahr::Arnt”) and any additional motif variations for the same TF (e.g. “var.2”) were excluded.

### 3.4. Metadata for TF interactions

For validation of TF pairs, we obtained a variety of metadata. Known PPIs were obtained from Biogrid [22]. Protein sequences for individual TFs were obtained from UniProt [23] and pairwise protein similarity was calculated using EMBOSS Stretcher [24]. Literature association of TFs was calculated by querying PubMed abstracts and titles for the common presence of each TF pair. The global counts of publications containing either TF1, TF2, and/or TF1+TF2 were used to calculate the cosine similarity measure representing the PubMed association score. Transcription factor families were obtained from AnimalTFDB (v3.0) [25] and using this classification, TF pairs were estimated to be either same-family or different-family pairs. GO-term analysis of TF clusters were performed using the goatools python package [26].

### 3.5. TOBIAS footprinting analysis

Footprinting analysis was performed with the TOBIAS pipeline [11] with default parameters. The input peaks, ATAC-seq reads and motifs were obtained from ENCODE and JASPAR respectively, as explained above. In order to reduce the effect of repetitive elements on the co-occurrence analysis, we used the masked hg38 genome [27].

### 3.6. Comparison to existing tools

In order to compare TF-COMB to existing computational tools, we performed a literature search and obtained 12 *in silico* tools for investigation of co-occurring TFs, taking webservices and command line usage into account (Supplementary Table 2). Namely, these 12 tools were CENTDIST [28], iTFs [29], INSECT 2.0 [30], TICA [31], NAUTICA [32], PC-Traff [9], SpaMo [10], COPS [33], TACO [8], MCOT [34], CIS-Miner [35] and coTRaCTE [36]. Of these, most tools were discarded from comparison due to different reasons including unreachable weblinks. Briefly for all accessible methods, PC-Traff is limited to predefined motifs, CIS-Miner does not provide a functional example of the expected input, COPS is limited to *Drosophila melanogaster* and *Mus musculus* genomes, TACO needs at least two replicate experiments to run and coTRaCTE needs differentially regulated chromatin regions from multiple cell types. The remaining two tools, SpaMo and MCOT, were used for an exemplary analysis based on ChIP data from ENCODE (cell line GM12878; Supplementary Table 1) on 74 TFs and hg38 genome version. We recorded the total runtime on a VM with 64 GB RAM and 8 cores at 2.6 GHz CPU for each tested tool individually. Identified TF pairs were ranked for each tool independently. Resulting lists of co-occurring TFs were aligned where applicable, and top ten exclusively found pairs per tool were manually evaluated via literature search in the PubMed database [37].

## 4. Results

### 4.1. TF-COMB: a universal tool to investigate grammar of enhancers

The typical workflow of a TF-COMB-based analysis is presented in Figure 1c-h. Briefly, the analysis starts with the initialization of an TF-COMB object with regions of interest. As genomic positions in standardized BED file format are supported, TF-COMB can handle (but is not limited to) binding sites from ChIP-seq, pre-calculated motif positions, histone modifications, locations of genes, enhancers, and open chromatin peaks (Figure 1c). In the next step, the genomic positions of the TF-COMB object are internally processed by a sliding window approach and an adjusted MBA is calculated in order to identify TF combinations (Figure 1d). At this stage, the framework provides a variety of analysis and visualization methods, as well as the functionality to compare different conditions with each other (Figure 1e). In order to further examine the TF combination data, the TF-COMB tools provide functionality to investigate TF binding grammar, which includes binding orientation (Figure 1f) and binding distance (Figure 1g), as well as the opportunity to investigate TF pair networks (Figure 1h).

In order to rate the features and performance of TF-COMB, we reviewed 12 existing implementations for TF co-occurrence prediction (Supplementary Table 2; methods segment 3.6). Only tools classified as comparable in functionality and able to run the test data were accepted for assessment. We identified two command-line tools, MCOT and SpaMo, as suitable for comparison with TF-COMB. Where applicable, we aligned individual tool functionality, and we identified a substantial set of features to be exclusively covered by TF-COMB (Supplementary Table 3). Briefly, TF network functionality, binding entity classification (orientation, distance) and quantitative support between conditions of TF-COMB render our software to be a significant extension of existing tools. In order to compare the quality of the results, we ran an exemplary analysis on public ChIP-data and validated the result to known interacting TFs from the BioGrid database [22]. Using a receiver operating characteristic (ROC) curve, we found that TF-COMB has the best predictive ability, with MCOT ranking second best (Supplementary Figure 1a). As expected, we found that SpaMo has a low ability to predict co-occurring TFs, as this tool is particularly focused on identifying motif spacing and not necessarily general motif co-occurrence. In addition to the ROC analysis, we manually rated the top ten candidates per tool via literature search, and found considerably more TF-COMB specific TF pairs verified by literature than for the other tools (Supplementary Table 4). In terms of runtime, we found TF-COMB to outperform the other tools, even though it generates an all-against-all analysis instead of using a single anchor TF (Supplementary Figure 1b). In conclusion, we present TF-COMB as a novel tool for the investigation of TF co-occurrence and TF binding grammar.

### 4.2. TF-COMB detects co-occurring TFs from ChIP-seq data

In order to illustrate the basic functionality of TF-COMB to detect co-occurring TFs, we have utilized the collection of high quality ChIP-seq datasets deposited by the ENCODE project [19, 38]. We collected a total of 1663 ChIP-seq experiments across 8 human cell lines (HepG2, K562, HEK293, GM12878, MCF-7, H1, A549, HeLa-S3) (Supplementary Table 1), and used TF-COMB to find TF associations.

By subsetting pairs based on cosine score and significance (Z-score) across all 8 cell lines, we were able to specify a total of 1938 (1877 unique) TF-TF co-occurring pairs, which correspond to 1-3% of all pairs per cell line (Figure 2a). Within the individual cell lines, TF-COMB predicted the top co-occurring TF pairs to be the well-known pairs MAX-MYC [39] in MCF-7, AP-1 (FOSL2-JUNB) [40] in A549, and CTCF-ZNF143 [41] in H1 cells (Figure 2b-c; Supplementary Table 5). Interestingly, besides highlighting significantly co-occurring pairs, the analysis is also informative in terms of establishing seemingly anti-co-occurring sites, as a negative z-score represents TFs with less co-occurrences than expected. In MCF-7, such pairs included CTCF-FOS, CTCF-GATA3 and CTCF-CEBPB, indicating that CTCF possibly avoids binding to certain partners (Figure 2b). The reason for this might be related to CTCF’s involvement in chromatin looping, as other TFs carry out separate functionalities, which should not interfere with chromatin organization. Thus, the TF-COMB co-occurrence analysis highlights both preferred and unpreferred TF pairs.

**Figure 2:**
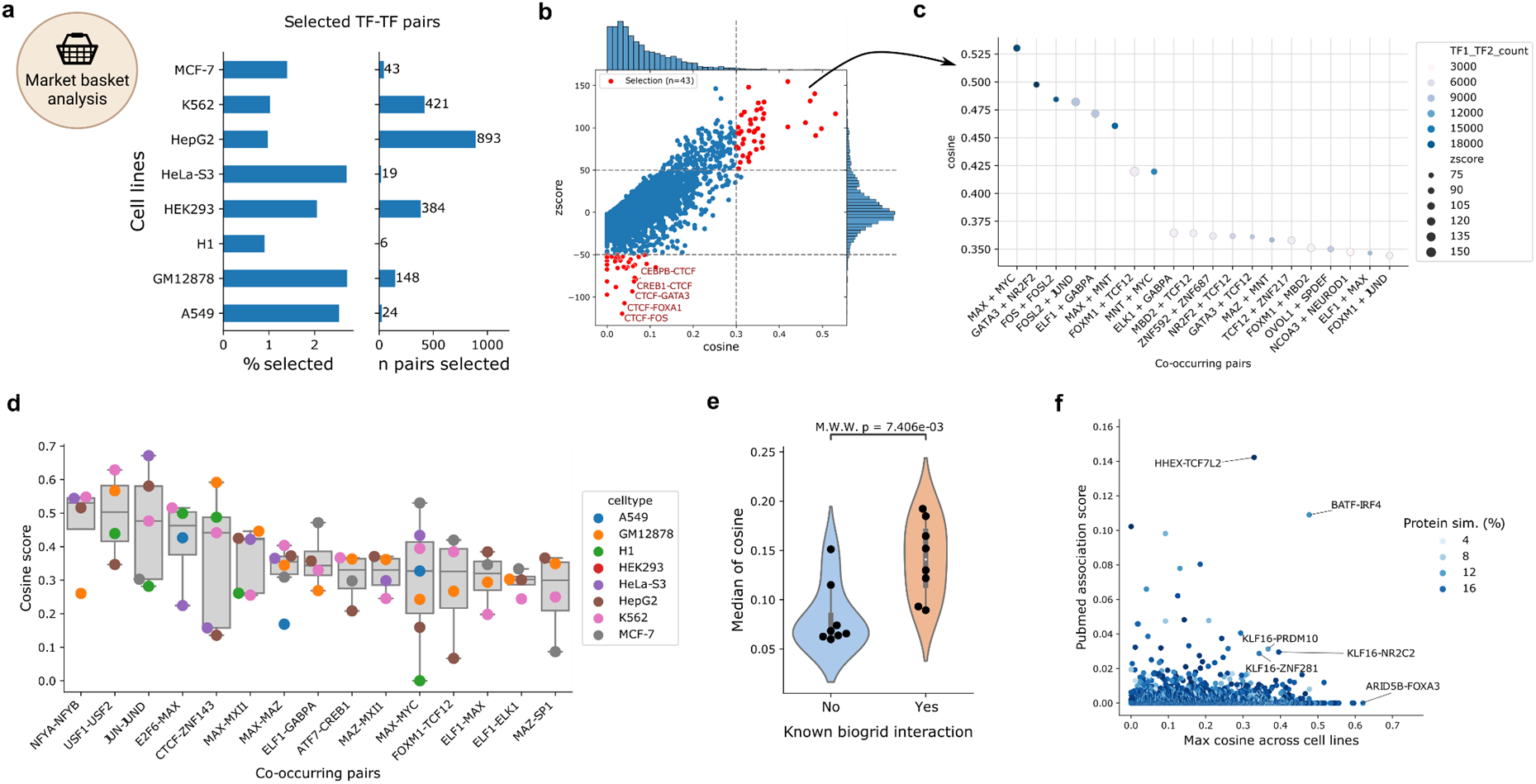
Co-occurrence of TF ChIP-seq peaks in ENCODE cell lines. a) Cell lines and their highly co-occurring TF pairs in percentage and numbers. b) Relationship between cosine and Z-score for each TF pair within cell line MCF-7. The upper right section (red) indicates pairs selected as highly co-occurring. The lower left section contains pairs with less-than-expected co-occurrence. c) Selected co-occurring TF pairs sorted by cosine score. Color indicates the number of TF1-TF2 occurrences, size illustrates the Z-score of the pair. d) The TF pairs identified to have the highest cosine across different cell lines. Only TF pairs present in at least 4 cell lines were included. Each point indicates the cousine score of the given TF pair (column), distribution of scores across cell lines is indicated by a boxplot. e) Violin plot of median cosine score for all TF interactions with (right) or without (left) known protein-protein interactions across the 8 cell lines. The significance of the difference in distributions is calculated using the Mann-Whitney U test. f) Correlation of maximum cosine (across cell lines) and known PubMed association score for TF pairs without known protein-protein interactions. Pairs with high cosine or high PubMed association are highlighted.

Because the list of available ChIP-seq experiments differs for each cell line (Supplementary Figure S2a), with the only TFs available in all cell lines being CTCF and REST, it is not possible to directly compare the top co-occurring pairs across cell lines. For co-occurring TF pairs present in at least 4 cell lines, we found NFYA-NFYB, USF1-USF2, JUN-JUND, E2F6-MAX and CTCF-ZNF143 pairs among others to have the highest median association scores across multiple cell lines (Figure 2d), indicating TF pairs of general importance. In confirmation of this result, we found all these pairs to either form a complex or to interact with each other [41-44]. In addition, by comparing association scores between cell lines, we found that the median spearman correlation is 0.49, suggesting the presence of both common, as well as cell-line specific co-occurring TF-pairs (Supplementary Figure S2b-c). Thus, we conclude TF-COMB to reliably identify known co-occurring TFs across different biological conditions.

In order to further validate the identified ChIP-based co-occurring TF-pairs within individual cell lines, we asked whether the found pairs recapitulate known PPIs from the BioGrid database [22]. Even though physical interactions are not a necessity for co-occurrence, we found that the majority of cell lines exhibit significantly higher association scores for the TF pairs with a known PPI in comparison to other pairs (Figure 2e; Supplementary Figure S2d). However, all cell lines also contain cases of high-association scores without known PPI. To ensure that the default window size of 100bp is sufficient to catch potential PPIs, we ran TF-COMB iteratively with increasing window sizes and correlated the results of each distance to the known PPIs. We found that while most physically interacting TF-TF pairs are collected at distances between 10-50bp(Supplementary Figure S2e), associations are still found for larger windows (Supplementary Figure S2f), meaning that there are other types of TF-TF co-occurrences than those explained by PPI. In this context, we observed that many of the top TF-COMB predictions without a known PPI are TFs from the same families such as FOXA1-FOXA2, MAFF-MAFK and SOX5-SOX13. With the purpose of ruling out cross-reactive ChIP-seq antibodies as the source of this effect, we correlated the association scores with the protein similarity within each TF pair. While we did find some examples of simultaneous high-similarity and high association score, there is no global effect of protein similarity on the co-occurrence analysis (Supplementary Figure S2g). Finally, to explain the association of low similarity TF pairs without known PPI, we also investigated the association of PubMed terms from literature. This analysis could confirm that some pairs, such as BATF-IRF4 were previously described, despite not being annotated with a known PPI in BioGrid (Figure 2f). However, this analysis still leaves a number of pairs including ARID5B-FOXA3, which has high co-occurrence association, but low PubMed association (Figure 2f; lower right). For these, more in-depth analysis will be needed in order to confirm the biological mechanisms of their observed association. In conclusion, we regard TF-COMB as a powerful tool to identify co-occurring TFs, both those physically interacting and those applying other modes of co-occurrence.

### 4.3. Integration of epigenetic marks reveals positional identity of TFs

Besides the expression of certain TFs in individual cell types, multiple epigenetic processes play a role in TF binding, such as the accessibility of chromatin, the presence of histone marks and the 3D chromatin organization. Thus, we sought to use TF-COMB to investigate the co-occurrence of TFs with other epigenetic signals by extending our ChIP-seq co-occurrence analysis with transcriptional co-factors and other DNA-binding proteins (e.g. DNA polymerase II), positions of known histone marks, positional information of genes, chromatin loop anchors from HiC data and open chromatin regions from ATAC-seq. While not all DNA-binding factors in this analysis are strictly identified as TFs, we will still use this term for simplicity.

First, we characterized TFBS in the context of chromatin accessibility and gene promoters. Although gene promoters make up only ∼3% of the entire genome, we found that the majority of factors have promoter association in the range of 10-20%, supporting the enrichment of TF binding to directly regulate gene expression (Figure 3a). In line with TF binding in enhancer regions, we find that the majority of TF binding sites are located in open chromatin outside of promoter regions. However, we also observed that the percentage of sites found in open chromatin is ranging from ∼95% for POU5F1 to below 10% for CBX8, which highlights the differences in functionality of individual TFs and co-factors. Interestingly, we found a number of known pioneer factors including PBX1, GATA3 and FOXA3 to be less associated with open chromatin than other factors, which reflects their ability to also bind to closed chromatin.

**Figure 3:**
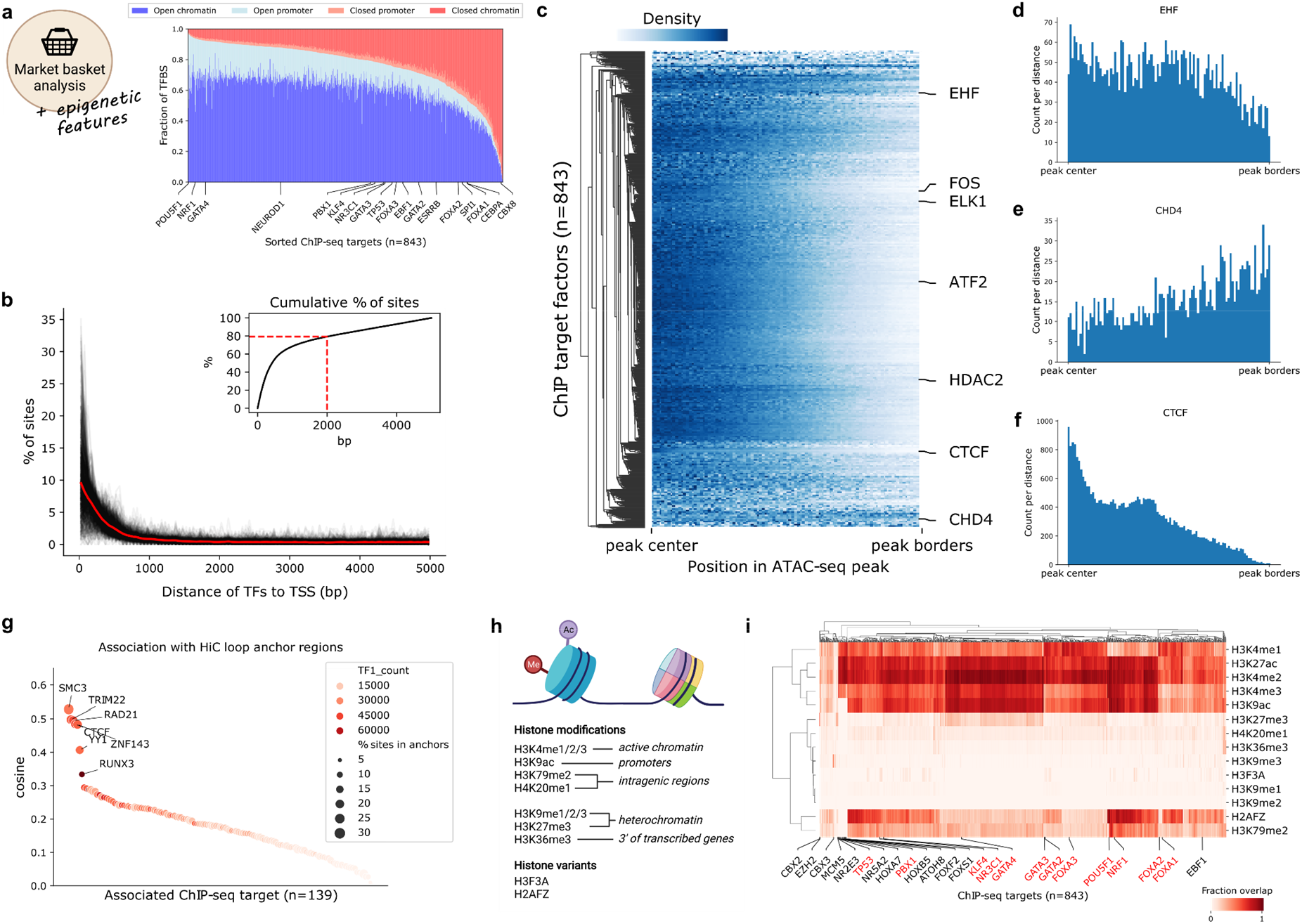
Integration of epigenetic marks reveals positional identity of TFs and co-factors. a) Percentage of ChIP-seq peaks (y-axis) in open/closed (defined by overlap with ATAC-seq peaks) promoter regions (defined as 5000bp upstream of the TSS) and chromatin regions (all regions not defined as promoter) for individual factors (x-axis). b) Distance of ChIP-seq peak summits to transcription start sites (TSS) of genes. Distributions for individual factors are shown in black, and the mean distance is shown in bold red. Upper right corner shows the cumulative distribution of sites with 80% of sites marked with a dashed line. c) Relative location of ChIP-seq peak summits in open chromatin regions. Counts for the same TF in different cell lines were merged by taking the mean at each distance. The colorbar represents the scaled number of positions found to co-localize at different percentages of the peak length. Counts to the left/right of the peak center are aggregated to a range from center to border. d-f) Relative TF binding positions in ATAC-seq peaks from peak-center (left) to outer peak-borders (right). g) Co-occurrence of TFs and HiC loop regions in the cell line GM12878. TF1_count represents the total count of ChIP-seq peaks per factor. h) Scheme of histone modifications and histone variants investigated. Created with BioRender.com i) Co-occurrence of TFs and histone marks as calculated by fractional overlap of regions. A selection of TFs from interesting clusters are annotated on the x-axis. Known pioneer factors are marked in red.

By performing a distance analysis between the TFBS and annotated genes, we found a strong enrichment of binding sites close to the TSS, with 80% of these binding sites occurring within 2000bp (Figure 3b). In contrast, when we investigated the more global TF binding patterns in the context of open chromatin, we found that the TF binding sites show locational preferences. As TFs are known to bind to open chromatin, it was not surprising that the vast majority of TFs are addressing the center region (+/-25%) of ATAC-seq peaks, such as shown for ATF2, FOS, ELK1 and the histone deacetylase HDAC2 (Figure 3c). However, we also found TFs with their binding sites located without preference across the whole open region (EHF; Figure 3d), as well as some TF candidates with a preference to the outer bounds of open chromatin regions (CHD4; Figure 3e). This localization of CHD4 is well explained by the fact that CHD4 has been shown to slide nucleosomes, which are found at the borders of open chromatin [45]. Overall, the relative locations of TFBS in open chromatin peaks showed a significant correlation between cell lines, which confirms that there are groups of TFs and co-factors with locational preferences regardless of cell type (Supplementary Figure S3a).

Within the group of TFs located at the center of ATAC-seq regions, we also observed CTCF, which has a highly centered peak around the +/-15% core of the peak (Figure 3f). As CTCF is known to mediate chromatin looping [46], we used TF-COMB to investigate the co-occurrence of ChIP-seq defined TF sites with chromatin loop anchor regions as defined by HiC. As expected, we found that CTCF had high co-occurrence with loop regions, and we additionally found SMC3, TRIM22, RAD21, ZNF143, YY1 and RUNX3 (Figure 3g). These results are perfectly in line with previous investigations showing that CTCF, RAD21, ZNF143, TRIM22 and RUNX3 can accurately predict the position of chromatin loops [47]. Additionally, SMC3 is an essential component of cohesin [48], and YY1 has been shown to mediate 3D chromatin interactions in collaboration with CTCF [49]. As seen for CTCF, these proteins likewise show strong positional specificities within open chromatin (Supplementary Figure S3b). As such, the relative positioning of these factors might be an important mechanism for higher chromatin organization.

Finally, we investigated the higher order binding patterns of TFs in the context of activating and repressing histone modifications (Figure 3h). Since the association of histone modifications with TFs is not necessarily symmetrical, we used the association ‘confidence’ score, which represents the fractional overlap between sites. Firstly, we confirmed the association of respective histone marks to chromatin, and not surprisingly, we found the active histone marks H3K4me1/2 and H3K27ac to have the highest association with open chromatin, and in contrast, the repressive histone marks H3K36me3 and H3K9me1/2/3 with the lowest association (Supplementary Figure S3c). Correspondingly, we found that H3K4me2 and H3K27ac have the highest overall association with ChIP-seq defined TF targets (Figure 3i), which has also been described previously [50]. In contrast, the repressing marks H3K27me3, H3K9me1/2/3, H3K36me3, H4K20me1 and histone variant H3F3A had a low overall association with TF binding. However, despite the minimal association with open chromatin and TF binding, we identified factors such as EZH2, CBX2/3/8, ZNF184, MCM3/5, XRCC3, ZNF280A, SRSF9 and PLRG1 to be prominently overlapping with H3K9me3 and H3K27me3, while simultaneously being depleted for association with active histone marks (Figure 3i). Thus, these proteins have an ability to bind in otherwise inactive chromatin, which the majority of other proteins do not. Indeed, a number of these TFs are members of the Polycomb Group of proteins (namely EZH2 and CBX family proteins), which assemble in multi-protein complexes to repress genes.

Interestingly, this analysis also highlighted another prominent cluster of TFs, which is defined by an overall strong association with active histone marks, but with an exclusive depletion of association with H3K9ac and H3K79me2, which are markers for active promoters and intragenic regions respectively. This cluster contains several HOX and FOX factors, nuclear receptors NR2E3 and NR5A2, as well as PBX1, ATOH8 and TP53 among others. Many of these factors are known to be pioneer factors (TP53 and PBX1) or part of families with many known pioneers (nuclear receptors and the FOX family) [51]. However, while some other known pioneers also show a decrease in association with H3K9ac (e.g. FOXA1/2), it is not an universal rule (e.g. NRF1), and the pioneer hypothesis is therefore not the only explanation for the depletion of H3K9ac for this cluster. Alternatively, the effect could be explained by the role of these factors in controlling lineage specification, as is well described for the HOX factors [52]. Thus, the discovery of TFs specifically co-occurring or restricting binding to certain histone modifications, can uncover hallmarks of TF binding to enhancers.

In conclusion, we find that TF-COMB analysis integrating epigenetic signatures uncovers DNA-binding proteins with locational specificity corresponding to individual biological functions.

### 4.4. Co-occurrence analysis utilizing TF footprinting

While we were able to use gold standard ChIP-seq data to identify positional locations of TFs within larger regulatory regions, this data comes with some fundamental challenges in the context of investigating local binding grammar. Mainly, TF ChIP-seq peaks are several hundred base pairs wide, and thus do not clearly indicate the exact location of TF binding sites. As a result, ChIP-seq will generally fail to find multiple TF sites from one factor in close proximity, and will lose the information of TF binding orientation, which impedes the investigation of a higher order of TF binding grammar from ChIP-seq data. In contrast, the identification of TFBS through methods such as motif prediction or digital genomic footprinting requires only one chromatin accessibility assay per cell type to estimate binding events for hundreds of TFs in parallel, while preserving location and orientation of the TFBS (Figure 4a). Thus, we obtained ATAC-seq experiments for cell lines A549, GM12878, HepG2, K562, MCF-7 from ENCODE, and ran our previously published ATAC-seq footprinting pipeline TOBIAS on the data [11]. The pipeline identifies bound TFBS on the basis of Tn5 insertion patterns, which were used to subsequently find co-occurring TFs with TF-COMB.

**Figure 4:**
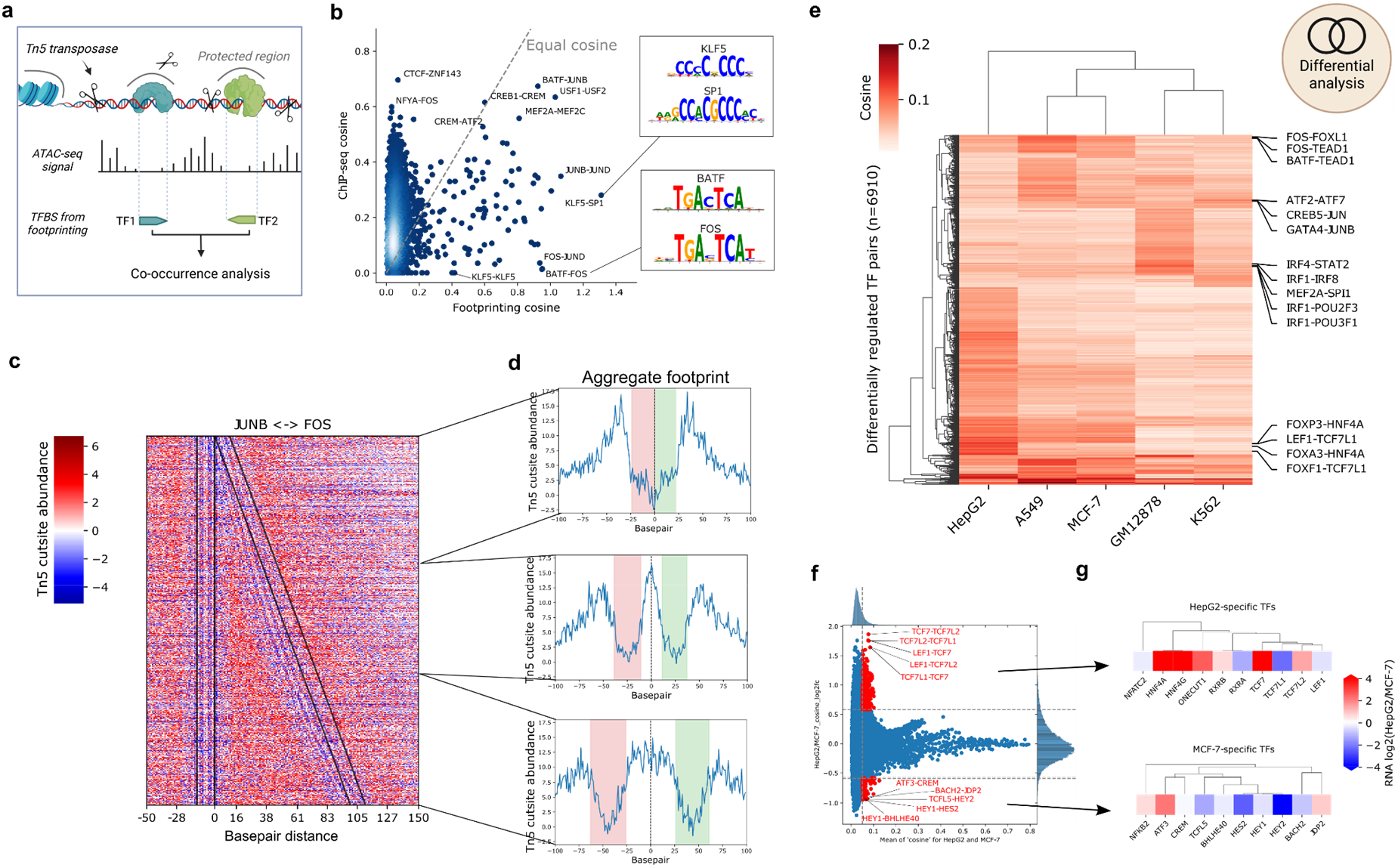
Footprinting data uncovers cell line specific TF co-occurrence. a) Scheme of TF co-occurrence and Tn5 mediated digital genomic footprinting. Prepared using BioRender.com. b) Direct comparison of scores derived for TF-pairs via ChIP-seq and ATAC-seq (footprinting) analysis. Two pairs with high footprinting scores are highlighted and corresponding motifs are illustrated. c) Footprinting heatmap of all co-occurring JUNB-FOS sites. Colored for Tn5 cutsites appearing more than expected (red) or less than expected (blue) if DNA is inaccessible. Black lines represent the edges of JUNB (left) and FOS (right) motifs. Edges show the binding strand of the respective TF. d) Aggregated views of the scores shown in e). Increasing distance (top to bottom) causes the combined TFs footprint to split into two distinct ones. e) Heatmap showing cosine scores for differentially co-occurring TF pairs across five cell lines. A subset of prominent cell line specific pairs are labeled on the right side. f) Activity of TF-pairs in direct comparison between HepG2 and MCF-7 cell lines. Significantly changed TF pairs are marked in red. g) Differential RNA expressions of the top 10 TFs selected in f) for each group. TFs are clustered by motif similarity.

Overall, we find both coincident and differing TF pairs when comparing the co-occurrence of ChIP-seq based and footprinting-derived TFBS (Figure 4b). For example, we observed that well-known factor pairs BATF-JUNB, USF1-USF2 and CREB1-CREM were found to have high cosine scores in both footprinting and ChIP-seq analysis. However, we also observed some pairs specifically found in ChIP-seq data including CTCF-ZNF143 and NFYA-FOS. As the footprinting analysis is dependent on the presence of sequence motifs, we performed a global analysis of the presence of motifs within ChIP-seq peaks, and found the match between ChIP and motifs to be very different across factors (Supplementary Figure S4a). For example, while a high percentage of CTCF sites contain at least one motif, less than 10% of ZNF143 ChIP-seq sites contain known motifs, thus making the CTCF-ZNF143 difficult to discover from motif-based data. As such, the lack of co-occurrence between ChIP-seq and footprinting-derived sites can be partially explained by a lack of identifiable motifs within these ChIP-seq peaks.

In contrast, there are also a number of co-occurring pairs which are more commonly found in footprinting data, including FOS-JUND, ATF3-ATF7 and KLF5-SP1, all of which have very similar motifs. We also observed self-pairs such as SP1-SP1, which we could not observe in ChIP-seq due to the lack of resolution of ChIP-seq peaks. When analyzing the underlying genomic sequence, we indeed found multiple SP1 motifs in the vicinity of one SP1 ChIP-seq peak, which naturally increases the cosine scores of this pair for footprinting data (Supplementary Figure S4b). In general, we found that the co-occurrence scores for footprinting data are correlated with motif similarity to a much higher degree than the ChIP-seq derived data (Supplementary Figure S4c). While this is expected due to motif overlap, the correlation to motif similarity persists, even when overlapping between sites is disallowed, which suggests that the effect is not only due to direct motif overlap. As is the case for SP1, this effect can arise due to multiple motifs within peaks, as exemplified by NFYA-NFYC (Supplementary Figure S4d). Whereas the association of two ChIP-seq peaks is only counted once, the co-occurrence of similar motifs will lead to high scoring pairs for NFYA-NFYA, NFYC-NFYC and NFYA-NFYC, as these are counted multiple times within the same window. Thus, TFs do in fact co-occur with similar motifs on two levels; firstly by direct motif overlap, and secondly by multiple copies of a motif in close proximity. The latter case suggests a certain importance of a genomic loci for an individual factor, as multiple binding sites provide an increased probability of binding - even in the event of mutations. In conclusion, co-occurrence of footprinting data reflects ChIP-seq derived data analysis, and additionally unravels genomic sequence compositions that utilize motif redundancy at target regions.

The gain of resolution by utilizing footprinting data additionally allows for a TF distance analysis as exemplarily shown for the TF TBP, which has a preferred distance of 16bp to the TSS (Supplementary Figure S4e), while the ChIP-based analysis did not show any preferred binding distances to TSS (Supplementary Figure S4f). We asked whether this increased resolution also enables the visualization of paired TF footprints and therefore utilized the TF-COMB plotting module for paired TF sites (Figure 4c). As exemplarily shown for the well characterized TF pair JUN-FOS, sites with close motif distances create one common footprint, and by increasing the motif distance of bound sites, the individual footprints appear distinguishable starting from distances above 20bp (Figure 4d). As footprints are generated by Tn5 transposase cutting patterns, we conclude that the Tn5 transposase is unable to insert adapters between closely bound TFs. Thus, in the context of footprinting analysis, individual footprints might not be directly mappable to single TFs, but might be the result of several closely bound TFs. This is an important factor to take into account for future footprinting algorithms.

As ChIP-seq data is limited to certain factors in each cell type, it can be tricky to compare co-occurrences between datasets. However, footprinting analysis contains the same TF motifs across all cell types, and just differs at the respective footprinting score levels. Thus, we used our previously calculated co-occurring TFs based on TOBIAS footprints and added a differential analysis to the TF-COMB object in order to quantify co-occurring TF pairs between cell lines in a global manner. As expected, we found the majority of TF pairs commonly active across cell lines (spearman correlations 0.8-0.9)(Supplementary Figure S4g), however, by selecting the enriched TF pairs from each contrast, we identified 3.2% (n=6910) of the potential TF pairs as differentially active between cell types (Figure 4e). Not surprisingly, the overall clustering of the cell lines reflected the respective cell origin, by grouping epithelial cells (A549 and MCF7), as well as the lymphocyte cell lines (GM12878 and K562). The cell line specific TF pairs nicely mapped to the biological background, which included FOXA3-HNF4A for HepG2 cells, which are well known liver TFs able to program fibroblasts into hepatocyte-like cells [53], and multiple pairs containing IRF for GM12878-cells, which supports the importance of these factors in lymphocyte differentiation [54].

Focusing on the prominent changes between HepG2 and MCF-7, we used TF-COMB to highlight ∼1% differential TF pairs specific for this contrast (Figure 4f). The changes in expression (log2FC RNA-seq) of the top 10 TFs between these cell types likewise showed the majority of the TFs to be upregulated in the cell type, for which they participate in co-occurring pairs (Figure 4g). In contrast, we observed TCF7L1 in many co-occurring pairs in HepG2, while it is actually downregulated on RNA level in comparison to MCF-7. This effect might be driven by motif similarity between TFs, as we see that both TCF7 and TCF7L2, which have highly similar motifs to TCF7L1, are upregulated in HepG2. Thus, the use of motifs makes it difficult to directly link motif activity with a certain TF, but integration of TF expression data might help to uncover which TF is most likely to be the participating partner in a co-occurring pair.

In summary, we conclude the application of co-occurrence analysis to digital genomic footprinting data to be a valuable approach for uncovering global changes of TF co-occurrence and TF binding grammar between biological conditions. In addition, the association of motifs allows to untangle TF relationships driven by motif similarity and motif redundancy.

### 4.5. TF binding grammar encodes biological relevance

Considering the higher resolution and completeness of TF binding activity for motif derived data, we asked if we are able to infer detailed information on TF binding grammar in the context of local binding site arrangement. Literature has several examples of highly regulated enhancers with specific distances and strandedness of TFs, such as the NFYB-USF1 pair, found at a preferred binding distance of 17-18bp in a converged orientation [55] or a preferred distance of 37bp for the CTCF-ZNF143 pair [41]. Therefore, we sought to use TF-COMB to investigate whether these exemplary binding characteristics are rare, or constitute a more global property of TF pairs that allow for a classification of TF pairs and enhancer organization. To be able to uncover the global presence of grammar in the genome, and not only for the sites which are predicted to be bound in the footprinting analysis, we used the full set of motif positions within the HepG2 ATAC-peak regions as input for TF-COMB.

Firstly we analyzed the global directionality of TF pairs, and found that a subset of ∼2% of TF pairs exhibit preferential directionality with more than 60% of sites in either the same or opposite orientations (Figure 5a). Of these, the orientations can be further divided into groups of TF1-TF2/TF2-TF1 and convergent/divergent for same and opposite groups respectively. While some TF pairs show equal distribution of the subdivided groups, the split highlights an additional preference for the exact order of binding (Figure 5b).

**Figure 5:**
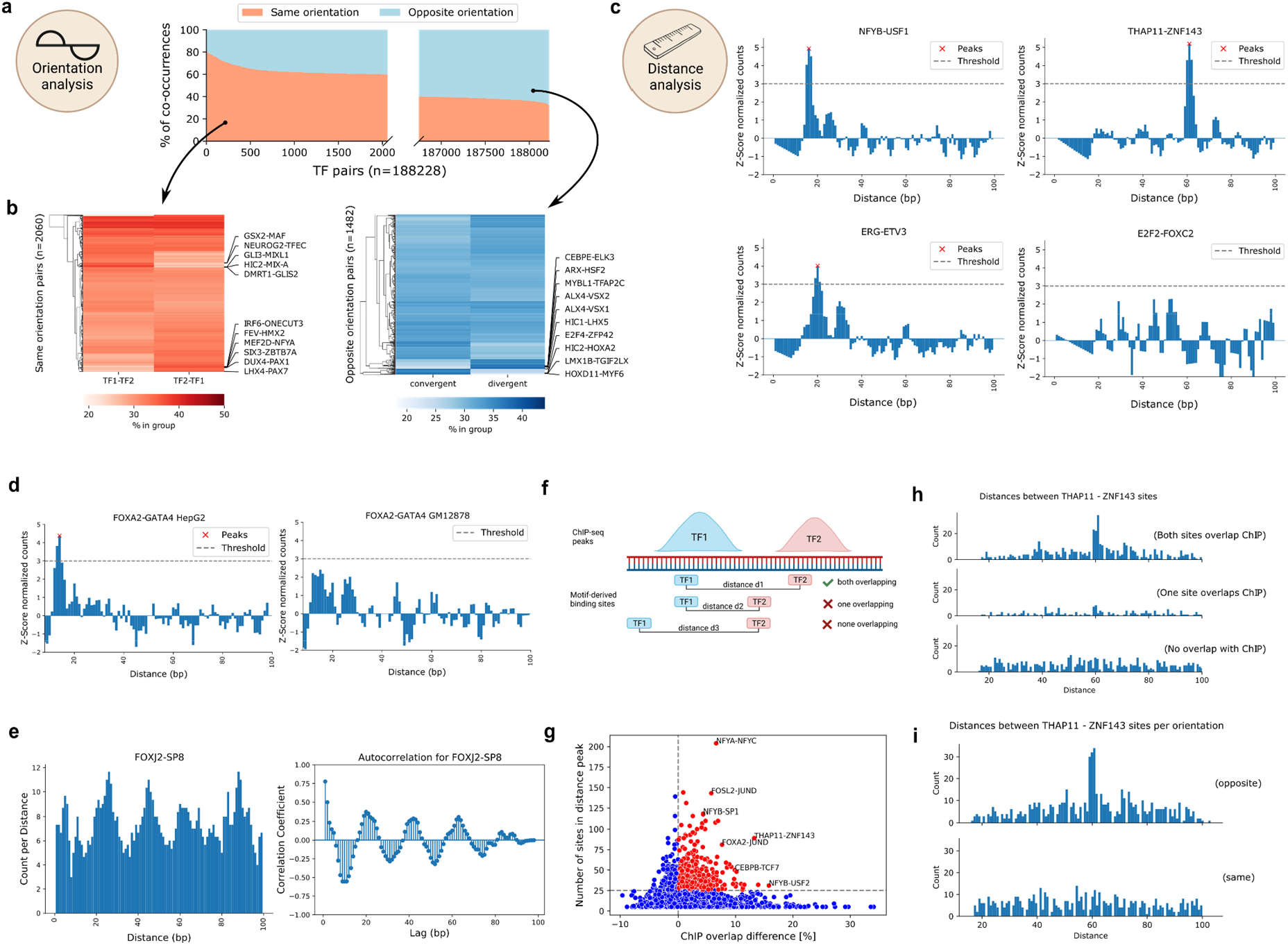
TF pairs exhibit local binding grammar. a) Percentage of TF-pair locations (y-axis) with both TFs on the same or opposite strand. X-axis gives the ranking of TF pairs with regards to orientation. Only pairs with more than 60% for either group are shown (axis is not continuous). b) Percentage of TF-pair locations derived from a), splitted by TFs orientation. Red heatmap highlights top pairs with orientation on the same strand, blue heatmap highlights top pairs with orientation on opposite strands. c) Z-score normalized TF-pair binding counts sorted by distance. Peaks above threshold are called by TF-COMB and considered preferred binding distance. d) Difference in binding distance distribution for FOXA2-GATA4 in HepG2 (left) and GM12878 (right) cells. e) Binding distance periodicity of the FOXJ2-SP8 pair. Left plot shows the distribution of binding site distances. Right plot shows the calculated autocorrelation for the signal, indicating a lag of 20bp. f) Scheme of motif-derived binding sites overlapping with ChIP peaks. Prepared using BioRender.com. g) Difference in ChIP-overlap fraction (x-axis) between preferred distance sites and no preferred distance sites per pair and power (y-axis). The most prominent pairs are annotated. h) Number of sites by distance between THAP11 and ZNF143 binding sites. Subplots indicate split of loci into groups as indicated in f). i) Number of sites by distance between THAP11 and ZNF143 binding sites. Upper and lower plots indicate strand orientation.

Next, we analyzed the preferred distance between TFs using TF-COMB, and found that 36.6% of all TF pairs exhibit at least one preferred binding distance (Supplementary Figure S5a). For the majority of pairs with a preferred binding distance, exactly one distance was predicted (35.5%), while the remaining pairs (1.1%) exhibited multiple distances (Supplementary Figure S5b). Among the candidates with predicted preferred binding distances, we found well established pairs, like THAP11-ZNF143 [56], NFY-USF members [57] (Figure 5c) and BATF-JUN [58, Murphy, 2013 #79] (Supplementary Figure S5c). In addition, among others, we found not yet described pairs like ETV3-ERG. In contrast, E2F2-FOXC2 is an example for a TF pair with no predicted preferred distance (Figure 5c). This highlights that preferred distances between TFs is a global property applicable when investigating binding grammar. Furthermore, this might also hint to additional characteristics of grammar, such as changes between biological conditions. Exemplary, the FOXA2-GATA4 pair, which was recently described as liver specific [4], differs between HepG2 and GM12878 cell lines (Figure 5d). In contrast, the ubiquitous pair NFYB-NFYC, which is known to form the trimeric NFY complex in collaboration with NFYA [42], remains similar between different cell lines (Supplementary Figure S5d). Thus, parallel to the general co-occurrence analysis, the individual TF binding distances are also indicative of cell line specific co-occurrence. In addition, by analyzing all distances per TF pair, we detected that many pairs have distributions of distances which seem to occur with certain periodicity. One such example is the FOXJ2-SP8 pair, which displays an apparent structure in the distribution of binding sites, as it translates to a period of ∼20bp between two peaks (Figure 5e). In contrast, we find other pairs with differing periods (Supplementary Figure S5e), indicating that individual pairs exhibit different binding preferences.

In order to evaluate the biological relevance of the preferred distance sites from a TF pair compared to the non preferred distance sites from the same pair, we split the data into three groups. The first group contains all sites corresponding to TF pairs classified to have no preferred binding distance, which we call “no preference sites”. The second group covers the sites for TF pairs classified to have a preferred binding distance, filtered for the “distance peak” sites, called “preferred distance sites”. Finally, the third group contains the remaining sites of the TF pairs from the prior group outside of the preferred binding distance, named “no preferred distance sites”. Firstly, we hypothesized that the preferred distance sites represent important functional units, and are therefore more likely to occur within regulatory features, such as gene promoters. Indeed, after annotating all paired sites with UROPA [59], we found a significant increase of the gene annotation rate for sites with a preferred distance compared to both groups without preferred distances (Supplementary Figure S5f). Next, we asked whether these motif-derived preferred distance sites can be used as a classifier to discriminate between real binding sites and potential binding sites. To this end, we overlapped the motif-derived sites with corresponding ChIP-seq peaks in HepG2 cells (Figure 5f). As suggested by our prior findings on the gene annotation level, we detected a significant increase of overlapping “true” ChIP-seq sites for the preferred distance sites group (Supplementary Figure S5g). Of note, the TF pairs THAP11-ZNF143 and NFY-USF, both well described in literature and already found in earlier sections (Figure 5c), were among the pairs showing the strongest differences (Figure 5g). This finding holds true when visualizing the distance plots for the ChIP-seq overlap of both TF sites, only one TF site, or no overlapping sites, respectively (Figure 5h). Finally, we combined distance and orientation analysis, and exemplarily found that the majority of preferred distance sites for THAP11-ZNF143 are located in opposite directions, exhibiting a preferred distance around 61bp (Figure 5i). This confirms that the preferred distance and orientation encodes for true co-occurrence of both factors.

In conclusion, we find that TF-COMB is able to globally uncover TF pairs that exhibit local binding grammar characteristics such as TF pair distance, and relative TF orientation. The overlap with ChIP-seq derived binding sites suggest biological relevance for the preferred binding distances, a finding that might contribute to future methods utilizing motifs as an approximation for DNA binding.

### 4.6. TF network analysis uncovers regulatory complexes

While we have investigated TF binding in pairs using TF-COMB, TFs often participate in multi-TF modules, which regulate complex biological networks through additive or synergistic binding [60]. Therefore, the discovery of these relationships is crucial for understanding gene regulation. As a result, TF-COMB can utilize the full set of TF rules to deduce a network in terms of nodes and edges. In order to identify potential protein complexes with the network module, we utilized the ChIP-seq data of TFs, co-factors and other DNA-binding proteins for the GM12878 cell line from ENCODE.

After initial filtering, TF-COMB generated a core network consisting of 329 edges (TF co-occurrences) and 93 nodes (TFs). The network view (Figure 6a) uncovers noticeable substructures, including isolated TF clusters not connected to the main network, some barely interconnected substructures with tight internal links, as well as dense subgroups, driven by highly interconnected nodes. In order to quantify this structure, TF-COMB uses the louvain method for community detection [61], which partitioned the exemplary network into 8 clusters (Figure 6a; Supplementary Figure S6a left). Not surprisingly when analyzing transcriptional regulators, GO-term analysis of the individual clusters revealed enrichment of terms such as “chromatin”, “transcription regulator complex” and “nucleus” among others (Figure 6b). However, besides GO-terms indicative of positive regulation, we also identified clusters, such as cluster 4, enriched for terms related to transcriptional repression. Further, we annotated cluster 1 to the cohesin complex, and cluster 3, containing BATF and JUN, to be enriched for “transcription factor AP-1 complex”. Of note, we observed many same-family TFs for cluster 3 (Supplementary Figure S6a right). Thus, we wanted to test whether same-family TF pairs are significantly enriched within individual clusters. Indeed, by randomly selecting pairs within and between clusters, we observed a significant increase in percentage of same-family pairs within network clusters (Supplementary Figure S6b). To summarize, the network analysis enabled us to identify protein clusters and complexes with particular biological functionality.

**Figure 6:**
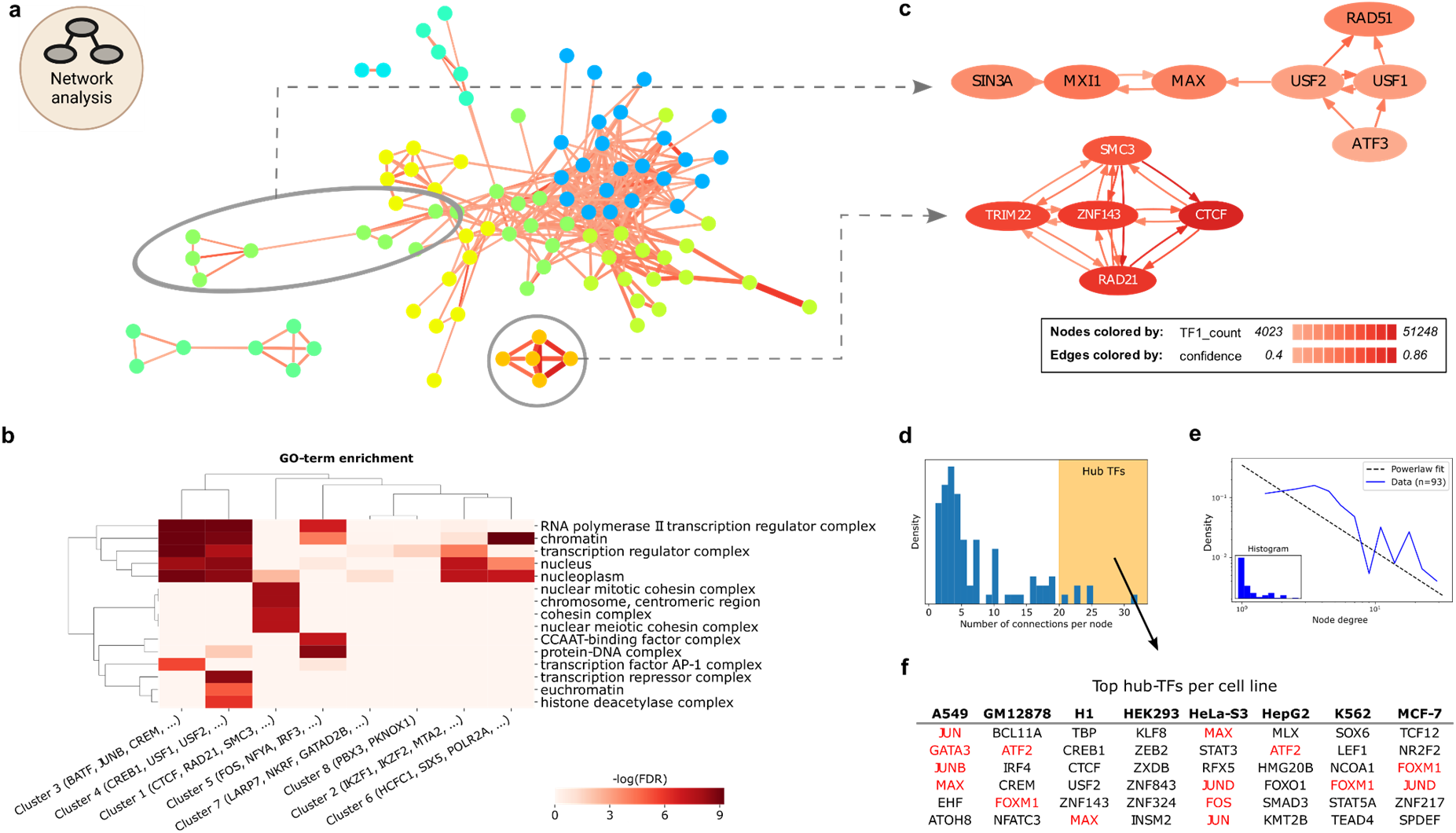
Network approach uncovers TF complexes and hubs. a) The GM12878 co-occurrence network. Nodes represent TFs, edges represent associated TF pairs. Edges are colored by cosine score and sized by the number of associated TF1-TF2 co-occurring sites. Coloring of nodes illustrates Louvain community clustering. Two sub-clusters are indicated for zooming in c). b) GO-term analysis on TF groups extracted from network clusters (columns). Coloring indicates enrichment for GO terms (rows). c) Directional sub-networks from a). Each arrow represents a dependency defined by the confidence score, e.g. ATF3 is dependent on USF2. Node coloring indicates the number of sites assigned to the respective TF. Only edges with confidence scores above 0.4 are shown. d) Distribution of node degrees. High node degree is shaded with yellow in the plot. e) Number of connections per TF (blue), approximates power-law distribution (grey dashed line). f) Table of TFs per cell line with the highest number of interactions (hub creators) from d). Marked TFs (red) are discussed in the main text.

Within the individual clusters, the TF-COMB network analysis can additionally uncover dependency relationships. For this, a directional network using the ‘confidence’ score, which represents the probability that TF2 is found, if TF1 is present, is used. For cluster 1, which includes ZNF143/CTCF/RAD21, all TFs are dependent on each other, which suggests that these factors bind in a protein complex (Figure 6c; upper). However, in the cluster of USF1/USF2/RAD51/ATF3, we found USF1 and USF2 to have directional relationships with ATF3 and RAD51 (Figure 6c; lower). The association of USF and RAD51 is supported by a recent study on the location of USF motifs at RAD51-bound elements [62], however, our analysis additionally indicates a significant number of RAD51 sites to have a completely independent role of USF. Interestingly, the network analysis also reveals ATF3 to be highly connected to USF factors, but not vice versa, with more than 50% of its binding sites in the vicinity of USF binding sites. Moreover, the lack of a link between ATF3 and RAD51 suggests that the interactions with USF are contained in independent regulatory circuits.

Finally, the network representation of TF co-occurrence also allows to draw conclusions on potential TF hubs, defined as TFs with many partners in the network. The distribution of node degrees has an apparent tail, indicating most TFs to have few partners, yet some TFs exhibit many co-occurrences with other factors (Figure 6d). Noteworthy, we found the networks of most cell types to follow a power-law distribution, which is a characteristic of biological networks [63] (Figure 6e, Supplementary Figure S6c). Taking all cell lines into account, the TFs with the highest node-degree include MAX, JUN, GATA3 and FOXM1 (Figure 6f). MAX is known to orchestrate a large network [64]. Likewise, TFs JUN, JUNB, JUND, FOS and ATF2 are all part of the AP-1 family of TFs, which can dimerize, thus explaining the hub characteristics of these TFs. Interestingly, we also find GATA3, which is a known pioneer factor, as well as FOXM1. While FOXM1 is not known to have pioneer activity, previous publication showing an overlap of 71% between FOXM1 and FOXA1 binding events [65], thus allows us to speculate that FOXM1 might indeed be able to function as a pioneer in line with FOXA1.

In conclusion, we found network analysis on TF co-occurrence as a highly flexible tool to explore relationships between transcriptional regulators in a wider perspective and to extract subgroups of factors as a valuable source for the hypothesis of potential TF complexes.

## Discussion & conclusion

This study was performed to demonstrate the functionality and usability of a new software framework, named TF-COMB, intended to gather, analyze, visualize and explore data in the field of co-occurrence and grammar of TF binding. Due to its generalized setup, TF-COMB is able to help to unravel epigenetic related aspects of TF binding grammar, such as the interplay of chromatin accessibility, histone modifications, and gene locations, as well as local grammar in terms of distance and orientation of TFs. In addition, TF-COMB allows a broader look at the interdependencies of TF co-occurrences in a global network context, which allows to draw conclusions on the binding of TFs not only in pairs, but also in larger compositions, including e.g. protein complexes.

The exemplary investigations on widely accepted gold-standard datasets from ENCODE are intended to illustrate potential analysis workflows provided via TF-COMB. To this end, we used TF binding data of both ChIP-seq peaks and motif sites derived from ATAC-seq based TF footprinting to identify commonalities, as well as unique aspects that can be derived for each data source. A major strength of ChIP-seq data was found to be the independence of TF sequence motifs [66] enabling to calculate co-occurrence of TFs with co-factors and other DNA-associating factors without known sequence motifs (e.g. RAD21, SMC3, TRIM22). In contrast, we demonstrated the advantage of TF footprinting and motif positions in general, which allow for global and more detailed analysis on TF binding grammar, as shown in the analysis of distances and orientations between individual TFs. Using the footprinting data, we found that TFs often co-occur with similar motifs at the same loci, a characteristic which has already been described to be important in both promoters and developmental enhancers [67]. These results hint towards a certain level of redundancy in TF binding, with similar transcription factors such as FOXA1 and FOXA2 substituting each other, as described previously [68]. In summary, all our findings illustrated that the selection of the input data plays a crucial role in the identification of resulting associations. Software design should ideally permit integration of data from various sources including newly emerging assays. For example, a recent method has improved on existing footprinting methods by applying methylation and bisulfite sequencing to interrogate single molecule footprints in single cells [69], which might therefore help to reduce the noise of footprinting in bulk samples. TF-COMB is ready to use such data, and with the advent of more technologies and available collections of TF binding positions, TF-COMB will help to further improve the investigation of co-occurrences and TF binding grammar.

By utilizing a pure motif analysis, we gained high resolution TFBS data, and could thereby isolate TF pairs which exhibit both preferred orientation and distance to each other. This, together with the observation of binding site periodicity, suggests the existence of a “Goldilocks distance” for many TF pairs, which we define by a set of locational parameters that probably optimizes complex building, binding duration or binding itself. The fact that these locations have significantly higher overlap with both regulatory features and “real” ChIP-seq derived binding sites, strongly supports this hypothesis. Of note, as only a minority of TFs are proven by wet lab based methods to physically interact, the preferred distance of TF pairs might also represent the right distance for comfortably fitting two proteins on the DNA without being sterically hindered by each other. In other cases, such as seen for the collective model, binding on the correct side of the DNA might also be essential in binding of co-factors. However, the majority of TFs do not show any particular binding grammar. This observation is not necessarily a rejection of their co-occurring status, but rather a sign of flexible TF binding. In fact, it has been shown that transient binding of multiple TFs is a mechanism to compete with nucleosome binding and keep DNA accessible - a mechanism known as ‘assisted loading’ [70]. In such cases, the exact location of TFs might be disregarded, as also observed in the billboard enhancer model.

Many areas of TF co-occurrence are still open for investigation, including the correlation of the size of open chromatin in relation to the number of TFs binding, and how TFs became hub proteins throughout evolution. For this study, we have focused on the co-occurrence of TFs within the same regulatory region, but our results have also identified a number of TFs, including CTCF and ZNF143, which are highly co-occurring at chromatin loop anchors, and are involved in connecting distant regulatory elements. There is thus an additional layer of 3D co-occurrences built between regulatory regions, as well as within looped enhancers, which we currently cannot track with our software. However, we were able to provide evidence for these structures in our co-occurrence analysis of histone modifications and TFs, where we found a number of TFs restricted to enhancer binding, avoiding cis regulatory regions. This suggests a model of e.g. differentiation, in which cell type specific TFs primarily control gene expression from regulatory enhancer elements *in trans*. This increases regulatory complexity, while simultaneously preventing spurious activation of target genes by TFs such as pioneers. Indeed, such regulation of enhancer activity by formation of topologically associated domains (TADs) is well-known for the HoxD cluster of genes, which are important for limb development [71]. Thus, the influence of chromatin organization should not be disregarded when discussing co-occurrence of transcription factors.

In conclusion, we have used TF-COMB to investigate a variety of aspects of TF binding grammar. Understanding the effect of TF co-occurrence is important for uncovering the direct targets of TFs, as multiple TFs create complicated AND/OR/XOR logic, as known from studies on systems biology. In particular, TFs such as pioneer factors can act as primers to subsequent binding of other TFs. It is therefore of great interest to discover potential sets of co-occurring TFs for individual cell lineages, and we believe that TF-COMB represents a valuable resource to identify, study and understand such TF co-occurrences in the context of gene regulation.

## Data availability

The TF-COMB tool is freely available under MIT License at Github: https://github.com/loosolab/TF-COMB. Full documentation and examples are available through ReadTheDocs at: https://tf-comb.readthedocs.io.

## Author contributions statement

M.B. and M.L conceived the study. M.B. designed TF-COMB. M.B., V.H. and H.S. implemented the framework. M.B, C.K. and V.H. preprocessed data. M.B., V.H. and H.S. performed the analysis. M.B., V.H., H.S., C.K. and M.L. wrote the manuscript. M.L. supervised the project.

## Figure attribution statement

Plots were produced using TF-COMB framework functionalities, and using matplotlib and seaborn packages in Python. The graphical abstract as well as explanatory (sub-)figures were created using BioRender.com as stated in the individual figure descriptions. Module icons as included in Figures 2-6 were taken from Figure 1.

## Supplementary data

**Supplementary Table 1:** An overview of ENCODE datasets used in this study. **Supplementary Table 2:** Overview on TF co-occurrence tools and evaluation on comparability.

**Supplementary Table 3:** Feature comparison of MCOT, SPAMO and TF-COMB. **Supplementary Table 4:** Literature search for top 10 exclusively found pairs for MCOT, SPAMO and TF-COMB.

**Supplementary Table 5:** Results of co-occurrence of ChIP-seq peaks across cell lines.

