## Supplementary information for "TF-COMB - discovering grammar of transcription factor binding sites"

### Supplementary Figures

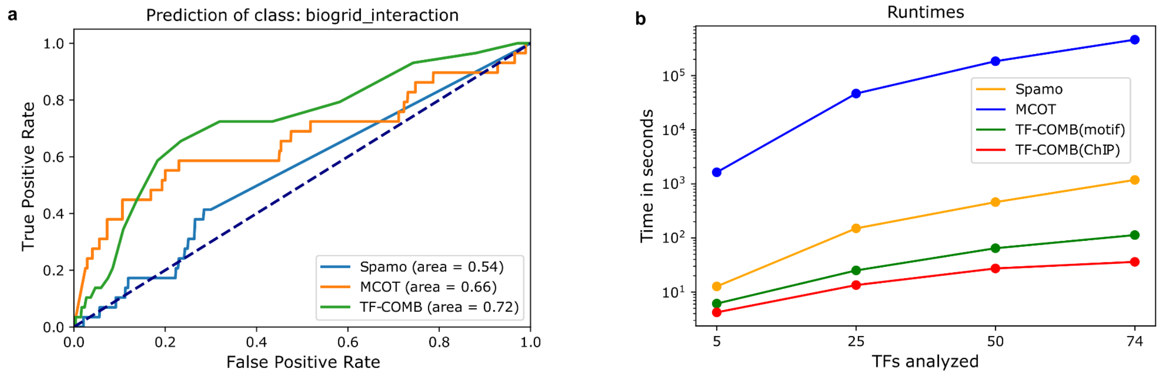

#### Supplementary Figure S1: Validation of TF-COMB and other tools

a) ROC curve of predictive ability of the assessed tools. Dashed line represents auROC=0.5.
b) Benchmark of runtime for assessed tools with increasing number of TFs analyzed.

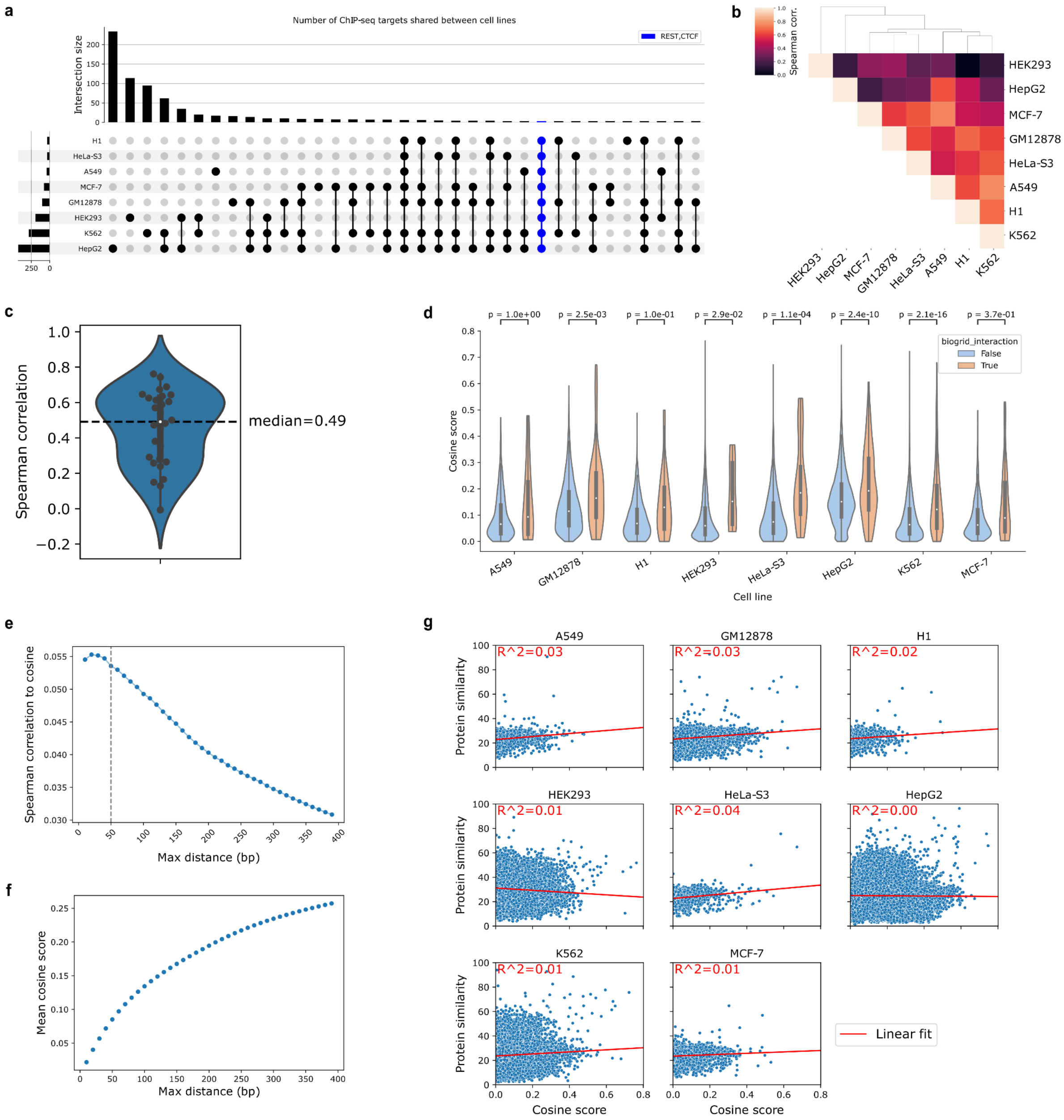

#### Supplementary Figure S2: Co-occurrence of ChIP-seq peaks across cell lines

a) Intersection count of available TFs ChIP-seq experiments for each cell line combination. Amount of TFs present in all cell lines highlighted in blue. Number of distinct TFs per cell line are on the left.
b) Correlation of TF-pair cosine values between cell lines.
c) Distribution of correlation values shown in b).
d) Distribution of cosine association scores for TF-pairs with and without PPIs across cell lines.
e) Correlation of cosine scores to PPIs with increasing allowed distance between TFs. Max distance of 50bp is marked with a dashed line.
f) Mean cosine association score with increasing allowed distance between TFs.
g) Cosine association compared to protein similarity of TF-pairs across cell lines. A linear fit is added in red and the R-squared is added at the left corner for each cell line.

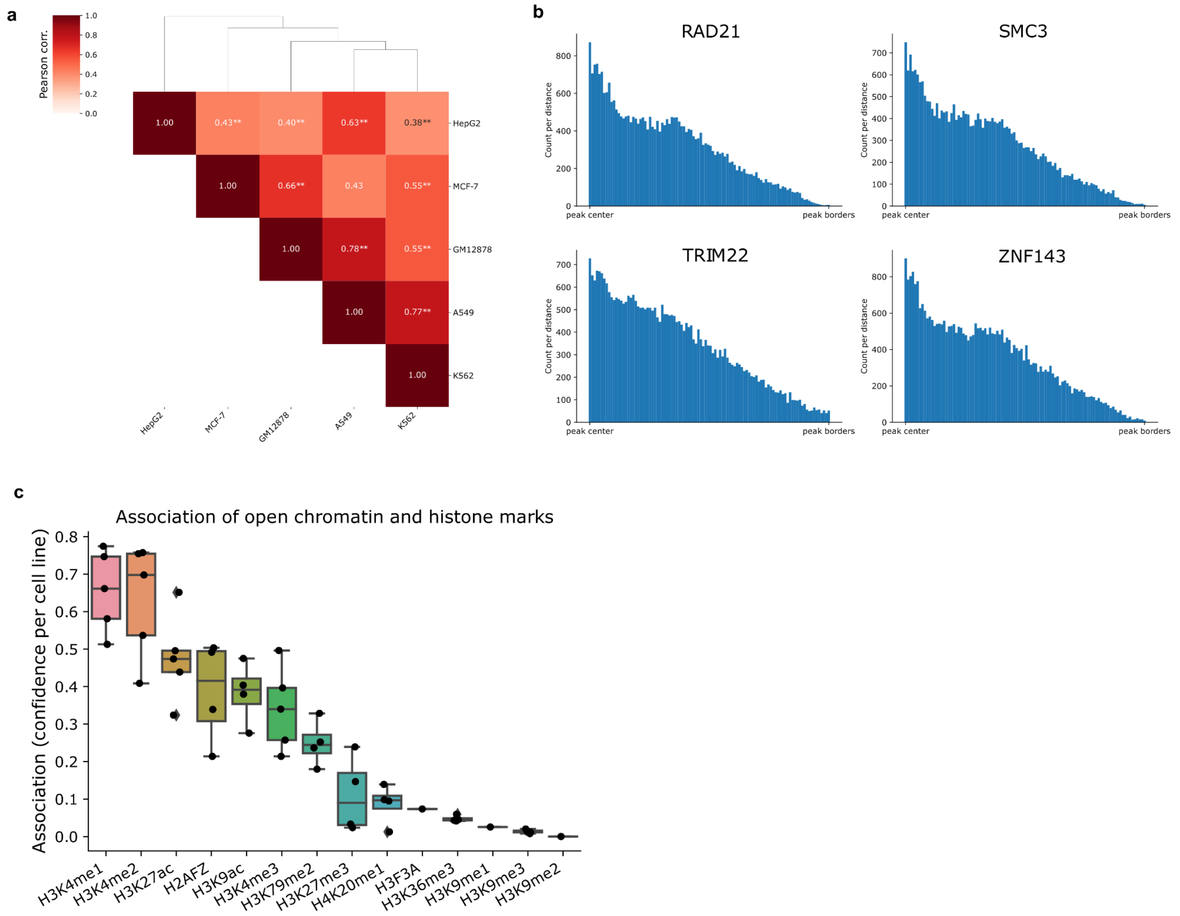

#### Supplementary Figure S3: Integration of epigenetic marks for co-occurrence

a) Correlation of TF binding distances(ChIP-seq) in relation to open chromatin (ATAC-seq) between cell lines. Significance of the correlation is marked with stars, where a p-value of less than 0.01 and 0.001 are marked with "*" and "**", respectively.

b) Examples of relative TF binding locations within peaks. The x-axis represents the binding location from center (left) to border (right) of open chromatin peaks.

c) Association of open chromatin to locations of histone modifications and variants. Each box shows the association for all cell lines to a specific histone modification.

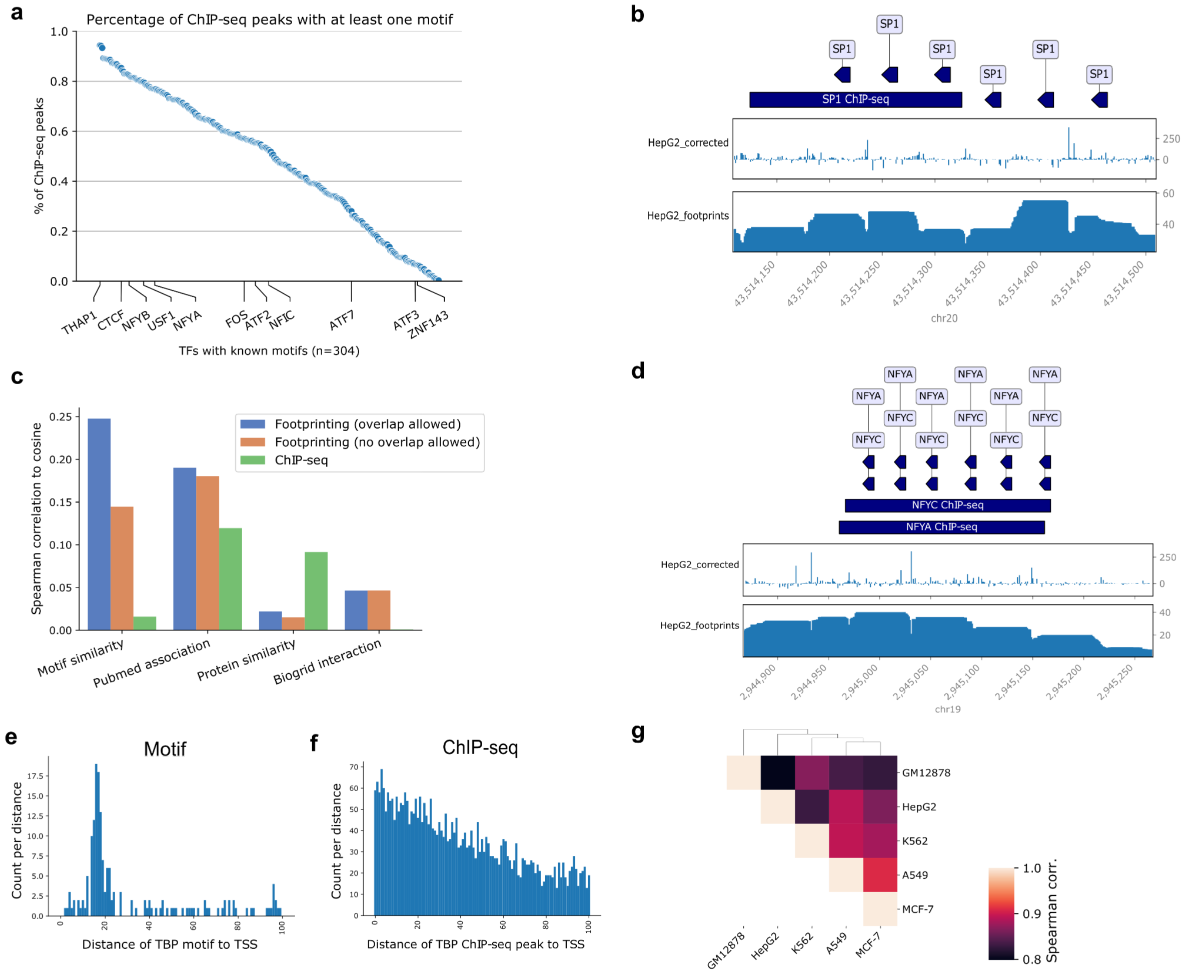

#### Supplementary Figure S4: Additional aspects of co-occurrence of footprinting

a) Fraction of ChIP-seq peaks containing sites of associated TF motifs.
b) Illustration of ChIP-seq peaks (upper blue boxes) in comparison to Tn5 signal (middle track) and TOBIAS derived footprinting score (lower track) for SP1. Individual TF motif locations are shown as triangles (upper track).
c) Correlation of cosine score with motif similarity, literature association score, protein similarity and protein interaction (BioGrid) score respectively per TF pair. Shown for ChIP-seq and footprinting with overlap allowed and excluded respectively.
d) Same plot as described in b) for NFYA/NFYC.
e-f) Distribution and gain of resolution shown for TBP binding site distances to TSS sites based on footprinting data analysis e) and ChIP-seq analysis f).
g) Correlation of TF pair cosine scores between cell lines from footprinting derived data.

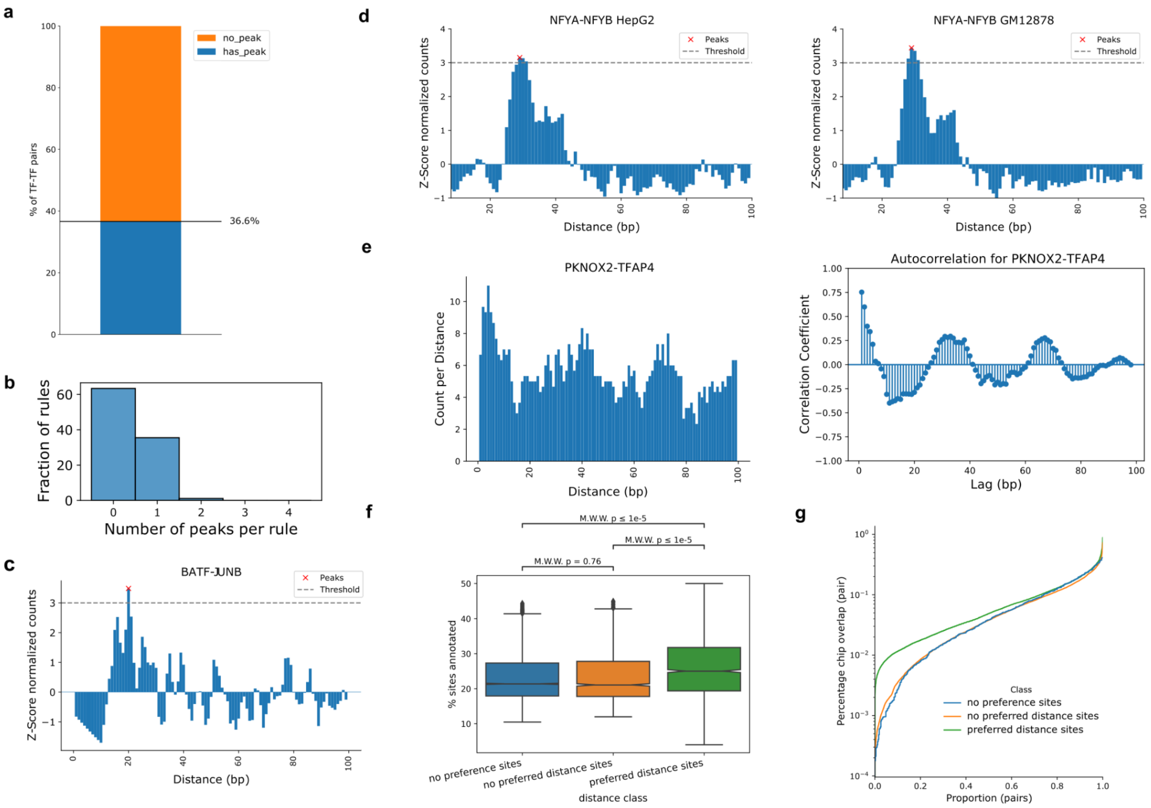

#### Supplementary Figure S5: Additional aspects of binding grammar

a) Percent of TF-pairs that exhibit at least one preferred binding distance (peak) as displayed in c-d).
b) Distribution of TF-pairs (rules) on their number of predicted preferred binding distances.
c) Z-score normalized TF-pair binding counts sorted by distance. Peaks above threshold are considered preferred binding distance.
d) Difference in binding distance distribution for NFYA-NFYB in HepG2 (left) and GM12878 (right).
e) Periodic binding distance preference for PKNOX2-TFAP4. Left plot shows the distribution of binding distances for all co-occurring sites. Right plot shows the calculated autocorrelation, i.e. lag of binding distances, for the pair.
f) Percentage of sites annotated to genes (using UROPA) per distance class.
g) Proportion of ChIP-seq overlap per distance class.

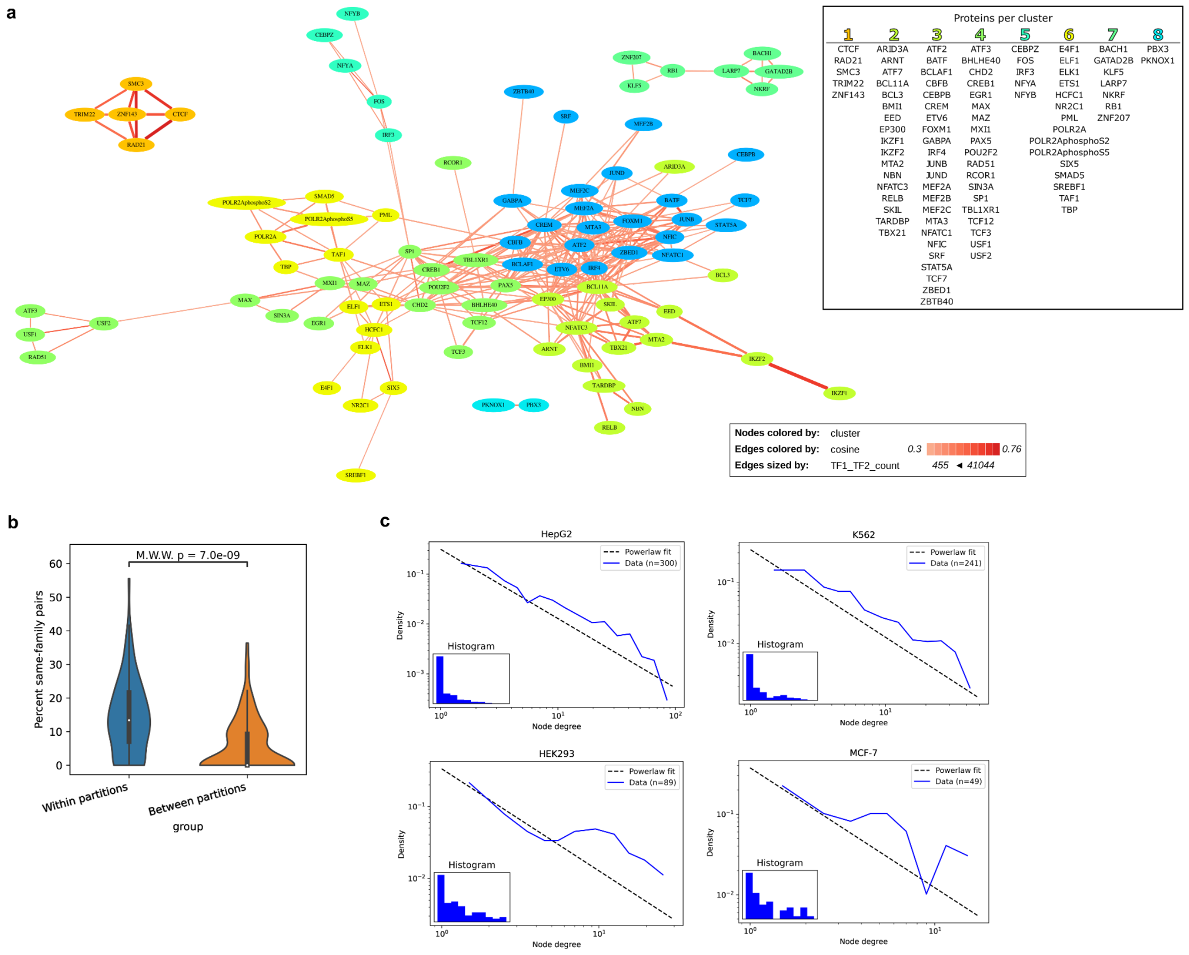

#### Supplementary Figure S6: Network analysis of co-occurrence

a) The GM12878 co-occurrence network. Nodes are colored on the basis of Louvain community clustering. On the right a list of TFs per cluster is shown.

b) Distribution of same-family pairs randomly picked from within or between partitions (clusters of a)). Percent same-family is estimated per subset for 20 randomly selected pairs across n=1000 iterations.

c) Node degree (count) for all TFs of different cell line networks. Node degrees follow power-law distribution.

#### Supplementary Table 1:
##### An overview of ENCODE datasets used in this study

| File accession | File format | Experiment target | Biosample term name | Biological replicate(s) | Assay |
|---|---|---|---|---|---|
| ENCFF607DTB | bam | | A549 | 1 | ATAC-seq |
| ENCFF701BDT | bam | | A549 | 2 | ATAC-seq |
| ENCFF616DYV | bam | | A549 | 3 | ATAC-seq |
| ENCFF981FXV | bam | | GM12878 | 1 | ATAC-seq |
| ENCFF962FMH | bam | | GM12878 | 2 | ATAC-seq |
| ENCFF440GRZ | bam | | GM12878 | 3 | ATAC-seq |
| ENCFF990VCP | bam | | HepG2 | 1 | ATAC-seq |
| ENCFF624SON | bam | | HepG2 | 2 | ATAC-seq |
| ENCFF926KFU | bam | | HepG2 | 3 | ATAC-seq |
| ENCFF534DCE | bam | | K562 | 1 | ATAC-seq |
| ENCFF128WZG | bam | | K562 | 2 | ATAC-seq |
| ENCFF077FBI | bam | | K562 | 3 | ATAC-seq |
| ENCFF607OSL | bam | | MCF-7 | 1 | ATAC-seq |
| ENCFF772EFK | bam | | MCF-7 | 2 | ATAC-seq |
| ENCFF678GBM | bed narrowPeak | | K562 | 1, 2, 3 | ATAC-seq |
| ENCFF915FZC | bed narrowPeak | | HepG2 | 1, 2, 3 | ATAC-seq |
| ENCFF871TST | bed narrowPeak | | GM12878 | 1, 2, 3 | ATAC-seq |
| ENCFF964AUZ | bed narrowPeak | | A549 | 1, 2, 3 | ATAC-seq |
| ENCFF291WTA | bed narrowPeak | | MCF-7 | 1, 2 | ATAC-seq |
| ENCFF918FET | bed narrowPeak | H3K27ac | A549 | 1, 2, 3 | Histone ChIP-seq |
| ENCFF594YDK | bed narrowPeak | H3K4me1 | A549 | 1, 2, 3 | Histone ChIP-seq |
| ENCFF794FQH | bed narrowPeak | H3K4me2 | A549 | 1, 2, 3 | Histone ChIP-seq |
| ENCFF535EYL | bed narrowPeak | H3K4me3 | A549 | 1, 2, 3 | Histone ChIP-seq |
| ENCFF164FDB | bed narrowPeak | H3K9me3 | A549 | 1, 2, 3 | Histone ChIP-seq |
| ENCFF377OJG | bed narrowPeak | H2AFZ | GM12878 | 1, 2 | Histone ChIP-seq |
| ENCFF023LTU | bed narrowPeak | H3K27ac | GM12878 | 1, 2 | Histone ChIP-seq |
| ENCFF695ETB | bed narrowPeak | H3K27me3 | GM12878 | 1, 2 | Histone ChIP-seq |
| ENCFF432EMI | bed narrowPeak | H3K36me3 | GM12878 | 1, 2 | Histone ChIP-seq |
| ENCFF321BVG | bed narrowPeak | H3K4me1 | GM12878 | 1, 2 | Histone ChIP-seq |
| ENCFF283LNH | bed narrowPeak | H3K4me2 | GM12878 | 1, 2 | Histone ChIP-seq |
| ENCFF587DVA | bed narrowPeak | H3K4me3 | GM12878 | 1, 2 | Histone ChIP-seq |
| ENCFF131ZGZ | bed narrowPeak | H3K79me2 | GM12878 | 1, 2 | Histone ChIP-seq |
| ENCFF069KAG | bed narrowPeak | H3K9ac | GM12878 | 1, 2 | Histone ChIP-seq |
| ENCFF725UFY | bed narrowPeak | H3K9me3 | GM12878 | 1, 2, 3 | Histone ChIP-seq |
| ENCFF469QYW | bed narrowPeak | H4K20me1 | GM12878 | 1, 2 | Histone ChIP-seq |
| ENCFF745GKP | bed narrowPeak | H2AFZ | H1 | 1, 2 | Histone ChIP-seq |
| ENCFF320JIP | bed narrowPeak | H2AK5ac | H1 | 1, 2 | Histone ChIP-seq |
| ENCFF808BIC | bed narrowPeak | H2BK120ac | H1 | 1, 2 | Histone ChIP-seq |
| ENCFF298VRY | bed narrowPeak | H2BK120ac | H1 | 1, 2 | Histone ChIP-seq |
| ENCFF237XZS | bed narrowPeak | H2BK12ac | H1 | 1, 2 | Histone ChIP-seq |
| ENCFF855ILJ | bed narrowPeak | H2BK15ac | H1 | 1, 2 | Histone ChIP-seq |
| ENCFF215EPH | bed narrowPeak | H2BK20ac | H1 | 1, 2 | Histone ChIP-seq |
| ENCFF929FTR | bed narrowPeak | H2BK5ac | H1 | 1, 2 | Histone ChIP-seq |
| ENCFF870OAL | bed narrowPeak | H3K14ac | H1 | 1, 2 | Histone ChIP-seq |
| ENCFF192GAX | bed narrowPeak | H3K18ac | H1 | 1, 2 | Histone ChIP-seq |
| ENCFF350XJF | bed narrowPeak | H3K23ac | H1 | 1, 2 | Histone ChIP-seq |

| | | | | | |
|---|---|---|---|---|---|
| ENCFF567WXW | bed narrowPeak | H3K23me2 | H1 | 1, 2 | Histone ChIP-seq |
| ENCFF689CJG | bed narrowPeak | H3K27ac | H1 | 1, 2 | Histone ChIP-seq |
| ENCFF254ACI | bed narrowPeak | H3K27me3 | H1 | 1, 2 | Histone ChIP-seq |
| ENCFF051UOL | bed narrowPeak | H3K36me3 | H1 | 1, 2 | Histone ChIP-seq |
| ENCFF711LQB | bed narrowPeak | H3K4ac | H1 | 1, 2 | Histone ChIP-seq |
| ENCFF327EZJ | bed narrowPeak | H3K4ac | H1 | 1, 2 | Histone ChIP-seq |
| ENCFF613QAB | bed narrowPeak | H3K4me1 | H1 | 1, 2 | Histone ChIP-seq |
| ENCFF583ABZ | bed narrowPeak | H3K4me2 | H1 | 1, 2 | Histone ChIP-seq |
| ENCFF041HYH | bed narrowPeak | H3K4me3 | H1 | 1, 2 | Histone ChIP-seq |
| ENCFF053HKF | bed narrowPeak | H3K56ac | H1 | 1, 2 | Histone ChIP-seq |
| ENCFF472HMD | bed narrowPeak | H3K56ac | H1 | 1, 2 | Histone ChIP-seq |
| ENCFF088PTH | bed narrowPeak | H3K79me1 | H1 | 1, 3 | Histone ChIP-seq |
| ENCFF620GIW | bed narrowPeak | H3K79me2 | H1 | 1, 2 | Histone ChIP-seq |
| ENCFF679LHF | bed narrowPeak | H3K9ac | H1 | 1, 2 | Histone ChIP-seq |
| ENCFF678VNN | bed narrowPeak | H3K9me3 | H1 | 1, 3 | Histone ChIP-seq |
| ENCFF439CWL | bed narrowPeak | H4K20me1 | H1 | 1, 2 | Histone ChIP-seq |
| ENCFF295UVV | bed narrowPeak | H4K20me1 | H1 | 1, 2 | Histone ChIP-seq |
| ENCFF097DDN | bed narrowPeak | H4K5ac | H1 | 1, 2 | Histone ChIP-seq |
| ENCFF760EFQ | bed narrowPeak | H4K8ac | H1 | 1, 2 | Histone ChIP-seq |
| ENCFF910SNO | bed narrowPeak | H4K91ac | H1 | 1, 2 | Histone ChIP-seq |
| ENCFF788KUH | bed narrowPeak | H4K91ac | H1 | 1, 2 | Histone ChIP-seq |
| ENCFF225FPN | bed narrowPeak | H3K27ac | HEK293 | 1, 2 | Histone ChIP-seq |
| ENCFF257ZFX | bed narrowPeak | H3K36me3 | HEK293 | 1, 2 | Histone ChIP-seq |
| ENCFF351MDO | bed narrowPeak | H3K4me1 | HEK293 | 1, 2 | Histone ChIP-seq |
| ENCFF617NUV | bed narrowPeak | H3K4me3 | HEK293 | 1, 2 | Histone ChIP-seq |
| ENCFF899MET | bed narrowPeak | H3K9me3 | HEK293 | 1, 2 | Histone ChIP-seq |
| ENCFF128TOM | bed narrowPeak | H2AFZ | HeLa-S3 | 1, 2 | Histone ChIP-seq |
| ENCFF831XSS | bed narrowPeak | H3K27ac | HeLa-S3 | 1, 2 | Histone ChIP-seq |
| ENCFF428MSN | bed narrowPeak | H3K27me3 | HeLa-S3 | 1, 2 | Histone ChIP-seq |
| ENCFF348FNV | bed narrowPeak | H3K36me3 | HeLa-S3 | 1, 2 | Histone ChIP-seq |
| ENCFF572NWJ | bed narrowPeak | H3K4me1 | HeLa-S3 | 1, 2 | Histone ChIP-seq |
| ENCFF429OQI | bed narrowPeak | H3K4me2 | HeLa-S3 | 1, 2 | Histone ChIP-seq |
| ENCFF932ZNX | bed narrowPeak | H3K4me3 | HeLa-S3 | 1, 2 | Histone ChIP-seq |
| ENCFF238XWI | bed narrowPeak | H3K79me2 | HeLa-S3 | 1, 2 | Histone ChIP-seq |
| ENCFF021PYM | bed narrowPeak | H3K9ac | HeLa-S3 | 1, 2 | Histone ChIP-seq |
| ENCFF872YCK | bed narrowPeak | H3K9me3 | HeLa-S3 | 1, 2 | Histone ChIP-seq |
| ENCFF406DAM | bed narrowPeak | H4K20me1 | HeLa-S3 | 1, 2 | Histone ChIP-seq |
| ENCFF303VCJ | bed narrowPeak | H2AFZ | HepG2 | 1 | Histone ChIP-seq |
| ENCFF749VEQ | bed narrowPeak | H3K27ac | HepG2 | 1, 2 | Histone ChIP-seq |
| ENCFF053AZC | bed narrowPeak | H3K27me3 | HepG2 | 1, 2 | Histone ChIP-seq |
| ENCFF296VZN | bed narrowPeak | H3K36me3 | HepG2 | 1, 2 | Histone ChIP-seq |
| ENCFF413EGR | bed narrowPeak | H3K4me1 | HepG2 | 1, 2 | Histone ChIP-seq |
| ENCFF946FQI | bed narrowPeak | H3K4me2 | HepG2 | 1, 2 | Histone ChIP-seq |
| ENCFF982DUT | bed narrowPeak | H3K4me3 | HepG2 | 1, 2 | Histone ChIP-seq |
| ENCFF500KVU | bed narrowPeak | H3K79me2 | HepG2 | 1 | Histone ChIP-seq |
| ENCFF044QMC | bed narrowPeak | H3K9ac | HepG2 | 1, 2 | Histone ChIP-seq |
| ENCFF372HCL | bed narrowPeak | H3K9me3 | HepG2 | 1, 2 | Histone ChIP-seq |
| ENCFF811ZMQ | bed narrowPeak | H4K20me1 | HepG2 | 1, 2 | Histone ChIP-seq |
| ENCFF213OTI | bed narrowPeak | H2AFZ | K562 | 1, 2 | Histone ChIP-seq |
| ENCFF864OSZ | bed narrowPeak | H3K27ac | K562 | 1, 2 | Histone ChIP-seq |
| ENCFF801AHF | bed narrowPeak | H3K27me3 | K562 | 1, 2 | Histone ChIP-seq |
| ENCFF561OUZ | bed narrowPeak | H3K36me3 | K562 | 1, 2 | Histone ChIP-seq |

| | | | | | |
|---|---|---|---|---|---|
| ENCFF759NWD | bed narrowPeak | H3K4me1 | K562 | 1, 2 | Histone ChIP-seq |
| ENCFF749KLQ | bed narrowPeak | H3K4me2 | K562 | 1, 2 | Histone ChIP-seq |
| ENCFF706WUF | bed narrowPeak | H3K4me3 | K562 | 1, 2 | Histone ChIP-seq |
| ENCFF945LQZ | bed narrowPeak | H3K79me2 | K562 | 1, 2 | Histone ChIP-seq |
| ENCFF148UQI | bed narrowPeak | H3K9ac | K562 | 1, 2 | Histone ChIP-seq |
| ENCFF462AVD | bed narrowPeak | H3K9me1 | K562 | 1 | Histone ChIP-seq |
| ENCFF963GZJ | bed narrowPeak | H3K9me3 | K562 | 1, 2 | Histone ChIP-seq |
| ENCFF909RKY | bed narrowPeak | H4K20me1 | K562 | 1, 2 | Histone ChIP-seq |
| ENCFF378IVZ | bed narrowPeak | H2AFZ | MCF-7 | 1, 2 | Histone ChIP-seq |
| ENCFF595UXF | bed narrowPeak | H3F3A | MCF-7 | 1, 2 | Histone ChIP-seq |
| ENCFF491LQY | bed narrowPeak | H3K27ac | MCF-7 | 1, 2 | Histone ChIP-seq |
| ENCFF669NUD | bed narrowPeak | H3K27me3 | MCF-7 | 1, 2 | Histone ChIP-seq |
| ENCFF739SBL | bed narrowPeak | H3K36me3 | MCF-7 | 1, 2 | Histone ChIP-seq |
| ENCFF991HJA | bed narrowPeak | H3K4me1 | MCF-7 | 1, 2 | Histone ChIP-seq |
| ENCFF188VRU | bed narrowPeak | H3K4me2 | MCF-7 | 1, 2 | Histone ChIP-seq |
| ENCFF268RXB | bed narrowPeak | H3K4me3 | MCF-7 | 1, 2 | Histone ChIP-seq |
| ENCFF660MSN | bed narrowPeak | H3K79me2 | MCF-7 | 1, 2 | Histone ChIP-seq |
| ENCFF348DEB | bed narrowPeak | H3K9ac | MCF-7 | 1, 2 | Histone ChIP-seq |
| ENCFF538BHD | bed narrowPeak | H3K9me2 | MCF-7 | 1, 2 | Histone ChIP-seq |
| ENCFF501UHK | bed narrowPeak | H3K9me3 | MCF-7 | 1, 2 | Histone ChIP-seq |
| ENCFF714DEQ | bed narrowPeak | H4K20me1 | MCF-7 | 1, 2 | Histone ChIP-seq |
| ENCFF863QWG | | | HepG2 | 1 | RNA-seq |
| ENCFF721BRA | | | MCF-7 | 1 | RNA-seq |
| ENCFF520POQ | bed narrowPeak | ATOH8 | A549 | 1, 2 | TF ChIP-seq |
| ENCFF008SZM | bed narrowPeak | BCL3 | A549 | 1, 2, 3 | TF ChIP-seq |
| ENCFF808XJN | bed narrowPeak | BHLHE40 | A549 | 1, 2 | TF ChIP-seq |
| ENCFF208AXT | bed narrowPeak | CBX2 | A549 | 1, 2 | TF ChIP-seq |
| ENCFF330OCU | bed narrowPeak | CBX8 | A549 | 1, 2 | TF ChIP-seq |
| ENCFF064WDQ | bed narrowPeak | CEBPB | A549 | 1, 2 | TF ChIP-seq |
| ENCFF757QNZ | bed narrowPeak | CHD2 | A549 | 1, 3 | TF ChIP-seq |
| ENCFF766YPH | bed narrowPeak | CHD4 | A549 | 1, 2 | TF ChIP-seq |
| ENCFF624ZSR | bed narrowPeak | CTCF | A549 | 1, 2, 3 | TF ChIP-seq |
| ENCFF786NAO | bed narrowPeak | E2F6 | A549 | 1, 2 | TF ChIP-seq |
| ENCFF507RLU | bed narrowPeak | EHF | A549 | 1, 2 | TF ChIP-seq |
| ENCFF199OOU | bed narrowPeak | EHMT2 | A549 | 1, 2 | TF ChIP-seq |
| ENCFF486IUH | bed narrowPeak | ELK1 | A549 | 1, 3 | TF ChIP-seq |
| ENCFF143OQP | bed narrowPeak | EP300 | A549 | 1, 2, 3 | TF ChIP-seq |
| ENCFF039TWK | bed narrowPeak | ESRRA | A549 | 1, 2 | TF ChIP-seq |
| ENCFF133ZSB | bed narrowPeak | FOSB | A549 | 1, 2 | TF ChIP-seq |
| ENCFF232TVP | bed narrowPeak | FOSL2 | A549 | 1, 2 | TF ChIP-seq |
| ENCFF109ADF | bed narrowPeak | FOXF2 | A549 | 1, 2 | TF ChIP-seq |
| ENCFF016TWL | bed narrowPeak | FOXS1 | A549 | 1, 2 | TF ChIP-seq |
| ENCFF517TKD | bed narrowPeak | GATA3 | A549 | 1, 2 | TF ChIP-seq |
| ENCFF814DAF | bed narrowPeak | HDAC2 | A549 | 1, 2 | TF ChIP-seq |
| ENCFF209ZVL | bed narrowPeak | HES2 | A549 | 1, 2 | TF ChIP-seq |
| ENCFF708HLM | bed narrowPeak | HMGXB4 | A549 | 1, 2 | TF ChIP-seq |
| ENCFF670UIE | bed narrowPeak | HOXA7 | A549 | 1, 2 | TF ChIP-seq |
| ENCFF641ZFM | bed narrowPeak | HOXB13 | A549 | 1, 2 | TF ChIP-seq |
| ENCFF634TZG | bed narrowPeak | HOXB5 | A549 | 1, 2 | TF ChIP-seq |
| ENCFF329CUQ | bed narrowPeak | JUN | A549 | 1, 2, 3 | TF ChIP-seq |
| ENCFF988QUL | bed narrowPeak | JUNB | A549 | 2, 3 | TF ChIP-seq |
| ENCFF316CBQ | bed narrowPeak | KDM1A | A549 | 1, 2 | TF ChIP-seq |

| | | | | |
|---|---|---|---|---|
| ENCFF702XIF | bed narrowPeak | KDM5A | A549 | 1, 2 TF ChIP-seq |
| ENCFF813WJW | bed narrowPeak | MAFK | A549 | 2, 3 TF ChIP-seq |
| ENCFF713RHL | bed narrowPeak | MAX | A549 | 1, 2 TF ChIP-seq |
| ENCFF661NNJ | bed narrowPeak | MAZ | A549 | 2, 3 TF ChIP-seq |
| ENCFF598BZD | bed narrowPeak | MYC | A549 | 1, 2 TF ChIP-seq |
| ENCFF418TUX | bed narrowPeak | NFE2L2 | A549 | 1, 2 TF ChIP-seq |
| ENCFF404ASR | bed narrowPeak | NR2E3 | A549 | 1, 2 TF ChIP-seq |
| ENCFF562UOF | bed narrowPeak | NR3C1 | A549 | 1, 2, 3 TF ChIP-seq |
| ENCFF507JZI | bed narrowPeak | NR5A2 | A549 | 1, 2 TF ChIP-seq |
| ENCFF992LDJ | bed narrowPeak | PBX1 | A549 | 1, 2 TF ChIP-seq |
| ENCFF656HFI | bed narrowPeak | PBX3 | A549 | 1, 2 TF ChIP-seq |
| ENCFF307JCM | bed narrowPeak | PHF8 | A549 | 1, 2 TF ChIP-seq |
| ENCFF681MRC | bed narrowPeak | POLR2A | A549 | 1, 2 TF ChIP-seq |
| ENCFF156MIR | bed narrowPeak | POLR2AphosphoS2 | A549 | 1, 2 TF ChIP-seq |
| ENCFF719BHI | bed narrowPeak | PRDM1 | A549 | 1, 2 TF ChIP-seq |
| ENCFF695MMQ | bed narrowPeak | RAD21 | A549 | 1, 2 TF ChIP-seq |
| ENCFF871ZOH | bed narrowPeak | RARB | A549 | 1, 2 TF ChIP-seq |
| ENCFF993WZP | bed narrowPeak | RCOR1 | A549 | 1, 2 TF ChIP-seq |
| ENCFF814JWH | bed narrowPeak | REST | A549 | 1, 2 TF ChIP-seq |
| ENCFF179WDI | bed narrowPeak | RFX5 | A549 | 2, 3 TF ChIP-seq |
| ENCFF110EOX | bed narrowPeak | RNF2 | A549 | 1, 2 TF ChIP-seq |
| ENCFF567BJI | bed narrowPeak | SIN3A | A549 | 2, 3 TF ChIP-seq |
| ENCFF146MNV | bed narrowPeak | SMC3 | A549 | 1, 2, 3 TF ChIP-seq |
| ENCFF624DDK | bed narrowPeak | SREBF1 | A549 | 2, 3 TF ChIP-seq |
| ENCFF483YCC | bed narrowPeak | SREBF2 | A549 | 2, 3 TF ChIP-seq |
| ENCFF766VUQ | bed narrowPeak | TEAD4 | A549 | 1, 2 TF ChIP-seq |
| ENCFF679WUD | bed narrowPeak | TFCP2L1 | A549 | 1, 2 TF ChIP-seq |
| ENCFF699UTZ | bed narrowPeak | TP53 | A549 | 1, 2 TF ChIP-seq |
| ENCFF294QSH | bed narrowPeak | TP63 | A549 | 1, 2 TF ChIP-seq |
| ENCFF593EOW | bed narrowPeak | USF2 | A549 | 2, 3 TF ChIP-seq |
| ENCFF390AZA | bed narrowPeak | ZC3H11A | A549 | 2, 3 TF ChIP-seq |
| ENCFF137JHO | bed narrowPeak | ZFP36 | A549 | 2, 3 TF ChIP-seq |
| ENCFF857LEC | bed narrowPeak | ZNF302 | A549 | 1, 2 TF ChIP-seq |
| ENCFF439SWX | bed narrowPeak | ZNF624 | A549 | 1, 2 TF ChIP-seq |
| ENCFF003VDB | bed narrowPeak | ARID3A | GM12878 | 5, 6 TF ChIP-seq |
| ENCFF758RQJ | bed narrowPeak | ARNT | GM12878 | 2, 3 TF ChIP-seq |
| ENCFF096XRG | bed narrowPeak | ASH2L | GM12878 | 1, 2 TF ChIP-seq |
| ENCFF806KKM | bed narrowPeak | ATF2 | GM12878 | 1, 2 TF ChIP-seq |
| ENCFF882AEU | bed narrowPeak | ATF3 | GM12878 | 1, 2 TF ChIP-seq |
| ENCFF495PWL | bed narrowPeak | ATF7 | GM12878 | 1, 2 TF ChIP-seq |
| ENCFF725YZH | bed narrowPeak | BACH1 | GM12878 | 3, 4 TF ChIP-seq |
| ENCFF832YIE | bed narrowPeak | BATF | GM12878 | 1, 2 TF ChIP-seq |
| ENCFF824QXX | bed narrowPeak | BCL11A | GM12878 | 1, 2 TF ChIP-seq |
| ENCFF452NTN | bed narrowPeak | BCL3 | GM12878 | 1, 2 TF ChIP-seq |
| ENCFF587BJK | bed narrowPeak | BCLAF1 | GM12878 | 1, 2 TF ChIP-seq |
| ENCFF633YUQ | bed narrowPeak | BHLHE40 | GM12878 | 1, 2 TF ChIP-seq |
| ENCFF592LPO | bed narrowPeak | BMI1 | GM12878 | 1, 2 TF ChIP-seq |
| ENCFF005JKU | bed narrowPeak | BRCA1 | GM12878 | 1, 2 TF ChIP-seq |
| ENCFF070SOX | bed narrowPeak | CBFB | GM12878 | 1, 2 TF ChIP-seq |
| ENCFF552QOA | bed narrowPeak | CBX3 | GM12878 | 1, 2 TF ChIP-seq |
| ENCFF864AYJ | bed narrowPeak | CBX5 | GM12878 | 1, 2 TF ChIP-seq |
| ENCFF955YFB | bed narrowPeak | CEBPB | GM12878 | 1, 2 TF ChIP-seq |

| | | | | | |
|---|---|---|---|---|---|
| ENCFF243GOG | bed narrowPeak | CEBPZ | GM12878 | 1, 3 | TF ChIP-seq |
| ENCFF863CTN | bed narrowPeak | CHD1 | GM12878 | 1, 2 | TF ChIP-seq |
| ENCFF351VXQ | bed narrowPeak | CHD2 | GM12878 | 1, 2 | TF ChIP-seq |
| ENCFF249SIN | bed narrowPeak | CHD4 | GM12878 | 1, 2 | TF ChIP-seq |
| ENCFF053UZX | bed narrowPeak | CREB1 | GM12878 | 1, 2 | TF ChIP-seq |
| ENCFF091YID | bed narrowPeak | CREM | GM12878 | 1, 2 | TF ChIP-seq |
| ENCFF827JRI | bed narrowPeak | CTCF | GM12878 | 1, 2, 3 | TF ChIP-seq |
| ENCFF567NFS | bed narrowPeak | CUX1 | GM12878 | 1, 2 | TF ChIP-seq |
| ENCFF003ZLG | bed narrowPeak | DPF2 | GM12878 | 1, 2 | TF ChIP-seq |
| ENCFF687SFB | bed narrowPeak | E2F4 | GM12878 | 1, 2 | TF ChIP-seq |
| ENCFF412GFI | bed narrowPeak | E2F8 | GM12878 | 1, 2 | TF ChIP-seq |
| ENCFF035GFS | bed narrowPeak | E4F1 | GM12878 | 1, 2 | TF ChIP-seq |
| ENCFF895MHN | bed narrowPeak | EBF1 | GM12878 | 1, 2 | TF ChIP-seq |
| ENCFF023ALY | bed narrowPeak | EED | GM12878 | 1, 2 | TF ChIP-seq |
| ENCFF637UJN | bed narrowPeak | EGR1 | GM12878 | 1, 2 | TF ChIP-seq |
| ENCFF146SYU | bed narrowPeak | ELF1 | GM12878 | 1, 2 | TF ChIP-seq |
| ENCFF432AQP | bed narrowPeak | ELK1 | GM12878 | 1, 2 | TF ChIP-seq |
| ENCFF718TWQ | bed narrowPeak | EP300 | GM12878 | 1, 2 | TF ChIP-seq |
| ENCFF722LJP | bed narrowPeak | ESRRA | GM12878 | 1, 2 | TF ChIP-seq |
| ENCFF980VOD | bed narrowPeak | ETS1 | GM12878 | 1, 2 | TF ChIP-seq |
| ENCFF745ANU | bed narrowPeak | ETV6 | GM12878 | 1, 2 | TF ChIP-seq |
| ENCFF615NYO | bed narrowPeak | EZH2 | GM12878 | 1, 2 | TF ChIP-seq |
| ENCFF571DGT | bed narrowPeak | FOS | GM12878 | 1, 2, 3 | TF ChIP-seq |
| ENCFF990MTR | bed narrowPeak | FOXK2 | GM12878 | 1, 2 | TF ChIP-seq |
| ENCFF549GKZ | bed narrowPeak | FOXM1 | GM12878 | 1, 2 | TF ChIP-seq |
| ENCFF093KLR | bed narrowPeak | GABPA | GM12878 | 1, 2 | TF ChIP-seq |
| ENCFF298AIX | bed narrowPeak | GATAD2B | GM12878 | 1, 2 | TF ChIP-seq |
| ENCFF722QBB | bed narrowPeak | HCFC1 | GM12878 | 1, 2 | TF ChIP-seq |
| ENCFF299UPZ | bed narrowPeak | HDAC2 | GM12878 | 1, 2 | TF ChIP-seq |
| ENCFF588SBL | bed narrowPeak | HDAC6 | GM12878 | 1 | TF ChIP-seq |
| ENCFF442WRJ | bed narrowPeak | HDGF | GM12878 | 1, 2 | TF ChIP-seq |
| ENCFF603BID | bed narrowPeak | HSF1 | GM12878 | 1, 2 | TF ChIP-seq |
| ENCFF819VMH | bed narrowPeak | IKZF1 | GM12878 | 1, 2 | TF ChIP-seq |
| ENCFF511HZB | bed narrowPeak | IKZF2 | GM12878 | 1, 2 | TF ChIP-seq |
| ENCFF604AZX | bed narrowPeak | IRF3 | GM12878 | 1, 2 | TF ChIP-seq |
| ENCFF113VGD | bed narrowPeak | IRF4 | GM12878 | 1, 2 | TF ChIP-seq |
| ENCFF843HDK | bed narrowPeak | IRF5 | GM12878 | 1, 2 | TF ChIP-seq |
| ENCFF478XNA | bed narrowPeak | JUNB | GM12878 | 1, 2 | TF ChIP-seq |
| ENCFF134BQO | bed narrowPeak | JUND | GM12878 | 1, 2 | TF ChIP-seq |
| ENCFF710ROZ | bed narrowPeak | KAT2A | GM12878 | 1, 2 | TF ChIP-seq |
| ENCFF996NUD | bed narrowPeak | KDM1A | GM12878 | 1, 2 | TF ChIP-seq |
| ENCFF417WPC | bed narrowPeak | KLF5 | GM12878 | 1, 2 | TF ChIP-seq |
| ENCFF305SLO | bed narrowPeak | LARP7 | GM12878 | 1, 2 | TF ChIP-seq |
| ENCFF436SJS | bed narrowPeak | MAFK | GM12878 | 1, 2 | TF ChIP-seq |
| ENCFF270NAL | bed narrowPeak | MAX | GM12878 | 1, 2 | TF ChIP-seq |
| ENCFF348STZ | bed narrowPeak | MAZ | GM12878 | 1, 2 | TF ChIP-seq |
| ENCFF826GQU | bed narrowPeak | MEF2A | GM12878 | 1, 2 | TF ChIP-seq |
| ENCFF623FAW | bed narrowPeak | MEF2B | GM12878 | 1, 2 | TF ChIP-seq |
| ENCFF830BRO | bed narrowPeak | MEF2C | GM12878 | 1, 2 | TF ChIP-seq |
| ENCFF652GOE | bed narrowPeak | MLLT1 | GM12878 | 1, 2 | TF ChIP-seq |
| ENCFF587POH | bed narrowPeak | MTA2 | GM12878 | 1, 2 | TF ChIP-seq |
| ENCFF661FMB | bed narrowPeak | MTA3 | GM12878 | 1, 2 | TF ChIP-seq |

| | | | | | |
|---|---|---|---|---|---|
| ENCFF199HGX | bed narrowPeak | MXI1 | GM12878 | 1, 2 | TF ChIP-seq |
| ENCFF402TSJ | bed narrowPeak | MYB | GM12878 | 1, 3 | TF ChIP-seq |
| ENCFF214XPD | bed narrowPeak | MYC | GM12878 | 1, 2 | TF ChIP-seq |
| ENCFF811VEN | bed narrowPeak | NBN | GM12878 | 1, 2 | TF ChIP-seq |
| ENCFF138ZBJ | bed narrowPeak | NFATC1 | GM12878 | 1, 2 | TF ChIP-seq |
| ENCFF002XEC | bed narrowPeak | NFATC3 | GM12878 | 1, 2 | TF ChIP-seq |
| ENCFF480WDX | bed narrowPeak | NFIC | GM12878 | 1, 2 | TF ChIP-seq |
| ENCFF860IXB | bed narrowPeak | NFXL1 | GM12878 | 1, 2 | TF ChIP-seq |
| ENCFF278GJK | bed narrowPeak | NFYA | GM12878 | 1, 2 | TF ChIP-seq |
| ENCFF156MUM | bed narrowPeak | NFYB | GM12878 | 1, 2 | TF ChIP-seq |
| ENCFF084NXU | bed narrowPeak | NKRF | GM12878 | 1, 2 | TF ChIP-seq |
| ENCFF462AKP | bed narrowPeak | NR2C1 | GM12878 | 1, 2 | TF ChIP-seq |
| ENCFF782KRV | bed narrowPeak | NR2C2 | GM12878 | 1, 2 | TF ChIP-seq |
| ENCFF531KOV | bed narrowPeak | NR2F1 | GM12878 | 1, 2 | TF ChIP-seq |
| ENCFF652BRY | bed narrowPeak | NRF1 | GM12878 | 1, 2 | TF ChIP-seq |
| ENCFF192WNJ | bed narrowPeak | PAX5 | GM12878 | 1, 2 | TF ChIP-seq |
| ENCFF992JWY | bed narrowPeak | PAX8 | GM12878 | 1, 2 | TF ChIP-seq |
| ENCFF402DQD | bed narrowPeak | PBX3 | GM12878 | 1, 2 | TF ChIP-seq |
| ENCFF335ADU | bed narrowPeak | PKNOX1 | GM12878 | 1, 2 | TF ChIP-seq |
| ENCFF242KVU | bed narrowPeak | PML | GM12878 | 1, 2 | TF ChIP-seq |
| ENCFF886PSD | bed narrowPeak | POLR2A | GM12878 | 1, 2, 3, 4, 5, 6, 7 | TF ChIP-seq |
| ENCFF847DXY | bed narrowPeak | POLR2AphosphoS2 | GM12878 | 1, 2 | TF ChIP-seq |
| ENCFF107UMF | bed narrowPeak | POLR2AphosphoS5 | GM12878 | 1, 2 | TF ChIP-seq |
| ENCFF934JFA | bed narrowPeak | POU2F2 | GM12878 | 1, 2, 3 | TF ChIP-seq |
| ENCFF157HQD | bed narrowPeak | PRDM15 | GM12878 | 1, 2 | TF ChIP-seq |
| ENCFF940IMA | bed narrowPeak | RAD21 | GM12878 | 1, 2 | TF ChIP-seq |
| ENCFF996NBR | bed narrowPeak | RAD51 | GM12878 | 1, 2 | TF ChIP-seq |
| ENCFF034OSV | bed narrowPeak | RB1 | GM12878 | 3, 4 | TF ChIP-seq |
| ENCFF687SSY | bed narrowPeak | RBBP5 | GM12878 | 1, 2 | TF ChIP-seq |
| ENCFF470ZMK | bed narrowPeak | RCOR1 | GM12878 | 1, 2 | TF ChIP-seq |
| ENCFF105YDI | bed narrowPeak | RELB | GM12878 | 1, 2 | TF ChIP-seq |
| ENCFF313CII | bed narrowPeak | REST | GM12878 | 1, 2 | TF ChIP-seq |
| ENCFF259LNG | bed narrowPeak | RFX5 | GM12878 | 1, 2 | TF ChIP-seq |
| ENCFF346JDW | bed narrowPeak | RUNX3 | GM12878 | 1, 2 | TF ChIP-seq |
| ENCFF313BDA | bed narrowPeak | RXRA | GM12878 | 1, 2 | TF ChIP-seq |
| ENCFF050CYK | bed narrowPeak | SIN3A | GM12878 | 1, 2 | TF ChIP-seq |
| ENCFF878FBM | bed narrowPeak | SIX5 | GM12878 | 1, 2 | TF ChIP-seq |
| ENCFF903KEI | bed narrowPeak | SKIL | GM12878 | 1, 2 | TF ChIP-seq |
| ENCFF987PGY | bed narrowPeak | SMAD1 | GM12878 | 1, 2 | TF ChIP-seq |
| ENCFF855SJG | bed narrowPeak | SMAD5 | GM12878 | 1, 2 | TF ChIP-seq |
| ENCFF550FDV | bed narrowPeak | SMARCA5 | GM12878 | 1, 2 | TF ChIP-seq |
| ENCFF837YJA | bed narrowPeak | SMC3 | GM12878 | 1, 2 | TF ChIP-seq |
| ENCFF038AVV | bed narrowPeak | SP1 | GM12878 | 2 | TF ChIP-seq |
| ENCFF071ZMW | bed narrowPeak | SPI1 | GM12878 | 2, 3 | TF ChIP-seq |
| ENCFF397YRO | bed narrowPeak | SREBF1 | GM12878 | 1, 2 | TF ChIP-seq |
| ENCFF896MPE | bed narrowPeak | SREBF2 | GM12878 | 1, 2 | TF ChIP-seq |
| ENCFF114CWH | bed narrowPeak | SRF | GM12878 | 1, 2 | TF ChIP-seq |
| ENCFF896SEI | bed narrowPeak | STAT1 | GM12878 | 1, 2 | TF ChIP-seq |
| ENCFF923CHO | bed narrowPeak | STAT3 | GM12878 | 1, 2 | TF ChIP-seq |
| ENCFF383YEA | bed narrowPeak | STAT5A | GM12878 | 1, 2 | TF ChIP-seq |
| ENCFF069YVD | bed narrowPeak | SUPT20H | GM12878 | 1, 2 | TF ChIP-seq |
| ENCFF894BBO | bed narrowPeak | SUZ12 | GM12878 | 1 | TF ChIP-seq |

| | | | | | |
|---|---|---|---|---|---|
| ENCFF540AAP | bed narrowPeak | TAF1 | GM12878 | 1, 2 | TF ChIP-seq |
| ENCFF668JHK | bed narrowPeak | TARDBP | GM12878 | 1, 2 | TF ChIP-seq |
| ENCFF392JWA | bed narrowPeak | TBL1XR1 | GM12878 | 1, 2 | TF ChIP-seq |
| ENCFF896UZB | bed narrowPeak | TBP | GM12878 | 1, 2 | TF ChIP-seq |
| ENCFF971VHK | bed narrowPeak | TBX21 | GM12878 | 1, 2 | TF ChIP-seq |
| ENCFF768VSH | bed narrowPeak | TCF12 | GM12878 | 1, 2 | TF ChIP-seq |
| ENCFF700TAS | bed narrowPeak | TCF3 | GM12878 | 1, 2 | TF ChIP-seq |
| ENCFF152RNE | bed narrowPeak | TCF7 | GM12878 | 1, 2 | TF ChIP-seq |
| ENCFF552WAH | bed narrowPeak | TRIM22 | GM12878 | 1, 2 | TF ChIP-seq |
| ENCFF295ZLM | bed narrowPeak | UBTF | GM12878 | 1, 2 | TF ChIP-seq |
| ENCFF879TPT | bed narrowPeak | USF1 | GM12878 | 1, 2 | TF ChIP-seq |
| ENCFF514SWA | bed narrowPeak | USF2 | GM12878 | 1, 2 | TF ChIP-seq |
| ENCFF514DDI | bed narrowPeak | WRNIP1 | GM12878 | 1, 2 | TF ChIP-seq |
| ENCFF500RBO | bed narrowPeak | YBX1 | GM12878 | 1, 2 | TF ChIP-seq |
| ENCFF446ZSX | bed narrowPeak | YY1 | GM12878 | 1, 2 | TF ChIP-seq |
| ENCFF630FLK | bed narrowPeak | ZBED1 | GM12878 | 1, 2 | TF ChIP-seq |
| ENCFF475DID | bed narrowPeak | ZBTB33 | GM12878 | 1, 4 | TF ChIP-seq |
| ENCFF084IUW | bed narrowPeak | ZBTB40 | GM12878 | 1, 2 | TF ChIP-seq |
| ENCFF204LCG | bed narrowPeak | ZEB1 | GM12878 | 1, 2 | TF ChIP-seq |
| ENCFF627JQX | bed narrowPeak | ZFP36 | GM12878 | 4, 5 | TF ChIP-seq |
| ENCFF012FCL | bed narrowPeak | ZNF143 | GM12878 | 1, 2 | TF ChIP-seq |
| ENCFF676BIG | bed narrowPeak | ZNF207 | GM12878 | 1, 2 | TF ChIP-seq |
| ENCFF200SLC | bed narrowPeak | ZNF217 | GM12878 | 1, 2 | TF ChIP-seq |
| ENCFF313HBL | bed narrowPeak | ZNF24 | GM12878 | 1, 2 | TF ChIP-seq |
| ENCFF955FRU | bed narrowPeak | ZNF384 | GM12878 | 1, 2 | TF ChIP-seq |
| ENCFF615DTQ | bed narrowPeak | ZNF592 | GM12878 | 1, 2 | TF ChIP-seq |
| ENCFF777DVJ | bed narrowPeak | ZNF622 | GM12878 | 1, 3 | TF ChIP-seq |
| ENCFF137BRA | bed narrowPeak | ZNF687 | GM12878 | 1, 2 | TF ChIP-seq |
| ENCFF214NJL | bed narrowPeak | ZSCAN29 | GM12878 | 1, 2 | TF ChIP-seq |
| ENCFF535WUH | bed narrowPeak | ZZZ3 | GM12878 | 1, 2 | TF ChIP-seq |
| ENCFF693FGQ | bed narrowPeak | ASH2L | H1 | 2 | TF ChIP-seq |
| ENCFF352KLD | bed narrowPeak | ATF2 | H1 | 1, 2 | TF ChIP-seq |
| ENCFF487GLV | bed narrowPeak | ATF3 | H1 | 1, 2 | TF ChIP-seq |
| ENCFF851YHG | bed narrowPeak | BACH1 | H1 | 1, 2 | TF ChIP-seq |
| ENCFF847HXU | bed narrowPeak | BCL11A | H1 | 1 | TF ChIP-seq |
| ENCFF721TNS | bed narrowPeak | BRCA1 | H1 | 1, 2 | TF ChIP-seq |
| ENCFF218OXB | bed narrowPeak | CBX5 | H1 | 1, 2 | TF ChIP-seq |
| ENCFF483UZG | bed narrowPeak | CBX8 | H1 | 1, 2 | TF ChIP-seq |
| ENCFF823KCM | bed narrowPeak | CEBPB | H1 | 1, 2 | TF ChIP-seq |
| ENCFF962NKC | bed narrowPeak | CHD1 | H1 | 1, 2 | TF ChIP-seq |
| ENCFF726GBF | bed narrowPeak | CHD2 | H1 | 1, 2 | TF ChIP-seq |
| ENCFF338IDU | bed narrowPeak | CHD7 | H1 | 1, 2 | TF ChIP-seq |
| ENCFF978KGB | bed narrowPeak | CREB1 | H1 | 1, 2 | TF ChIP-seq |
| ENCFF482FLE | bed narrowPeak | CTBP2 | H1 | 1, 2 | TF ChIP-seq |
| ENCFF023LAA | bed narrowPeak | CTCF | H1 | 1, 2 | TF ChIP-seq |
| ENCFF174AVU | bed narrowPeak | E2F6 | H1 | 1, 2 | TF ChIP-seq |
| ENCFF477ANT | bed narrowPeak | EGR1 | H1 | 1, 2 | TF ChIP-seq |
| ENCFF583FUX | bed narrowPeak | EP300 | H1 | 1 | TF ChIP-seq |
| ENCFF414CAB | bed narrowPeak | EZH2 | H1 | 1, 2 | TF ChIP-seq |
| ENCFF428RHR | bed narrowPeak | FOSL1 | H1 | 1, 2 | TF ChIP-seq |
| ENCFF225GFQ | bed narrowPeak | GABPA | H1 | 1, 2 | TF ChIP-seq |
| ENCFF493PRB | bed narrowPeak | GTF2F1 | H1 | 1, 2 | TF ChIP-seq |

| | | | | | |
|---|---|---|---|---|---|
| ENCFF923TXH | bed narrowPeak | HDAC2 | H1 | 1, 2 | TF ChIP-seq |
| ENCFF802HUU | bed narrowPeak | HDAC6 | H1 | 1 | TF ChIP-seq |
| ENCFF821GUI | bed narrowPeak | JUN | H1 | 1, 2 | TF ChIP-seq |
| ENCFF646IUA | bed narrowPeak | JUND | H1 | 1, 2 | TF ChIP-seq |
| ENCFF759CSN | bed narrowPeak | KDM1A | H1 | 1, 2 | TF ChIP-seq |
| ENCFF021QGZ | bed narrowPeak | KDM4A | H1 | 1 | TF ChIP-seq |
| ENCFF342EEV | bed narrowPeak | KDM5A | H1 | 1, 2 | TF ChIP-seq |
| ENCFF710YJK | bed narrowPeak | MAFK | H1 | 1, 2 | TF ChIP-seq |
| ENCFF460LNS | bed narrowPeak | MAX | H1 | 1, 2 | TF ChIP-seq |
| ENCFF727RNJ | bed narrowPeak | MXI1 | H1 | 1, 2 | TF ChIP-seq |
| ENCFF049SMR | bed narrowPeak | MYC | H1 | 1 | TF ChIP-seq |
| ENCFF435DTC | bed narrowPeak | NANOG | H1 | 1, 2 | TF ChIP-seq |
| ENCFF414RES | bed narrowPeak | NRF1 | H1 | 1, 2 | TF ChIP-seq |
| ENCFF651QOL | bed narrowPeak | PHF8 | H1 | 1, 2 | TF ChIP-seq |
| ENCFF918UHB | bed narrowPeak | POLR2A | H1 | 1 | TF ChIP-seq |
| ENCFF418QVJ | bed narrowPeak | POLR2AphosphoS5 | H1 | 1, 2 | TF ChIP-seq |
| ENCFF383EYO | bed narrowPeak | POU5F1 | H1 | 1, 2 | TF ChIP-seq |
| ENCFF883FUW | bed narrowPeak | RAD21 | H1 | 1, 2 | TF ChIP-seq |
| ENCFF607WCG | bed narrowPeak | RBBP5 | H1 | 1, 2 | TF ChIP-seq |
| ENCFF649VNE | bed narrowPeak | REST | H1 | 1, 2 | TF ChIP-seq |
| ENCFF062WBN | bed narrowPeak | RFX5 | H1 | 1, 2 | TF ChIP-seq |
| ENCFF283MNG | bed narrowPeak | RNF2 | H1 | 1, 2 | TF ChIP-seq |
| ENCFF745EBL | bed narrowPeak | RXRA | H1 | 1, 2 | TF ChIP-seq |
| ENCFF193TFR | bed narrowPeak | SAP30 | H1 | 1, 2 | TF ChIP-seq |
| ENCFF432EYM | bed narrowPeak | SIN3A | H1 | 1, 2 | TF ChIP-seq |
| ENCFF219XEX | bed narrowPeak | SIRT6 | H1 | 1 | TF ChIP-seq |
| ENCFF384KWP | bed narrowPeak | SIX5 | H1 | 1, 2 | TF ChIP-seq |
| ENCFF284JVS | bed narrowPeak | SP1 | H1 | 1, 2 | TF ChIP-seq |
| ENCFF309QRC | bed narrowPeak | SP2 | H1 | 1, 2 | TF ChIP-seq |
| ENCFF257FUV | bed narrowPeak | SP4 | H1 | 1, 2 | TF ChIP-seq |
| ENCFF648QJE | bed narrowPeak | SRF | H1 | 1, 2 | TF ChIP-seq |
| ENCFF233GVJ | bed narrowPeak | SUZ12 | H1 | 1, 3 | TF ChIP-seq |
| ENCFF870SFJ | bed narrowPeak | TAF1 | H1 | 1, 2 | TF ChIP-seq |
| ENCFF243PSJ | bed narrowPeak | TAF7 | H1 | 1, 2 | TF ChIP-seq |
| ENCFF748YXF | bed narrowPeak | TBP | H1 | 1, 2 | TF ChIP-seq |
| ENCFF959HJP | bed narrowPeak | TCF12 | H1 | 1, 2 | TF ChIP-seq |
| ENCFF885PQR | bed narrowPeak | TEAD4 | H1 | 1, 2 | TF ChIP-seq |
| ENCFF699HXL | bed narrowPeak | USF1 | H1 | 1, 2 | TF ChIP-seq |
| ENCFF346KIW | bed narrowPeak | USF2 | H1 | 1, 2 | TF ChIP-seq |
| ENCFF376FVJ | bed narrowPeak | YY1 | H1 | 1, 2 | TF ChIP-seq |
| ENCFF235ROG | bed narrowPeak | ZNF143 | H1 | 1, 2 | TF ChIP-seq |
| ENCFF718OGI | bed narrowPeak | ZNF274 | H1 | 1, 2 | TF ChIP-seq |
| ENCFF913PSL | bed narrowPeak | AEBP2 | HEK293 | 1, 2 | TF ChIP-seq |
| ENCFF225VCG | bed narrowPeak | ATF2 | HEK293 | 1, 2 | TF ChIP-seq |
| ENCFF775QVY | bed narrowPeak | BCL11A | HEK293 | 1, 2 | TF ChIP-seq |
| ENCFF860DSJ | bed narrowPeak | BCL11B | HEK293 | 1, 2 | TF ChIP-seq |
| ENCFF165BAG | bed narrowPeak | BCL6B | HEK293 | 1, 2 | TF ChIP-seq |
| ENCFF071FSR | bed narrowPeak | CTCF | HEK293 | 1, 2 | TF ChIP-seq |
| ENCFF238OFT | bed narrowPeak | EGR2 | HEK293 | 1, 2 | TF ChIP-seq |
| ENCFF226QHL | bed narrowPeak | ELK4 | HEK293 | 1, 2 | TF ChIP-seq |
| ENCFF324ETS | bed narrowPeak | FEZF1 | HEK293 | 1, 2 | TF ChIP-seq |
| ENCFF457HJS | bed narrowPeak | GFI1B | HEK293 | 1, 2 | TF ChIP-seq |

| | | | | |
|---|---|---|---|---|
| ENCFF617QGK | bed narrowPeak | GLI2 | HEK293 | 1, 2 TF ChIP-seq |
| ENCFF322YKQ | bed narrowPeak | GLI4 | HEK293 | 1, 2 TF ChIP-seq |
| ENCFF452YBB | bed narrowPeak | GLIS1 | HEK293 | 1, 2 TF ChIP-seq |
| ENCFF855XBY | bed narrowPeak | GLIS2 | HEK293 | 1, 2 TF ChIP-seq |
| ENCFF473AGH | bed narrowPeak | HIC1 | HEK293 | 1, 2 TF ChIP-seq |
| ENCFF999MOT | bed narrowPeak | IKZF3 | HEK293 | 2, 3 TF ChIP-seq |
| ENCFF929SUL | bed narrowPeak | INSM2 | HEK293 | 1, 2 TF ChIP-seq |
| ENCFF884RBR | bed narrowPeak | KLF1 | HEK293 | 1, 2 TF ChIP-seq |
| ENCFF753PHO | bed narrowPeak | KLF10 | HEK293 | 2, 3 TF ChIP-seq |
| ENCFF303TSY | bed narrowPeak | KLF16 | HEK293 | 1, 2 TF ChIP-seq |
| ENCFF661EXV | bed narrowPeak | KLF17 | HEK293 | 1, 2 TF ChIP-seq |
| ENCFF903AMR | bed narrowPeak | KLF7 | HEK293 | 1, 2 TF ChIP-seq |
| ENCFF399DXM | bed narrowPeak | KLF8 | HEK293 | 1, 2 TF ChIP-seq |
| ENCFF868TRF | bed narrowPeak | KLF9 | HEK293 | 1, 2 TF ChIP-seq |
| ENCFF529XTF | bed narrowPeak | MAZ | HEK293 | 1, 2 TF ChIP-seq |
| ENCFF132UAB | bed narrowPeak | MYNN | HEK293 | 1, 2 TF ChIP-seq |
| ENCFF347IBU | bed narrowPeak | MZF1 | HEK293 | 1, 2 TF ChIP-seq |
| ENCFF784QUI | bed narrowPeak | OSR2 | HEK293 | 1, 2 TF ChIP-seq |
| ENCFF293RRZ | bed narrowPeak | OVOL3 | HEK293 | 1, 2 TF ChIP-seq |
| ENCFF375JQI | bed narrowPeak | PATZ1 | HEK293 | 1, 2 TF ChIP-seq |
| ENCFF138HMJ | bed narrowPeak | PRDM1 | HEK293 | 1, 2 TF ChIP-seq |
| ENCFF680HBX | bed narrowPeak | PRDM10 | HEK293 | 1, 2 TF ChIP-seq |
| ENCFF395BHY | bed narrowPeak | PRDM2 | HEK293 | 1, 2 TF ChIP-seq |
| ENCFF756AYX | bed narrowPeak | PRDM4 | HEK293 | 2, 3 TF ChIP-seq |
| ENCFF540USW | bed narrowPeak | PRDM6 | HEK293 | 1, 2 TF ChIP-seq |
| ENCFF357ZHI | bed narrowPeak | RBAK | HEK293 | 1, 2 TF ChIP-seq |
| ENCFF565NIW | bed narrowPeak | REPIN1 | HEK293 | 1, 3 TF ChIP-seq |
| ENCFF653SCL | bed narrowPeak | REST | HEK293 | 1, 2 TF ChIP-seq |
| ENCFF076TXN | bed narrowPeak | SALL2 | HEK293 | 1, 2 TF ChIP-seq |
| ENCFF799CAL | bed narrowPeak | SCRT1 | HEK293 | 1, 2 TF ChIP-seq |
| ENCFF916ITG | bed narrowPeak | SCRT2 | HEK293 | 1, 2 TF ChIP-seq |
| ENCFF812YDP | bed narrowPeak | SETDB1 | HEK293 | 1 TF ChIP-seq |
| ENCFF676YIL | bed narrowPeak | SP2 | HEK293 | 1, 2 TF ChIP-seq |
| ENCFF663PCO | bed narrowPeak | SP3 | HEK293 | 1, 2 TF ChIP-seq |
| ENCFF925KAK | bed narrowPeak | SP7 | HEK293 | 1, 2 TF ChIP-seq |
| ENCFF136SOA | bed narrowPeak | TCF7L2 | HEK293 | 1, 2 TF ChIP-seq |
| ENCFF211HPP | bed narrowPeak | TRIM28 | HEK293 | 1, 2 TF ChIP-seq |
| ENCFF691JUF | bed narrowPeak | TSHZ1 | HEK293 | 1, 2 TF ChIP-seq |
| ENCFF572OEF | bed narrowPeak | WT1 | HEK293 | 1, 2 TF ChIP-seq |
| ENCFF437JVZ | bed narrowPeak | YY1 | HEK293 | 1, 2 TF ChIP-seq |
| ENCFF814YLC | bed narrowPeak | YY2 | HEK293 | 1, 2 TF ChIP-seq |
| ENCFF765JWP | bed narrowPeak | ZBTB1 | HEK293 | 1, 2 TF ChIP-seq |
| ENCFF511BOQ | bed narrowPeak | ZBTB10 | HEK293 | 1, 2 TF ChIP-seq |
| ENCFF988PII | bed narrowPeak | ZBTB11 | HEK293 | 1, 2 TF ChIP-seq |
| ENCFF904ANO | bed narrowPeak | ZBTB12 | HEK293 | 1, 2 TF ChIP-seq |
| ENCFF299VRF | bed narrowPeak | ZBTB17 | HEK293 | 1, 2 TF ChIP-seq |
| ENCFF076WDH | bed narrowPeak | ZBTB20 | HEK293 | 1, 2 TF ChIP-seq |
| ENCFF313DUS | bed narrowPeak | ZBTB21 | HEK293 | 4, 5 TF ChIP-seq |
| ENCFF668ZRO | bed narrowPeak | ZBTB26 | HEK293 | 1, 2 TF ChIP-seq |
| ENCFF922WQN | bed narrowPeak | ZBTB44 | HEK293 | 2, 3 TF ChIP-seq |
| ENCFF233DCN | bed narrowPeak | ZBTB48 | HEK293 | 1, 2 TF ChIP-seq |
| ENCFF827YPG | bed narrowPeak | ZBTB49 | HEK293 | 1, 2 TF ChIP-seq |

| | | | | |
|---|---|---|---|---|
| ENCFF773KEL | bed narrowPeak | ZBTB6 | HEK293 | 1, 2 TF ChIP-seq |
| ENCFF928KRQ | bed narrowPeak | ZBTB7A | HEK293 | 1, 2 TF ChIP-seq |
| ENCFF480WCB | bed narrowPeak | ZBTB8A | HEK293 | 1, 2 TF ChIP-seq |
| ENCFF662TMD | bed narrowPeak | ZEB1 | HEK293 | 1, 2 TF ChIP-seq |
| ENCFF270KRH | bed narrowPeak | ZEB2 | HEK293 | 1, 2 TF ChIP-seq |
| ENCFF175UAG | bed narrowPeak | ZFHX2 | HEK293 | 1, 2 TF ChIP-seq |
| ENCFF348HKK | bed narrowPeak | ZFP3 | HEK293 | 1, 2 TF ChIP-seq |
| ENCFF668BSL | bed narrowPeak | ZFP37 | HEK293 | 1, 2 TF ChIP-seq |
| ENCFF720MYN | bed narrowPeak | ZFP41 | HEK293 | 1, 2 TF ChIP-seq |
| ENCFF963VBB | bed narrowPeak | ZFP69B | HEK293 | 1, 2 TF ChIP-seq |
| ENCFF187CEY | bed narrowPeak | ZIC2 | HEK293 | 1, 2 TF ChIP-seq |
| ENCFF315FFJ | bed narrowPeak | ZNF10 | HEK293 | 1, 2 TF ChIP-seq |
| ENCFF831DRU | bed narrowPeak | ZNF112 | HEK293 | 1, 2 TF ChIP-seq |
| ENCFF728QYS | bed narrowPeak | ZNF114 | HEK293 | 2, 3 TF ChIP-seq |
| ENCFF496WZS | bed narrowPeak | ZNF121 | HEK293 | 1, 2 TF ChIP-seq |
| ENCFF526AOS | bed narrowPeak | ZNF132 | HEK293 | 1, 2 TF ChIP-seq |
| ENCFF113OWH | bed narrowPeak | ZNF133 | HEK293 | 1, 2 TF ChIP-seq |
| ENCFF334TBU | bed narrowPeak | ZNF138 | HEK293 | 1, 2 TF ChIP-seq |
| ENCFF285WQB | bed narrowPeak | ZNF140 | HEK293 | 1, 2 TF ChIP-seq |
| ENCFF825PWD | bed narrowPeak | ZNF146 | HEK293 | 1, 2 TF ChIP-seq |
| ENCFF960ZJT | bed narrowPeak | ZNF148 | HEK293 | 1, 2 TF ChIP-seq |
| ENCFF091JWQ | bed narrowPeak | ZNF155 | HEK293 | 1, 2 TF ChIP-seq |
| ENCFF498XCM | bed narrowPeak | ZNF157 | HEK293 | 1, 2 TF ChIP-seq |
| ENCFF978XMG | bed narrowPeak | ZNF16 | HEK293 | 1, 2 TF ChIP-seq |
| ENCFF236ZPP | bed narrowPeak | ZNF169 | HEK293 | 1, 2 TF ChIP-seq |
| ENCFF924JMH | bed narrowPeak | ZNF174 | HEK293 | 1, 2 TF ChIP-seq |
| ENCFF004JFD | bed narrowPeak | ZNF175 | HEK293 | 1, 2 TF ChIP-seq |
| ENCFF195OUI | bed narrowPeak | ZNF18 | HEK293 | 1, 2 TF ChIP-seq |
| ENCFF045FIY | bed narrowPeak | ZNF184 | HEK293 | 2, 3 TF ChIP-seq |
| ENCFF824YMX | bed narrowPeak | ZNF189 | HEK293 | 1, 2 TF ChIP-seq |
| ENCFF020TLE | bed narrowPeak | ZNF19 | HEK293 | 1, 2 TF ChIP-seq |
| ENCFF145USV | bed narrowPeak | ZNF2 | HEK293 | 1, 2 TF ChIP-seq |
| ENCFF107CPY | bed narrowPeak | ZNF202 | HEK293 | 1, 2 TF ChIP-seq |
| ENCFF407RMG | bed narrowPeak | ZNF211 | HEK293 | 1, 2 TF ChIP-seq |
| ENCFF786PVX | bed narrowPeak | ZNF214 | HEK293 | 1, 2 TF ChIP-seq |
| ENCFF219VBE | bed narrowPeak | ZNF221 | HEK293 | 1, 2 TF ChIP-seq |
| ENCFF863YLX | bed narrowPeak | ZNF223 | HEK293 | 1, 2 TF ChIP-seq |
| ENCFF655PMG | bed narrowPeak | ZNF23 | HEK293 | 1, 2 TF ChIP-seq |
| ENCFF901CWB | bed narrowPeak | ZNF239 | HEK293 | 1, 2 TF ChIP-seq |
| ENCFF216ALU | bed narrowPeak | ZNF24 | HEK293 | 1, 2 TF ChIP-seq |
| ENCFF982VOB | bed narrowPeak | ZNF248 | HEK293 | 1, 2 TF ChIP-seq |
| ENCFF665NYZ | bed narrowPeak | ZNF26 | HEK293 | 1, 2 TF ChIP-seq |
| ENCFF827SZZ | bed narrowPeak | ZNF263 | HEK293 | 1, 2 TF ChIP-seq |
| ENCFF053AVA | bed narrowPeak | ZNF266 | HEK293 | 1, 2 TF ChIP-seq |
| ENCFF428SMH | bed narrowPeak | ZNF274 | HEK293 | 1, 2 TF ChIP-seq |
| ENCFF120CMN | bed narrowPeak | ZNF280C | HEK293 | 1, 2 TF ChIP-seq |
| ENCFF085OEQ | bed narrowPeak | ZNF280D | HEK293 | 1, 2 TF ChIP-seq |
| ENCFF639RPS | bed narrowPeak | ZNF292 | HEK293 | 1, 2 TF ChIP-seq |
| ENCFF966YLG | bed narrowPeak | ZNF302 | HEK293 | 1, 2 TF ChIP-seq |
| ENCFF975JSZ | bed narrowPeak | ZNF311 | HEK293 | 1, 2 TF ChIP-seq |
| ENCFF902ZKV | bed narrowPeak | ZNF324 | HEK293 | 1, 2 TF ChIP-seq |
| ENCFF887JQV | bed narrowPeak | ZNF335 | HEK293 | 1, 2 TF ChIP-seq |

| | | | | |
|---|---|---|---|---|
| ENCFF123BYP | bed narrowPeak | ZNF34 | HEK293 | 1, 2 TF ChIP-seq |
| ENCFF482CYW | bed narrowPeak | ZNF341 | HEK293 | 1, 2 TF ChIP-seq |
| ENCFF032IPA | bed narrowPeak | ZNF350 | HEK293 | 1, 2 TF ChIP-seq |
| ENCFF036AEX | bed narrowPeak | ZNF354C | HEK293 | 1, 2 TF ChIP-seq |
| ENCFF409GGL | bed narrowPeak | ZNF362 | HEK293 | 1, 2 TF ChIP-seq |
| ENCFF738CQK | bed narrowPeak | ZNF366 | HEK293 | 1, 2 TF ChIP-seq |
| ENCFF328HFF | bed narrowPeak | ZNF37A | HEK293 | 2, 3 TF ChIP-seq |
| ENCFF032CZR | bed narrowPeak | ZNF391 | HEK293 | 1, 2 TF ChIP-seq |
| ENCFF886YDF | bed narrowPeak | ZNF394 | HEK293 | 1, 2 TF ChIP-seq |
| ENCFF772OQU | bed narrowPeak | ZNF398 | HEK293 | 1, 2 TF ChIP-seq |
| ENCFF444CQQ | bed narrowPeak | ZNF404 | HEK293 | 1, 2 TF ChIP-seq |
| ENCFF126IZP | bed narrowPeak | ZNF416 | HEK293 | 1, 2 TF ChIP-seq |
| ENCFF311IXZ | bed narrowPeak | ZNF423 | HEK293 | 1, 2 TF ChIP-seq |
| ENCFF193XBL | bed narrowPeak | ZNF426 | HEK293 | 1, 2 TF ChIP-seq |
| ENCFF139XGF | bed narrowPeak | ZNF433 | HEK293 | 1, 2 TF ChIP-seq |
| ENCFF886WTI | bed narrowPeak | ZNF449 | HEK293 | 1, 2 TF ChIP-seq |
| ENCFF604AVO | bed narrowPeak | ZNF473 | HEK293 | 1, 2 TF ChIP-seq |
| ENCFF265LNU | bed narrowPeak | ZNF48 | HEK293 | 1, 2 TF ChIP-seq |
| ENCFF684WPJ | bed narrowPeak | ZNF488 | HEK293 | 1, 2 TF ChIP-seq |
| ENCFF090VNO | bed narrowPeak | ZNF501 | HEK293 | 1, 2 TF ChIP-seq |
| ENCFF193LIK | bed narrowPeak | ZNF510 | HEK293 | 1, 2 TF ChIP-seq |
| ENCFF970URF | bed narrowPeak | ZNF513 | HEK293 | 2, 3 TF ChIP-seq |
| ENCFF596KQL | bed narrowPeak | ZNF514 | HEK293 | 1, 2 TF ChIP-seq |
| ENCFF834NNK | bed narrowPeak | ZNF518A | HEK293 | 1, 2 TF ChIP-seq |
| ENCFF996QGV | bed narrowPeak | ZNF521 | HEK293 | 1, 2 TF ChIP-seq |
| ENCFF806WJB | bed narrowPeak | ZNF529 | HEK293 | 1, 2 TF ChIP-seq |
| ENCFF729VVD | bed narrowPeak | ZNF530 | HEK293 | 1, 2 TF ChIP-seq |
| ENCFF673JLB | bed narrowPeak | ZNF544 | HEK293 | 1, 2 TF ChIP-seq |
| ENCFF576BGV | bed narrowPeak | ZNF547 | HEK293 | 1, 2 TF ChIP-seq |
| ENCFF980EGS | bed narrowPeak | ZNF548 | HEK293 | 1, 2 TF ChIP-seq |
| ENCFF237SGW | bed narrowPeak | ZNF549 | HEK293 | 2, 3 TF ChIP-seq |
| ENCFF852JPW | bed narrowPeak | ZNF555 | HEK293 | 1, 2 TF ChIP-seq |
| ENCFF755FMU | bed narrowPeak | ZNF558 | HEK293 | 1, 2 TF ChIP-seq |
| ENCFF173MMC | bed narrowPeak | ZNF560 | HEK293 | 2, 3 TF ChIP-seq |
| ENCFF127PKO | bed narrowPeak | ZNF561 | HEK293 | 2, 3 TF ChIP-seq |
| ENCFF388XSN | bed narrowPeak | ZNF571 | HEK293 | 1, 2 TF ChIP-seq |
| ENCFF631BJV | bed narrowPeak | ZNF580 | HEK293 | 1, 2 TF ChIP-seq |
| ENCFF955EQI | bed narrowPeak | ZNF585B | HEK293 | 1, 2 TF ChIP-seq |
| ENCFF709SOO | bed narrowPeak | ZNF596 | HEK293 | 2, 3 TF ChIP-seq |
| ENCFF326MMO | bed narrowPeak | ZNF600 | HEK293 | 1, 2 TF ChIP-seq |
| ENCFF339XZT | bed narrowPeak | ZNF610 | HEK293 | 1, 2 TF ChIP-seq |
| ENCFF258PUP | bed narrowPeak | ZNF621 | HEK293 | 1, 2 TF ChIP-seq |
| ENCFF269NKC | bed narrowPeak | ZNF623 | HEK293 | 1, 2 TF ChIP-seq |
| ENCFF580LRK | bed narrowPeak | ZNF624 | HEK293 | 1, 2 TF ChIP-seq |
| ENCFF361TIN | bed narrowPeak | ZNF626 | HEK293 | 2, 3 TF ChIP-seq |
| ENCFF167ZZY | bed narrowPeak | ZNF629 | HEK293 | 2, 3 TF ChIP-seq |
| ENCFF747QYQ | bed narrowPeak | ZNF639 | HEK293 | 1, 2 TF ChIP-seq |
| ENCFF657RPA | bed narrowPeak | ZNF645 | HEK293 | 1, 2 TF ChIP-seq |
| ENCFF149OME | bed narrowPeak | ZNF654 | HEK293 | 1, 2 TF ChIP-seq |
| ENCFF894CGQ | bed narrowPeak | ZNF658 | HEK293 | 1, 2 TF ChIP-seq |
| ENCFF197LYS | bed narrowPeak | ZNF660 | HEK293 | 2, 3 TF ChIP-seq |
| ENCFF337BRJ | bed narrowPeak | ZNF664 | HEK293 | 1, 2 TF ChIP-seq |

| | | | | | |
|---|---|---|---|---|---|
| ENCFF869NMB | bed narrowPeak | ZNF670 | HEK293 | 2, 3 | TF ChIP-seq |
| ENCFF153JJQ | bed narrowPeak | ZNF677 | HEK293 | 2, 3 | TF ChIP-seq |
| ENCFF359XJQ | bed narrowPeak | ZNF680 | HEK293 | 1, 2 | TF ChIP-seq |
| ENCFF167MHN | bed narrowPeak | ZNF692 | HEK293 | 1, 2 | TF ChIP-seq |
| ENCFF832OST | bed narrowPeak | ZNF701 | HEK293 | 1, 2 | TF ChIP-seq |
| ENCFF137FMN | bed narrowPeak | ZNF704 | HEK293 | 1, 2 | TF ChIP-seq |
| ENCFF735QIK | bed narrowPeak | ZNF707 | HEK293 | 1, 2 | TF ChIP-seq |
| ENCFF029DNH | bed narrowPeak | ZNF747 | HEK293 | 1, 2 | TF ChIP-seq |
| ENCFF170VVR | bed narrowPeak | ZNF76 | HEK293 | 1, 2 | TF ChIP-seq |
| ENCFF713BAD | bed narrowPeak | ZNF768 | HEK293 | 1, 2 | TF ChIP-seq |
| ENCFF554RDX | bed narrowPeak | ZNF770 | HEK293 | 1, 2 | TF ChIP-seq |
| ENCFF836LWG | bed narrowPeak | ZNF776 | HEK293 | 1, 2 | TF ChIP-seq |
| ENCFF804ZLF | bed narrowPeak | ZNF777 | HEK293 | 1, 2 | TF ChIP-seq |
| ENCFF261IAG | bed narrowPeak | ZNF785 | HEK293 | 1, 2 | TF ChIP-seq |
| ENCFF576BRL | bed narrowPeak | ZNF791 | HEK293 | 1, 2 | TF ChIP-seq |
| ENCFF652HBV | bed narrowPeak | ZNF792 | HEK293 | 1, 2 | TF ChIP-seq |
| ENCFF667JVF | bed narrowPeak | ZNF830 | HEK293 | 2, 3 | TF ChIP-seq |
| ENCFF045YGV | bed narrowPeak | ZNF837 | HEK293 | 1, 2 | TF ChIP-seq |
| ENCFF381VJM | bed narrowPeak | ZNF843 | HEK293 | 1, 2 | TF ChIP-seq |
| ENCFF799OTE | bed narrowPeak | ZSCAN16 | HEK293 | 1, 2 | TF ChIP-seq |
| ENCFF509YKU | bed narrowPeak | ZSCAN18 | HEK293 | 1, 2 | TF ChIP-seq |
| ENCFF676QQC | bed narrowPeak | ZSCAN21 | HEK293 | 1, 2 | TF ChIP-seq |
| ENCFF312JKD | bed narrowPeak | ZSCAN23 | HEK293 | 1, 2 | TF ChIP-seq |
| ENCFF500ABD | bed narrowPeak | ZSCAN26 | HEK293 | 1, 2 | TF ChIP-seq |
| ENCFF530LIW | bed narrowPeak | ZSCAN30 | HEK293 | 2, 3 | TF ChIP-seq |
| ENCFF815ZYA | bed narrowPeak | ZSCAN4 | HEK293 | 1, 2 | TF ChIP-seq |
| ENCFF280IWS | bed narrowPeak | ZSCAN5A | HEK293 | 1, 2 | TF ChIP-seq |
| ENCFF679DYR | bed narrowPeak | ZSCAN5C | HEK293 | 1, 2 | TF ChIP-seq |
| ENCFF427HLT | bed narrowPeak | ZXDB | HEK293 | 1, 2 | TF ChIP-seq |
| ENCFF143SAQ | bed narrowPeak | BRCA1 | HeLa-S3 | 1, 2 | TF ChIP-seq |
| ENCFF621LKI | bed narrowPeak | BRF1 | HeLa-S3 | 1, 2 | TF ChIP-seq |
| ENCFF645KWQ | bed narrowPeak | BRF2 | HeLa-S3 | 1, 2 | TF ChIP-seq |
| ENCFF014TJO | bed narrowPeak | CEBPB | HeLa-S3 | 1, 2 | TF ChIP-seq |
| ENCFF644XCT | bed narrowPeak | CHD2 | HeLa-S3 | 1, 2 | TF ChIP-seq |
| ENCFF548ARU | bed narrowPeak | CTCF | HeLa-S3 | 1, 2 | TF ChIP-seq |
| ENCFF049NRL | bed narrowPeak | DEK | HeLa-S3 | 1, 2 | TF ChIP-seq |
| ENCFF872RSG | bed narrowPeak | E2F1 | HeLa-S3 | 1, 2 | TF ChIP-seq |
| ENCFF718JTU | bed narrowPeak | E2F4 | HeLa-S3 | 1, 2 | TF ChIP-seq |
| ENCFF160MCO | bed narrowPeak | E2F6 | HeLa-S3 | 1, 2 | TF ChIP-seq |
| ENCFF279OML | bed narrowPeak | ELK1 | HeLa-S3 | 1, 2 | TF ChIP-seq |
| ENCFF582BHA | bed narrowPeak | ELK4 | HeLa-S3 | 1, 2 | TF ChIP-seq |
| ENCFF631WOD | bed narrowPeak | EP300 | HeLa-S3 | 1, 2 | TF ChIP-seq |
| ENCFF568QUD | bed narrowPeak | EZH2 | HeLa-S3 | 1, 2 | TF ChIP-seq |
| ENCFF494YNM | bed narrowPeak | FOS | HeLa-S3 | 1, 2 | TF ChIP-seq |
| ENCFF168BDU | bed narrowPeak | GABPA | HeLa-S3 | 1, 2 | TF ChIP-seq |
| ENCFF394JHN | bed narrowPeak | GTF2F1 | HeLa-S3 | 1, 2 | TF ChIP-seq |
| ENCFF094AXB | bed narrowPeak | HCFC1 | HeLa-S3 | 1, 2 | TF ChIP-seq |
| ENCFF291VFT | bed narrowPeak | IRF3 | HeLa-S3 | 1, 2 | TF ChIP-seq |
| ENCFF806AQE | bed narrowPeak | JUN | HeLa-S3 | 1, 2 | TF ChIP-seq |
| ENCFF995QMN | bed narrowPeak | JUND | HeLa-S3 | 1, 2 | TF ChIP-seq |
| ENCFF013GAH | bed narrowPeak | KAT2A | HeLa-S3 | 1, 2 | TF ChIP-seq |
| ENCFF280WZN | bed narrowPeak | MAFF | HeLa-S3 | 2, 3 | TF ChIP-seq |

| | | | | | |
|---|---|---|---|---|---|
| ENCFF328IZQ | bed narrowPeak | MAFK | HeLa-S3 | 1, 2 | TF ChIP-seq |
| ENCFF666XCS | bed narrowPeak | MAX | HeLa-S3 | 1, 2 | TF ChIP-seq |
| ENCFF424JHD | bed narrowPeak | MAZ | HeLa-S3 | 1, 2 | TF ChIP-seq |
| ENCFF420AQP | bed narrowPeak | MXI1 | HeLa-S3 | 1, 2 | TF ChIP-seq |
| ENCFF515BTV | bed narrowPeak | MYC | HeLa-S3 | 1, 2 | TF ChIP-seq |
| ENCFF664AZO | bed narrowPeak | NFE2L2 | HeLa-S3 | 1, 2 | TF ChIP-seq |
| ENCFF865QSV | bed narrowPeak | NFYA | HeLa-S3 | 1, 2 | TF ChIP-seq |
| ENCFF191KVV | bed narrowPeak | NFYB | HeLa-S3 | 1, 2 | TF ChIP-seq |
| ENCFF703LZR | bed narrowPeak | NR2C2 | HeLa-S3 | 1, 2 | TF ChIP-seq |
| ENCFF730BLH | bed narrowPeak | NRF1 | HeLa-S3 | 1, 2 | TF ChIP-seq |
| ENCFF245CAL | bed narrowPeak | POLR2A | HeLa-S3 | 1, 2, 3 | TF ChIP-seq |
| ENCFF867RKD | bed narrowPeak | POLR2AphosphoS2 | HeLa-S3 | 1, 2 | TF ChIP-seq |
| ENCFF767DMU | bed narrowPeak | POLR3A | HeLa-S3 | 1, 2 | TF ChIP-seq |
| ENCFF721NYV | bed narrowPeak | PRDM1 | HeLa-S3 | 1, 2 | TF ChIP-seq |
| ENCFF239FBO | bed narrowPeak | RAD21 | HeLa-S3 | 1, 2 | TF ChIP-seq |
| ENCFF753CEB | bed narrowPeak | RCOR1 | HeLa-S3 | 1, 2 | TF ChIP-seq |
| ENCFF096JDA | bed narrowPeak | REST | HeLa-S3 | 1, 2 | TF ChIP-seq |
| ENCFF438ZEN | bed narrowPeak | RFX5 | HeLa-S3 | 1, 2 | TF ChIP-seq |
| ENCFF216YDM | bed narrowPeak | SMARCA4 | HeLa-S3 | 1, 2, 3 | TF ChIP-seq |
| ENCFF572FHR | bed narrowPeak | SMARCB1 | HeLa-S3 | 1, 2, 3 | TF ChIP-seq |
| ENCFF492BST | bed narrowPeak | SMARCC1 | HeLa-S3 | 1, 2, 3 | TF ChIP-seq |
| ENCFF253UAA | bed narrowPeak | SMARCC2 | HeLa-S3 | 1, 2, 3 | TF ChIP-seq |
| ENCFF141KLW | bed narrowPeak | SMC3 | HeLa-S3 | 1, 2 | TF ChIP-seq |
| ENCFF785YII | bed narrowPeak | SREBF2 | HeLa-S3 | 2, 3 | TF ChIP-seq |
| ENCFF746VUX | bed narrowPeak | STAT3 | HeLa-S3 | 1, 2 | TF ChIP-seq |
| ENCFF044DFE | bed narrowPeak | SUPT20H | HeLa-S3 | 1, 2 | TF ChIP-seq |
| ENCFF045BFG | bed narrowPeak | TAF1 | HeLa-S3 | 1, 2 | TF ChIP-seq |
| ENCFF363NQY | bed narrowPeak | TBP | HeLa-S3 | 1, 2 | TF ChIP-seq |
| ENCFF936XIP | bed narrowPeak | TCF7L2 | HeLa-S3 | 1, 2 | TF ChIP-seq |
| ENCFF834LQR | bed narrowPeak | UBTF | HeLa-S3 | 1, 2 | TF ChIP-seq |
| ENCFF405SDO | bed narrowPeak | USF2 | HeLa-S3 | 1, 2 | TF ChIP-seq |
| ENCFF346ZGY | bed narrowPeak | ZFP36 | HeLa-S3 | 1, 3 | TF ChIP-seq |
| ENCFF267DZF | bed narrowPeak | ZHX1 | HeLa-S3 | 1, 2 | TF ChIP-seq |
| ENCFF945JJR | bed narrowPeak | ZKSCAN1 | HeLa-S3 | 1, 2 | TF ChIP-seq |
| ENCFF236NLF | bed narrowPeak | ZNF143 | HeLa-S3 | 1, 2 | TF ChIP-seq |
| ENCFF546RIC | bed narrowPeak | ZNF274 | HeLa-S3 | 1, 2 | TF ChIP-seq |
| ENCFF675ARG | bed narrowPeak | ZZZ3 | HeLa-S3 | 1, 2 | TF ChIP-seq |
| ENCFF162UZN | bed narrowPeak | ADNP | HepG2 | 1, 2 | TF ChIP-seq |
| ENCFF026PPP | bed narrowPeak | AFF4 | HepG2 | 1, 2 | TF ChIP-seq |
| ENCFF627BHP | bed narrowPeak | AGO1 | HepG2 | 1, 2 | TF ChIP-seq |
| ENCFF465FII | bed narrowPeak | AGO2 | HepG2 | 1, 2 | TF ChIP-seq |
| ENCFF048HNT | bed narrowPeak | AHDC1 | HepG2 | 1, 2 | TF ChIP-seq |
| ENCFF683QEX | bed narrowPeak | AHR | HepG2 | 1, 2 | TF ChIP-seq |
| ENCFF716HAB | bed narrowPeak | AKAP8 | HepG2 | 1, 2 | TF ChIP-seq |
| ENCFF411GOS | bed narrowPeak | AKNA | HepG2 | 1, 2 | TF ChIP-seq |
| ENCFF247GXE | bed narrowPeak | ARID3A | HepG2 | 1, 2 | TF ChIP-seq |
| ENCFF258MNE | bed narrowPeak | ARID4A | HepG2 | 1, 2 | TF ChIP-seq |
| ENCFF492BXK | bed narrowPeak | ARID4B | HepG2 | 1, 2 | TF ChIP-seq |
| ENCFF477OUI | bed narrowPeak | ARID5B | HepG2 | 1, 2 | TF ChIP-seq |
| ENCFF616WXJ | bed narrowPeak | ARNT | HepG2 | 1, 2 | TF ChIP-seq |
| ENCFF588WBI | bed narrowPeak | ARNTL | HepG2 | 1, 2 | TF ChIP-seq |
| ENCFF230OBH | bed narrowPeak | ASH2L | HepG2 | 1, 2 | TF ChIP-seq |

| | | | | | |
|---|---|---|---|---|---|
| ENCFF585SZT | bed narrowPeak | ATAD3A | HepG2 | 1, 2 | TF ChIP-seq |
| ENCFF683OFC | bed narrowPeak | ATF1 | HepG2 | 1, 2 | TF ChIP-seq |
| ENCFF522LHU | bed narrowPeak | ATF2 | HepG2 | 1, 2 | TF ChIP-seq |
| ENCFF349HIP | bed narrowPeak | ATF3 | HepG2 | 1, 2 | TF ChIP-seq |
| ENCFF266GBP | bed narrowPeak | ATF4 | HepG2 | 1, 2 | TF ChIP-seq |
| ENCFF482BEV | bed narrowPeak | ATF5 | HepG2 | 1, 2 | TF ChIP-seq |
| ENCFF894YFW | bed narrowPeak | ATF7 | HepG2 | 1, 2 | TF ChIP-seq |
| ENCFF906FVB | bed narrowPeak | ATM | HepG2 | 1, 2 | TF ChIP-seq |
| ENCFF998FLA | bed narrowPeak | ATRX | HepG2 | 1, 2 | TF ChIP-seq |
| ENCFF424FZS | bed narrowPeak | BCL3 | HepG2 | 1, 2 | TF ChIP-seq |
| ENCFF014UTU | bed narrowPeak | BCL6 | HepG2 | 1, 2 | TF ChIP-seq |
| ENCFF506PXL | bed narrowPeak | BCLAF1 | HepG2 | 1, 2 | TF ChIP-seq |
| ENCFF992EBV | bed narrowPeak | BHLHE40 | HepG2 | 1, 2 | TF ChIP-seq |
| ENCFF791SNR | bed narrowPeak | BRCA1 | HepG2 | 1, 2 | TF ChIP-seq |
| ENCFF440ODC | bed narrowPeak | BRD4 | HepG2 | 1, 2 | TF ChIP-seq |
| ENCFF333PIQ | bed narrowPeak | CAMTA2 | HepG2 | 1, 2 | TF ChIP-seq |
| ENCFF951CCL | bed narrowPeak | CBFB | HepG2 | 1, 2 | TF ChIP-seq |
| ENCFF566PBY | bed narrowPeak | CBX1 | HepG2 | 1 | TF ChIP-seq |
| ENCFF501QII | bed narrowPeak | CBX2 | HepG2 | 1, 2 | TF ChIP-seq |
| ENCFF486LNN | bed narrowPeak | CBX5 | HepG2 | 1, 2 | TF ChIP-seq |
| ENCFF039LHY | bed narrowPeak | CCAR2 | HepG2 | 1, 2 | TF ChIP-seq |
| ENCFF874PIO | bed narrowPeak | CEBPA | HepG2 | 1, 2 | TF ChIP-seq |
| ENCFF157DSR | bed narrowPeak | CEBPB | HepG2 | 1, 2 | TF ChIP-seq |
| ENCFF929XBL | bed narrowPeak | CEBPD | HepG2 | 1, 2 | TF ChIP-seq |
| ENCFF094SJQ | bed narrowPeak | CEBPG | HepG2 | 1, 2 | TF ChIP-seq |
| ENCFF195BKI | bed narrowPeak | CEBPZ | HepG2 | 1, 2 | TF ChIP-seq |
| ENCFF821HRB | bed narrowPeak | CENPX | HepG2 | 1, 2 | TF ChIP-seq |
| ENCFF209MHH | bed narrowPeak | CERS6 | HepG2 | 1, 2 | TF ChIP-seq |
| ENCFF102ZOU | bed narrowPeak | CHD2 | HepG2 | 1, 2 | TF ChIP-seq |
| ENCFF691DTG | bed narrowPeak | CHD4 | HepG2 | 1, 2 | TF ChIP-seq |
| ENCFF955MFB | bed narrowPeak | CREB1 | HepG2 | 1, 2 | TF ChIP-seq |
| ENCFF767ZRB | bed narrowPeak | CREB3 | HepG2 | 1, 2 | TF ChIP-seq |
| ENCFF871PDU | bed narrowPeak | CREM | HepG2 | 1, 2 | TF ChIP-seq |
| ENCFF664UGR | bed narrowPeak | CTCF | HepG2 | 1, 2 | TF ChIP-seq |
| ENCFF544WSX | bed narrowPeak | DBP | HepG2 | 1, 2 | TF ChIP-seq |
| ENCFF659PQJ | bed narrowPeak | DDIT3 | HepG2 | 1, 2 | TF ChIP-seq |
| ENCFF707SJC | bed narrowPeak | DLX6 | HepG2 | 1, 2 | TF ChIP-seq |
| ENCFF092ZIZ | bed narrowPeak | DMAP1 | HepG2 | 1, 2 | TF ChIP-seq |
| ENCFF598CBU | bed narrowPeak | DMTF1 | HepG2 | 1, 2 | TF ChIP-seq |
| ENCFF622GUA | bed narrowPeak | DNMT1 | HepG2 | 1, 2 | TF ChIP-seq |
| ENCFF186YGH | bed narrowPeak | DNMT3B | HepG2 | 1, 2 | TF ChIP-seq |
| ENCFF007COI | bed narrowPeak | DPF2 | HepG2 | 1, 2 | TF ChIP-seq |
| ENCFF089DNE | bed narrowPeak | DR1 | HepG2 | 1, 2 | TF ChIP-seq |
| ENCFF052EOE | bed narrowPeak | DRAP1 | HepG2 | 1, 2 | TF ChIP-seq |
| ENCFF015STD | bed narrowPeak | DZIP1 | HepG2 | 1, 2 | TF ChIP-seq |
| ENCFF049BWK | bed narrowPeak | E2F1 | HepG2 | 1, 2 | TF ChIP-seq |
| ENCFF372PXU | bed narrowPeak | E2F2 | HepG2 | 1, 2 | TF ChIP-seq |
| ENCFF245XOL | bed narrowPeak | E2F4 | HepG2 | 1, 2 | TF ChIP-seq |
| ENCFF217STG | bed narrowPeak | EEA1 | HepG2 | 1, 2 | TF ChIP-seq |
| ENCFF405LDS | bed narrowPeak | EED | HepG2 | 1, 2 | TF ChIP-seq |
| ENCFF603PUS | bed narrowPeak | EGR1 | HepG2 | 1, 2 | TF ChIP-seq |
| ENCFF855VUF | bed narrowPeak | EHMT2 | HepG2 | 1, 2 | TF ChIP-seq |

| | | | | |
|---|---|---|---|---|
| ENCFF215GPK | bed narrowPeak | ELF1 | HepG2 | 1, 2 TF ChIP-seq |
| ENCFF080FAU | bed narrowPeak | ELF3 | HepG2 | 1, 2 TF ChIP-seq |
| ENCFF488DML | bed narrowPeak | ELF4 | HepG2 | 1, 2 TF ChIP-seq |
| ENCFF399YOO | bed narrowPeak | ELK1 | HepG2 | 1, 2 TF ChIP-seq |
| ENCFF890ZHY | bed narrowPeak | ELK4 | HepG2 | 1, 2 TF ChIP-seq |
| ENCFF393LUP | bed narrowPeak | EP300 | HepG2 | 1, 2 TF ChIP-seq |
| ENCFF554LPO | bed narrowPeak | ERF | HepG2 | 1, 2 TF ChIP-seq |
| ENCFF700DMQ | bed narrowPeak | ESRRA | HepG2 | 1, 2 TF ChIP-seq |
| ENCFF191NVV | bed narrowPeak | ETS1 | HepG2 | 1, 2 TF ChIP-seq |
| ENCFF624RGO | bed narrowPeak | ETV4 | HepG2 | 1, 2 TF ChIP-seq |
| ENCFF712HHN | bed narrowPeak | ETV5 | HepG2 | 1, 2 TF ChIP-seq |
| ENCFF504QZJ | bed narrowPeak | EZH2 | HepG2 | 1, 2 TF ChIP-seq |
| ENCFF031LBW | bed narrowPeak | FIP1L1 | HepG2 | 1, 2 TF ChIP-seq |
| ENCFF497SIC | bed narrowPeak | FOSL1 | HepG2 | 1, 2 TF ChIP-seq |
| ENCFF772SCR | bed narrowPeak | FOSL2 | HepG2 | 1, 2 TF ChIP-seq |
| ENCFF466RZX | bed narrowPeak | FOXA1 | HepG2 | 1, 2 TF ChIP-seq |
| ENCFF759YCY | bed narrowPeak | FOXA2 | HepG2 | 1, 2 TF ChIP-seq |
| ENCFF251TUO | bed narrowPeak | FOXA3 | HepG2 | 1, 2 TF ChIP-seq |
| ENCFF548ZQR | bed narrowPeak | FOXJ3 | HepG2 | 1, 2 TF ChIP-seq |
| ENCFF605GXY | bed narrowPeak | FOXK1 | HepG2 | 1, 2 TF ChIP-seq |
| ENCFF315CHX | bed narrowPeak | FOXK2 | HepG2 | 1, 2 TF ChIP-seq |
| ENCFF567QWP | bed narrowPeak | FOXM1 | HepG2 | 1, 2 TF ChIP-seq |
| ENCFF331BSI | bed narrowPeak | FOXO1 | HepG2 | 1, 2 TF ChIP-seq |
| ENCFF460UOW | bed narrowPeak | FOXP1 | HepG2 | 1, 2 TF ChIP-seq |
| ENCFF163LGN | bed narrowPeak | FOXP4 | HepG2 | 1, 2 TF ChIP-seq |
| ENCFF139XIZ | bed narrowPeak | FUBP1 | HepG2 | 1, 2 TF ChIP-seq |
| ENCFF953FNE | bed narrowPeak | FUBP3 | HepG2 | 1, 2 TF ChIP-seq |
| ENCFF216YZI | bed narrowPeak | FUS | HepG2 | 1, 2 TF ChIP-seq |
| ENCFF890WJZ | bed narrowPeak | GABPA | HepG2 | 1, 2 TF ChIP-seq |
| ENCFF217XUD | bed narrowPeak | GABPB1 | HepG2 | 1, 2 TF ChIP-seq |
| ENCFF409YKQ | bed narrowPeak | GATA2 | HepG2 | 1, 2 TF ChIP-seq |
| ENCFF097OXR | bed narrowPeak | GATA4 | HepG2 | 1, 2 TF ChIP-seq |
| ENCFF928HFL | bed narrowPeak | GATAD1 | HepG2 | 1, 2 TF ChIP-seq |
| ENCFF158BJH | bed narrowPeak | GATAD2A | HepG2 | 1, 2 TF ChIP-seq |
| ENCFF999HNJ | bed narrowPeak | GFI1 | HepG2 | 1, 2 TF ChIP-seq |
| ENCFF802VTG | bed narrowPeak | GLI4 | HepG2 | 1, 2 TF ChIP-seq |
| ENCFF745JFN | bed narrowPeak | GLMP | HepG2 | 1, 2 TF ChIP-seq |
| ENCFF004QJZ | bed narrowPeak | GMEB1 | HepG2 | 1, 2 TF ChIP-seq |
| ENCFF219KQD | bed narrowPeak | GMEB2 | HepG2 | 1, 2 TF ChIP-seq |
| ENCFF019SZJ | bed narrowPeak | GPBP1L1 | HepG2 | 1, 2 TF ChIP-seq |
| ENCFF955LND | bed narrowPeak | GPN1 | HepG2 | 1, 2 TF ChIP-seq |
| ENCFF599TWF | bed narrowPeak | GTF2F1 | HepG2 | 1, 2 TF ChIP-seq |
| ENCFF190LEQ | bed narrowPeak | GZF1 | HepG2 | 1, 2 TF ChIP-seq |
| ENCFF894AWT | bed narrowPeak | HBP1 | HepG2 | 1, 2 TF ChIP-seq |
| ENCFF485SRU | bed narrowPeak | HCFC1 | HepG2 | 1, 2 TF ChIP-seq |
| ENCFF705AOW | bed narrowPeak | HDAC1 | HepG2 | 1, 2 TF ChIP-seq |
| ENCFF221AOL | bed narrowPeak | HDAC2 | HepG2 | 1, 2 TF ChIP-seq |
| ENCFF109EXK | bed narrowPeak | HDAC6 | HepG2 | 1, 2 TF ChIP-seq |
| ENCFF714AQQ | bed narrowPeak | HES4 | HepG2 | 1, 2 TF ChIP-seq |
| ENCFF207CLF | bed narrowPeak | HHEX | HepG2 | 1, 2 TF ChIP-seq |
| ENCFF445FCL | bed narrowPeak | HINFP | HepG2 | 1, 2 TF ChIP-seq |
| ENCFF076ESL | bed narrowPeak | HIVEP1 | HepG2 | 1, 2 TF ChIP-seq |

| | | | | |
|---|---|---|---|---|
| ENCFF685UQE | bed narrowPeak | HLF | HepG2 | 1, 2 TF ChIP-seq |
| ENCFF914TSO | bed narrowPeak | HMG20A | HepG2 | 1, 2 TF ChIP-seq |
| ENCFF727GHV | bed narrowPeak | HMG20B | HepG2 | 1, 2 TF ChIP-seq |
| ENCFF684UHQ | bed narrowPeak | HMGXB3 | HepG2 | 1, 2 TF ChIP-seq |
| ENCFF305WWR | bed narrowPeak | HMGXB4 | HepG2 | 1, 2 TF ChIP-seq |
| ENCFF767MSS | bed narrowPeak | HNF1A | HepG2 | 1, 2 TF ChIP-seq |
| ENCFF022QCK | bed narrowPeak | HNF1B | HepG2 | 1, 2 TF ChIP-seq |
| ENCFF704BPD | bed narrowPeak | HNF4A | HepG2 | 1, 2 TF ChIP-seq |
| ENCFF123FQW | bed narrowPeak | HNF4G | HepG2 | 1, 2 TF ChIP-seq |
| ENCFF046NUR | bed narrowPeak | HNRNPH1 | HepG2 | 1, 2 TF ChIP-seq |
| ENCFF035OPG | bed narrowPeak | HNRNPK | HepG2 | 1, 2 TF ChIP-seq |
| ENCFF039CUI | bed narrowPeak | HNRNPL | HepG2 | 1, 2 TF ChIP-seq |
| ENCFF890KTX | bed narrowPeak | HNRNPLL | HepG2 | 1, 2 TF ChIP-seq |
| ENCFF509YFF | bed narrowPeak | HNRNPUL1 | HepG2 | 1, 2 TF ChIP-seq |
| ENCFF478OVU | bed narrowPeak | HOMEZ | HepG2 | 1, 2 TF ChIP-seq |
| ENCFF753OCA | bed narrowPeak | HOXA3 | HepG2 | 1, 2 TF ChIP-seq |
| ENCFF385DPO | bed narrowPeak | HOXA5 | HepG2 | 1, 2 TF ChIP-seq |
| ENCFF647VJL | bed narrowPeak | HOXD1 | HepG2 | 1, 2 TF ChIP-seq |
| ENCFF062UNZ | bed narrowPeak | HSF2 | HepG2 | 1, 2 TF ChIP-seq |
| ENCFF969BZA | bed narrowPeak | IKZF1 | HepG2 | 1, 2 TF ChIP-seq |
| ENCFF948XVV | bed narrowPeak | IKZF4 | HepG2 | 1, 2 TF ChIP-seq |
| ENCFF371LUB | bed narrowPeak | IKZF5 | HepG2 | 1, 2 TF ChIP-seq |
| ENCFF368IYV | bed narrowPeak | IRF1 | HepG2 | 1, 2 TF ChIP-seq |
| ENCFF095ZXG | bed narrowPeak | IRF2 | HepG2 | 1, 2 TF ChIP-seq |
| ENCFF619GRL | bed narrowPeak | IRF3 | HepG2 | 1, 2 TF ChIP-seq |
| ENCFF827VMW | bed narrowPeak | IRF5 | HepG2 | 1, 2 TF ChIP-seq |
| ENCFF308RXD | bed narrowPeak | ISL2 | HepG2 | 1, 2 TF ChIP-seq |
| ENCFF674BRV | bed narrowPeak | JRK | HepG2 | 1, 2 TF ChIP-seq |
| ENCFF079UOJ | bed narrowPeak | JUN | HepG2 | 1, 2 TF ChIP-seq |
| ENCFF562YSV | bed narrowPeak | JUND | HepG2 | 1, 2 TF ChIP-seq |
| ENCFF132RKR | bed narrowPeak | KAT2B | HepG2 | 1, 2 TF ChIP-seq |
| ENCFF847BWZ | bed narrowPeak | KAT7 | HepG2 | 1, 2 TF ChIP-seq |
| ENCFF602VMT | bed narrowPeak | KAT8 | HepG2 | 1, 2 TF ChIP-seq |
| ENCFF439CGG | bed narrowPeak | KDM1A | HepG2 | 1, 2 TF ChIP-seq |
| ENCFF902PJA | bed narrowPeak | KDM2A | HepG2 | 1, 2 TF ChIP-seq |
| ENCFF729ZHC | bed narrowPeak | KDM3A | HepG2 | 1, 2 TF ChIP-seq |
| ENCFF361JRX | bed narrowPeak | KDM4B | HepG2 | 1, 2 TF ChIP-seq |
| ENCFF574LUU | bed narrowPeak | KDM5A | HepG2 | 1, 2 TF ChIP-seq |
| ENCFF595TJC | bed narrowPeak | KDM5B | HepG2 | 1, 2 TF ChIP-seq |
| ENCFF376CUV | bed narrowPeak | KDM6A | HepG2 | 1, 2 TF ChIP-seq |
| ENCFF854ZPK | bed narrowPeak | KIAA2018 | HepG2 | 1, 2 TF ChIP-seq |
| ENCFF443QKY | bed narrowPeak | KLF11 | HepG2 | 1, 2 TF ChIP-seq |
| ENCFF124UUN | bed narrowPeak | KLF12 | HepG2 | 1, 2 TF ChIP-seq |
| ENCFF494FXL | bed narrowPeak | KLF13 | HepG2 | 1, 2 TF ChIP-seq |
| ENCFF257BGF | bed narrowPeak | KLF16 | HepG2 | 1, 2 TF ChIP-seq |
| ENCFF980OTF | bed narrowPeak | KLF6 | HepG2 | 1, 2 TF ChIP-seq |
| ENCFF529UJY | bed narrowPeak | KLF9 | HepG2 | 1, 2 TF ChIP-seq |
| ENCFF644EPI | bed narrowPeak | KMT2A | HepG2 | 1, 2 TF ChIP-seq |
| ENCFF253EID | bed narrowPeak | KMT2B | HepG2 | 1, 2 TF ChIP-seq |
| ENCFF918TGZ | bed narrowPeak | LBX2 | HepG2 | 1, 2 TF ChIP-seq |
| ENCFF453UVV | bed narrowPeak | LCOR | HepG2 | 1, 2 TF ChIP-seq |
| ENCFF606UCO | bed narrowPeak | LCORL | HepG2 | 1, 2 TF ChIP-seq |

| | | | | | |
|---|---|---|---|---|---|
| ENCFF976LBY | bed narrowPeak | MAF1 | HepG2 | 1, 2 | TF ChIP-seq |
| ENCFF005YUC | bed narrowPeak | MAFF | HepG2 | 1, 2 | TF ChIP-seq |
| ENCFF944IDN | bed narrowPeak | MAFG | HepG2 | 1, 2 | TF ChIP-seq |
| ENCFF654RRF | bed narrowPeak | MAFK | HepG2 | 1, 2 | TF ChIP-seq |
| ENCFF241YGM | bed narrowPeak | MATR3 | HepG2 | 1, 2 | TF ChIP-seq |
| ENCFF254ZDA | bed narrowPeak | MAX | HepG2 | 1, 2 | TF ChIP-seq |
| ENCFF341OLL | bed narrowPeak | MAZ | HepG2 | 1, 2 | TF ChIP-seq |
| ENCFF960FKQ | bed narrowPeak | MBD1 | HepG2 | 1, 2 | TF ChIP-seq |
| ENCFF551CCD | bed narrowPeak | MBD4 | HepG2 | 1, 2 | TF ChIP-seq |
| ENCFF493UFO | bed narrowPeak | MED1 | HepG2 | 1, 2 | TF ChIP-seq |
| ENCFF003HBS | bed narrowPeak | MED13 | HepG2 | 1, 2 | TF ChIP-seq |
| ENCFF938PFW | bed narrowPeak | MEF2A | HepG2 | 1, 2 | TF ChIP-seq |
| ENCFF407WPM | bed narrowPeak | MEF2D | HepG2 | 1, 2 | TF ChIP-seq |
| ENCFF546OSE | bed narrowPeak | MEIS1 | HepG2 | 1, 2 | TF ChIP-seq |
| ENCFF647OBG | bed narrowPeak | MEIS2 | HepG2 | 1, 2 | TF ChIP-seq |
| ENCFF713BVZ | bed narrowPeak | MIER2 | HepG2 | 1, 2 | TF ChIP-seq |
| ENCFF186JAO | bed narrowPeak | MIER3 | HepG2 | 1, 2 | TF ChIP-seq |
| ENCFF963OLI | bed narrowPeak | MIXL1 | HepG2 | 1, 2 | TF ChIP-seq |
| ENCFF748SGP | bed narrowPeak | MLLT10 | HepG2 | 1, 2 | TF ChIP-seq |
| ENCFF232WMR | bed narrowPeak | MLX | HepG2 | 1, 2 | TF ChIP-seq |
| ENCFF780JTP | bed narrowPeak | MLXIP | HepG2 | 1, 2 | TF ChIP-seq |
| ENCFF562FMQ | bed narrowPeak | MNT | HepG2 | 1, 2 | TF ChIP-seq |
| ENCFF958FYM | bed narrowPeak | MNX1 | HepG2 | 1, 2 | TF ChIP-seq |
| ENCFF381WCL | bed narrowPeak | MTA1 | HepG2 | 1, 2 | TF ChIP-seq |
| ENCFF093AZT | bed narrowPeak | MTA3 | HepG2 | 1, 2 | TF ChIP-seq |
| ENCFF502QZR | bed narrowPeak | MTF1 | HepG2 | 1, 2 | TF ChIP-seq |
| ENCFF620YHG | bed narrowPeak | MTF2 | HepG2 | 1, 2 | TF ChIP-seq |
| ENCFF835WKW | bed narrowPeak | MXD1 | HepG2 | 1, 2 | TF ChIP-seq |
| ENCFF985DMC | bed narrowPeak | MXD3 | HepG2 | 1, 2 | TF ChIP-seq |
| ENCFF160XKY | bed narrowPeak | MXD4 | HepG2 | 1, 2 | TF ChIP-seq |
| ENCFF599KZZ | bed narrowPeak | MXI1 | HepG2 | 1, 2, 3 | TF ChIP-seq |
| ENCFF904DOZ | bed narrowPeak | MYBL2 | HepG2 | 1, 2 | TF ChIP-seq |
| ENCFF735PKA | bed narrowPeak | MYC | HepG2 | 1, 2, 3 | TF ChIP-seq |
| ENCFF048GXO | bed narrowPeak | MYNN | HepG2 | 1, 2 | TF ChIP-seq |
| ENCFF158PPW | bed narrowPeak | MYRF | HepG2 | 1, 2 | TF ChIP-seq |
| ENCFF287TUA | bed narrowPeak | NAIF1 | HepG2 | 1, 2 | TF ChIP-seq |
| ENCFF516UWH | bed narrowPeak | NBN | HepG2 | 3, 4 | TF ChIP-seq |
| ENCFF399LOS | bed narrowPeak | NCOA1 | HepG2 | 1, 2 | TF ChIP-seq |
| ENCFF312YZB | bed narrowPeak | NCOA5 | HepG2 | 1, 2 | TF ChIP-seq |
| ENCFF878YXY | bed narrowPeak | NCOR1 | HepG2 | 1, 2 | TF ChIP-seq |
| ENCFF069YFV | bed narrowPeak | NFAT5 | HepG2 | 1, 2 | TF ChIP-seq |
| ENCFF482JEG | bed narrowPeak | NFATC3 | HepG2 | 1, 2 | TF ChIP-seq |
| ENCFF306ZFX | bed narrowPeak | NFE2 | HepG2 | 1, 2 | TF ChIP-seq |
| ENCFF620OUT | bed narrowPeak | NFE2L1 | HepG2 | 1, 2 | TF ChIP-seq |
| ENCFF130LVT | bed narrowPeak | NFE2L2 | HepG2 | 1, 2 | TF ChIP-seq |
| ENCFF926VIT | bed narrowPeak | NFIA | HepG2 | 1, 2 | TF ChIP-seq |
| ENCFF127ZZG | bed narrowPeak | NFIB | HepG2 | 1, 2 | TF ChIP-seq |
| ENCFF611FIV | bed narrowPeak | NFIC | HepG2 | 1, 2 | TF ChIP-seq |
| ENCFF278ZNM | bed narrowPeak | NFIL3 | HepG2 | 1, 2 | TF ChIP-seq |
| ENCFF687TJX | bed narrowPeak | NFKB2 | HepG2 | 1, 2 | TF ChIP-seq |
| ENCFF505JMW | bed narrowPeak | NFKBIZ | HepG2 | 1, 2 | TF ChIP-seq |
| ENCFF162TPR | bed narrowPeak | NFRKB | HepG2 | 3, 4 | TF ChIP-seq |

| | | | | | |
|---|---|---|---|---|---|
| ENCFF166QJY | bed narrowPeak | NFYA | HepG2 | 1, 2 | TF ChIP-seq |
| ENCFF257CCO | bed narrowPeak | NFYB | HepG2 | 1, 2 | TF ChIP-seq |
| ENCFF124ITH | bed narrowPeak | NFYC | HepG2 | 1, 2 | TF ChIP-seq |
| ENCFF435AEQ | bed narrowPeak | NKX3-1 | HepG2 | 1, 2 | TF ChIP-seq |
| ENCFF420QKI | bed narrowPeak | NONO | HepG2 | 1, 2 | TF ChIP-seq |
| ENCFF996LQH | bed narrowPeak | NR0B2 | HepG2 | 1, 2 | TF ChIP-seq |
| ENCFF042QSE | bed narrowPeak | NR2C2 | HepG2 | 1, 2 | TF ChIP-seq |
| ENCFF852EYZ | bed narrowPeak | NR2F1 | HepG2 | 1, 2 | TF ChIP-seq |
| ENCFF804MKP | bed narrowPeak | NR2F2 | HepG2 | 1, 2 | TF ChIP-seq |
| ENCFF786BSA | bed narrowPeak | NR2F6 | HepG2 | 1, 2 | TF ChIP-seq |
| ENCFF514SEQ | bed narrowPeak | NR3C1 | HepG2 | 1, 2 | TF ChIP-seq |
| ENCFF510SAT | bed narrowPeak | NR5A1 | HepG2 | 1, 2 | TF ChIP-seq |
| ENCFF982PLG | bed narrowPeak | NRF1 | HepG2 | 1, 2 | TF ChIP-seq |
| ENCFF233BHX | bed narrowPeak | NRL | HepG2 | 1, 2 | TF ChIP-seq |
| ENCFF919ZSN | bed narrowPeak | ONECUT1 | HepG2 | 1, 2 | TF ChIP-seq |
| ENCFF417HLM | bed narrowPeak | ONECUT2 | HepG2 | 1, 2 | TF ChIP-seq |
| ENCFF252CVY | bed narrowPeak | PAF1 | HepG2 | 1, 2 | TF ChIP-seq |
| ENCFF174JHS | bed narrowPeak | PATZ1 | HepG2 | 1, 2 | TF ChIP-seq |
| ENCFF165IYE | bed narrowPeak | PAWR | HepG2 | 1, 2 | TF ChIP-seq |
| ENCFF700GPU | bed narrowPeak | PBX2 | HepG2 | 1, 2 | TF ChIP-seq |
| ENCFF487WAN | bed narrowPeak | PCBP1 | HepG2 | 1, 2 | TF ChIP-seq |
| ENCFF642XRH | bed narrowPeak | PCBP2 | HepG2 | 1, 2 | TF ChIP-seq |
| ENCFF882RPA | bed narrowPeak | PHB2 | HepG2 | 1, 2 | TF ChIP-seq |
| ENCFF635XBX | bed narrowPeak | PHF20 | HepG2 | 1, 2 | TF ChIP-seq |
| ENCFF682RGD | bed narrowPeak | PHF21A | HepG2 | 1, 2 | TF ChIP-seq |
| ENCFF944XYP | bed narrowPeak | PHF5A | HepG2 | 2 | TF ChIP-seq |
| ENCFF558WZE | bed narrowPeak | PHF8 | HepG2 | 1, 2 | TF ChIP-seq |
| ENCFF954EXR | bed narrowPeak | PITX1 | HepG2 | 1, 2 | TF ChIP-seq |
| ENCFF873OHG | bed narrowPeak | PLRG1 | HepG2 | 1, 2 | TF ChIP-seq |
| ENCFF991XSG | bed narrowPeak | PLSCR1 | HepG2 | 1, 2 | TF ChIP-seq |
| ENCFF175FFY | bed narrowPeak | POGZ | HepG2 | 1, 2 | TF ChIP-seq |
| ENCFF043FHJ | bed narrowPeak | POLR2A | HepG2 | 1, 2, 3 | TF ChIP-seq |
| ENCFF652YVO | bed narrowPeak | POLR2AphosphoS2 | HepG2 | 1, 2 | TF ChIP-seq |
| ENCFF520SGM | bed narrowPeak | POLR2AphosphoS5 | HepG2 | 1, 2 | TF ChIP-seq |
| ENCFF551IJP | bed narrowPeak | POLR2G | HepG2 | 1, 2 | TF ChIP-seq |
| ENCFF736PPV | bed narrowPeak | PPARG | HepG2 | 1, 2 | TF ChIP-seq |
| ENCFF668GIP | bed narrowPeak | PRDM10 | HepG2 | 1, 2 | TF ChIP-seq |
| ENCFF486MHG | bed narrowPeak | PRDM15 | HepG2 | 1, 2 | TF ChIP-seq |
| ENCFF666ISN | bed narrowPeak | PRDM4 | HepG2 | 1, 2 | TF ChIP-seq |
| ENCFF249NCB | bed narrowPeak | PREB | HepG2 | 1, 2 | TF ChIP-seq |
| ENCFF908QCS | bed narrowPeak | PRPF4 | HepG2 | 1, 2 | TF ChIP-seq |
| ENCFF875ZPV | bed narrowPeak | PTBP1 | HepG2 | 1, 2 | TF ChIP-seq |
| ENCFF052UCF | bed narrowPeak | RAD21 | HepG2 | 1, 2 | TF ChIP-seq |
| ENCFF859MBC | bed narrowPeak | RAD51 | HepG2 | 1, 2 | TF ChIP-seq |
| ENCFF458WPZ | bed narrowPeak | RARA | HepG2 | 1, 2 | TF ChIP-seq |
| ENCFF909XQR | bed narrowPeak | RARG | HepG2 | 1, 2 | TF ChIP-seq |
| ENCFF080TIX | bed narrowPeak | RBAK | HepG2 | 1, 2 | TF ChIP-seq |
| ENCFF871YRG | bed narrowPeak | RBFOX2 | HepG2 | 1, 2 | TF ChIP-seq |
| ENCFF305WYD | bed narrowPeak | RBM22 | HepG2 | 1, 2 | TF ChIP-seq |
| ENCFF420ALF | bed narrowPeak | RBM39 | HepG2 | 1, 2 | TF ChIP-seq |
| ENCFF153ODZ | bed narrowPeak | RBPJ | HepG2 | 1, 2 | TF ChIP-seq |
| ENCFF987VKU | bed narrowPeak | RCOR1 | HepG2 | 1, 2 | TF ChIP-seq |

| | | | | |
|---|---|---|---|---|
| ENCFF523CUM | bed narrowPeak | RCOR2 | HepG2 | 1, 2 TF ChIP-seq |
| ENCFF089XCL | bed narrowPeak | RELA | HepG2 | 1, 2 TF ChIP-seq |
| ENCFF499ZBA | bed narrowPeak | RERE | HepG2 | 1, 2 TF ChIP-seq |
| ENCFF669XCW | bed narrowPeak | REST | HepG2 | 1, 2 TF ChIP-seq |
| ENCFF788CJF | bed narrowPeak | RFX1 | HepG2 | 2, 3 TF ChIP-seq |
| ENCFF431YWW | bed narrowPeak | RFX3 | HepG2 | 1, 2 TF ChIP-seq |
| ENCFF059GWW | bed narrowPeak | RFX5 | HepG2 | 1, 2 TF ChIP-seq |
| ENCFF125UKX | bed narrowPeak | RFXANK | HepG2 | 1, 2 TF ChIP-seq |
| ENCFF322CXC | bed narrowPeak | RFXAP | HepG2 | 1, 2 TF ChIP-seq |
| ENCFF380SYL | bed narrowPeak | RNF2 | HepG2 | 1, 3 TF ChIP-seq |
| ENCFF926UHP | bed narrowPeak | RNF219 | HepG2 | 1, 2 TF ChIP-seq |
| ENCFF044VTI | bed narrowPeak | RORA | HepG2 | 1, 2 TF ChIP-seq |
| ENCFF625WYP | bed narrowPeak | RREB1 | HepG2 | 1, 2 TF ChIP-seq |
| ENCFF328SRN | bed narrowPeak | RXRA | HepG2 | 1, 2 TF ChIP-seq |
| ENCFF499COS | bed narrowPeak | RXRB | HepG2 | 1, 2 TF ChIP-seq |
| ENCFF710TSK | bed narrowPeak | SAFB2 | HepG2 | 1, 2 TF ChIP-seq |
| ENCFF182FNR | bed narrowPeak | SAP130 | HepG2 | 1, 2 TF ChIP-seq |
| ENCFF004LNW | bed narrowPeak | SATB2 | HepG2 | 1, 2 TF ChIP-seq |
| ENCFF782EKY | bed narrowPeak | SCMH1 | HepG2 | 1, 2 TF ChIP-seq |
| ENCFF615OJH | bed narrowPeak | SETDB1 | HepG2 | 1, 2 TF ChIP-seq |
| ENCFF779CUH | bed narrowPeak | SFPQ | HepG2 | 1, 2 TF ChIP-seq |
| ENCFF635YMI | bed narrowPeak | SIN3A | HepG2 | 1, 2 TF ChIP-seq |
| ENCFF193DQZ | bed narrowPeak | SIN3B | HepG2 | 1, 2 TF ChIP-seq |
| ENCFF069QZX | bed narrowPeak | SIX1 | HepG2 | 1, 2 TF ChIP-seq |
| ENCFF643LFY | bed narrowPeak | SIX4 | HepG2 | 1, 2 TF ChIP-seq |
| ENCFF035ZFO | bed narrowPeak | SKI | HepG2 | 2, 3 TF ChIP-seq |
| ENCFF243JXS | bed narrowPeak | SMAD1 | HepG2 | 1, 2 TF ChIP-seq |
| ENCFF206KPV | bed narrowPeak | SMAD3 | HepG2 | 1, 2 TF ChIP-seq |
| ENCFF541EKJ | bed narrowPeak | SMAD4 | HepG2 | 1, 2 TF ChIP-seq |
| ENCFF339ZPY | bed narrowPeak | SMAD9 | HepG2 | 1, 2 TF ChIP-seq |
| ENCFF150NHK | bed narrowPeak | SMARCC2 | HepG2 | 1, 2 TF ChIP-seq |
| ENCFF210HAA | bed narrowPeak | SMARCE1 | HepG2 | 1, 2 TF ChIP-seq |
| ENCFF035YWE | bed narrowPeak | SMC3 | HepG2 | 1, 2 TF ChIP-seq |
| ENCFF179LFO | bed narrowPeak | SNAI1 | HepG2 | 1, 2 TF ChIP-seq |
| ENCFF754CRG | bed narrowPeak | SNAPC4 | HepG2 | 1, 2 TF ChIP-seq |
| ENCFF455YMK | bed narrowPeak | SNRNP70 | HepG2 | 1, 2 TF ChIP-seq |
| ENCFF517EJH | bed narrowPeak | SOX13 | HepG2 | 1, 2 TF ChIP-seq |
| ENCFF338YFQ | bed narrowPeak | SOX18 | HepG2 | 1, 2 TF ChIP-seq |
| ENCFF703VBH | bed narrowPeak | SOX5 | HepG2 | 1, 2 TF ChIP-seq |
| ENCFF863JMY | bed narrowPeak | SOX6 | HepG2 | 1, 2 TF ChIP-seq |
| ENCFF333SWC | bed narrowPeak | SP1 | HepG2 | 1, 2 TF ChIP-seq |
| ENCFF342ETP | bed narrowPeak | SP110 | HepG2 | 1, 2 TF ChIP-seq |
| ENCFF467WXB | bed narrowPeak | SP140L | HepG2 | 1, 2 TF ChIP-seq |
| ENCFF480YAW | bed narrowPeak | SP2 | HepG2 | 1, 2 TF ChIP-seq |
| ENCFF185BHD | bed narrowPeak | SP5 | HepG2 | 1, 2 TF ChIP-seq |
| ENCFF198OOM | bed narrowPeak | SPEN | HepG2 | 1, 2 TF ChIP-seq |
| ENCFF050QMU | bed narrowPeak | SRF | HepG2 | 1, 2 TF ChIP-seq |
| ENCFF131DDP | bed narrowPeak | SRSF1 | HepG2 | 1, 2 TF ChIP-seq |
| ENCFF122FVR | bed narrowPeak | SRSF4 | HepG2 | 1, 2 TF ChIP-seq |
| ENCFF851DRP | bed narrowPeak | SRSF9 | HepG2 | 1, 2 TF ChIP-seq |
| ENCFF046ODH | bed narrowPeak | SSRP1 | HepG2 | 1, 2 TF ChIP-seq |
| ENCFF477JRZ | bed narrowPeak | STAG1 | HepG2 | 1, 2 TF ChIP-seq |

| | | | | |
|---|---|---|---|---|
| ENCFF370XOM | bed narrowPeak | STAT5B | HepG2 | 1, 2 TF ChIP-seq |
| ENCFF861MUF | bed narrowPeak | STAT6 | HepG2 | 1, 2 TF ChIP-seq |
| ENCFF239LRW | bed narrowPeak | SUZ12 | HepG2 | 1, 2 TF ChIP-seq |
| ENCFF157IIV | bed narrowPeak | SYNCRIP | HepG2 | 1, 2 TF ChIP-seq |
| ENCFF006LKS | bed narrowPeak | TAF1 | HepG2 | 1, 2 TF ChIP-seq |
| ENCFF718RXL | bed narrowPeak | TAF15 | HepG2 | 1, 2 TF ChIP-seq |
| ENCFF696QPP | bed narrowPeak | TARDBP | HepG2 | 1, 2 TF ChIP-seq |
| ENCFF126KGW | bed narrowPeak | TBL1XR1 | HepG2 | 1, 2 TF ChIP-seq |
| ENCFF198GZQ | bed narrowPeak | TBP | HepG2 | 1, 2 TF ChIP-seq |
| ENCFF946LHJ | bed narrowPeak | TBX2 | HepG2 | 1, 2 TF ChIP-seq |
| ENCFF887DUY | bed narrowPeak | TBX3 | HepG2 | 1, 2 TF ChIP-seq |
| ENCFF299JYV | bed narrowPeak | TCF12 | HepG2 | 1, 2 TF ChIP-seq |
| ENCFF902EKZ | bed narrowPeak | TCF3 | HepG2 | 1, 2 TF ChIP-seq |
| ENCFF715QJD | bed narrowPeak | TCF7 | HepG2 | 1, 2 TF ChIP-seq |
| ENCFF726SWQ | bed narrowPeak | TCF7L2 | HepG2 | 1, 2 TF ChIP-seq |
| ENCFF460HAG | bed narrowPeak | TEAD1 | HepG2 | 1, 2 TF ChIP-seq |
| ENCFF680JFD | bed narrowPeak | TEAD2 | HepG2 | 1, 2 TF ChIP-seq |
| ENCFF314CDT | bed narrowPeak | TEAD3 | HepG2 | 1, 2 TF ChIP-seq |
| ENCFF355HRT | bed narrowPeak | TEAD4 | HepG2 | 1, 2 TF ChIP-seq |
| ENCFF475HLY | bed narrowPeak | TEF | HepG2 | 1, 2 TF ChIP-seq |
| ENCFF645QRU | bed narrowPeak | TFAP4 | HepG2 | 1, 2 TF ChIP-seq |
| ENCFF245GRN | bed narrowPeak | TFDP1 | HepG2 | 1, 2 TF ChIP-seq |
| ENCFF340VOQ | bed narrowPeak | TFDP2 | HepG2 | 1, 2 TF ChIP-seq |
| ENCFF277NOY | bed narrowPeak | TFE3 | HepG2 | 1, 2 TF ChIP-seq |
| ENCFF661WFA | bed narrowPeak | TGIF2 | HepG2 | 1, 2 TF ChIP-seq |
| ENCFF889SYG | bed narrowPeak | THAP11 | HepG2 | 1, 2 TF ChIP-seq |
| ENCFF723WCO | bed narrowPeak | THAP8 | HepG2 | 1, 2 TF ChIP-seq |
| ENCFF858SAF | bed narrowPeak | THAP9 | HepG2 | 1, 2 TF ChIP-seq |
| ENCFF737MXG | bed narrowPeak | THRA | HepG2 | 1, 2 TF ChIP-seq |
| ENCFF671MXD | bed narrowPeak | THRB | HepG2 | 1, 2 TF ChIP-seq |
| ENCFF003AMQ | bed narrowPeak | TIGD3 | HepG2 | 1, 2 TF ChIP-seq |
| ENCFF667FDX | bed narrowPeak | TIGD6 | HepG2 | 1, 2 TF ChIP-seq |
| ENCFF167IAP | bed narrowPeak | TOE1 | HepG2 | 1, 2 TF ChIP-seq |
| ENCFF503IOR | bed narrowPeak | TRAFD1 | HepG2 | 1, 2 TF ChIP-seq |
| ENCFF063GDN | bed narrowPeak | TRIM22 | HepG2 | 1, 2 TF ChIP-seq |
| ENCFF278CIM | bed narrowPeak | TSC22D2 | HepG2 | 1, 2 TF ChIP-seq |
| ENCFF034KUO | bed narrowPeak | U2AF1 | HepG2 | 1, 2 TF ChIP-seq |
| ENCFF556EWI | bed narrowPeak | U2AF2 | HepG2 | 1, 2 TF ChIP-seq |
| ENCFF810LSF | bed narrowPeak | UBTF | HepG2 | 1, 2 TF ChIP-seq |
| ENCFF442JPI | bed narrowPeak | USF1 | HepG2 | 1, 2 TF ChIP-seq |
| ENCFF458AZS | bed narrowPeak | USF2 | HepG2 | 1, 2 TF ChIP-seq |
| ENCFF985SRX | bed narrowPeak | WIZ | HepG2 | 1, 2 TF ChIP-seq |
| ENCFF472DLA | bed narrowPeak | XBP1 | HepG2 | 1, 2 TF ChIP-seq |
| ENCFF790ZAQ | bed narrowPeak | XRCC5 | HepG2 | 1, 2 TF ChIP-seq |
| ENCFF332FUE | bed narrowPeak | YBX1 | HepG2 | 1, 2 TF ChIP-seq |
| ENCFF481VPQ | bed narrowPeak | YEATS2 | HepG2 | 1, 2 TF ChIP-seq |
| ENCFF931JDS | bed narrowPeak | YEATS4 | HepG2 | 1, 2 TF ChIP-seq |
| ENCFF177YDT | bed narrowPeak | YY1 | HepG2 | 1, 2 TF ChIP-seq |
| ENCFF021YBN | bed narrowPeak | ZBED4 | HepG2 | 1, 2 TF ChIP-seq |
| ENCFF160DNR | bed narrowPeak | ZBED5 | HepG2 | 1, 2 TF ChIP-seq |
| ENCFF643FQU | bed narrowPeak | ZBTB1 | HepG2 | 1, 2 TF ChIP-seq |
| ENCFF825GVH | bed narrowPeak | ZBTB10 | HepG2 | 1, 2 TF ChIP-seq |

| | | | | |
|---|---|---|---|---|
| ENCFF428ZPS | bed narrowPeak | ZBTB14 | HepG2 | 1, 2 TF ChIP-seq |
| ENCFF819NNH | bed narrowPeak | ZBTB21 | HepG2 | 1, 2 TF ChIP-seq |
| ENCFF148RPN | bed narrowPeak | ZBTB24 | HepG2 | 1, 2 TF ChIP-seq |
| ENCFF330FGQ | bed narrowPeak | ZBTB25 | HepG2 | 1, 2 TF ChIP-seq |
| ENCFF053GKA | bed narrowPeak | ZBTB26 | HepG2 | 1, 2 TF ChIP-seq |
| ENCFF149QHA | bed narrowPeak | ZBTB3 | HepG2 | 1, 2 TF ChIP-seq |
| ENCFF260XMT | bed narrowPeak | ZBTB33 | HepG2 | 1, 2 TF ChIP-seq |
| ENCFF180WMP | bed narrowPeak | ZBTB37 | HepG2 | 1, 2 TF ChIP-seq |
| ENCFF349OOT | bed narrowPeak | ZBTB38 | HepG2 | 1, 2 TF ChIP-seq |
| ENCFF482QHM | bed narrowPeak | ZBTB39 | HepG2 | 1, 2 TF ChIP-seq |
| ENCFF374HXP | bed narrowPeak | ZBTB4 | HepG2 | 1, 2 TF ChIP-seq |
| ENCFF624WDI | bed narrowPeak | ZBTB40 | HepG2 | 1, 2 TF ChIP-seq |
| ENCFF696QTC | bed narrowPeak | ZBTB42 | HepG2 | 1, 2 TF ChIP-seq |
| ENCFF458IBK | bed narrowPeak | ZBTB44 | HepG2 | 1, 2 TF ChIP-seq |
| ENCFF930FAL | bed narrowPeak | ZBTB46 | HepG2 | 1, 2 TF ChIP-seq |
| ENCFF906SCU | bed narrowPeak | ZBTB49 | HepG2 | 1, 2 TF ChIP-seq |
| ENCFF953JQD | bed narrowPeak | ZBTB7A | HepG2 | 1, 2 TF ChIP-seq |
| ENCFF725ZBA | bed narrowPeak | ZBTB7B | HepG2 | 1, 2 TF ChIP-seq |
| ENCFF421QJP | bed narrowPeak | ZBTB8A | HepG2 | 1, 2 TF ChIP-seq |
| ENCFF847EOB | bed narrowPeak | ZC3H4 | HepG2 | 1, 2 TF ChIP-seq |
| ENCFF049ISU | bed narrowPeak | ZC3H8 | HepG2 | 1, 2 TF ChIP-seq |
| ENCFF060IPX | bed narrowPeak | ZEB1 | HepG2 | 1, 2 TF ChIP-seq |
| ENCFF027UTC | bed narrowPeak | ZFP1 | HepG2 | 1, 2 TF ChIP-seq |
| ENCFF800VNW | bed narrowPeak | ZFP14 | HepG2 | 1, 2 TF ChIP-seq |
| ENCFF166GKK | bed narrowPeak | ZFP36 | HepG2 | 2, 3 TF ChIP-seq |
| ENCFF320EKJ | bed narrowPeak | ZFP36L2 | HepG2 | 1, 2 TF ChIP-seq |
| ENCFF743AUT | bed narrowPeak | ZFP37 | HepG2 | 1, 2 TF ChIP-seq |
| ENCFF853YIS | bed narrowPeak | ZFP41 | HepG2 | 1, 2 TF ChIP-seq |
| ENCFF224RFJ | bed narrowPeak | ZFP62 | HepG2 | 1, 2 TF ChIP-seq |
| ENCFF281KMO | bed narrowPeak | ZFP64 | HepG2 | 1, 2 TF ChIP-seq |
| ENCFF408CTC | bed narrowPeak | ZFP82 | HepG2 | 1, 2 TF ChIP-seq |
| ENCFF099SDO | bed narrowPeak | ZFX | HepG2 | 1, 2 TF ChIP-seq |
| ENCFF291LHW | bed narrowPeak | ZFY | HepG2 | 1, 2 TF ChIP-seq |
| ENCFF339GYF | bed narrowPeak | ZGPAT | HepG2 | 1, 2 TF ChIP-seq |
| ENCFF006LDU | bed narrowPeak | ZHX2 | HepG2 | 1, 2 TF ChIP-seq |
| ENCFF438MZI | bed narrowPeak | ZHX3 | HepG2 | 1, 2 TF ChIP-seq |
| ENCFF009LDQ | bed narrowPeak | ZIK1 | HepG2 | 1, 2 TF ChIP-seq |
| ENCFF721NEC | bed narrowPeak | ZKSCAN1 | HepG2 | 1, 3 TF ChIP-seq |
| ENCFF669MLA | bed narrowPeak | ZKSCAN5 | HepG2 | 1, 2 TF ChIP-seq |
| ENCFF024QKK | bed narrowPeak | ZKSCAN8 | HepG2 | 1, 2 TF ChIP-seq |
| ENCFF896EHN | bed narrowPeak | ZMAT5 | HepG2 | 1, 2 TF ChIP-seq |
| ENCFF640TCA | bed narrowPeak | ZMYM3 | HepG2 | 1, 2 TF ChIP-seq |
| ENCFF321MKV | bed narrowPeak | ZNF10 | HepG2 | 1, 2 TF ChIP-seq |
| ENCFF373KUZ | bed narrowPeak | ZNF101 | HepG2 | 1, 2 TF ChIP-seq |
| ENCFF409PJJ | bed narrowPeak | ZNF12 | HepG2 | 1, 2 TF ChIP-seq |
| ENCFF604THW | bed narrowPeak | ZNF121 | HepG2 | 1, 2 TF ChIP-seq |
| ENCFF683CTP | bed narrowPeak | ZNF124 | HepG2 | 1, 2 TF ChIP-seq |
| ENCFF389YCG | bed narrowPeak | ZNF138 | HepG2 | 1, 2 TF ChIP-seq |
| ENCFF868DTR | bed narrowPeak | ZNF142 | HepG2 | 1, 2 TF ChIP-seq |
| ENCFF698ITQ | bed narrowPeak | ZNF143 | HepG2 | 1, 2 TF ChIP-seq |
| ENCFF300AER | bed narrowPeak | ZNF17 | HepG2 | 1, 2 TF ChIP-seq |
| ENCFF405YXO | bed narrowPeak | ZNF180 | HepG2 | 1, 2 TF ChIP-seq |

| | | | | |
|---|---|---|---|---|
| ENCFF627DNU | bed narrowPeak | ZNF20 | HepG2 | 1, 2 TF ChIP-seq |
| ENCFF766KKU | bed narrowPeak | ZNF205 | HepG2 | 1, 2 TF ChIP-seq |
| ENCFF657ZXY | bed narrowPeak | ZNF207 | HepG2 | 1, 2 TF ChIP-seq |
| ENCFF130OKL | bed narrowPeak | ZNF217 | HepG2 | 1, 2 TF ChIP-seq |
| ENCFF708ZFL | bed narrowPeak | ZNF219 | HepG2 | 1, 2 TF ChIP-seq |
| ENCFF772BAU | bed narrowPeak | ZNF224 | HepG2 | 1, 2 TF ChIP-seq |
| ENCFF186RPU | bed narrowPeak | ZNF225 | HepG2 | 1, 2 TF ChIP-seq |
| ENCFF854DNO | bed narrowPeak | ZNF230 | HepG2 | 1, 2 TF ChIP-seq |
| ENCFF119UOZ | bed narrowPeak | ZNF232 | HepG2 | 1, 2 TF ChIP-seq |
| ENCFF858WPR | bed narrowPeak | ZNF24 | HepG2 | 2, 3 TF ChIP-seq |
| ENCFF706SJT | bed narrowPeak | ZNF25 | HepG2 | 1, 2 TF ChIP-seq |
| ENCFF878TPY | bed narrowPeak | ZNF251 | HepG2 | 1, 2 TF ChIP-seq |
| ENCFF612MQA | bed narrowPeak | ZNF260 | HepG2 | 1, 2 TF ChIP-seq |
| ENCFF927JRS | bed narrowPeak | ZNF263 | HepG2 | 1, 2 TF ChIP-seq |
| ENCFF404IUH | bed narrowPeak | ZNF264 | HepG2 | 1, 2 TF ChIP-seq |
| ENCFF823WVL | bed narrowPeak | ZNF274 | HepG2 | 1, 2 TF ChIP-seq |
| ENCFF451VTH | bed narrowPeak | ZNF276 | HepG2 | 1, 2 TF ChIP-seq |
| ENCFF428HNJ | bed narrowPeak | ZNF280B | HepG2 | 1, 2 TF ChIP-seq |
| ENCFF413OMR | bed narrowPeak | ZNF280D | HepG2 | 1, 2 TF ChIP-seq |
| ENCFF964FPZ | bed narrowPeak | ZNF281 | HepG2 | 1, 2 TF ChIP-seq |
| ENCFF482XNG | bed narrowPeak | ZNF282 | HepG2 | 1, 2 TF ChIP-seq |
| ENCFF845MDX | bed narrowPeak | ZNF3 | HepG2 | 1, 2 TF ChIP-seq |
| ENCFF290UUF | bed narrowPeak | ZNF30 | HepG2 | 1, 2 TF ChIP-seq |
| ENCFF865WCB | bed narrowPeak | ZNF317 | HepG2 | 1, 2 TF ChIP-seq |
| ENCFF720DYW | bed narrowPeak | ZNF318 | HepG2 | 1, 2 TF ChIP-seq |
| ENCFF541FQS | bed narrowPeak | ZNF326 | HepG2 | 1, 2 TF ChIP-seq |
| ENCFF731RBX | bed narrowPeak | ZNF329 | HepG2 | 1, 2 TF ChIP-seq |
| ENCFF623DVC | bed narrowPeak | ZNF331 | HepG2 | 1, 2 TF ChIP-seq |
| ENCFF486PIE | bed narrowPeak | ZNF333 | HepG2 | 1, 2 TF ChIP-seq |
| ENCFF034ASW | bed narrowPeak | ZNF335 | HepG2 | 1, 2 TF ChIP-seq |
| ENCFF272IPH | bed narrowPeak | ZNF337 | HepG2 | 1, 2 TF ChIP-seq |
| ENCFF237AST | bed narrowPeak | ZNF33A | HepG2 | 1, 2 TF ChIP-seq |
| ENCFF135VAJ | bed narrowPeak | ZNF33B | HepG2 | 1, 2 TF ChIP-seq |
| ENCFF841BQO | bed narrowPeak | ZNF34 | HepG2 | 1, 2 TF ChIP-seq |
| ENCFF415YWK | bed narrowPeak | ZNF343 | HepG2 | 1, 2 TF ChIP-seq |
| ENCFF518ZKO | bed narrowPeak | ZNF350 | HepG2 | 1, 2 TF ChIP-seq |
| ENCFF361PBY | bed narrowPeak | ZNF362 | HepG2 | 1, 2 TF ChIP-seq |
| ENCFF236CPT | bed narrowPeak | ZNF367 | HepG2 | 1, 2 TF ChIP-seq |
| ENCFF524CCS | bed narrowPeak | ZNF382 | HepG2 | 1, 2 TF ChIP-seq |
| ENCFF442ZRY | bed narrowPeak | ZNF383 | HepG2 | 1, 2 TF ChIP-seq |
| ENCFF308CTG | bed narrowPeak | ZNF384 | HepG2 | 1, 2 TF ChIP-seq |
| ENCFF409BJK | bed narrowPeak | ZNF414 | HepG2 | 1, 2 TF ChIP-seq |
| ENCFF632NSF | bed narrowPeak | ZNF430 | HepG2 | 1, 2 TF ChIP-seq |
| ENCFF748KIJ | bed narrowPeak | ZNF432 | HepG2 | 1, 2 TF ChIP-seq |
| ENCFF624TVT | bed narrowPeak | ZNF44 | HepG2 | 1, 2 TF ChIP-seq |
| ENCFF924VAH | bed narrowPeak | ZNF441 | HepG2 | 1, 2 TF ChIP-seq |
| ENCFF645VCZ | bed narrowPeak | ZNF446 | HepG2 | 1, 2 TF ChIP-seq |
| ENCFF628ZKC | bed narrowPeak | ZNF451 | HepG2 | 1, 2 TF ChIP-seq |
| ENCFF261NJZ | bed narrowPeak | ZNF460 | HepG2 | 1, 2 TF ChIP-seq |
| ENCFF543CJD | bed narrowPeak | ZNF48 | HepG2 | 1, 2 TF ChIP-seq |
| ENCFF616FXO | bed narrowPeak | ZNF483 | HepG2 | 1, 2 TF ChIP-seq |
| ENCFF388IZF | bed narrowPeak | ZNF484 | HepG2 | 1, 2 TF ChIP-seq |

| | | | | | |
|---|---|---|---|---|---|
| ENCFF837BSM | bed narrowPeak | ZNF485 | HepG2 | 1, 2 | TF ChIP-seq |
| ENCFF703APN | bed narrowPeak | ZNF490 | HepG2 | 1, 2 | TF ChIP-seq |
| ENCFF322IVF | bed narrowPeak | ZNF501 | HepG2 | 1, 2 | TF ChIP-seq |
| ENCFF622FUC | bed narrowPeak | ZNF503 | HepG2 | 1, 2 | TF ChIP-seq |
| ENCFF108PPK | bed narrowPeak | ZNF510 | HepG2 | 1, 2 | TF ChIP-seq |
| ENCFF390RLO | bed narrowPeak | ZNF511 | HepG2 | 1, 2 | TF ChIP-seq |
| ENCFF233OIC | bed narrowPeak | ZNF512 | HepG2 | 1, 2 | TF ChIP-seq |
| ENCFF050KYT | bed narrowPeak | ZNF512B | HepG2 | 1, 2 | TF ChIP-seq |
| ENCFF753CVD | bed narrowPeak | ZNF526 | HepG2 | 1, 2 | TF ChIP-seq |
| ENCFF468WZL | bed narrowPeak | ZNF530 | HepG2 | 1, 2 | TF ChIP-seq |
| ENCFF141UZG | bed narrowPeak | ZNF543 | HepG2 | 1, 2 | TF ChIP-seq |
| ENCFF825WJM | bed narrowPeak | ZNF547 | HepG2 | 1, 2 | TF ChIP-seq |
| ENCFF802DLC | bed narrowPeak | ZNF548 | HepG2 | 1, 2 | TF ChIP-seq |
| ENCFF259UJN | bed narrowPeak | ZNF549 | HepG2 | 1, 2 | TF ChIP-seq |
| ENCFF414GKA | bed narrowPeak | ZNF550 | HepG2 | 1, 2 | TF ChIP-seq |
| ENCFF340TNC | bed narrowPeak | ZNF556 | HepG2 | 1, 2 | TF ChIP-seq |
| ENCFF130ZHU | bed narrowPeak | ZNF557 | HepG2 | 1, 2 | TF ChIP-seq |
| ENCFF795VJZ | bed narrowPeak | ZNF569 | HepG2 | 1, 2 | TF ChIP-seq |
| ENCFF259PTV | bed narrowPeak | ZNF570 | HepG2 | 1, 2 | TF ChIP-seq |
| ENCFF841YHN | bed narrowPeak | ZNF572 | HepG2 | 1, 2 | TF ChIP-seq |
| ENCFF550VES | bed narrowPeak | ZNF574 | HepG2 | 1, 2 | TF ChIP-seq |
| ENCFF115ENN | bed narrowPeak | ZNF576 | HepG2 | 1, 2 | TF ChIP-seq |
| ENCFF535BYF | bed narrowPeak | ZNF577 | HepG2 | 1, 2 | TF ChIP-seq |
| ENCFF293UEL | bed narrowPeak | ZNF580 | HepG2 | 1, 2 | TF ChIP-seq |
| ENCFF464WKR | bed narrowPeak | ZNF597 | HepG2 | 1, 2 | TF ChIP-seq |
| ENCFF364DXQ | bed narrowPeak | ZNF598 | HepG2 | 1, 2 | TF ChIP-seq |
| ENCFF754OSO | bed narrowPeak | ZNF607 | HepG2 | 1, 2 | TF ChIP-seq |
| ENCFF778FIE | bed narrowPeak | ZNF608 | HepG2 | 1, 2 | TF ChIP-seq |
| ENCFF667MEL | bed narrowPeak | ZNF609 | HepG2 | 1, 2 | TF ChIP-seq |
| ENCFF241JAR | bed narrowPeak | ZNF614 | HepG2 | 1, 2 | TF ChIP-seq |
| ENCFF618JPG | bed narrowPeak | ZNF615 | HepG2 | 1, 2 | TF ChIP-seq |
| ENCFF255QEY | bed narrowPeak | ZNF616 | HepG2 | 1, 2 | TF ChIP-seq |
| ENCFF898ZHK | bed narrowPeak | ZNF619 | HepG2 | 1, 2 | TF ChIP-seq |
| ENCFF859VUR | bed narrowPeak | ZNF639 | HepG2 | 1, 2 | TF ChIP-seq |
| ENCFF623COM | bed narrowPeak | ZNF644 | HepG2 | 1, 2 | TF ChIP-seq |
| ENCFF496HIK | bed narrowPeak | ZNF646 | HepG2 | 1, 2 | TF ChIP-seq |
| ENCFF761HIZ | bed narrowPeak | ZNF652 | HepG2 | 1, 2 | TF ChIP-seq |
| ENCFF215UIS | bed narrowPeak | ZNF660 | HepG2 | 1, 2 | TF ChIP-seq |
| ENCFF450BWT | bed narrowPeak | ZNF674 | HepG2 | 1, 2 | TF ChIP-seq |
| ENCFF678IWM | bed narrowPeak | ZNF678 | HepG2 | 1, 2 | TF ChIP-seq |
| ENCFF581GZR | bed narrowPeak | ZNF687 | HepG2 | 1, 2 | TF ChIP-seq |
| ENCFF296WYX | bed narrowPeak | ZNF691 | HepG2 | 1, 2 | TF ChIP-seq |
| ENCFF995ZSA | bed narrowPeak | ZNF7 | HepG2 | 1, 2 | TF ChIP-seq |
| ENCFF898LPA | bed narrowPeak | ZNF703 | HepG2 | 1, 2 | TF ChIP-seq |
| ENCFF497RWP | bed narrowPeak | ZNF707 | HepG2 | 1, 2 | TF ChIP-seq |
| ENCFF371EUY | bed narrowPeak | ZNF709 | HepG2 | 1, 2 | TF ChIP-seq |
| ENCFF216JWP | bed narrowPeak | ZNF710 | HepG2 | 1, 2 | TF ChIP-seq |
| ENCFF262CSX | bed narrowPeak | ZNF713 | HepG2 | 1, 2 | TF ChIP-seq |
| ENCFF879KUL | bed narrowPeak | ZNF720 | HepG2 | 1, 2 | TF ChIP-seq |
| ENCFF024LQZ | bed narrowPeak | ZNF737 | HepG2 | 1, 2 | TF ChIP-seq |
| ENCFF854KWX | bed narrowPeak | ZNF740 | HepG2 | 1, 2 | TF ChIP-seq |
| ENCFF042SUM | bed narrowPeak | ZNF766 | HepG2 | 1, 2 | TF ChIP-seq |

| | | | | | |
|---|---|---|---|---|---|
| ENCFF079KQG | bed narrowPeak | ZNF768 | HepG2 | 1, 2 | TF ChIP-seq |
| ENCFF514IUU | bed narrowPeak | ZNF770 | HepG2 | 1, 2 | TF ChIP-seq |
| ENCFF874HXM | bed narrowPeak | ZNF772 | HepG2 | 1, 2 | TF ChIP-seq |
| ENCFF906WDG | bed narrowPeak | ZNF773 | HepG2 | 1, 2 | TF ChIP-seq |
| ENCFF230KZH | bed narrowPeak | ZNF775 | HepG2 | 1, 2 | TF ChIP-seq |
| ENCFF396ZSB | bed narrowPeak | ZNF776 | HepG2 | 1, 2 | TF ChIP-seq |
| ENCFF920PLY | bed narrowPeak | ZNF777 | HepG2 | 1, 2 | TF ChIP-seq |
| ENCFF078ATH | bed narrowPeak | ZNF778 | HepG2 | 1, 2 | TF ChIP-seq |
| ENCFF482XKY | bed narrowPeak | ZNF781 | HepG2 | 1, 2 | TF ChIP-seq |
| ENCFF462RVE | bed narrowPeak | ZNF782 | HepG2 | 1, 2 | TF ChIP-seq |
| ENCFF645BZY | bed narrowPeak | ZNF784 | HepG2 | 1, 2 | TF ChIP-seq |
| ENCFF663KMO | bed narrowPeak | ZNF788 | HepG2 | 1, 2 | TF ChIP-seq |
| ENCFF253CWZ | bed narrowPeak | ZNF792 | HepG2 | 1, 2 | TF ChIP-seq |
| ENCFF618NTX | bed narrowPeak | ZNF800 | HepG2 | 1, 2 | TF ChIP-seq |
| ENCFF621IOC | bed narrowPeak | ZNF83 | HepG2 | 1, 2 | TF ChIP-seq |
| ENCFF332MPQ | bed narrowPeak | ZNF839 | HepG2 | 1, 2 | TF ChIP-seq |
| ENCFF060MHX | bed narrowPeak | ZNF865 | HepG2 | 1, 2 | TF ChIP-seq |
| ENCFF189PKJ | bed narrowPeak | ZNF883 | HepG2 | 1, 2 | TF ChIP-seq |
| ENCFF903IZG | bed narrowPeak | ZNF891 | HepG2 | 1, 2 | TF ChIP-seq |
| ENCFF594DAV | bed narrowPeak | ZSCAN12 | HepG2 | 1, 2 | TF ChIP-seq |
| ENCFF505AEY | bed narrowPeak | ZSCAN20 | HepG2 | 1, 2 | TF ChIP-seq |
| ENCFF904LYG | bed narrowPeak | ZSCAN22 | HepG2 | 1, 2 | TF ChIP-seq |
| ENCFF560NWT | bed narrowPeak | ZSCAN30 | HepG2 | 1, 2 | TF ChIP-seq |
| ENCFF810TWJ | bed narrowPeak | ZSCAN31 | HepG2 | 1, 2 | TF ChIP-seq |
| ENCFF673YDK | bed narrowPeak | ZSCAN9 | HepG2 | 1, 2 | TF ChIP-seq |
| ENCFF594JNI | bed narrowPeak | ZUFSP | HepG2 | 1, 2 | TF ChIP-seq |
| ENCFF044XYZ | bed narrowPeak | ZXDC | HepG2 | 1, 2 | TF ChIP-seq |
| ENCFF791UUM | bed narrowPeak | ZZZ3 | HepG2 | 1, 2 | TF ChIP-seq |
| ENCFF739AJO | bed narrowPeak | ADNP | K562 | 1, 2 | TF ChIP-seq |
| ENCFF489SKQ | bed narrowPeak | AFF1 | K562 | 1, 2 | TF ChIP-seq |
| ENCFF100VYA | bed narrowPeak | AGO1 | K562 | 1, 2 | TF ChIP-seq |
| ENCFF089PKE | bed narrowPeak | ARHGAP35 | K562 | 1, 2 | TF ChIP-seq |
| ENCFF879NTL | bed narrowPeak | ARID1B | K562 | 1, 2 | TF ChIP-seq |
| ENCFF344MKI | bed narrowPeak | ARID2 | K562 | 1, 2 | TF ChIP-seq |
| ENCFF891OQP | bed narrowPeak | ARID3A | K562 | 1, 2 | TF ChIP-seq |
| ENCFF270TSN | bed narrowPeak | ARID3B | K562 | 1, 2 | TF ChIP-seq |
| ENCFF655EFA | bed narrowPeak | ARNT | K562 | 1, 2 | TF ChIP-seq |
| ENCFF958YSG | bed narrowPeak | ASH1L | K562 | 1, 2 | TF ChIP-seq |
| ENCFF206BGR | bed narrowPeak | ATF1 | K562 | 1, 2 | TF ChIP-seq |
| ENCFF803FHN | bed narrowPeak | ATF2 | K562 | 1, 2 | TF ChIP-seq |
| ENCFF718VHT | bed narrowPeak | ATF3 | K562 | 1, 2 | TF ChIP-seq |
| ENCFF182MNO | bed narrowPeak | ATF4 | K562 | 1, 2 | TF ChIP-seq |
| ENCFF863ZFH | bed narrowPeak | ATF6 | K562 | 1, 2 | TF ChIP-seq |
| ENCFF371SJR | bed narrowPeak | ATF7 | K562 | 1, 2 | TF ChIP-seq |
| ENCFF026BDA | bed narrowPeak | BACH1 | K562 | 1, 2 | TF ChIP-seq |
| ENCFF941EDY | bed narrowPeak | BCL6 | K562 | 1, 2 | TF ChIP-seq |
| ENCFF070ZTX | bed narrowPeak | BCLAF1 | K562 | 1, 2 | TF ChIP-seq |
| ENCFF186JKG | bed narrowPeak | BCOR | K562 | 1, 2 | TF ChIP-seq |
| ENCFF284FTY | bed narrowPeak | BDP1 | K562 | 1, 2, 3 | TF ChIP-seq |
| ENCFF154IVU | bed narrowPeak | BHLHE40 | K562 | 1, 2 | TF ChIP-seq |
| ENCFF352DRR | bed narrowPeak | BMI1 | K562 | 1, 2 | TF ChIP-seq |
| ENCFF620FIH | bed narrowPeak | BRCA1 | K562 | 1, 2 | TF ChIP-seq |

| | | | | | |
|---|---|---|---|---|---|
| ENCFF130JVF | bed narrowPeak | BRD4 | K562 | 1, 2 | TF ChIP-seq |
| ENCFF411RMT | bed narrowPeak | BRD9 | K562 | 1, 2 | TF ChIP-seq |
| ENCFF577LSK | bed narrowPeak | BRF1 | K562 | 1, 2, 3 | TF ChIP-seq |
| ENCFF944LWS | bed narrowPeak | BRF2 | K562 | 1, 2, 3 | TF ChIP-seq |
| ENCFF104MXG | bed narrowPeak | C11orf30 | K562 | 1, 2 | TF ChIP-seq |
| ENCFF419PEK | bed narrowPeak | CBFA2T2 | K562 | 1, 2 | TF ChIP-seq |
| ENCFF153IFH | bed narrowPeak | CBFA2T3 | K562 | 1, 2 | TF ChIP-seq |
| ENCFF802NHC | bed narrowPeak | CBFB | K562 | 1, 2 | TF ChIP-seq |
| ENCFF163FLA | bed narrowPeak | CBX1 | K562 | 1, 2 | TF ChIP-seq |
| ENCFF258XBJ | bed narrowPeak | CBX2 | K562 | 1 | TF ChIP-seq |
| ENCFF068OEJ | bed narrowPeak | CBX3 | K562 | 1 | TF ChIP-seq |
| ENCFF403TAE | bed narrowPeak | CBX5 | K562 | 1, 2 | TF ChIP-seq |
| ENCFF210GJE | bed narrowPeak | CBX8 | K562 | 1, 2 | TF ChIP-seq |
| ENCFF180TUM | bed narrowPeak | CC2D1A | K562 | 1, 2 | TF ChIP-seq |
| ENCFF704PGT | bed narrowPeak | CCAR2 | K562 | 1, 2 | TF ChIP-seq |
| ENCFF642ZYO | bed narrowPeak | CCNT2 | K562 | 1, 2 | TF ChIP-seq |
| ENCFF384ALH | bed narrowPeak | CDC5L | K562 | 1, 2 | TF ChIP-seq |
| ENCFF882ARK | bed narrowPeak | CEBPB | K562 | 1, 2 | TF ChIP-seq |
| ENCFF797MRW | bed narrowPeak | CEBPG | K562 | 1, 2 | TF ChIP-seq |
| ENCFF588GNU | bed narrowPeak | CEBPZ | K562 | 1, 4 | TF ChIP-seq |
| ENCFF771XZZ | bed narrowPeak | CGGBP1 | K562 | 1, 2 | TF ChIP-seq |
| ENCFF646MEF | bed narrowPeak | CHAMP1 | K562 | 1, 2 | TF ChIP-seq |
| ENCFF408NUX | bed narrowPeak | CHD1 | K562 | 1 | TF ChIP-seq |
| ENCFF947AEO | bed narrowPeak | CHD2 | K562 | 1, 2 | TF ChIP-seq |
| ENCFF985QBS | bed narrowPeak | CHD4 | K562 | 1, 2, 3 | TF ChIP-seq |
| ENCFF722UJW | bed narrowPeak | CHD7 | K562 | 1 | TF ChIP-seq |
| ENCFF648UJW | bed narrowPeak | CLOCK | K562 | 1, 2 | TF ChIP-seq |
| ENCFF552EBC | bed narrowPeak | COPS2 | K562 | 1, 2 | TF ChIP-seq |
| ENCFF193LLN | bed narrowPeak | CREB1 | K562 | 1, 2 | TF ChIP-seq |
| ENCFF606EUI | bed narrowPeak | CREB3 | K562 | 1, 2 | TF ChIP-seq |
| ENCFF566HGU | bed narrowPeak | CREB3L1 | K562 | 1, 2 | TF ChIP-seq |
| ENCFF875JMR | bed narrowPeak | CREB5 | K562 | 1, 2 | TF ChIP-seq |
| ENCFF532VPN | bed narrowPeak | CREBBP | K562 | 1 | TF ChIP-seq |
| ENCFF021XJN | bed narrowPeak | CREM | K562 | 1, 2 | TF ChIP-seq |
| ENCFF351DBE | bed narrowPeak | CSDE1 | K562 | 1, 2 | TF ChIP-seq |
| ENCFF349UTF | bed narrowPeak | CTBP1 | K562 | 1, 2 | TF ChIP-seq |
| ENCFF221SKA | bed narrowPeak | CTCF | K562 | 1, 2, 3 | TF ChIP-seq |
| ENCFF630YVJ | bed narrowPeak | CTCFL | K562 | 1, 2 | TF ChIP-seq |
| ENCFF136KLM | bed narrowPeak | CUX1 | K562 | 1, 2 | TF ChIP-seq |
| ENCFF870LJV | bed narrowPeak | DACH1 | K562 | 1, 2 | TF ChIP-seq |
| ENCFF869RFC | bed narrowPeak | DDIT3 | K562 | 1, 2 | TF ChIP-seq |
| ENCFF536LKB | bed narrowPeak | DDX20 | K562 | 1, 2 | TF ChIP-seq |
| ENCFF030AXK | bed narrowPeak | DEAF1 | K562 | 1, 2 | TF ChIP-seq |
| ENCFF108JFW | bed narrowPeak | DIDO1 | K562 | 1, 2 | TF ChIP-seq |
| ENCFF832DBU | bed narrowPeak | DLX4 | K562 | 1, 2 | TF ChIP-seq |
| ENCFF549TVW | bed narrowPeak | DNMT1 | K562 | 1, 2 | TF ChIP-seq |
| ENCFF542HJS | bed narrowPeak | DPF2 | K562 | 1, 2 | TF ChIP-seq |
| ENCFF104VFF | bed narrowPeak | E2F1 | K562 | 1, 2 | TF ChIP-seq |
| ENCFF710UAZ | bed narrowPeak | E2F3 | K562 | 1, 2 | TF ChIP-seq |
| ENCFF225TLP | bed narrowPeak | E2F4 | K562 | 1, 2 | TF ChIP-seq |
| ENCFF468VJV | bed narrowPeak | E2F5 | K562 | 1, 2 | TF ChIP-seq |
| ENCFF831NDQ | bed narrowPeak | E2F6 | K562 | 1, 2 | TF ChIP-seq |

| | | | | | |
|---|---|---|---|---|---|
| ENCFF013EHI | bed narrowPeak | E2F7 | K562 | 1, 2 | TF ChIP-seq |
| ENCFF171WWF | bed narrowPeak | E2F8 | K562 | 1, 2 | TF ChIP-seq |
| ENCFF683WRK | bed narrowPeak | E4F1 | K562 | 1, 2 | TF ChIP-seq |
| ENCFF566PRZ | bed narrowPeak | EGR1 | K562 | 1, 2 | TF ChIP-seq |
| ENCFF682XPD | bed narrowPeak | EHMT2 | K562 | 1, 2 | TF ChIP-seq |
| ENCFF133TSU | bed narrowPeak | ELF1 | K562 | 1, 2 | TF ChIP-seq |
| ENCFF695PDY | bed narrowPeak | ELF2 | K562 | 1, 2 | TF ChIP-seq |
| ENCFF539SXG | bed narrowPeak | ELF4 | K562 | 1, 2 | TF ChIP-seq |
| ENCFF119SCQ | bed narrowPeak | ELK1 | K562 | 1, 2 | TF ChIP-seq |
| ENCFF198NGY | bed narrowPeak | ELK3 | K562 | 1, 2 | TF ChIP-seq |
| ENCFF821MKR | bed narrowPeak | EP300 | K562 | 1, 2 | TF ChIP-seq |
| ENCFF167CQF | bed narrowPeak | EP400 | K562 | 1, 2 | TF ChIP-seq |
| ENCFF462ZIG | bed narrowPeak | ERF | K562 | 1, 2 | TF ChIP-seq |
| ENCFF592GWM | bed narrowPeak | ESRRA | K562 | 1, 2 | TF ChIP-seq |
| ENCFF106ECV | bed narrowPeak | ESRRB | K562 | 1, 2 | TF ChIP-seq |
| ENCFF461PRP | bed narrowPeak | ETS1 | K562 | 1, 2 | TF ChIP-seq |
| ENCFF687OMH | bed narrowPeak | ETS2 | K562 | 1, 2 | TF ChIP-seq |
| ENCFF804KSB | bed narrowPeak | ETV1 | K562 | 1, 2 | TF ChIP-seq |
| ENCFF604LXR | bed narrowPeak | ETV5 | K562 | 1, 2 | TF ChIP-seq |
| ENCFF426GSY | bed narrowPeak | ETV6 | K562 | 1, 2 | TF ChIP-seq |
| ENCFF560CYG | bed narrowPeak | EWSR1 | K562 | 1, 2 | TF ChIP-seq |
| ENCFF804RVA | bed narrowPeak | EZH2 | K562 | 1, 2 | TF ChIP-seq |
| ENCFF285TMA | bed narrowPeak | FIP1L1 | K562 | 1, 2 | TF ChIP-seq |
| ENCFF258PLH | bed narrowPeak | FOS | K562 | 1, 2, 3 | TF ChIP-seq |
| ENCFF004HXL | bed narrowPeak | FOSL1 | K562 | 1, 2 | TF ChIP-seq |
| ENCFF765NAN | bed narrowPeak | FOXA1 | K562 | 1, 2 | TF ChIP-seq |
| ENCFF957CVJ | bed narrowPeak | FOXJ2 | K562 | 1, 2 | TF ChIP-seq |
| ENCFF567CPM | bed narrowPeak | FOXJ3 | K562 | 1, 2 | TF ChIP-seq |
| ENCFF066CWG | bed narrowPeak | FOXK2 | K562 | 1, 2 | TF ChIP-seq |
| ENCFF778PWE | bed narrowPeak | FOXM1 | K562 | 1, 2 | TF ChIP-seq |
| ENCFF750AGR | bed narrowPeak | FOXO4 | K562 | 1, 2 | TF ChIP-seq |
| ENCFF688ARM | bed narrowPeak | FUS | K562 | 1, 2 | TF ChIP-seq |
| ENCFF489EME | bed narrowPeak | GABPA | K562 | 1, 2 | TF ChIP-seq |
| ENCFF700DXR | bed narrowPeak | GABPB1 | K562 | 1, 2 | TF ChIP-seq |
| ENCFF657CTC | bed narrowPeak | GATA1 | K562 | 1, 2 | TF ChIP-seq |
| ENCFF165ZEP | bed narrowPeak | GATA2 | K562 | 1, 2 | TF ChIP-seq |
| ENCFF950ZWP | bed narrowPeak | GATAD2A | K562 | 1, 2 | TF ChIP-seq |
| ENCFF569CMJ | bed narrowPeak | GATAD2B | K562 | 1, 2 | TF ChIP-seq |
| ENCFF365ETH | bed narrowPeak | GMEB1 | K562 | 1, 2 | TF ChIP-seq |
| ENCFF839QEL | bed narrowPeak | GTF2A2 | K562 | 1, 2 | TF ChIP-seq |
| ENCFF970ULG | bed narrowPeak | GTF2B | K562 | 1, 2 | TF ChIP-seq |
| ENCFF564KIU | bed narrowPeak | GTF2E2 | K562 | 1, 2 | TF ChIP-seq |
| ENCFF843UHP | bed narrowPeak | GTF2F1 | K562 | 1, 2 | TF ChIP-seq |
| ENCFF866OZW | bed narrowPeak | GTF2I | K562 | 1, 2 | TF ChIP-seq |
| ENCFF496PIQ | bed narrowPeak | GTF3C2 | K562 | 1, 2, 3 | TF ChIP-seq |
| ENCFF139FMU | bed narrowPeak | HCFC1 | K562 | 1, 2 | TF ChIP-seq |
| ENCFF188TBM | bed narrowPeak | HDAC1 | K562 | 1, 2 | TF ChIP-seq |
| ENCFF519RWJ | bed narrowPeak | HDAC2 | K562 | 1, 2 | TF ChIP-seq |
| ENCFF742LSD | bed narrowPeak | HDAC3 | K562 | 3, 4 | TF ChIP-seq |
| ENCFF295GBP | bed narrowPeak | HDAC6 | K562 | 1, 2 | TF ChIP-seq |
| ENCFF349GBE | bed narrowPeak | HDAC8 | K562 | 1, 2 | TF ChIP-seq |
| ENCFF297WMY | bed narrowPeak | HDGF | K562 | 1, 2 | TF ChIP-seq |

| | | | | | |
|---|---|---|---|---|---|
| ENCFF010OOE | bed narrowPeak | HES1 | K562 | 1, 2 | TF ChIP-seq |
| ENCFF180KAV | bed narrowPeak | HEY1 | K562 | 1, 2 | TF ChIP-seq |
| ENCFF558VAL | bed narrowPeak | HINFP | K562 | 1, 2 | TF ChIP-seq |
| ENCFF449JMB | bed narrowPeak | HIVEP1 | K562 | 1, 2 | TF ChIP-seq |
| ENCFF689LNQ | bed narrowPeak | HLTF | K562 | 1, 2 | TF ChIP-seq |
| ENCFF718DFX | bed narrowPeak | HMBOX1 | K562 | 1, 2 | TF ChIP-seq |
| ENCFF099OSI | bed narrowPeak | HMG20B | K562 | 1, 2 | TF ChIP-seq |
| ENCFF615AKB | bed narrowPeak | HMGN3 | K562 | 1, 2 | TF ChIP-seq |
| ENCFF844QFF | bed narrowPeak | HNRNPH1 | K562 | 1, 2 | TF ChIP-seq |
| ENCFF984QUV | bed narrowPeak | HNRNPK | K562 | 1, 2 | TF ChIP-seq |
| ENCFF984ESZ | bed narrowPeak | HNRNPL | K562 | 1, 2 | TF ChIP-seq |
| ENCFF662WPN | bed narrowPeak | HNRNPLL | K562 | 1, 2 | TF ChIP-seq |
| ENCFF991ZSC | bed narrowPeak | HNRNPUL1 | K562 | 1, 2 | TF ChIP-seq |
| ENCFF601LMD | bed narrowPeak | HOMEZ | K562 | 1, 2 | TF ChIP-seq |
| ENCFF794SCQ | bed narrowPeak | ID3 | K562 | 1, 2 | TF ChIP-seq |
| ENCFF994OQH | bed narrowPeak | IKZF1 | K562 | 1, 2 | TF ChIP-seq |
| ENCFF368AAQ | bed narrowPeak | ILF3 | K562 | 1, 4 | TF ChIP-seq |
| ENCFF406XWT | bed narrowPeak | ILK | K562 | 1, 2 | TF ChIP-seq |
| ENCFF203LRV | bed narrowPeak | IRF1 | K562 | 1, 2 | TF ChIP-seq |
| ENCFF886EVL | bed narrowPeak | IRF2 | K562 | 1, 3 | TF ChIP-seq |
| ENCFF863WCN | bed narrowPeak | IRF9 | K562 | 1, 2 | TF ChIP-seq |
| ENCFF589QXC | bed narrowPeak | JUN | K562 | 1, 2 | TF ChIP-seq |
| ENCFF362UGH | bed narrowPeak | JUNB | K562 | 1, 2 | TF ChIP-seq |
| ENCFF273KIA | bed narrowPeak | JUND | K562 | 1, 2, 3 | TF ChIP-seq |
| ENCFF349VSP | bed narrowPeak | KAT2B | K562 | 1 | TF ChIP-seq |
| ENCFF207ZEK | bed narrowPeak | KAT8 | K562 | 1, 2 | TF ChIP-seq |
| ENCFF054XCG | bed narrowPeak | KDM1A | K562 | 1, 2, 3 | TF ChIP-seq |
| ENCFF955AOD | bed narrowPeak | KDM4B | K562 | 1, 2 | TF ChIP-seq |
| ENCFF807FNB | bed narrowPeak | KDM5B | K562 | 1, 2 | TF ChIP-seq |
| ENCFF317QKH | bed narrowPeak | KHSRP | K562 | 3, 4 | TF ChIP-seq |
| ENCFF287GDT | bed narrowPeak | KLF1 | K562 | 1, 2 | TF ChIP-seq |
| ENCFF142ZTD | bed narrowPeak | KLF10 | K562 | 1, 2 | TF ChIP-seq |
| ENCFF381GEK | bed narrowPeak | KLF13 | K562 | 1, 2 | TF ChIP-seq |
| ENCFF488OTN | bed narrowPeak | KLF16 | K562 | 1, 2 | TF ChIP-seq |
| ENCFF563ZJM | bed narrowPeak | KLF6 | K562 | 1, 2 | TF ChIP-seq |
| ENCFF423LPW | bed narrowPeak | L3MBTL2 | K562 | 1, 2 | TF ChIP-seq |
| ENCFF668ZTJ | bed narrowPeak | LARP7 | K562 | 1, 2 | TF ChIP-seq |
| ENCFF134HQP | bed narrowPeak | LEF1 | K562 | 3, 4 | TF ChIP-seq |
| ENCFF119AHD | bed narrowPeak | MAFF | K562 | 1, 2 | TF ChIP-seq |
| ENCFF393CCU | bed narrowPeak | MAFG | K562 | 1, 2 | TF ChIP-seq |
| ENCFF439TJM | bed narrowPeak | MAFK | K562 | 1, 2 | TF ChIP-seq |
| ENCFF822FKQ | bed narrowPeak | MAX | K562 | 1, 2 | TF ChIP-seq |
| ENCFF837NNR | bed narrowPeak | MAZ | K562 | 1, 2 | TF ChIP-seq |
| ENCFF617QSK | bed narrowPeak | MBD2 | K562 | 1, 3 | TF ChIP-seq |
| ENCFF571REC | bed narrowPeak | MCM2 | K562 | 1, 2 | TF ChIP-seq |
| ENCFF672PYP | bed narrowPeak | MCM3 | K562 | 1, 2 | TF ChIP-seq |
| ENCFF658SJY | bed narrowPeak | MCM5 | K562 | 1, 2 | TF ChIP-seq |
| ENCFF914ELA | bed narrowPeak | MCM7 | K562 | 1, 2 | TF ChIP-seq |
| ENCFF031NTF | bed narrowPeak | MEF2A | K562 | 1, 2 | TF ChIP-seq |
| ENCFF436DME | bed narrowPeak | MEF2D | K562 | 1, 2 | TF ChIP-seq |
| ENCFF937UEE | bed narrowPeak | MEIS2 | K562 | 1, 2 | TF ChIP-seq |
| ENCFF524ZER | bed narrowPeak | MGA | K562 | 1, 2 | TF ChIP-seq |

| | | | | | |
|---|---|---|---|---|---|
| ENCFF163YZB | bed narrowPeak | MIER1 | K562 | 1, 2 | TF ChIP-seq |
| ENCFF262TMM | bed narrowPeak | MITF | K562 | 3, 4 | TF ChIP-seq |
| ENCFF010AIG | bed narrowPeak | MLLT1 | K562 | 1, 2 | TF ChIP-seq |
| ENCFF926CRV | bed narrowPeak | MNT | K562 | 1, 2 | TF ChIP-seq |
| ENCFF801KEW | bed narrowPeak | MTA1 | K562 | 1, 2 | TF ChIP-seq |
| ENCFF080IBW | bed narrowPeak | MTA2 | K562 | 1, 2 | TF ChIP-seq |
| ENCFF459XLR | bed narrowPeak | MTA3 | K562 | 2, 3 | TF ChIP-seq |
| ENCFF068IGH | bed narrowPeak | MXI1 | K562 | 1, 2 | TF ChIP-seq |
| ENCFF637PFI | bed narrowPeak | MYBL2 | K562 | 1, 2 | TF ChIP-seq |
| ENCFF566CTX | bed narrowPeak | MYC | K562 | 1, 2, 3 | TF ChIP-seq |
| ENCFF516ZEQ | bed narrowPeak | MYNN | K562 | 1, 2 | TF ChIP-seq |
| ENCFF728KKP | bed narrowPeak | NBN | K562 | 1, 2 | TF ChIP-seq |
| ENCFF474QDS | bed narrowPeak | NCOA1 | K562 | 1, 2 | TF ChIP-seq |
| ENCFF584SNZ | bed narrowPeak | NCOA2 | K562 | 1, 2 | TF ChIP-seq |
| ENCFF749HKV | bed narrowPeak | NCOA4 | K562 | 1, 2 | TF ChIP-seq |
| ENCFF438BWN | bed narrowPeak | NCOA6 | K562 | 1, 2 | TF ChIP-seq |
| ENCFF816AEF | bed narrowPeak | NCOR1 | K562 | 1 | TF ChIP-seq |
| ENCFF265KKE | bed narrowPeak | NELFE | K562 | 1, 2 | TF ChIP-seq |
| ENCFF625QHR | bed narrowPeak | NEUROD1 | K562 | 1, 2 | TF ChIP-seq |
| ENCFF430JFH | bed narrowPeak | NFATC3 | K562 | 1, 2 | TF ChIP-seq |
| ENCFF286HTI | bed narrowPeak | NFE2 | K562 | 1, 2 | TF ChIP-seq |
| ENCFF462UJN | bed narrowPeak | NFE2L1 | K562 | 1, 2 | TF ChIP-seq |
| ENCFF169MAE | bed narrowPeak | NFIC | K562 | 1, 2 | TF ChIP-seq |
| ENCFF726LLI | bed narrowPeak | NFIX | K562 | 1, 2 | TF ChIP-seq |
| ENCFF158FUG | bed narrowPeak | NFRKB | K562 | 1, 2 | TF ChIP-seq |
| ENCFF201WFM | bed narrowPeak | NFXL1 | K562 | 1, 2 | TF ChIP-seq |
| ENCFF908HSL | bed narrowPeak | NFYA | K562 | 1, 2 | TF ChIP-seq |
| ENCFF718ZFY | bed narrowPeak | NFYB | K562 | 1, 2 | TF ChIP-seq |
| ENCFF520QSR | bed narrowPeak | NKRF | K562 | 1, 2 | TF ChIP-seq |
| ENCFF211TTD | bed narrowPeak | NONO | K562 | 1, 2 | TF ChIP-seq |
| ENCFF305OOU | bed narrowPeak | NR0B1 | K562 | 1, 2 | TF ChIP-seq |
| ENCFF798KXN | bed narrowPeak | NR1H2 | K562 | 1, 2 | TF ChIP-seq |
| ENCFF431GJK | bed narrowPeak | NR2C1 | K562 | 1, 2 | TF ChIP-seq |
| ENCFF789AXP | bed narrowPeak | NR2C2 | K562 | 1, 2 | TF ChIP-seq |
| ENCFF363IQN | bed narrowPeak | NR2F1 | K562 | 1, 2 | TF ChIP-seq |
| ENCFF118HUH | bed narrowPeak | NR2F2 | K562 | 1, 2 | TF ChIP-seq |
| ENCFF835FPJ | bed narrowPeak | NR2F6 | K562 | 1, 2 | TF ChIP-seq |
| ENCFF821YMC | bed narrowPeak | NR3C1 | K562 | 1, 2 | TF ChIP-seq |
| ENCFF137TCL | bed narrowPeak | NR4A1 | K562 | 1, 2 | TF ChIP-seq |
| ENCFF410RJD | bed narrowPeak | NRF1 | K562 | 1, 2 | TF ChIP-seq |
| ENCFF885JMZ | bed narrowPeak | NUFIP1 | K562 | 1, 2 | TF ChIP-seq |
| ENCFF743JJF | bed narrowPeak | PATZ1 | K562 | 1, 2 | TF ChIP-seq |
| ENCFF685CMQ | bed narrowPeak | PBX2 | K562 | 1, 2 | TF ChIP-seq |
| ENCFF956MGE | bed narrowPeak | PCBP1 | K562 | 1, 2 | TF ChIP-seq |
| ENCFF941XZW | bed narrowPeak | PCBP2 | K562 | 1, 2 | TF ChIP-seq |
| ENCFF424OWM | bed narrowPeak | PHB | K562 | 1, 2 | TF ChIP-seq |
| ENCFF988OXX | bed narrowPeak | PHB2 | K562 | 1, 2 | TF ChIP-seq |
| ENCFF259HUS | bed narrowPeak | PHF20 | K562 | 1, 2 | TF ChIP-seq |
| ENCFF657UVA | bed narrowPeak | PHF21A | K562 | 1, 2 | TF ChIP-seq |
| ENCFF981ISM | bed narrowPeak | PHF8 | K562 | 2, 3 | TF ChIP-seq |
| ENCFF130WQT | bed narrowPeak | PHTF2 | K562 | 1, 2 | TF ChIP-seq |
| ENCFF062VBB | bed narrowPeak | PKNOX1 | K562 | 1, 2 | TF ChIP-seq |

| | | | | |
|---|---|---|---|---|
| ENCFF800QDU | bed narrowPeak | PML | K562 | 1, 2 TF ChIP-seq |
| ENCFF634JRD | bed narrowPeak | POLR2A | K562 | 1, 2 TF ChIP-seq |
| ENCFF950YZE | bed narrowPeak | POLR2AphosphoS2 | K562 | 1, 2 TF ChIP-seq |
| ENCFF060MMW | bed narrowPeak | POLR2AphosphoS5 | K562 | 1, 2 TF ChIP-seq |
| ENCFF821QOS | bed narrowPeak | POLR2B | K562 | 1, 2 TF ChIP-seq |
| ENCFF413LLO | bed narrowPeak | POLR2G | K562 | 1, 2 TF ChIP-seq |
| ENCFF084KHS | bed narrowPeak | POLR2H | K562 | 1, 2 TF ChIP-seq |
| ENCFF627DJP | bed narrowPeak | POLR3A | K562 | 1, 2 TF ChIP-seq |
| ENCFF307GHJ | bed narrowPeak | POLR3G | K562 | 1, 2 TF ChIP-seq |
| ENCFF814QPF | bed narrowPeak | POU5F1 | K562 | 1, 2 TF ChIP-seq |
| ENCFF600HPZ | bed narrowPeak | PRDM10 | K562 | 1, 2 TF ChIP-seq |
| ENCFF018TNP | bed narrowPeak | PRMT5 | K562 | 1, 2 TF ChIP-seq |
| ENCFF417RQZ | bed narrowPeak | PRPF4 | K562 | 1, 2 TF ChIP-seq |
| ENCFF835NOD | bed narrowPeak | PTBP1 | K562 | 1, 2 TF ChIP-seq |
| ENCFF229BQZ | bed narrowPeak | PTRF | K562 | 1, 2 TF ChIP-seq |
| ENCFF587PBU | bed narrowPeak | PTTG1 | K562 | 1, 2 TF ChIP-seq |
| ENCFF500BWO | bed narrowPeak | PURB | K562 | 1, 2 TF ChIP-seq |
| ENCFF720FJY | bed narrowPeak | PYGO2 | K562 | 1, 2 TF ChIP-seq |
| ENCFF930WPG | bed narrowPeak | RAD21 | K562 | 1, 2 TF ChIP-seq |
| ENCFF740OPF | bed narrowPeak | RAD51 | K562 | 1, 2 TF ChIP-seq |
| ENCFF328QZM | bed narrowPeak | RB1 | K562 | 3, 4 TF ChIP-seq |
| ENCFF942VGF | bed narrowPeak | RBBP5 | K562 | 1, 2 TF ChIP-seq |
| ENCFF538ACI | bed narrowPeak | RBFOX2 | K562 | 1, 2 TF ChIP-seq |
| ENCFF465UMU | bed narrowPeak | RBM14 | K562 | 1, 2 TF ChIP-seq |
| ENCFF563WDZ | bed narrowPeak | RBM15 | K562 | 1, 2 TF ChIP-seq |
| ENCFF056OIG | bed narrowPeak | RBM17 | K562 | 1, 2 TF ChIP-seq |
| ENCFF420IBN | bed narrowPeak | RBM22 | K562 | 1, 2 TF ChIP-seq |
| ENCFF849VEO | bed narrowPeak | RBM25 | K562 | 1, 2 TF ChIP-seq |
| ENCFF670ILH | bed narrowPeak | RBM34 | K562 | 1, 2 TF ChIP-seq |
| ENCFF503DIK | bed narrowPeak | RBM39 | K562 | 1, 2 TF ChIP-seq |
| ENCFF691JZW | bed narrowPeak | RBPJ | K562 | 1, 2 TF ChIP-seq |
| ENCFF627PLM | bed narrowPeak | RCOR1 | K562 | 1, 2 TF ChIP-seq |
| ENCFF931SVP | bed narrowPeak | RELA | K562 | 1, 2 TF ChIP-seq |
| ENCFF539MIO | bed narrowPeak | REST | K562 | 1, 2, 3 TF ChIP-seq |
| ENCFF193PVX | bed narrowPeak | RFX1 | K562 | 3, 4 TF ChIP-seq |
| ENCFF201YKU | bed narrowPeak | RFX5 | K562 | 1, 2 TF ChIP-seq |
| ENCFF774NZZ | bed narrowPeak | RFX7 | K562 | 1, 2 TF ChIP-seq |
| ENCFF599CBB | bed narrowPeak | RLF | K562 | 2, 3 TF ChIP-seq |
| ENCFF271MJE | bed narrowPeak | RNF2 | K562 | 1, 2 TF ChIP-seq |
| ENCFF057RJK | bed narrowPeak | RREB1 | K562 | 1, 2 TF ChIP-seq |
| ENCFF003LPE | bed narrowPeak | RUNX1 | K562 | 1, 2 TF ChIP-seq |
| ENCFF537RNU | bed narrowPeak | SAFB | K562 | 1, 2 TF ChIP-seq |
| ENCFF087DKT | bed narrowPeak | SAFB2 | K562 | 1, 2 TF ChIP-seq |
| ENCFF383IEP | bed narrowPeak | SAP30 | K562 | 1, 2 TF ChIP-seq |
| ENCFF752EMN | bed narrowPeak | SETDB1 | K562 | 1, 2 TF ChIP-seq |
| ENCFF652ZEN | bed narrowPeak | SFPQ | K562 | 1, 2 TF ChIP-seq |
| ENCFF078EPC | bed narrowPeak | SHOX2 | K562 | 1, 2 TF ChIP-seq |
| ENCFF884MNF | bed narrowPeak | SIN3A | K562 | 1, 2 TF ChIP-seq |
| ENCFF543INR | bed narrowPeak | SIN3B | K562 | 5, 6 TF ChIP-seq |
| ENCFF217ACS | bed narrowPeak | SIRT6 | K562 | 1, 2 TF ChIP-seq |
| ENCFF754ZWP | bed narrowPeak | SIX5 | K562 | 1, 2 TF ChIP-seq |
| ENCFF254QDM | bed narrowPeak | SKIL | K562 | 2, 3 TF ChIP-seq |

| | | | | | |
|---|---|---|---|---|---|
| ENCFF084BUP | bed narrowPeak | SMAD1 | K562 | 1, 2 | TF ChIP-seq |
| ENCFF186MFI | bed narrowPeak | SMAD2 | K562 | 1, 2 | TF ChIP-seq |
| ENCFF069AAY | bed narrowPeak | SMAD5 | K562 | 1, 2 | TF ChIP-seq |
| ENCFF267OGF | bed narrowPeak | SMARCA4 | K562 | 1, 2 | TF ChIP-seq |
| ENCFF481TNF | bed narrowPeak | SMARCA5 | K562 | 1, 2 | TF ChIP-seq |
| ENCFF308QHX | bed narrowPeak | SMARCB1 | K562 | 1, 2 | TF ChIP-seq |
| ENCFF751ZVX | bed narrowPeak | SMARCC2 | K562 | 1, 3 | TF ChIP-seq |
| ENCFF148YMC | bed narrowPeak | SMARCE1 | K562 | 1, 2 | TF ChIP-seq |
| ENCFF175UEE | bed narrowPeak | SMC3 | K562 | 1, 2 | TF ChIP-seq |
| ENCFF656THP | bed narrowPeak | SNAPC5 | K562 | 1, 2 | TF ChIP-seq |
| ENCFF529BDW | bed narrowPeak | SNIP1 | K562 | 1, 2 | TF ChIP-seq |
| ENCFF206MJS | bed narrowPeak | SNRNP70 | K562 | 1, 2 | TF ChIP-seq |
| ENCFF431STY | bed narrowPeak | SOX6 | K562 | 1, 2 | TF ChIP-seq |
| ENCFF171NEU | bed narrowPeak | SP1 | K562 | 1, 2 | TF ChIP-seq |
| ENCFF741FZG | bed narrowPeak | SP2 | K562 | 1, 2 | TF ChIP-seq |
| ENCFF888CKG | bed narrowPeak | SPI1 | K562 | 1, 2 | TF ChIP-seq |
| ENCFF777MYW | bed narrowPeak | SREBF1 | K562 | 1, 2 | TF ChIP-seq |
| ENCFF710YFH | bed narrowPeak | SREBF2 | K562 | 1, 2 | TF ChIP-seq |
| ENCFF087EVW | bed narrowPeak | SRF | K562 | 1, 2 | TF ChIP-seq |
| ENCFF792SXS | bed narrowPeak | SRSF1 | K562 | 1, 2 | TF ChIP-seq |
| ENCFF926XGK | bed narrowPeak | SRSF3 | K562 | 1, 2 | TF ChIP-seq |
| ENCFF550VUN | bed narrowPeak | SRSF7 | K562 | 1, 2 | TF ChIP-seq |
| ENCFF217HAW | bed narrowPeak | SRSF9 | K562 | 1, 2 | TF ChIP-seq |
| ENCFF921BXP | bed narrowPeak | STAG1 | K562 | 1, 2 | TF ChIP-seq |
| ENCFF517IXK | bed narrowPeak | STAT5A | K562 | 1, 2 | TF ChIP-seq |
| ENCFF122FTW | bed narrowPeak | STAT5B | K562 | 1, 2 | TF ChIP-seq |
| ENCFF738XMN | bed narrowPeak | SUPT5H | K562 | 1, 2 | TF ChIP-seq |
| ENCFF889QYR | bed narrowPeak | SUZ12 | K562 | 1, 2 | TF ChIP-seq |
| ENCFF784BTI | bed narrowPeak | TAF1 | K562 | 1, 2 | TF ChIP-seq |
| ENCFF547PES | bed narrowPeak | TAF15 | K562 | 1, 2 | TF ChIP-seq |
| ENCFF792CKI | bed narrowPeak | TAF7 | K562 | 1, 2 | TF ChIP-seq |
| ENCFF223HDM | bed narrowPeak | TAF9B | K562 | 1, 2 | TF ChIP-seq |
| ENCFF852ZRK | bed narrowPeak | TAL1 | K562 | 1, 2 | TF ChIP-seq |
| ENCFF448YOS | bed narrowPeak | TARDBP | K562 | 1, 2 | TF ChIP-seq |
| ENCFF868SWL | bed narrowPeak | TBL1XR1 | K562 | 1, 2 | TF ChIP-seq |
| ENCFF370YGS | bed narrowPeak | TBP | K562 | 1, 2 | TF ChIP-seq |
| ENCFF927NEU | bed narrowPeak | TBPL1 | K562 | 1, 2 | TF ChIP-seq |
| ENCFF405EGZ | bed narrowPeak | TBX18 | K562 | 1, 2 | TF ChIP-seq |
| ENCFF912LXU | bed narrowPeak | TCF12 | K562 | 1, 2 | TF ChIP-seq |
| ENCFF773RNU | bed narrowPeak | TCF3 | K562 | 1, 2 | TF ChIP-seq |
| ENCFF512IAI | bed narrowPeak | TCF7 | K562 | 1, 2 | TF ChIP-seq |
| ENCFF556FYF | bed narrowPeak | TCF7L2 | K562 | 1, 2 | TF ChIP-seq |
| ENCFF692HUD | bed narrowPeak | TEAD1 | K562 | 1, 2 | TF ChIP-seq |
| ENCFF563LGJ | bed narrowPeak | TEAD2 | K562 | 1, 2 | TF ChIP-seq |
| ENCFF547MLB | bed narrowPeak | TEAD4 | K562 | 1, 2 | TF ChIP-seq |
| ENCFF392MAR | bed narrowPeak | TFCP2 | K562 | 1, 2 | TF ChIP-seq |
| ENCFF362JHQ | bed narrowPeak | TFDP1 | K562 | 1, 2 | TF ChIP-seq |
| ENCFF592NJN | bed narrowPeak | TFE3 | K562 | 1, 2 | TF ChIP-seq |
| ENCFF130TPD | bed narrowPeak | THAP1 | K562 | 1, 2 | TF ChIP-seq |
| ENCFF206LMB | bed narrowPeak | THAP12 | K562 | 1, 2 | TF ChIP-seq |
| ENCFF372YFU | bed narrowPeak | THAP7 | K562 | 1, 2 | TF ChIP-seq |
| ENCFF309DMZ | bed narrowPeak | THRA | K562 | 1, 2 | TF ChIP-seq |

| | | | | | |
|---|---|---|---|---|---|
| ENCFF067FJF | bed narrowPeak | THRAP3 | K562 | 1, 2 | TF ChIP-seq |
| ENCFF563WUP | bed narrowPeak | TOE1 | K562 | 1, 2 | TF ChIP-seq |
| ENCFF063NXI | bed narrowPeak | TRIM24 | K562 | 1, 2 | TF ChIP-seq |
| ENCFF514DDG | bed narrowPeak | TRIM25 | K562 | 1, 2 | TF ChIP-seq |
| ENCFF634OOU | bed narrowPeak | TRIM28 | K562 | 1, 2 | TF ChIP-seq |
| ENCFF534VQL | bed narrowPeak | TRIP13 | K562 | 1, 2 | TF ChIP-seq |
| ENCFF265QXK | bed narrowPeak | TSC22D4 | K562 | 1, 2 | TF ChIP-seq |
| ENCFF144ZLB | bed narrowPeak | TSHZ1 | K562 | 1, 2 | TF ChIP-seq |
| ENCFF482DRO | bed narrowPeak | U2AF1 | K562 | 1, 2 | TF ChIP-seq |
| ENCFF134HBP | bed narrowPeak | U2AF2 | K562 | 1, 2 | TF ChIP-seq |
| ENCFF403TAF | bed narrowPeak | UBTF | K562 | 1, 2 | TF ChIP-seq |
| ENCFF310CCS | bed narrowPeak | USF1 | K562 | 1, 2 | TF ChIP-seq |
| ENCFF289ZIR | bed narrowPeak | USF2 | K562 | 1, 2 | TF ChIP-seq |
| ENCFF433ERT | bed narrowPeak | VEZF1 | K562 | 3, 4 | TF ChIP-seq |
| ENCFF862UUR | bed narrowPeak | WHSC1 | K562 | 1 | TF ChIP-seq |
| ENCFF115PGE | bed narrowPeak | XRCC3 | K562 | 1, 2 | TF ChIP-seq |
| ENCFF929TWP | bed narrowPeak | XRCC5 | K562 | 1, 2 | TF ChIP-seq |
| ENCFF520DIY | bed narrowPeak | YBX1 | K562 | 1, 2 | TF ChIP-seq |
| ENCFF508WCC | bed narrowPeak | YBX3 | K562 | 1, 2 | TF ChIP-seq |
| ENCFF398UQZ | bed narrowPeak | YY1 | K562 | 1, 2 | TF ChIP-seq |
| ENCFF388TYU | bed narrowPeak | ZBED1 | K562 | 1, 2 | TF ChIP-seq |
| ENCFF554GSE | bed narrowPeak | ZBTB11 | K562 | 1, 2 | TF ChIP-seq |
| ENCFF830MTX | bed narrowPeak | ZBTB12 | K562 | 1, 2 | TF ChIP-seq |
| ENCFF292HSS | bed narrowPeak | ZBTB17 | K562 | 1, 2 | TF ChIP-seq |
| ENCFF189WAO | bed narrowPeak | ZBTB2 | K562 | 1, 2 | TF ChIP-seq |
| ENCFF146GZZ | bed narrowPeak | ZBTB33 | K562 | 1, 2 | TF ChIP-seq |
| ENCFF624PTB | bed narrowPeak | ZBTB40 | K562 | 1, 2 | TF ChIP-seq |
| ENCFF382IYX | bed narrowPeak | ZBTB5 | K562 | 1, 2 | TF ChIP-seq |
| ENCFF245LRG | bed narrowPeak | ZBTB7A | K562 | 1, 2 | TF ChIP-seq |
| ENCFF328SSL | bed narrowPeak | ZBTB8A | K562 | 1, 2 | TF ChIP-seq |
| ENCFF588CXX | bed narrowPeak | ZBTB9 | K562 | 1, 2 | TF ChIP-seq |
| ENCFF478PGJ | bed narrowPeak | ZC3H11A | K562 | 1, 2 | TF ChIP-seq |
| ENCFF270TMW | bed narrowPeak | ZC3H4 | K562 | 1, 2 | TF ChIP-seq |
| ENCFF045AOZ | bed narrowPeak | ZC3H8 | K562 | 1, 2 | TF ChIP-seq |
| ENCFF808NWU | bed narrowPeak | ZEB2 | K562 | 1, 2 | TF ChIP-seq |
| ENCFF795AWO | bed narrowPeak | ZFP1 | K562 | 1, 2 | TF ChIP-seq |
| ENCFF839GAS | bed narrowPeak | ZFP36 | K562 | 1, 2 | TF ChIP-seq |
| ENCFF150ZBH | bed narrowPeak | ZFP91 | K562 | 1, 2 | TF ChIP-seq |
| ENCFF910JTR | bed narrowPeak | ZFX | K562 | 1, 2 | TF ChIP-seq |
| ENCFF495BPY | bed narrowPeak | ZHX1 | K562 | 1, 3 | TF ChIP-seq |
| ENCFF560UGR | bed narrowPeak | ZKSCAN1 | K562 | 1, 2 | TF ChIP-seq |
| ENCFF697DRN | bed narrowPeak | ZKSCAN3 | K562 | 1, 2 | TF ChIP-seq |
| ENCFF908SNB | bed narrowPeak | ZKSCAN8 | K562 | 1, 2 | TF ChIP-seq |
| ENCFF881DAT | bed narrowPeak | ZMIZ1 | K562 | 1, 2 | TF ChIP-seq |
| ENCFF117XRE | bed narrowPeak | ZMYM3 | K562 | 1, 2 | TF ChIP-seq |
| ENCFF021JCJ | bed narrowPeak | ZNF12 | K562 | 1, 2 | TF ChIP-seq |
| ENCFF810OHB | bed narrowPeak | ZNF133 | K562 | 1, 2 | TF ChIP-seq |
| ENCFF700GZI | bed narrowPeak | ZNF143 | K562 | 1, 2 | TF ChIP-seq |
| ENCFF932FVX | bed narrowPeak | ZNF146 | K562 | 1 | TF ChIP-seq |
| ENCFF836QWY | bed narrowPeak | ZNF148 | K562 | 3, 4 | TF ChIP-seq |
| ENCFF169QYL | bed narrowPeak | ZNF165 | K562 | 1, 2 | TF ChIP-seq |
| ENCFF822FXR | bed narrowPeak | ZNF174 | K562 | 1, 2 | TF ChIP-seq |

| | | | | |
|---|---|---|---|---|
| ENCFF192MEM | bed narrowPeak | ZNF175 | K562 | 1, 2 TF ChIP-seq |
| ENCFF882XTP | bed narrowPeak | ZNF184 | K562 | 1, 2 TF ChIP-seq |
| ENCFF987UBO | bed narrowPeak | ZNF197 | K562 | 1, 2 TF ChIP-seq |
| ENCFF435MHH | bed narrowPeak | ZNF212 | K562 | 1, 2 TF ChIP-seq |
| ENCFF922RHM | bed narrowPeak | ZNF215 | K562 | 1, 2 TF ChIP-seq |
| ENCFF063VRD | bed narrowPeak | ZNF23 | K562 | 1, 2 TF ChIP-seq |
| ENCFF569TJP | bed narrowPeak | ZNF239 | K562 | 1, 2 TF ChIP-seq |
| ENCFF268YOM | bed narrowPeak | ZNF24 | K562 | 1, 2 TF ChIP-seq |
| ENCFF598QEL | bed narrowPeak | ZNF250 | K562 | 1, 2 TF ChIP-seq |
| ENCFF738IQL | bed narrowPeak | ZNF257 | K562 | 1, 2 TF ChIP-seq |
| ENCFF579ETM | bed narrowPeak | ZNF263 | K562 | 1, 2 TF ChIP-seq |
| ENCFF045CFL | bed narrowPeak | ZNF274 | K562 | 1, 2 TF ChIP-seq |
| ENCFF074WRG | bed narrowPeak | ZNF280A | K562 | 1, 2 TF ChIP-seq |
| ENCFF253UAF | bed narrowPeak | ZNF280B | K562 | 1, 2 TF ChIP-seq |
| ENCFF648ORA | bed narrowPeak | ZNF281 | K562 | 1, 2 TF ChIP-seq |
| ENCFF596JDS | bed narrowPeak | ZNF282 | K562 | 1, 2 TF ChIP-seq |
| ENCFF798AFV | bed narrowPeak | ZNF3 | K562 | 1, 2 TF ChIP-seq |
| ENCFF080KWE | bed narrowPeak | ZNF311 | K562 | 1, 2 TF ChIP-seq |
| ENCFF451AEQ | bed narrowPeak | ZNF316 | K562 | 1, 2 TF ChIP-seq |
| ENCFF577LQR | bed narrowPeak | ZNF318 | K562 | 1, 2 TF ChIP-seq |
| ENCFF873VFI | bed narrowPeak | ZNF319 | K562 | 1, 2 TF ChIP-seq |
| ENCFF265MQC | bed narrowPeak | ZNF324 | K562 | 1, 2 TF ChIP-seq |
| ENCFF843NBV | bed narrowPeak | ZNF347 | K562 | 1, 2 TF ChIP-seq |
| ENCFF712NHB | bed narrowPeak | ZNF354B | K562 | 1, 2 TF ChIP-seq |
| ENCFF864XZP | bed narrowPeak | ZNF384 | K562 | 1, 2 TF ChIP-seq |
| ENCFF275HDU | bed narrowPeak | ZNF395 | K562 | 1, 3 TF ChIP-seq |
| ENCFF193BGB | bed narrowPeak | ZNF397 | K562 | 1, 2 TF ChIP-seq |
| ENCFF538GSS | bed narrowPeak | ZNF407 | K562 | 1, 2 TF ChIP-seq |
| ENCFF207MLR | bed narrowPeak | ZNF408 | K562 | 1, 2 TF ChIP-seq |
| ENCFF318ZUN | bed narrowPeak | ZNF436 | K562 | 1, 2 TF ChIP-seq |
| ENCFF295XCB | bed narrowPeak | ZNF444 | K562 | 1, 2 TF ChIP-seq |
| ENCFF416TFM | bed narrowPeak | ZNF445 | K562 | 1, 2 TF ChIP-seq |
| ENCFF878SVX | bed narrowPeak | ZNF507 | K562 | 1, 2 TF ChIP-seq |
| ENCFF825ZIU | bed narrowPeak | ZNF512 | K562 | 1, 2 TF ChIP-seq |
| ENCFF096AYV | bed narrowPeak | ZNF518B | K562 | 1, 2 TF ChIP-seq |
| ENCFF913YMX | bed narrowPeak | ZNF551 | K562 | 1, 2 TF ChIP-seq |
| ENCFF945IST | bed narrowPeak | ZNF57 | K562 | 1, 2 TF ChIP-seq |
| ENCFF053ABJ | bed narrowPeak | ZNF583 | K562 | 1, 2 TF ChIP-seq |
| ENCFF005MBI | bed narrowPeak | ZNF584 | K562 | 1, 2 TF ChIP-seq |
| ENCFF345IHK | bed narrowPeak | ZNF589 | K562 | 1, 2 TF ChIP-seq |
| ENCFF847QJI | bed narrowPeak | ZNF592 | K562 | 1, 2 TF ChIP-seq |
| ENCFF017JQJ | bed narrowPeak | ZNF639 | K562 | 1, 2 TF ChIP-seq |
| ENCFF773XPT | bed narrowPeak | ZNF644 | K562 | 1, 2 TF ChIP-seq |
| ENCFF156EZP | bed narrowPeak | ZNF655 | K562 | 1, 2 TF ChIP-seq |
| ENCFF493LZB | bed narrowPeak | ZNF695 | K562 | 1, 2 TF ChIP-seq |
| ENCFF018HWM | bed narrowPeak | ZNF7 | K562 | 1, 2 TF ChIP-seq |
| ENCFF208OME | bed narrowPeak | ZNF700 | K562 | 1, 2 TF ChIP-seq |
| ENCFF159END | bed narrowPeak | ZNF717 | K562 | 1, 2 TF ChIP-seq |
| ENCFF219NIA | bed narrowPeak | ZNF740 | K562 | 1, 2 TF ChIP-seq |
| ENCFF307UZC | bed narrowPeak | ZNF75A | K562 | 1, 2 TF ChIP-seq |
| ENCFF305XQM | bed narrowPeak | ZNF76 | K562 | 1, 2 TF ChIP-seq |
| ENCFF049MDN | bed narrowPeak | ZNF764 | K562 | 1, 2 TF ChIP-seq |

| | | | | | |
|---|---|---|---|---|---|
| ENCFF168PYI | bed narrowPeak | ZNF766 | K562 | 1, 2 | TF ChIP-seq |
| ENCFF809JNC | bed narrowPeak | ZNF77 | K562 | 1, 2 | TF ChIP-seq |
| ENCFF165JQS | bed narrowPeak | ZNF778 | K562 | 1, 2 | TF ChIP-seq |
| ENCFF794OSY | bed narrowPeak | ZNF780A | K562 | 1, 2 | TF ChIP-seq |
| ENCFF352NGK | bed narrowPeak | ZNF785 | K562 | 1, 2 | TF ChIP-seq |
| ENCFF670RLH | bed narrowPeak | ZNF79 | K562 | 1, 2 | TF ChIP-seq |
| ENCFF791JWI | bed narrowPeak | ZNF83 | K562 | 1, 2 | TF ChIP-seq |
| ENCFF150ZBY | bed narrowPeak | ZNF830 | K562 | 1, 2 | TF ChIP-seq |
| ENCFF014HYS | bed narrowPeak | ZNF84 | K562 | 1, 2 | TF ChIP-seq |
| ENCFF908ZLN | bed narrowPeak | ZSCAN29 | K562 | 1, 2 | TF ChIP-seq |
| ENCFF537HHU | bed narrowPeak | ZSCAN32 | K562 | 1, 2 | TF ChIP-seq |
| ENCFF945HJR | bed narrowPeak | ZZZ3 | K562 | 1, 2 | TF ChIP-seq |
| ENCFF618NVV | bed narrowPeak | ARID3A | MCF-7 | 1, 2 | TF ChIP-seq |
| ENCFF760ZVI | bed narrowPeak | ATF7 | MCF-7 | 1, 2 | TF ChIP-seq |
| ENCFF414LXZ | bed narrowPeak | BMI1 | MCF-7 | 1, 2 | TF ChIP-seq |
| ENCFF740MEN | bed narrowPeak | BRCA2 | MCF-7 | 1, 2 | TF ChIP-seq |
| ENCFF240ZHW | bed narrowPeak | CEBPB | MCF-7 | 1, 2 | TF ChIP-seq |
| ENCFF930PBH | bed narrowPeak | CEBPG | MCF-7 | 1, 2 | TF ChIP-seq |
| ENCFF730UAD | bed narrowPeak | CHD1 | MCF-7 | 1, 3 | TF ChIP-seq |
| ENCFF025SMR | bed narrowPeak | CLOCK | MCF-7 | 1, 2 | TF ChIP-seq |
| ENCFF682WFF | bed narrowPeak | COPS2 | MCF-7 | 1, 2 | TF ChIP-seq |
| ENCFF495PCJ | bed narrowPeak | CREB1 | MCF-7 | 1, 2 | TF ChIP-seq |
| ENCFF982QZR | bed narrowPeak | CSDE1 | MCF-7 | 1, 2 | TF ChIP-seq |
| ENCFF456MGR | bed narrowPeak | CTBP1 | MCF-7 | 1, 2 | TF ChIP-seq |
| ENCFF785NTC | bed narrowPeak | CTCF | MCF-7 | 1, 2 | TF ChIP-seq |
| ENCFF762CDY | bed narrowPeak | CUX1 | MCF-7 | 2, 3 | TF ChIP-seq |
| ENCFF089GNH | bed narrowPeak | DDX20 | MCF-7 | 1, 2 | TF ChIP-seq |
| ENCFF042AWM | bed narrowPeak | DPF2 | MCF-7 | 1, 2 | TF ChIP-seq |
| ENCFF115KRO | bed narrowPeak | E2F1 | MCF-7 | 1, 2 | TF ChIP-seq |
| ENCFF090CEP | bed narrowPeak | E2F4 | MCF-7 | 1, 2 | TF ChIP-seq |
| ENCFF072VGV | bed narrowPeak | E2F8 | MCF-7 | 1, 2 | TF ChIP-seq |
| ENCFF347USC | bed narrowPeak | E4F1 | MCF-7 | 1, 2 | TF ChIP-seq |
| ENCFF427RSA | bed narrowPeak | EGR1 | MCF-7 | 1, 2 | TF ChIP-seq |
| ENCFF091NQE | bed narrowPeak | ELF1 | MCF-7 | 1, 2 | TF ChIP-seq |
| ENCFF408TWV | bed narrowPeak | ELK1 | MCF-7 | 1, 3 | TF ChIP-seq |
| ENCFF290DJX | bed narrowPeak | EP300 | MCF-7 | 1, 2 | TF ChIP-seq |
| ENCFF541DRZ | bed narrowPeak | ESRRA | MCF-7 | 1, 2 | TF ChIP-seq |
| ENCFF170POB | bed narrowPeak | FOS | MCF-7 | 1, 3 | TF ChIP-seq |
| ENCFF423KOE | bed narrowPeak | FOSL2 | MCF-7 | 1, 2 | TF ChIP-seq |
| ENCFF160RLI | bed narrowPeak | FOXA1 | MCF-7 | 1, 2 | TF ChIP-seq |
| ENCFF899MQW | bed narrowPeak | FOXK2 | MCF-7 | 1, 2 | TF ChIP-seq |
| ENCFF563MHZ | bed narrowPeak | FOXM1 | MCF-7 | 1, 2 | TF ChIP-seq |
| ENCFF678DJM | bed narrowPeak | GABPA | MCF-7 | 1, 2 | TF ChIP-seq |
| ENCFF565QWY | bed narrowPeak | GATA3 | MCF-7 | 1, 2 | TF ChIP-seq |
| ENCFF191SBE | bed narrowPeak | GATAD2B | MCF-7 | 1, 2 | TF ChIP-seq |
| ENCFF343QQE | bed narrowPeak | GTF2F1 | MCF-7 | 1, 2 | TF ChIP-seq |
| ENCFF401IAI | bed narrowPeak | HCFC1 | MCF-7 | 1, 3 | TF ChIP-seq |
| ENCFF661VXV | bed narrowPeak | HDAC2 | MCF-7 | 1, 2 | TF ChIP-seq |
| ENCFF161SFU | bed narrowPeak | HDGF | MCF-7 | 1, 2 | TF ChIP-seq |
| ENCFF144OPN | bed narrowPeak | HES1 | MCF-7 | 1, 2 | TF ChIP-seq |
| ENCFF708ACK | bed narrowPeak | HSF1 | MCF-7 | 1, 2 | TF ChIP-seq |
| ENCFF907UNK | bed narrowPeak | JUN | MCF-7 | 1, 2 | TF ChIP-seq |

| | | | | |
|---|---|---|---|---|
| ENCFF569ZCY | bed narrowPeak | JUND | MCF-7 | 1, 2 TF ChIP-seq |
| ENCFF461CLM | bed narrowPeak | KLF4 | MCF-7 | 1, 2 TF ChIP-seq |
| ENCFF891BPE | bed narrowPeak | KLF9 | MCF-7 | 3, 4 TF ChIP-seq |
| ENCFF736DZL | bed narrowPeak | LARP7 | MCF-7 | 1, 2 TF ChIP-seq |
| ENCFF873SVI | bed narrowPeak | MAFK | MCF-7 | 1, 2 TF ChIP-seq |
| ENCFF624CRN | bed narrowPeak | MAX | MCF-7 | 1, 2 TF ChIP-seq |
| ENCFF666YGQ | bed narrowPeak | MAZ | MCF-7 | 1, 2 TF ChIP-seq |
| ENCFF464QAL | bed narrowPeak | MBD2 | MCF-7 | 1, 2 TF ChIP-seq |
| ENCFF578NMN | bed narrowPeak | MLLT1 | MCF-7 | 1, 2 TF ChIP-seq |
| ENCFF432GSK | bed narrowPeak | MNT | MCF-7 | 1, 2 TF ChIP-seq |
| ENCFF283FBB | bed narrowPeak | MSX2 | MCF-7 | 1, 2 TF ChIP-seq |
| ENCFF225VFR | bed narrowPeak | MTA1 | MCF-7 | 1, 2 TF ChIP-seq |
| ENCFF180XXZ | bed narrowPeak | MTA2 | MCF-7 | 1, 2 TF ChIP-seq |
| ENCFF083AZM | bed narrowPeak | MTA3 | MCF-7 | 1, 2 TF ChIP-seq |
| ENCFF370EQJ | bed narrowPeak | MYC | MCF-7 | 1, 2 TF ChIP-seq |
| ENCFF209WRW | bed narrowPeak | NBN | MCF-7 | 1, 2 TF ChIP-seq |
| ENCFF320TAN | bed narrowPeak | NCOA3 | MCF-7 | 1, 2 TF ChIP-seq |
| ENCFF059LJD | bed narrowPeak | NEUROD1 | MCF-7 | 1, 2 TF ChIP-seq |
| ENCFF519XTN | bed narrowPeak | NFIB | MCF-7 | 1, 2 TF ChIP-seq |
| ENCFF895MJB | bed narrowPeak | NFRKB | MCF-7 | 1, 2 TF ChIP-seq |
| ENCFF927DIO | bed narrowPeak | NFXL1 | MCF-7 | 1, 2 TF ChIP-seq |
| ENCFF800CDQ | bed narrowPeak | NONO | MCF-7 | 1, 2 TF ChIP-seq |
| ENCFF386FQQ | bed narrowPeak | NR2F2 | MCF-7 | 1, 2 TF ChIP-seq |
| ENCFF269RME | bed narrowPeak | NRF1 | MCF-7 | 1, 3 TF ChIP-seq |
| ENCFF024BHX | bed narrowPeak | OVOL1 | MCF-7 | 1, 2 TF ChIP-seq |
| ENCFF473UHQ | bed narrowPeak | PAX8 | MCF-7 | 1, 2 TF ChIP-seq |
| ENCFF105PFS | bed narrowPeak | PKNOX1 | MCF-7 | 1, 2 TF ChIP-seq |
| ENCFF569JWM | bed narrowPeak | PML | MCF-7 | 1, 2 TF ChIP-seq |
| ENCFF456FTV | bed narrowPeak | POLR2A | MCF-7 | 1, 2 TF ChIP-seq |
| ENCFF508JRB | bed narrowPeak | PPP1R10 | MCF-7 | 1, 2 TF ChIP-seq |
| ENCFF682UUL | bed narrowPeak | RAD21 | MCF-7 | 1, 2 TF ChIP-seq |
| ENCFF091AYX | bed narrowPeak | RAD51 | MCF-7 | 1, 2 TF ChIP-seq |
| ENCFF838LXI | bed narrowPeak | RCOR1 | MCF-7 | 3, 4 TF ChIP-seq |
| ENCFF680JMZ | bed narrowPeak | REST | MCF-7 | 1, 2 TF ChIP-seq |
| ENCFF150PTQ | bed narrowPeak | RFX1 | MCF-7 | 1, 3 TF ChIP-seq |
| ENCFF103MPW | bed narrowPeak | RFX5 | MCF-7 | 1, 2 TF ChIP-seq |
| ENCFF803WYI | bed narrowPeak | SIN3A | MCF-7 | 1, 2 TF ChIP-seq |
| ENCFF441UHA | bed narrowPeak | SIX4 | MCF-7 | 1, 2 TF ChIP-seq |
| ENCFF618JNX | bed narrowPeak | SMARCA5 | MCF-7 | 1, 2 TF ChIP-seq |
| ENCFF761NKP | bed narrowPeak | SMARCE1 | MCF-7 | 1, 2 TF ChIP-seq |
| ENCFF455HWV | bed narrowPeak | SNIP1 | MCF-7 | 1, 2 TF ChIP-seq |
| ENCFF577EMC | bed narrowPeak | SP1 | MCF-7 | 1, 2 TF ChIP-seq |
| ENCFF464GFC | bed narrowPeak | SPDEF | MCF-7 | 1, 2 TF ChIP-seq |
| ENCFF275WAD | bed narrowPeak | SREBF1 | MCF-7 | 1, 2 TF ChIP-seq |
| ENCFF131TYZ | bed narrowPeak | SRF | MCF-7 | 1, 2 TF ChIP-seq |
| ENCFF258ZVN | bed narrowPeak | SUZ12 | MCF-7 | 1, 2 TF ChIP-seq |
| ENCFF075TIU | bed narrowPeak | TAF1 | MCF-7 | 1, 2 TF ChIP-seq |
| ENCFF233RBO | bed narrowPeak | TARDBP | MCF-7 | 1, 2 TF ChIP-seq |
| ENCFF987QLI | bed narrowPeak | TCF12 | MCF-7 | 1, 2 TF ChIP-seq |
| ENCFF332AZX | bed narrowPeak | TCF7L2 | MCF-7 | 1, 2 TF ChIP-seq |
| ENCFF144VMM | bed narrowPeak | TOE1 | MCF-7 | 1, 2 TF ChIP-seq |
| ENCFF452VLA | bed narrowPeak | TRIM22 | MCF-7 | 1, 3 TF ChIP-seq |

| | | | | | |
|---|---|---|---|---|---|
| ENCFF247VVK | bed narrowPeak | YBX1 | MCF-7 | 1, 2 | TF ChIP-seq |
| ENCFF589MVU | bed narrowPeak | ZBTB1 | MCF-7 | 1, 2 | TF ChIP-seq |
| ENCFF496RVC | bed narrowPeak | ZBTB11 | MCF-7 | 1, 2 | TF ChIP-seq |
| ENCFF780WLS | bed narrowPeak | ZBTB33 | MCF-7 | 1, 2 | TF ChIP-seq |
| ENCFF932XEU | bed narrowPeak | ZBTB40 | MCF-7 | 1, 2 | TF ChIP-seq |
| ENCFF794UEM | bed narrowPeak | ZBTB7B | MCF-7 | 1, 2 | TF ChIP-seq |
| ENCFF775BWJ | bed narrowPeak | ZFX | MCF-7 | 1, 2 | TF ChIP-seq |
| ENCFF694ZRC | bed narrowPeak | ZHX2 | MCF-7 | 1, 2 | TF ChIP-seq |
| ENCFF687REM | bed narrowPeak | ZKSCAN1 | MCF-7 | 3, 4 | TF ChIP-seq |
| ENCFF621ZSK | bed narrowPeak | ZNF207 | MCF-7 | 1, 2 | TF ChIP-seq |
| ENCFF083LCM | bed narrowPeak | ZNF217 | MCF-7 | 1, 2 | TF ChIP-seq |
| ENCFF699IZY | bed narrowPeak | ZNF227 | MCF-7 | 2, 3 | TF ChIP-seq |
| ENCFF619BFO | bed narrowPeak | ZNF24 | MCF-7 | 1, 2 | TF ChIP-seq |
| ENCFF611WRQ | bed narrowPeak | ZNF331 | MCF-7 | 1, 2 | TF ChIP-seq |
| ENCFF786XJV | bed narrowPeak | ZNF444 | MCF-7 | 1, 2 | TF ChIP-seq |
| ENCFF675SAG | bed narrowPeak | ZNF507 | MCF-7 | 1, 2 | TF ChIP-seq |
| ENCFF414EYO | bed narrowPeak | ZNF512B | MCF-7 | 1, 2 | TF ChIP-seq |
| ENCFF290LSS | bed narrowPeak | ZNF574 | MCF-7 | 1, 2 | TF ChIP-seq |
| ENCFF306PBX | bed narrowPeak | ZNF579 | MCF-7 | 1, 2 | TF ChIP-seq |
| ENCFF541HRT | bed narrowPeak | ZNF592 | MCF-7 | 1, 2 | TF ChIP-seq |
| ENCFF329QYZ | bed narrowPeak | ZNF687 | MCF-7 | 1, 2 | TF ChIP-seq |
| ENCFF525RRP | bed narrowPeak | ZNF8 | MCF-7 | 1, 2 | TF ChIP-seq |

#### Supplementary Table 2:

##### Overview on TF co-occurrence tools and evaluation on comparability

| Tool name | Maintained | Chosen for comparison | Year | Last updated | Source | Note |
|---|---|---|---|---|---|---|
| coTRaCTE | yes | no | 2018 | - | [1] | Precalculated data needed |
| PC-TraFF | yes | no | 2015 | - | [2] | Predefined motifs only |
| TACO | yes | no | 2014 | 2016 | [3] | Replicated data needed |
| CIS Miner | no | no | 2014 | 2016 | [4] | No input example |
| INSECT 2.0 | yes | no | 2015 | 2015 | [5] | Disregarded ( too different approach) |
| SpaMo | yes | yes | 2012 | 2021 | [6] | Comparable |
| iTFs | no | no | 2013 | - | [7] | Link not reachable |
| COPS | no | no | 2012 | 2012 | [8] | Limited to Drosophila & Mus musculus |
| CENDIST | no | no | 2011 | - | [9] | Link not reachable |
| TICA | no | no | 2018 | - | [10] | Link not reachable |
| Nautica | no | no | 2020 | - | [11] | Link not reachable |
| MCOT | yes | yes | 2019 | 2021 | [12] | Comparable |

#### Supplementary Table 3:

##### Feature comparison of MCOT, SPAMO and TF-COMB

| Tool name | MCOT | SPAMO | TF-COMB |
|---|---|---|---|
| **Overview** | | | |
| Input | ChIP-seq; Anchor motif; Partner motif(s); | ChIP-seq; Meme motifs | ChIP-seq; Meme motifs; Peaks with regions of interest |
| Type | One-against-all | One-against-all | All-against-all |
| ChIP-seq mandatory | Yes | Yes | No |
| Genome reference free | No | Yes | Yes |
| Programming language | C++ | C | Python, Cython |
| Output | plethora of result files (intermediate files, different files for metrics) | .html summary, result .tsv | result tsv, optional plots |
| **Features** | | | |
| Scoring | Yes | Yes | Yes |
| Differential analysis | No | No | Yes |
| Finding Hubs | No | No | Yes |
| Preferred Distance analysis | No | (Yes) | Yes |
| Dynamic parameters for distance analysis | (Yes) | No | Yes |
| Orientation analysis | forced | forced | optional |
| Anchor free motif prediction | No | No | Yes |
| Deriving networks | No | No | Yes |
| Network clustering | No | No | Yes |
| Overlapping motifs | Yes | No | Yes |
| **Plotting** | | | |
| Weighted networks | No | No | Yes |
| Network substructures | No | No | Yes |
| Preferred spacing | No | Yes | Yes |
| Paired footprint visualization | No | No | Yes |

### Supplementary Table 4:

#### Literature search for top 10 exclusively found pairs for MCOT, SPAMO and TF-COMB

| **Mcot Only** | | | | weak evidence |
|---|---|---|---|---|
| Anchor | Partner | Sources | | no evidence |
| ATF2 | SREBF2 | [13] | | strong evidence |
| NR2C1 | ESRRA | - | | |
| MEF2A | SMAD5 | [14] | | |
| SMAD5 | MEF2B | [14] | | |
| BHLHE40 | MYC | [15] | | |
| SPI1 | ETS1 | [16] | | |
| NR2F1 | USF1 | - | | |
| STAT3 | STAT5A | [17] | | |
| USF2 | SREBF1 | [18] | | |
| STAT3 | STAT1 | [16] | | |
| **Spamo Only** | | | | |
| Anchor | Partner | Sources | | |
| MEF2C | PBX3 | [18] | | |
| PAX5 | ZNF384 | - | | |
| ZNF384 | PBX3 | - | | |
| NFYB | MXI1 | | | |
| MEF2C | EGR1 | - | | |
| NFYB | MAX | [19] | | |
| NFIC | ZNF384 | - | | |
| PBX3 | CEBPB | - | | |
| PAX5 | CEBPB | - | | |
| ZNF384 | MAZ | - | | |
| **TF-COMB Only** | | | | |
| Anchor | Partner | Sources | | |
| JUNB | BATF | [20],[21],[22] | | |
| JUNB | NFIC | [23] | | |
| NFATC3 | TBX21 | [24] | | |
| IRF4 | CREM | [25] | | |
| JUNB | ATF2 | [26] | | |
| NFIC | BATF | * | | |
| MEF2B | EBF1 | [27] | | |
| CREM | ETV6 | - | | |
| JUNB | IRF4 | [20] | | |
| SP1 | CREM | [16]** | | |

* We found the pairs NFIC-JUNB and JUNB-BATF (top2)  with strong evidence. This hints on a connection between BATF-NFIC connected by JUNB

** Only reported in C. familiaris (BioGrid)

#### Supplementary Table 5:

##### Results of co-occurrence of ChIP-seq peaks across cell lines

| TF1 | TF2 | TF1_TF2_count | TF1_count | TF2_count | cosine | zscore | cell_line |
|---|---|---|---|---|---|---|---|
| FOSL2 | JUNB | 14274 | 43162 | 20675 | 0.478 | 90.383 | A549 |
| GATA3 | JUNB | 10288 | 23062 | 20675 | 0.471 | 115.365 | A549 |
| E2F6 | MAX | 12050 | 18250 | 43738 | 0.427 | 84.097 | A549 |
| JUN | MAX | 17173 | 39252 | 43738 | 0.414 | 51.693 | A549 |
| GATA3 | TEAD4 | 7251 | 23062 | 14330 | 0.399 | 111.333 | A549 |
| FOSL2 | GATA3 | 12237 | 43162 | 23062 | 0.388 | 60.506 | A549 |
| FOSB | HOXA7 | 2719 | 6489 | 7586 | 0.388 | 129.043 | A549 |
| EHF | HOXB13 | 2725 | 6247 | 8237 | 0.380 | 140.920 | A549 |
| ATOH8 | HOXB13 | 2816 | 7005 | 8237 | 0.371 | 114.163 | A549 |
| NR2E3 | TP53 | 2642 | 8105 | 6281 | 0.370 | 120.110 | A549 |
| ATOH8 | HOXB5 | 3177 | 7005 | 10516 | 0.370 | 133.932 | A549 |
| GATA3 | JUN | 11011 | 23062 | 39252 | 0.366 | 57.757 | A549 |
| HOXB13 | HOXB5 | 3360 | 8237 | 10516 | 0.361 | 152.978 | A549 |
| ATOH8 | EHF | 2373 | 7005 | 6247 | 0.359 | 128.919 | A549 |
| EHF | HOXB5 | 2785 | 6247 | 10516 | 0.344 | 120.340 | A549 |
| JUNB | TEAD4 | 5741 | 20675 | 14330 | 0.334 | 84.682 | A549 |
| EHF | ZNF302 | 1388 | 6247 | 2822 | 0.331 | 117.198 | A549 |
| MAX | MYC | 6673 | 43738 | 9501 | 0.327 | 68.723 | A549 |
| FOXF2 | FOXS1 | 2082 | 9371 | 4405 | 0.324 | 110.059 | A549 |
| EHF | FOXS1 | 1685 | 6247 | 4405 | 0.321 | 115.844 | A549 |
| JUN | JUNB | 8945 | 39252 | 20675 | 0.314 | 57.775 | A549 |
| HES2 | JUNB | 2858 | 4245 | 20675 | 0.305 | 114.575 | A549 |
| HOXB13 | ZNF302 | 1467 | 8237 | 2822 | 0.304 | 99.763 | A549 |
| ATOH8 | FOXS1 | 1680 | 7005 | 4405 | 0.302 | 118.804 | A549 |
| IKZF1 | IKZF2 | 41044 | 69901 | 53515 | 0.671 | 50.628 | GM12878 |
| BATF | JUNB | 22204 | 36828 | 34714 | 0.621 | 101.763 | GM12878 |
| CTCF | ZNF143 | 27606 | 51248 | 42469 | 0.592 | 63.633 | GM12878 |
| CREB1 | CREM | 18020 | 28275 | 33311 | 0.587 | 104.822 | GM12878 |
| FOS | NFYA | 893 | 2042 | 1141 | 0.585 | 215.378 | GM12878 |
| USF1 | USF2 | 4570 | 8460 | 7690 | 0.567 | 188.573 | GM12878 |
| MEF2A | MEF2C | 8899 | 22587 | 11826 | 0.544 | 173.504 | GM12878 |
| ATF2 | NFIC | 18466 | 30795 | 38532 | 0.536 | 83.961 | GM12878 |
| TCF12 | TCF3 | 11380 | 25135 | 18871 | 0.523 | 132.995 | GM12878 |
| BCL11A | IRF4 | 11306 | 21231 | 23042 | 0.511 | 119.652 | GM12878 |
| ATF2 | CREM | 15774 | 30795 | 33311 | 0.493 | 77.627 | GM12878 |
| FOXM1 | NFIC | 14109 | 22379 | 38532 | 0.480 | 77.692 | GM12878 |
| BATF | BCL11A | 13347 | 36828 | 21231 | 0.477 | 85.542 | GM12878 |
| BATF | IRF4 | 13893 | 36828 | 23042 | 0.477 | 85.525 | GM12878 |
| JUNB | NFIC | 17258 | 34714 | 38532 | 0.472 | 56.653 | GM12878 |
| ATF2 | BCL11A | 12031 | 30795 | 21231 | 0.471 | 81.716 | GM12878 |
| ATF2 | FOXM1 | 12275 | 30795 | 22379 | 0.468 | 93.800 | GM12878 |
| NFATC3 | TBX21 | 15597 | 25994 | 43559 | 0.464 | 60.361 | GM12878 |
| FOS | IRF3 | 1430 | 2042 | 4772 | 0.458 | 147.571 | GM12878 |
| ATF2 | MEF2A | 12064 | 30795 | 22587 | 0.457 | 85.072 | GM12878 |
| BCL11A | NFIC | 13057 | 21231 | 38532 | 0.457 | 69.868 | GM12878 |
| CREM | NFIC | 16262 | 33311 | 38532 | 0.454 | 58.064 | GM12878 |

| | | | | | | | |
|---|---|---|---|---|---|---|---|
| BCL11A | CREM | 11958 | 21231 | 33311 | 0.450 | 88.028 | GM12878 |
| NFATC3 | RELB | 15879 | 25994 | 48090 | 0.449 | 60.897 | GM12878 |
| MEF2A | NFIC | 13234 | 22587 | 38532 | 0.449 | 58.780 | GM12878 |
| BCL11A | PAX5 | 10180 | 21231 | 24348 | 0.448 | 88.344 | GM12878 |
| MAX | MXI1 | 9231 | 18647 | 22963 | 0.446 | 100.319 | GM12878 |
| ATF2 | IRF4 | 11834 | 30795 | 23042 | 0.444 | 93.399 | GM12878 |
| ATF7 | NFATC3 | 14894 | 44211 | 25994 | 0.439 | 54.479 | GM12878 |
| CREM | IRF4 | 11950 | 33311 | 23042 | 0.431 | 71.023 | GM12878 |
| NFATC3 | SKIL | 11116 | 25994 | 25597 | 0.431 | 78.641 | GM12878 |
| ATF2 | JUNB | 14059 | 30795 | 34714 | 0.430 | 53.038 | GM12878 |
| FOXM1 | MEF2A | 9662 | 22379 | 22587 | 0.430 | 82.026 | GM12878 |
| BCL11A | FOXM1 | 9345 | 21231 | 22379 | 0.429 | 90.880 | GM12878 |
| BCL11A | JUNB | 11626 | 21231 | 34714 | 0.428 | 89.335 | GM12878 |
| BCL11A | MEF2A | 9326 | 21231 | 22587 | 0.426 | 91.259 | GM12878 |
| CREM | MEF2A | 11547 | 33311 | 22587 | 0.421 | 73.078 | GM12878 |
| BCL11A | IKZF2 | 14121 | 21231 | 53515 | 0.419 | 50.507 | GM12878 |
| IRF4 | JUNB | 11848 | 23042 | 34714 | 0.419 | 54.249 | GM12878 |
| ETV6 | NFIC | 12204 | 22142 | 38532 | 0.418 | 69.087 | GM12878 |
| SKIL | TBX21 | 13911 | 25597 | 43559 | 0.417 | 55.436 | GM12878 |
| CREM | ETV6 | 11210 | 33311 | 22142 | 0.413 | 75.251 | GM12878 |
| ATF2 | ETV6 | 10758 | 30795 | 22142 | 0.412 | 53.301 | GM12878 |
| IRF4 | NFIC | 12259 | 23042 | 38532 | 0.411 | 62.798 | GM12878 |
| BCL11A | ETV6 | 8918 | 21231 | 22142 | 0.411 | 92.463 | GM12878 |
| BCL11A | TBX21 | 12501 | 21231 | 43559 | 0.411 | 60.838 | GM12878 |
| CREM | SP1 | 9330 | 33311 | 15591 | 0.409 | 71.293 | GM12878 |
| PBX3 | PKNOX1 | 8759 | 12586 | 36603 | 0.408 | 85.282 | GM12878 |
| CREM | PAX5 | 11621 | 33311 | 24348 | 0.408 | 50.252 | GM12878 |
| CREM | FOXM1 | 11137 | 33311 | 22379 | 0.408 | 63.742 | GM12878 |
| ARNT | NFATC3 | 9556 | 21468 | 25994 | 0.405 | 81.866 | GM12878 |
| BCL11A | NFATC3 | 9438 | 21231 | 25994 | 0.402 | 85.135 | GM12878 |
| ETV6 | IRF4 | 9017 | 22142 | 23042 | 0.399 | 92.367 | GM12878 |
| IRF4 | PAX5 | 9445 | 23042 | 24348 | 0.399 | 62.505 | GM12878 |
| CREB1 | SP1 | 8334 | 28275 | 15591 | 0.397 | 84.781 | GM12878 |
| PAX5 | TCF12 | 9819 | 24348 | 25135 | 0.397 | 69.445 | GM12878 |
| ATF2 | CREB1 | 11705 | 30795 | 28275 | 0.397 | 58.372 | GM12878 |
| ARNT | SKIL | 9262 | 21468 | 25597 | 0.395 | 69.879 | GM12878 |
| ATF3 | USF2 | 2196 | 4023 | 7690 | 0.395 | 141.728 | GM12878 |
| FOXM1 | JUNB | 10847 | 22379 | 34714 | 0.389 | 65.906 | GM12878 |
| FOXM1 | IRF4 | 8836 | 22379 | 23042 | 0.389 | 72.260 | GM12878 |
| ATF3 | USF1 | 2264 | 4023 | 8460 | 0.388 | 123.052 | GM12878 |
| IRF3 | NFYA | 900 | 4772 | 1141 | 0.386 | 116.270 | GM12878 |
| ATF2 | NFATC3 | 10879 | 30795 | 25994 | 0.385 | 51.135 | GM12878 |
| BCL11A | SKIL | 8951 | 21231 | 25597 | 0.384 | 74.381 | GM12878 |
| CREM | GABPA | 8220 | 33311 | 13947 | 0.381 | 71.965 | GM12878 |
| ETS1 | SIX5 | 3200 | 12891 | 5476 | 0.381 | 137.331 | GM12878 |
| IRF4 | MEF2A | 8688 | 23042 | 22587 | 0.381 | 83.756 | GM12878 |
| ELF1 | ETS1 | 7124 | 27368 | 12891 | 0.379 | 82.845 | GM12878 |
| ATF2 | ZBED1 | 7953 | 30795 | 14421 | 0.377 | 71.244 | GM12878 |
| BHLHE40 | PAX5 | 9955 | 28650 | 24348 | 0.377 | 60.443 | GM12878 |
| PAX5 | POU2F2 | 9828 | 24348 | 27931 | 0.377 | 55.202 | GM12878 |
| PAX5 | SP1 | 7332 | 24348 | 15591 | 0.376 | 86.380 | GM12878 |
| JUNB | MEF2A | 10485 | 34714 | 22587 | 0.374 | 56.027 | GM12878 |

| | | | | | | | |
|---|---|---|---|---|---|---|---|
| ATF2 | PAX5 | 10248 | 30795 | 24348 | 0.374 | 61.745 | GM12878 |
| CEBPZ | NFYA | 455 | 1300 | 1141 | 0.374 | 139.245 | GM12878 |
| ETV6 | FOXM1 | 8277 | 22142 | 22379 | 0.372 | 59.497 | GM12878 |
| CREB1 | GABPA | 7271 | 28275 | 13947 | 0.366 | 80.582 | GM12878 |
| FOXM1 | SKIL | 8745 | 22379 | 25597 | 0.365 | 62.327 | GM12878 |
| CEBPZ | FOS | 595 | 1300 | 2042 | 0.365 | 135.477 | GM12878 |
| BCL11A | TCF12 | 8428 | 21231 | 25135 | 0.365 | 62.289 | GM12878 |
| ATF2 | NFATC1 | 8030 | 30795 | 15744 | 0.365 | 59.969 | GM12878 |
| FOXM1 | ZBED1 | 6531 | 22379 | 14421 | 0.364 | 83.681 | GM12878 |
| ATF2 | MEF2C | 6937 | 30795 | 11826 | 0.364 | 81.517 | GM12878 |
| CREM | TCF12 | 10512 | 33311 | 25135 | 0.363 | 55.576 | GM12878 |
| MAZ | MXI1 | 8446 | 23716 | 22963 | 0.362 | 53.640 | GM12878 |
| ETV6 | MEF2A | 8049 | 22142 | 22587 | 0.360 | 51.844 | GM12878 |
| NFIC | ZBED1 | 8470 | 38532 | 14421 | 0.359 | 63.012 | GM12878 |
| BCL11A | ZBED1 | 6280 | 21231 | 14421 | 0.359 | 76.137 | GM12878 |
| POU2F2 | TCF12 | 9507 | 27931 | 25135 | 0.359 | 54.207 | GM12878 |
| ARNT | BCL11A | 7602 | 21468 | 21231 | 0.356 | 56.207 | GM12878 |
| IRF4 | NFATC3 | 8695 | 23042 | 25994 | 0.355 | 54.676 | GM12878 |
| BHLHE40 | SP1 | 7457 | 28650 | 15591 | 0.353 | 61.053 | GM12878 |
| IRF4 | SKIL | 8555 | 23042 | 25597 | 0.352 | 51.389 | GM12878 |
| BCL11A | BHLHE40 | 8664 | 21231 | 28650 | 0.351 | 54.046 | GM12878 |
| NFATC1 | STAT5A | 4641 | 15744 | 11090 | 0.351 | 96.922 | GM12878 |
| MAZ | SP1 | 6720 | 23716 | 15591 | 0.349 | 63.245 | GM12878 |
| ETV6 | ZBED1 | 6219 | 22142 | 14421 | 0.348 | 66.676 | GM12878 |
| MAX | MAZ | 7247 | 18647 | 23716 | 0.345 | 54.031 | GM12878 |
| BCL11A | MEF2C | 5451 | 21231 | 11826 | 0.344 | 63.513 | GM12878 |
| FOXM1 | MEF2C | 5569 | 22379 | 11826 | 0.342 | 80.516 | GM12878 |
| MEF2C | NFIC | 7291 | 11826 | 38532 | 0.342 | 64.748 | GM12878 |
| NFATC1 | ZBED1 | 5146 | 15744 | 14421 | 0.342 | 74.728 | GM12878 |
| ELF1 | SP1 | 7048 | 27368 | 15591 | 0.341 | 53.591 | GM12878 |
| ETV6 | NFATC3 | 8184 | 22142 | 25994 | 0.341 | 54.011 | GM12878 |
| POU2F2 | SP1 | 7116 | 27931 | 15591 | 0.341 | 61.211 | GM12878 |
| PAX5 | TCF3 | 7301 | 24348 | 18871 | 0.341 | 65.719 | GM12878 |
| NR2C1 | SIX5 | 2621 | 10830 | 5476 | 0.340 | 105.347 | GM12878 |
| MEF2A | ZBED1 | 6130 | 22587 | 14421 | 0.340 | 67.647 | GM12878 |
| SP1 | TCF12 | 6718 | 15591 | 25135 | 0.339 | 56.924 | GM12878 |
| KLF5 | ZNF207 | 5124 | 13891 | 16441 | 0.339 | 79.248 | GM12878 |
| ETS1 | GABPA | 4533 | 12891 | 13947 | 0.338 | 79.040 | GM12878 |
| MEF2A | PAX5 | 7925 | 22587 | 24348 | 0.338 | 50.931 | GM12878 |
| BHLHE40 | IRF4 | 8671 | 28650 | 23042 | 0.337 | 53.755 | GM12878 |
| CREM | MEF2C | 6692 | 33311 | 11826 | 0.337 | 59.231 | GM12878 |
| IRF4 | TCF12 | 8041 | 23042 | 25135 | 0.334 | 61.099 | GM12878 |
| IRF4 | JUND | 4415 | 23042 | 7601 | 0.334 | 101.168 | GM12878 |
| SIX5 | SREBF1 | 1456 | 5476 | 3497 | 0.333 | 109.016 | GM12878 |
| FOXM1 | NFATC1 | 6243 | 22379 | 15744 | 0.333 | 54.665 | GM12878 |
| ATF2 | GABPA | 6864 | 30795 | 13947 | 0.331 | 53.605 | GM12878 |
| ETV6 | GABPA | 5807 | 22142 | 13947 | 0.330 | 63.720 | GM12878 |
| BACH1 | NKRF | 7750 | 24260 | 22769 | 0.330 | 52.998 | GM12878 |
| BCL11A | NFATC1 | 6014 | 21231 | 15744 | 0.329 | 59.043 | GM12878 |
| BCL11A | JUND | 4139 | 21231 | 7601 | 0.326 | 79.920 | GM12878 |
| FOS | NFYB | 1778 | 2042 | 14677 | 0.325 | 84.996 | GM12878 |
| MXI1 | TBP | 7006 | 22963 | 20320 | 0.324 | 60.992 | GM12878 |

| | | | | | | | |
|---|---|---|---|---|---|---|---|
| IRF4 | ZBED1 | 5877 | 23042 | 14421 | 0.322 | 59.224 | GM12878 |
| ETV6 | SKIL | 7560 | 22142 | 25597 | 0.318 | 53.493 | GM12878 |
| IRF3 | SP1 | 2721 | 4772 | 15591 | 0.315 | 83.094 | GM12878 |
| ETS1 | SP1 | 4457 | 12891 | 15591 | 0.314 | 63.319 | GM12878 |
| MEF2C | ZBED1 | 4103 | 11826 | 14421 | 0.314 | 67.459 | GM12878 |
| CEBPB | FOXM1 | 3370 | 5173 | 22379 | 0.313 | 97.187 | GM12878 |
| CREM | JUND | 4976 | 33311 | 7601 | 0.313 | 57.835 | GM12878 |
| FOS | SP1 | 1761 | 2042 | 15591 | 0.312 | 91.117 | GM12878 |
| IRF4 | NFATC1 | 5867 | 23042 | 15744 | 0.308 | 65.727 | GM12878 |
| FOXM1 | TCF7 | 4555 | 22379 | 9835 | 0.307 | 57.818 | GM12878 |
| ELK1 | ETS1 | 3033 | 7580 | 12891 | 0.307 | 76.652 | GM12878 |
| MAX | USF2 | 3674 | 18647 | 7690 | 0.307 | 81.050 | GM12878 |
| ATF2 | JUND | 4685 | 30795 | 7601 | 0.306 | 51.371 | GM12878 |
| MEF2A | NFATC1 | 5741 | 22587 | 15744 | 0.304 | 57.213 | GM12878 |
| STAT5A | ZBED1 | 3850 | 11090 | 14421 | 0.304 | 64.427 | GM12878 |
| IRF4 | SP1 | 5760 | 23042 | 15591 | 0.304 | 54.024 | GM12878 |
| NFATC3 | ZBED1 | 5883 | 25994 | 14421 | 0.304 | 53.888 | GM12878 |
| ETV6 | NFATC1 | 5673 | 22142 | 15744 | 0.304 | 50.701 | GM12878 |
| ATF2 | STAT5A | 5614 | 30795 | 11090 | 0.304 | 54.557 | GM12878 |
| NFIC | TCF7 | 5907 | 38532 | 9835 | 0.303 | 52.114 | GM12878 |
| ELF1 | ELK1 | 4364 | 27368 | 7580 | 0.303 | 64.108 | GM12878 |
| BATF | JUND | 5042 | 36828 | 7601 | 0.301 | 69.481 | GM12878 |
| E2F6 | MAX | 18090 | 31301 | 41947 | 0.499 | 84.430 | H1 |
| CTCF | ZNF143 | 22202 | 57335 | 36137 | 0.488 | 53.554 | H1 |
| USF1 | USF2 | 7047 | 30940 | 8328 | 0.439 | 134.339 | H1 |
| ATF3 | USF2 | 2423 | 5657 | 8328 | 0.353 | 129.753 | H1 |
| CREB1 | TBP | 6510 | 19981 | 22252 | 0.309 | 53.809 | H1 |
| BACH1 | MAFK | 3974 | 12876 | 13405 | 0.302 | 73.055 | H1 |
| ZNF34 | ZNF426 | 7205 | 10050 | 8879 | 0.763 | 363.424 | HEK293 |
| ZNF426 | ZNF555 | 5652 | 8879 | 6703 | 0.733 | 274.994 | HEK293 |
| EGR2 | ZFHX2 | 45127 | 62403 | 66444 | 0.701 | 120.041 | HEK293 |
| ZNF34 | ZNF555 | 5657 | 10050 | 6703 | 0.689 | 350.019 | HEK293 |
| ZNF433 | ZNF555 | 3019 | 3798 | 6703 | 0.598 | 272.116 | HEK293 |
| ZNF433 | ZNF560 | 2175 | 3798 | 3492 | 0.597 | 213.803 | HEK293 |
| ZNF140 | ZNF555 | 4174 | 7978 | 6703 | 0.571 | 231.953 | HEK293 |
| ZNF426 | ZNF433 | 3293 | 8879 | 3798 | 0.567 | 209.693 | HEK293 |
| ZNF140 | ZNF433 | 3042 | 7978 | 3798 | 0.553 | 200.831 | HEK293 |
| ZNF140 | ZNF426 | 4644 | 7978 | 8879 | 0.552 | 212.828 | HEK293 |
| ZFP69B | ZNF692 | 15511 | 24305 | 34826 | 0.533 | 142.055 | HEK293 |
| ZNF34 | ZNF433 | 3285 | 10050 | 3798 | 0.532 | 203.805 | HEK293 |
| MAZ | PATZ1 | 22402 | 42804 | 41663 | 0.530 | 112.255 | HEK293 |
| SP2 | SP3 | 16013 | 31446 | 29750 | 0.524 | 124.987 | HEK293 |
| ZNF140 | ZNF34 | 4580 | 7978 | 10050 | 0.511 | 210.740 | HEK293 |
| ZNF555 | ZNF560 | 2474 | 6703 | 3492 | 0.511 | 211.233 | HEK293 |
| ZBTB21 | ZNF629 | 14800 | 20325 | 43843 | 0.496 | 143.132 | HEK293 |
| ZNF266 | ZNF433 | 2191 | 5240 | 3798 | 0.491 | 210.418 | HEK293 |
| ZNF426 | ZNF623 | 5527 | 8879 | 15029 | 0.478 | 192.118 | HEK293 |
| ZNF555 | ZNF623 | 4760 | 6703 | 15029 | 0.474 | 205.913 | HEK293 |
| ZNF426 | ZNF560 | 2638 | 8879 | 3492 | 0.474 | 172.049 | HEK293 |
| ZNF140 | ZNF560 | 2487 | 7978 | 3492 | 0.471 | 193.379 | HEK293 |
| ZEB2 | ZNF629 | 19682 | 40116 | 43843 | 0.469 | 78.431 | HEK293 |
| SCRT1 | SCRT2 | 19336 | 26938 | 64216 | 0.465 | 95.085 | HEK293 |

| | | | | | | | |
|---|---|---|---|---|---|---|---|
| ZNF518A | ZNF555 | 5188 | 18621 | 6703 | 0.464 | 189.979 | HEK293 |
| ZNF34 | ZNF518A | 6320 | 10050 | 18621 | 0.462 | 172.578 | HEK293 |
| SP7 | ZEB2 | 20699 | 50285 | 40116 | 0.461 | 87.137 | HEK293 |
| ZNF292 | ZNF433 | 1237 | 1922 | 3798 | 0.458 | 203.536 | HEK293 |
| ZNF34 | ZNF623 | 5599 | 10050 | 15029 | 0.456 | 157.679 | HEK293 |
| ZNF426 | ZNF518A | 5848 | 8879 | 18621 | 0.455 | 141.287 | HEK293 |
| SP7 | ZNF629 | 21080 | 50285 | 43843 | 0.449 | 68.597 | HEK293 |
| KLF8 | SP3 | 13817 | 32219 | 29750 | 0.446 | 100.003 | HEK293 |
| ZNF34 | ZNF560 | 2638 | 10050 | 3492 | 0.445 | 208.955 | HEK293 |
| MAZ | SP3 | 15876 | 42804 | 29750 | 0.445 | 96.056 | HEK293 |
| INSM2 | TSHZ1 | 6408 | 17714 | 12240 | 0.435 | 127.254 | HEK293 |
| ZNF266 | ZNF560 | 1860 | 5240 | 3492 | 0.435 | 216.384 | HEK293 |
| INSM2 | ZNF639 | 7707 | 17714 | 18233 | 0.429 | 136.496 | HEK293 |
| ZNF433 | ZNF623 | 3232 | 3798 | 15029 | 0.428 | 169.390 | HEK293 |
| ZNF292 | ZNF560 | 1105 | 1922 | 3492 | 0.427 | 208.590 | HEK293 |
| ZNF266 | ZNF555 | 2525 | 5240 | 6703 | 0.426 | 155.408 | HEK293 |
| ZFP37 | ZNF501 | 8903 | 19451 | 22541 | 0.425 | 133.099 | HEK293 |
| ZNF24 | ZNF639 | 8387 | 21595 | 18233 | 0.423 | 147.566 | HEK293 |
| KLF9 | SP3 | 11534 | 25095 | 29750 | 0.422 | 97.516 | HEK293 |
| GLIS2 | PATZ1 | 17295 | 40291 | 41663 | 0.422 | 74.465 | HEK293 |
| KLF8 | ZEB2 | 15164 | 32219 | 40116 | 0.422 | 73.216 | HEK293 |
| ZEB2 | ZXDB | 16221 | 40116 | 36901 | 0.422 | 88.559 | HEK293 |
| ZNF843 | ZXDB | 12944 | 25751 | 36901 | 0.420 | 93.030 | HEK293 |
| FEZF1 | ZNF843 | 12360 | 33696 | 25751 | 0.420 | 128.471 | HEK293 |
| ZNF629 | ZXDB | 16869 | 43843 | 36901 | 0.419 | 62.313 | HEK293 |
| ZNF292 | ZNF404 | 880 | 1922 | 2306 | 0.418 | 179.710 | HEK293 |
| ZBTB11 | ZNF518A | 8760 | 23626 | 18621 | 0.418 | 108.575 | HEK293 |
| ZNF324 | ZNF639 | 7274 | 16663 | 18233 | 0.417 | 121.288 | HEK293 |
| ZNF24 | ZNF324 | 7877 | 21595 | 16663 | 0.415 | 124.847 | HEK293 |
| ZNF292 | ZNF354C | 742 | 1922 | 1678 | 0.413 | 187.573 | HEK293 |
| SP7 | ZXDB | 17787 | 50285 | 36901 | 0.413 | 64.834 | HEK293 |
| ZNF404 | ZNF433 | 1222 | 2306 | 3798 | 0.413 | 177.894 | HEK293 |
| MAZ | ZBTB26 | 15950 | 42804 | 35149 | 0.411 | 79.211 | HEK293 |
| ZNF629 | ZNF639 | 11579 | 43843 | 18233 | 0.410 | 116.937 | HEK293 |
| YY1 | YY2 | 4718 | 12504 | 10615 | 0.410 | 140.995 | HEK293 |
| GLIS1 | GLIS2 | 20697 | 64378 | 40291 | 0.406 | 58.803 | HEK293 |
| MAZ | SP2 | 14880 | 42804 | 31446 | 0.406 | 80.939 | HEK293 |
| INSM2 | ZNF24 | 7926 | 17714 | 21595 | 0.405 | 132.903 | HEK293 |
| KLF8 | KLF9 | 11460 | 32219 | 25095 | 0.403 | 89.522 | HEK293 |
| ZNF24 | ZNF629 | 12387 | 21595 | 43843 | 0.403 | 84.461 | HEK293 |
| KLF8 | PATZ1 | 14731 | 32219 | 41663 | 0.402 | 76.724 | HEK293 |
| KLF8 | ZXDB | 13857 | 32219 | 36901 | 0.402 | 90.165 | HEK293 |
| ZNF433 | ZNF510 | 1840 | 3798 | 5540 | 0.401 | 182.187 | HEK293 |
| ZEB2 | ZNF24 | 11791 | 40116 | 21595 | 0.401 | 109.385 | HEK293 |
| KLF16 | KLF8 | 9578 | 18018 | 32219 | 0.398 | 116.349 | HEK293 |
| ZNF354C | ZNF404 | 780 | 1678 | 2306 | 0.397 | 170.156 | HEK293 |
| ZEB2 | ZNF843 | 12734 | 40116 | 25751 | 0.396 | 98.961 | HEK293 |
| KLF8 | MAZ | 14705 | 32219 | 42804 | 0.396 | 83.038 | HEK293 |
| INSM2 | ZNF629 | 11023 | 17714 | 43843 | 0.396 | 86.961 | HEK293 |
| KLF8 | SP7 | 15909 | 32219 | 50285 | 0.395 | 79.800 | HEK293 |
| ZNF140 | ZNF623 | 4313 | 7978 | 15029 | 0.394 | 145.867 | HEK293 |
| ZNF324 | ZNF391 | 5996 | 16663 | 13956 | 0.393 | 127.321 | HEK293 |

| | | | | | | | |
|---|---|---|---|---|---|---|---|
| ZNF266 | ZNF292 | 1247 | 5240 | 1922 | 0.393 | 172.969 | HEK293 |
| ZNF266 | ZNF426 | 2675 | 5240 | 8879 | 0.392 | 148.802 | HEK293 |
| INSM2 | ZNF843 | 8346 | 17714 | 25751 | 0.391 | 105.230 | HEK293 |
| KLF8 | ZNF629 | 14653 | 32219 | 43843 | 0.390 | 67.902 | HEK293 |
| ZNF140 | ZNF266 | 2515 | 7978 | 5240 | 0.389 | 148.776 | HEK293 |
| ZBTB8A | ZXDB | 15180 | 41359 | 36901 | 0.389 | 76.169 | HEK293 |
| ZNF639 | ZSCAN23 | 5019 | 18233 | 9154 | 0.388 | 133.712 | HEK293 |
| ZEB2 | ZNF692 | 14516 | 40116 | 34826 | 0.388 | 71.431 | HEK293 |
| KLF17 | KLF8 | 11799 | 28900 | 32219 | 0.387 | 87.527 | HEK293 |
| KLF16 | KLF9 | 8220 | 18018 | 25095 | 0.387 | 108.426 | HEK293 |
| INSM2 | ZEB2 | 10253 | 17714 | 40116 | 0.385 | 86.875 | HEK293 |
| ZNF394 | ZNF518A | 9430 | 32293 | 18621 | 0.385 | 114.955 | HEK293 |
| SP7 | ZNF843 | 13831 | 50285 | 25751 | 0.384 | 74.100 | HEK293 |
| PATZ1 | SP3 | 13513 | 41663 | 29750 | 0.384 | 72.839 | HEK293 |
| PATZ1 | ZEB2 | 15671 | 41663 | 40116 | 0.383 | 74.709 | HEK293 |
| TSHZ1 | ZNF639 | 5710 | 12240 | 18233 | 0.382 | 108.508 | HEK293 |
| ZNF138 | ZNF292 | 680 | 1647 | 1922 | 0.382 | 168.007 | HEK293 |
| OSR2 | ZNF843 | 11590 | 35833 | 25751 | 0.382 | 97.476 | HEK293 |
| ZNF266 | ZNF404 | 1326 | 5240 | 2306 | 0.381 | 125.249 | HEK293 |
| KLF9 | SP2 | 10702 | 25095 | 31446 | 0.381 | 97.888 | HEK293 |
| KLF10 | KLF8 | 10220 | 22350 | 32219 | 0.381 | 74.868 | HEK293 |
| KLF8 | SP2 | 12098 | 32219 | 31446 | 0.380 | 92.211 | HEK293 |
| ZNF404 | ZNF560 | 1078 | 2306 | 3492 | 0.380 | 172.784 | HEK293 |
| ZNF433 | ZNF518A | 3194 | 3798 | 18621 | 0.380 | 130.216 | HEK293 |
| ZNF391 | ZNF639 | 6052 | 13956 | 18233 | 0.379 | 106.854 | HEK293 |
| SP7 | ZNF24 | 12490 | 50285 | 21595 | 0.379 | 84.568 | HEK293 |
| INSM2 | ZNF324 | 6509 | 17714 | 16663 | 0.379 | 109.823 | HEK293 |
| ZEB2 | ZNF324 | 9787 | 40116 | 16663 | 0.379 | 90.684 | HEK293 |
| GLIS2 | MAZ | 15699 | 40291 | 42804 | 0.378 | 67.580 | HEK293 |
| ZNF34 | ZNF394 | 6806 | 10050 | 32293 | 0.378 | 120.017 | HEK293 |
| ZNF24 | ZNF843 | 8899 | 21595 | 25751 | 0.377 | 110.981 | HEK293 |
| KLF17 | SP7 | 14382 | 28900 | 50285 | 0.377 | 67.111 | HEK293 |
| TSHZ1 | ZNF843 | 6697 | 12240 | 25751 | 0.377 | 125.447 | HEK293 |
| ZNF629 | ZNF843 | 12666 | 43843 | 25751 | 0.377 | 85.702 | HEK293 |
| ZBTB21 | ZNF639 | 7222 | 20325 | 18233 | 0.375 | 108.646 | HEK293 |
| KLF8 | ZNF639 | 9086 | 32219 | 18233 | 0.375 | 87.518 | HEK293 |
| ZNF561 | ZSCAN30 | 9449 | 24194 | 26304 | 0.375 | 85.440 | HEK293 |
| ZEB2 | ZNF639 | 10117 | 40116 | 18233 | 0.374 | 85.924 | HEK293 |
| ZNF266 | ZNF34 | 2713 | 5240 | 10050 | 0.374 | 145.201 | HEK293 |
| ZEB2 | ZNF2 | 12794 | 40116 | 29224 | 0.374 | 79.928 | HEK293 |
| KLF9 | MAZ | 12214 | 25095 | 42804 | 0.373 | 78.568 | HEK293 |
| ZNF2 | ZNF324 | 8200 | 29224 | 16663 | 0.372 | 100.728 | HEK293 |
| ZNF510 | ZNF560 | 1627 | 5540 | 3492 | 0.370 | 166.130 | HEK293 |
| KLF17 | ZNF324 | 8108 | 28900 | 16663 | 0.369 | 91.715 | HEK293 |
| BCL11A | BCL11B | 4906 | 14204 | 12552 | 0.367 | 137.479 | HEK293 |
| ZNF202 | ZNF433 | 1853 | 6698 | 3798 | 0.367 | 159.276 | HEK293 |
| KLF1 | KLF9 | 11404 | 38403 | 25095 | 0.367 | 74.338 | HEK293 |
| KLF16 | ZNF324 | 6357 | 18018 | 16663 | 0.367 | 119.485 | HEK293 |
| ZNF354C | ZNF433 | 926 | 1678 | 3798 | 0.367 | 153.049 | HEK293 |
| ZBTB11 | ZNF555 | 4607 | 23626 | 6703 | 0.366 | 145.030 | HEK293 |
| ZNF2 | ZNF391 | 7391 | 29224 | 13956 | 0.366 | 127.481 | HEK293 |
| KLF8 | ZNF324 | 8478 | 32219 | 16663 | 0.366 | 97.437 | HEK293 |

| | | | | | | | |
|---|---|---|---|---|---|---|---|
| ZNF560 | ZNF623 | 2649 | 3492 | 15029 | 0.366 | 126.588 | HEK293 |
| ZNF324 | ZSCAN23 | 4514 | 16663 | 9154 | 0.365 | 127.795 | HEK293 |
| ZNF138 | ZNF404 | 712 | 1647 | 2306 | 0.365 | 146.054 | HEK293 |
| KLF8 | ZNF692 | 12225 | 32219 | 34826 | 0.365 | 75.494 | HEK293 |
| INSM2 | SP7 | 10876 | 17714 | 50285 | 0.364 | 83.876 | HEK293 |
| PRDM4 | ZNF843 | 8660 | 22102 | 25751 | 0.363 | 96.840 | HEK293 |
| TSHZ1 | ZNF24 | 5899 | 12240 | 21595 | 0.363 | 91.776 | HEK293 |
| ZNF692 | ZXDB | 13007 | 34826 | 36901 | 0.363 | 61.544 | HEK293 |
| KLF17 | ZEB2 | 12334 | 28900 | 40116 | 0.362 | 62.722 | HEK293 |
| ZFP69B | ZNF324 | 7289 | 24305 | 16663 | 0.362 | 106.626 | HEK293 |
| PATZ1 | SP2 | 13073 | 41663 | 31446 | 0.361 | 63.530 | HEK293 |
| ZNF501 | ZXDB | 10405 | 22541 | 36901 | 0.361 | 90.709 | HEK293 |
| SP7 | ZNF639 | 10919 | 50285 | 18233 | 0.361 | 64.850 | HEK293 |
| INSM2 | ZNF391 | 5664 | 17714 | 13956 | 0.360 | 130.425 | HEK293 |
| KLF1 | KLF8 | 12652 | 38403 | 32219 | 0.360 | 73.174 | HEK293 |
| INSM2 | ZSCAN23 | 4577 | 17714 | 9154 | 0.359 | 118.696 | HEK293 |
| FEZF1 | OSR2 | 12485 | 33696 | 35833 | 0.359 | 71.528 | HEK293 |
| ZNF394 | ZNF555 | 5280 | 32293 | 6703 | 0.359 | 106.523 | HEK293 |
| INSM2 | KLF8 | 8569 | 17714 | 32219 | 0.359 | 90.604 | HEK293 |
| ZNF19 | ZNF404 | 780 | 2055 | 2306 | 0.358 | 133.145 | HEK293 |
| ZNF561 | ZNF660 | 10019 | 24194 | 32350 | 0.358 | 92.924 | HEK293 |
| ZNF394 | ZNF426 | 6057 | 32293 | 8879 | 0.358 | 106.448 | HEK293 |
| PATZ1 | ZXDB | 14025 | 41663 | 36901 | 0.358 | 56.220 | HEK293 |
| OVOL3 | ZNF843 | 5634 | 9640 | 25751 | 0.358 | 97.775 | HEK293 |
| ZNF140 | ZNF518A | 4349 | 7978 | 18621 | 0.357 | 99.920 | HEK293 |
| SP3 | ZXDB | 11786 | 29750 | 36901 | 0.356 | 69.025 | HEK293 |
| ZNF518A | ZNF623 | 5936 | 18621 | 15029 | 0.355 | 97.435 | HEK293 |
| ZNF292 | ZNF555 | 1273 | 1922 | 6703 | 0.355 | 169.923 | HEK293 |
| ZNF138 | ZNF433 | 887 | 1647 | 3798 | 0.355 | 160.104 | HEK293 |
| ZNF24 | ZXDB | 10004 | 21595 | 36901 | 0.354 | 75.386 | HEK293 |
| ZNF324 | ZNF692 | 8535 | 16663 | 34826 | 0.354 | 98.662 | HEK293 |
| KLF8 | ZNF24 | 9339 | 32219 | 21595 | 0.354 | 80.106 | HEK293 |
| ZNF639 | ZNF843 | 7657 | 18233 | 25751 | 0.353 | 92.206 | HEK293 |
| ZNF24 | ZNF391 | 6129 | 21595 | 13956 | 0.353 | 117.372 | HEK293 |
| ZNF510 | ZNF555 | 2151 | 5540 | 6703 | 0.353 | 161.021 | HEK293 |
| WT1 | ZNF843 | 9613 | 28823 | 25751 | 0.353 | 71.373 | HEK293 |
| SP7 | ZNF324 | 10187 | 50285 | 16663 | 0.352 | 80.488 | HEK293 |
| KLF16 | KLF17 | 8024 | 18018 | 28900 | 0.352 | 94.330 | HEK293 |
| MAZ | ZNF398 | 12423 | 42804 | 29168 | 0.352 | 65.695 | HEK293 |
| ZNF19 | ZNF354C | 651 | 2055 | 1678 | 0.351 | 152.056 | HEK293 |
| ZNF2 | ZNF501 | 8992 | 29224 | 22541 | 0.350 | 72.938 | HEK293 |
| ZNF660 | ZSCAN30 | 10212 | 32350 | 26304 | 0.350 | 84.853 | HEK293 |
| ZNF266 | ZNF354C | 1037 | 5240 | 1678 | 0.350 | 146.590 | HEK293 |
| ZBTB11 | ZXDB | 10324 | 23626 | 36901 | 0.350 | 71.701 | HEK293 |
| ZNF394 | ZXDB | 12063 | 32293 | 36901 | 0.349 | 55.279 | HEK293 |
| ZNF639 | ZXDB | 9063 | 18233 | 36901 | 0.349 | 80.906 | HEK293 |
| ZEB1 | ZNF639 | 5590 | 14079 | 18233 | 0.349 | 101.834 | HEK293 |
| TSHZ1 | ZNF629 | 8080 | 12240 | 43843 | 0.349 | 75.165 | HEK293 |
| PATZ1 | ZNF398 | 12156 | 41663 | 29168 | 0.349 | 72.786 | HEK293 |
| ZNF324 | ZNF629 | 9412 | 16663 | 43843 | 0.348 | 75.833 | HEK293 |
| ZBTB26 | ZBTB8A | 13270 | 35149 | 41359 | 0.348 | 63.154 | HEK293 |
| KLF9 | PATZ1 | 11252 | 25095 | 41663 | 0.348 | 56.490 | HEK293 |

| | | | | | | | |
|---|---|---|---|---|---|---|---|
| KLF1 | KLF17 | 11585 | 38403 | 28900 | 0.348 | 65.245 | HEK293 |
| GLI2 | OVOL3 | 2582 | 5719 | 9640 | 0.348 | 98.572 | HEK293 |
| ZNF391 | ZSCAN23 | 3930 | 13956 | 9154 | 0.348 | 128.801 | HEK293 |
| BCL11A | TSHZ1 | 4582 | 14204 | 12240 | 0.348 | 134.444 | HEK293 |
| ZNF138 | ZNF354C | 577 | 1647 | 1678 | 0.347 | 147.672 | HEK293 |
| ZBTB21 | ZNF24 | 7264 | 20325 | 21595 | 0.347 | 82.534 | HEK293 |
| KLF16 | KLF7 | 5246 | 18018 | 12712 | 0.347 | 115.623 | HEK293 |
| ZNF24 | ZNF692 | 9494 | 21595 | 34826 | 0.346 | 79.046 | HEK293 |
| ZBTB11 | ZNF426 | 5007 | 23626 | 8879 | 0.346 | 118.170 | HEK293 |
| ZNF324 | ZNF843 | 7159 | 16663 | 25751 | 0.346 | 88.882 | HEK293 |
| KLF17 | ZNF629 | 12278 | 28900 | 43843 | 0.345 | 59.900 | HEK293 |
| KLF17 | ZNF639 | 7906 | 28900 | 18233 | 0.344 | 85.190 | HEK293 |
| SP7 | ZNF2 | 13174 | 50285 | 29224 | 0.344 | 55.594 | HEK293 |
| KLF17 | PATZ1 | 11922 | 28900 | 41663 | 0.344 | 66.161 | HEK293 |
| ZEB1 | ZEB2 | 8153 | 14079 | 40116 | 0.343 | 91.141 | HEK293 |
| BCL11A | INSM2 | 5435 | 14204 | 17714 | 0.343 | 95.817 | HEK293 |
| ZBTB11 | ZNF394 | 9439 | 23626 | 32293 | 0.342 | 68.879 | HEK293 |
| ZNF354C | ZNF560 | 827 | 1678 | 3492 | 0.342 | 134.928 | HEK293 |
| AEBP2 | ZNF292 | 742 | 2455 | 1922 | 0.342 | 148.797 | HEK293 |
| KLF10 | ZNF561 | 7936 | 22350 | 24194 | 0.341 | 78.790 | HEK293 |
| ZBTB11 | ZBTB8A | 10659 | 23626 | 41359 | 0.341 | 73.841 | HEK293 |
| KLF8 | ZNF843 | 9816 | 32219 | 25751 | 0.341 | 64.553 | HEK293 |
| KLF17 | ZNF391 | 6826 | 28900 | 13956 | 0.340 | 84.963 | HEK293 |
| ZNF24 | ZSCAN23 | 4777 | 21595 | 9154 | 0.340 | 107.634 | HEK293 |
| KLF17 | ZNF24 | 8485 | 28900 | 21595 | 0.340 | 74.535 | HEK293 |
| TSHZ1 | ZSCAN23 | 3595 | 12240 | 9154 | 0.340 | 113.422 | HEK293 |
| ZNF19 | ZNF292 | 674 | 2055 | 1922 | 0.339 | 171.967 | HEK293 |
| ZBTB11 | ZNF34 | 5218 | 23626 | 10050 | 0.339 | 96.479 | HEK293 |
| OVOL3 | TSHZ1 | 3676 | 9640 | 12240 | 0.338 | 120.104 | HEK293 |
| INSM2 | KLF17 | 7656 | 17714 | 28900 | 0.338 | 95.163 | HEK293 |
| ZNF2 | ZNF24 | 8490 | 29224 | 21595 | 0.338 | 77.828 | HEK293 |
| KLF8 | ZNF76 | 8080 | 32219 | 17824 | 0.337 | 96.952 | HEK293 |
| SP3 | ZNF76 | 7760 | 29750 | 17824 | 0.337 | 80.988 | HEK293 |
| ZNF324 | ZXDB | 8354 | 16663 | 36901 | 0.337 | 80.296 | HEK293 |
| KLF16 | SP3 | 7797 | 18018 | 29750 | 0.337 | 74.397 | HEK293 |
| ZEB2 | ZNF501 | 10117 | 40116 | 22541 | 0.336 | 69.093 | HEK293 |
| ZEB2 | ZFP69B | 10504 | 40116 | 24305 | 0.336 | 78.198 | HEK293 |
| KLF17 | ZNF692 | 10655 | 28900 | 34826 | 0.336 | 59.552 | HEK293 |
| KLF10 | KLF9 | 7952 | 22350 | 25095 | 0.336 | 95.460 | HEK293 |
| KLF8 | ZNF391 | 7115 | 32219 | 13956 | 0.336 | 92.381 | HEK293 |
| SP3 | ZBTB26 | 10846 | 29750 | 35149 | 0.335 | 63.683 | HEK293 |
| INSM2 | ZBTB21 | 6354 | 17714 | 20325 | 0.335 | 99.977 | HEK293 |
| ZFP69B | ZNF24 | 7671 | 24305 | 21595 | 0.335 | 91.411 | HEK293 |
| AEBP2 | ZNF354C | 679 | 2455 | 1678 | 0.335 | 146.359 | HEK293 |
| ZNF2 | ZNF639 | 7714 | 29224 | 18233 | 0.334 | 96.775 | HEK293 |
| ZNF433 | ZNF680 | 2136 | 3798 | 10768 | 0.334 | 125.714 | HEK293 |
| PATZ1 | ZBTB10 | 9201 | 41663 | 18260 | 0.334 | 73.715 | HEK293 |
| KLF16 | ZXDB | 8594 | 18018 | 36901 | 0.333 | 76.398 | HEK293 |
| KLF17 | MAZ | 11718 | 28900 | 42804 | 0.333 | 56.315 | HEK293 |
| ZNF266 | ZNF510 | 1795 | 5240 | 5540 | 0.333 | 122.209 | HEK293 |
| KLF16 | ZNF391 | 5274 | 18018 | 13956 | 0.333 | 94.272 | HEK293 |
| ZNF324 | ZSCAN21 | 6370 | 16663 | 22039 | 0.332 | 95.750 | HEK293 |

| | | | | | | | |
|---|---|---|---|---|---|---|---|
| KLF16 | ZEB2 | 8920 | 18018 | 40116 | 0.332 | 66.135 | HEK293 |
| PATZ1 | ZBTB26 | 12676 | 41663 | 35149 | 0.331 | 55.867 | HEK293 |
| KLF7 | KLF8 | 6702 | 12712 | 32219 | 0.331 | 76.056 | HEK293 |
| ZNF140 | ZNF292 | 1296 | 7978 | 1922 | 0.331 | 147.441 | HEK293 |
| FEZF1 | PRDM6 | 12964 | 33696 | 45657 | 0.331 | 54.179 | HEK293 |
| ZNF24 | ZSCAN21 | 7210 | 21595 | 22039 | 0.330 | 79.996 | HEK293 |
| ZNF76 | ZXDB | 8457 | 17824 | 36901 | 0.330 | 67.841 | HEK293 |
| ZBTB21 | ZNF324 | 6067 | 20325 | 16663 | 0.330 | 90.649 | HEK293 |
| KLF10 | ZEB2 | 9864 | 22350 | 40116 | 0.329 | 59.057 | HEK293 |
| ZNF292 | ZNF510 | 1074 | 1922 | 5540 | 0.329 | 125.341 | HEK293 |
| KLF10 | SP3 | 8481 | 22350 | 29750 | 0.329 | 83.180 | HEK293 |
| TSHZ1 | ZEB2 | 7274 | 12240 | 40116 | 0.328 | 79.585 | HEK293 |
| ZNF2 | ZNF692 | 10464 | 29224 | 34826 | 0.328 | 62.179 | HEK293 |
| ZNF2 | ZXDB | 10761 | 29224 | 36901 | 0.328 | 64.694 | HEK293 |
| IKZF3 | ZSCAN30 | 9379 | 31196 | 26304 | 0.327 | 68.319 | HEK293 |
| ZNF324 | ZNF610 | 5149 | 16663 | 14846 | 0.327 | 97.651 | HEK293 |
| INSM2 | ZXDB | 8361 | 17714 | 36901 | 0.327 | 72.238 | HEK293 |
| ZNF394 | ZNF501 | 8814 | 32293 | 22541 | 0.327 | 62.136 | HEK293 |
| KLF17 | ZNF2 | 9490 | 28900 | 29224 | 0.327 | 72.770 | HEK293 |
| ZNF2 | ZSCAN21 | 8283 | 29224 | 22039 | 0.326 | 80.781 | HEK293 |
| KLF16 | ZNF692 | 8162 | 18018 | 34826 | 0.326 | 71.725 | HEK293 |
| ZNF138 | ZNF560 | 781 | 1647 | 3492 | 0.326 | 143.036 | HEK293 |
| ZBTB8A | ZNF394 | 11879 | 41359 | 32293 | 0.325 | 58.123 | HEK293 |
| KLF8 | ZFP69B | 9087 | 32219 | 24305 | 0.325 | 61.129 | HEK293 |
| KLF16 | MAZ | 9017 | 18018 | 42804 | 0.325 | 63.823 | HEK293 |
| BCL11A | ZNF843 | 6206 | 14204 | 25751 | 0.324 | 103.262 | HEK293 |
| ZNF391 | ZNF843 | 6142 | 13956 | 25751 | 0.324 | 92.497 | HEK293 |
| ZBTB20 | ZBTB26 | 12532 | 42580 | 35149 | 0.324 | 52.515 | HEK293 |
| INSM2 | WT1 | 7319 | 17714 | 28823 | 0.324 | 68.202 | HEK293 |
| ZNF202 | ZNF560 | 1566 | 6698 | 3492 | 0.324 | 128.485 | HEK293 |
| KLF16 | PATZ1 | 8869 | 18018 | 41663 | 0.324 | 82.142 | HEK293 |
| GLI2 | TSHZ1 | 2708 | 5719 | 12240 | 0.324 | 145.264 | HEK293 |
| ZNF639 | ZSCAN21 | 6487 | 18233 | 22039 | 0.324 | 87.174 | HEK293 |
| SP3 | ZEB2 | 11173 | 29750 | 40116 | 0.323 | 59.389 | HEK293 |
| ZNF518A | ZNF560 | 2603 | 18621 | 3492 | 0.323 | 118.127 | HEK293 |
| ZEB2 | ZNF391 | 7637 | 40116 | 13956 | 0.323 | 77.416 | HEK293 |
| BCL11A | ZNF639 | 5184 | 14204 | 18233 | 0.322 | 107.154 | HEK293 |
| ZFP69B | ZNF639 | 6780 | 24305 | 18233 | 0.322 | 73.332 | HEK293 |
| ZNF692 | ZNF843 | 9642 | 34826 | 25751 | 0.322 | 66.406 | HEK293 |
| ZNF292 | ZNF426 | 1328 | 1922 | 8879 | 0.321 | 121.670 | HEK293 |
| KLF10 | ZNF629 | 10047 | 22350 | 43843 | 0.321 | 55.138 | HEK293 |
| INSM2 | OVOL3 | 4190 | 17714 | 9640 | 0.321 | 132.862 | HEK293 |
| ZEB1 | ZNF324 | 4910 | 14079 | 16663 | 0.321 | 81.299 | HEK293 |
| KLF17 | ZFP69B | 8487 | 28900 | 24305 | 0.320 | 80.791 | HEK293 |
| SP7 | ZNF501 | 10773 | 50285 | 22541 | 0.320 | 51.451 | HEK293 |
| ZNF202 | ZNF555 | 2143 | 6698 | 6703 | 0.320 | 114.225 | HEK293 |
| KLF8 | ZBTB26 | 10741 | 32219 | 35149 | 0.319 | 58.417 | HEK293 |
| ZNF354C | ZNF514 | 536 | 1678 | 1682 | 0.319 | 155.699 | HEK293 |
| ZNF404 | ZNF555 | 1254 | 2306 | 6703 | 0.319 | 116.380 | HEK293 |
| ZNF324 | ZNF501 | 6178 | 16663 | 22541 | 0.319 | 89.116 | HEK293 |
| KLF1 | KLF7 | 7034 | 38403 | 12712 | 0.318 | 76.637 | HEK293 |
| INSM2 | ZFP69B | 6602 | 17714 | 24305 | 0.318 | 85.994 | HEK293 |

| | | | | | | | |
|---|---|---|---|---|---|---|---|
| KLF17 | WT1 | 9177 | 28900 | 28823 | 0.318 | 66.685 | HEK293 |
| MAZ | ZNF76 | 8778 | 42804 | 17824 | 0.318 | 59.930 | HEK293 |
| KLF8 | ZNF2 | 9747 | 32219 | 29224 | 0.318 | 56.575 | HEK293 |
| ZBTB21 | ZEB2 | 9066 | 20325 | 40116 | 0.317 | 73.586 | HEK293 |
| ZNF391 | ZNF501 | 5625 | 13956 | 22541 | 0.317 | 93.778 | HEK293 |
| TSHZ1 | ZNF324 | 4529 | 12240 | 16663 | 0.317 | 88.856 | HEK293 |
| ZNF202 | ZNF266 | 1877 | 6698 | 5240 | 0.317 | 134.002 | HEK293 |
| ZEB2 | ZNF660 | 11410 | 40116 | 32350 | 0.317 | 54.037 | HEK293 |
| KLF17 | ZXDB | 10341 | 28900 | 36901 | 0.317 | 61.892 | HEK293 |
| ZFP69B | ZNF391 | 5827 | 24305 | 13956 | 0.316 | 78.630 | HEK293 |
| ZNF518A | ZXDB | 8289 | 18621 | 36901 | 0.316 | 64.093 | HEK293 |
| ZEB2 | ZNF561 | 9840 | 40116 | 24194 | 0.316 | 61.551 | HEK293 |
| AEBP2 | ZNF404 | 751 | 2455 | 2306 | 0.316 | 131.082 | HEK293 |
| ZNF138 | ZNF266 | 927 | 1647 | 5240 | 0.316 | 128.442 | HEK293 |
| ZBTB11 | ZNF433 | 2985 | 23626 | 3798 | 0.315 | 108.601 | HEK293 |
| SP7 | TSHZ1 | 7809 | 50285 | 12240 | 0.315 | 64.818 | HEK293 |
| AEBP2 | ZNF433 | 961 | 2455 | 3798 | 0.315 | 142.289 | HEK293 |
| ZNF426 | ZNF510 | 2206 | 8879 | 5540 | 0.315 | 121.857 | HEK293 |
| ZNF34 | ZNF510 | 2345 | 10050 | 5540 | 0.314 | 116.294 | HEK293 |
| ZNF140 | ZNF510 | 2087 | 7978 | 5540 | 0.314 | 135.400 | HEK293 |
| ZEB2 | ZSCAN21 | 9309 | 40116 | 22039 | 0.313 | 61.556 | HEK293 |
| ZBTB26 | ZNF76 | 7825 | 35149 | 17824 | 0.313 | 73.836 | HEK293 |
| TSHZ1 | WT1 | 5866 | 12240 | 28823 | 0.312 | 86.219 | HEK293 |
| ZNF266 | ZNF623 | 2768 | 5240 | 15029 | 0.312 | 94.433 | HEK293 |
| ZNF639 | ZNF692 | 7859 | 18233 | 34826 | 0.312 | 68.789 | HEK293 |
| ZFP37 | ZNF394 | 7808 | 19451 | 32293 | 0.312 | 75.220 | HEK293 |
| BCL11A | ZEB2 | 7427 | 14204 | 40116 | 0.311 | 79.141 | HEK293 |
| ZNF580 | ZXDB | 9456 | 25124 | 36901 | 0.311 | 66.464 | HEK293 |
| ZNF2 | ZNF843 | 8515 | 29224 | 25751 | 0.310 | 84.447 | HEK293 |
| BCL11A | ZNF24 | 5436 | 14204 | 21595 | 0.310 | 123.288 | HEK293 |
| ZNF580 | ZNF843 | 7893 | 25124 | 25751 | 0.310 | 65.122 | HEK293 |
| KLF16 | SP7 | 9334 | 18018 | 50285 | 0.310 | 51.384 | HEK293 |
| INSM2 | ZNF692 | 7702 | 17714 | 34826 | 0.310 | 78.861 | HEK293 |
| KLF7 | KLF9 | 5538 | 12712 | 25095 | 0.310 | 97.496 | HEK293 |
| ZNF19 | ZNF433 | 866 | 2055 | 3798 | 0.310 | 122.206 | HEK293 |
| KLF10 | ZXDB | 8897 | 22350 | 36901 | 0.310 | 50.713 | HEK293 |
| ZNF561 | ZXDB | 9247 | 24194 | 36901 | 0.309 | 54.287 | HEK293 |
| ZNF394 | ZNF639 | 7509 | 32293 | 18233 | 0.309 | 68.529 | HEK293 |
| ZNF518A | ZNF558 | 5197 | 18621 | 15148 | 0.309 | 86.391 | HEK293 |
| ZFP69B | ZNF2 | 8244 | 24305 | 29224 | 0.309 | 65.548 | HEK293 |
| INSM2 | ZSCAN21 | 6111 | 17714 | 22039 | 0.309 | 98.901 | HEK293 |
| ZEB1 | ZNF24 | 5388 | 14079 | 21595 | 0.309 | 90.343 | HEK293 |
| KLF9 | ZBTB26 | 9173 | 25095 | 35149 | 0.309 | 76.058 | HEK293 |
| ATF2 | ZBTB21 | 7718 | 30766 | 20325 | 0.309 | 74.128 | HEK293 |
| AEBP2 | ZNF266 | 1106 | 2455 | 5240 | 0.308 | 126.874 | HEK293 |
| ZNF404 | ZNF510 | 1102 | 2306 | 5540 | 0.308 | 128.594 | HEK293 |
| PRDM4 | TSHZ1 | 5070 | 22102 | 12240 | 0.308 | 89.505 | HEK293 |
| INSM2 | ZNF2 | 7011 | 17714 | 29224 | 0.308 | 75.789 | HEK293 |
| ZNF391 | ZSCAN21 | 5403 | 13956 | 22039 | 0.308 | 99.778 | HEK293 |
| ZNF2 | ZNF629 | 11012 | 29224 | 43843 | 0.308 | 55.180 | HEK293 |
| SP2 | ZBTB26 | 10227 | 31446 | 35149 | 0.308 | 53.625 | HEK293 |
| ZBTB10 | ZBTB7A | 4273 | 18260 | 10584 | 0.307 | 101.128 | HEK293 |

| | | | | | | | |
|---|---|---|---|---|---|---|---|
| ZFP69B | ZNF629 | 10029 | 24305 | 43843 | 0.307 | 53.676 | HEK293 |
| KLF8 | ZNF561 | 8577 | 32219 | 24194 | 0.307 | 68.979 | HEK293 |
| KLF16 | ZNF76 | 5505 | 18018 | 17824 | 0.307 | 91.064 | HEK293 |
| ZNF561 | ZNF692 | 8913 | 24194 | 34826 | 0.307 | 59.960 | HEK293 |
| SP7 | ZNF391 | 8128 | 50285 | 13956 | 0.307 | 68.892 | HEK293 |
| OVOL3 | ZXDB | 5774 | 9640 | 36901 | 0.306 | 83.152 | HEK293 |
| GFI1B | GLI2 | 2425 | 10973 | 5719 | 0.306 | 91.035 | HEK293 |
| TSHZ1 | ZNF391 | 3997 | 12240 | 13956 | 0.306 | 110.270 | HEK293 |
| OSR2 | TSHZ1 | 6401 | 35833 | 12240 | 0.306 | 73.281 | HEK293 |
| ZNF629 | ZSCAN21 | 9490 | 43843 | 22039 | 0.305 | 52.872 | HEK293 |
| ZSCAN30 | ZXDB | 9511 | 26304 | 36901 | 0.305 | 51.230 | HEK293 |
| TSHZ1 | ZBTB21 | 4815 | 12240 | 20325 | 0.305 | 82.229 | HEK293 |
| KLF16 | ZNF2 | 7002 | 18018 | 29224 | 0.305 | 77.052 | HEK293 |
| KLF16 | ZNF24 | 6012 | 18018 | 21595 | 0.305 | 83.196 | HEK293 |
| ZNF292 | ZNF34 | 1338 | 1922 | 10050 | 0.304 | 127.138 | HEK293 |
| ZNF391 | ZXDB | 6905 | 13956 | 36901 | 0.304 | 82.686 | HEK293 |
| KLF16 | ZFP69B | 6367 | 18018 | 24305 | 0.304 | 83.569 | HEK293 |
| WT1 | ZNF24 | 7587 | 28823 | 21595 | 0.304 | 67.431 | HEK293 |
| KLF7 | ZNF324 | 4425 | 12712 | 16663 | 0.304 | 106.008 | HEK293 |
| KLF16 | ZNF843 | 6544 | 18018 | 25751 | 0.304 | 76.796 | HEK293 |
| INSM2 | KLF10 | 6043 | 17714 | 22350 | 0.304 | 76.133 | HEK293 |
| BCL11A | KLF8 | 6497 | 14204 | 32219 | 0.304 | 77.893 | HEK293 |
| ZNF324 | ZNF76 | 5232 | 16663 | 17824 | 0.304 | 77.066 | HEK293 |
| ZNF202 | ZNF292 | 1088 | 6698 | 1922 | 0.303 | 152.044 | HEK293 |
| ZBTB21 | ZNF518A | 5898 | 20325 | 18621 | 0.303 | 85.421 | HEK293 |
| ZNF394 | ZNF623 | 6675 | 32293 | 15029 | 0.303 | 63.173 | HEK293 |
| KLF17 | KLF7 | 5804 | 28900 | 12712 | 0.303 | 68.958 | HEK293 |
| ZNF292 | ZNF530 | 631 | 1922 | 2262 | 0.303 | 113.348 | HEK293 |
| KLF10 | MAZ | 9360 | 22350 | 42804 | 0.303 | 56.717 | HEK293 |
| ZNF391 | ZNF692 | 6664 | 13956 | 34826 | 0.302 | 74.704 | HEK293 |
| KLF10 | ZNF843 | 7251 | 22350 | 25751 | 0.302 | 80.217 | HEK293 |
| KLF8 | ZNF501 | 8144 | 32219 | 22541 | 0.302 | 67.890 | HEK293 |
| OVOL3 | PRDM4 | 4410 | 9640 | 22102 | 0.302 | 95.662 | HEK293 |
| SALL2 | ZNF148 | 963 | 2650 | 3836 | 0.302 | 113.835 | HEK293 |
| ZNF560 | ZNF680 | 1851 | 3492 | 10768 | 0.302 | 147.715 | HEK293 |
| ATF2 | ZNF629 | 11086 | 30766 | 43843 | 0.302 | 50.387 | HEK293 |
| KLF8 | WT1 | 9185 | 32219 | 28823 | 0.301 | 52.633 | HEK293 |
| ZNF138 | ZNF19 | 554 | 1647 | 2055 | 0.301 | 122.642 | HEK293 |
| ZNF555 | ZNF680 | 2558 | 6703 | 10768 | 0.301 | 121.036 | HEK293 |
| KLF10 | ZNF660 | 8095 | 22350 | 32350 | 0.301 | 66.420 | HEK293 |
| ZBTB44 | ZNF561 | 6680 | 20353 | 24194 | 0.301 | 76.391 | HEK293 |
| ZNF140 | ZNF404 | 1291 | 7978 | 2306 | 0.301 | 122.507 | HEK293 |
| ZNF610 | ZNF843 | 5880 | 14846 | 25751 | 0.301 | 84.123 | HEK293 |
| ZBTB44 | ZNF843 | 6880 | 20353 | 25751 | 0.301 | 79.586 | HEK293 |
| ZBTB21 | ZXDB | 8230 | 20325 | 36901 | 0.301 | 55.457 | HEK293 |
| PATZ1 | ZFP69B | 9562 | 41663 | 24305 | 0.300 | 56.205 | HEK293 |
| KLF8 | TSHZ1 | 5963 | 32219 | 12240 | 0.300 | 65.469 | HEK293 |
| ZEB2 | ZNF76 | 8028 | 40116 | 17824 | 0.300 | 69.512 | HEK293 |
| JUN | JUND | 18801 | 22995 | 34106 | 0.671 | 173.426 | HeLa-S3 |
| MAFF | MAFK | 12021 | 25547 | 16601 | 0.584 | 136.704 | HeLa-S3 |
| NFYA | NFYB | 4131 | 7179 | 8033 | 0.544 | 194.226 | HeLa-S3 |
| FOS | JUN | 7804 | 10227 | 22995 | 0.509 | 187.956 | HeLa-S3 |

| | | | | | | | |
|---|---|---|---|---|---|---|---|
| JUN | STAT3 | 9270 | 22995 | 17088 | 0.468 | 125.151 | HeLa-S3 |
| FOS | JUND | 8621 | 10227 | 34106 | 0.462 | 122.582 | HeLa-S3 |
| MAX | MYC | 8704 | 34096 | 11821 | 0.434 | 108.792 | HeLa-S3 |
| JUND | STAT3 | 10369 | 34106 | 17088 | 0.430 | 80.718 | HeLa-S3 |
| ELK1 | ELK4 | 2811 | 6090 | 7250 | 0.423 | 141.927 | HeLa-S3 |
| MAX | MXI1 | 9623 | 34096 | 15255 | 0.422 | 103.843 | HeLa-S3 |
| ELK1 | GABPA | 2819 | 6090 | 7553 | 0.416 | 111.645 | HeLa-S3 |
| ELK4 | GABPA | 3038 | 7250 | 7553 | 0.411 | 116.017 | HeLa-S3 |
| IRF3 | NFYA | 1425 | 1848 | 7179 | 0.391 | 126.619 | HeLa-S3 |
| IRF3 | NFYB | 1429 | 1848 | 8033 | 0.371 | 140.041 | HeLa-S3 |
| MAX | MAZ | 8678 | 34096 | 16564 | 0.365 | 63.397 | HeLa-S3 |
| FOS | STAT3 | 4697 | 10227 | 17088 | 0.355 | 93.733 | HeLa-S3 |
| MXI1 | MYC | 4749 | 15255 | 11821 | 0.354 | 94.686 | HeLa-S3 |
| RFX5 | TBP | 7861 | 24378 | 23581 | 0.328 | 56.083 | HeLa-S3 |
| RFX5 | STAT3 | 6581 | 24378 | 17088 | 0.322 | 51.954 | HeLa-S3 |
| FOXA1 | FOXA2 | 40029 | 58770 | 48956 | 0.746 | 61.743 | HepG2 |
| SOX13 | SOX5 | 30692 | 50089 | 41113 | 0.676 | 50.224 | HepG2 |
| NR2F6 | RARA | 25665 | 39697 | 44851 | 0.608 | 50.925 | HepG2 |
| NFYB | NFYC | 9642 | 15448 | 16430 | 0.605 | 159.930 | HepG2 |
| FOXA3 | FOXP1 | 21688 | 47916 | 28473 | 0.587 | 56.148 | HepG2 |
| FOXP1 | FOXP4 | 18581 | 28473 | 35653 | 0.583 | 65.819 | HepG2 |
| JUN | JUND | 16036 | 18375 | 41511 | 0.581 | 72.573 | HepG2 |
| ETV4 | ETV5 | 15113 | 21788 | 32099 | 0.571 | 85.483 | HepG2 |
| HNF4A | HNF4G | 20715 | 53121 | 24914 | 0.569 | 64.522 | HepG2 |
| TFDP1 | TFDP2 | 12872 | 23515 | 22516 | 0.559 | 84.727 | HepG2 |
| FOXP1 | SOX5 | 19096 | 28473 | 41113 | 0.558 | 57.631 | HepG2 |
| FOXP4 | SOX5 | 21285 | 35653 | 41113 | 0.556 | 50.948 | HepG2 |
| RXRB | THRA | 19065 | 41739 | 28190 | 0.556 | 55.849 | HepG2 |
| HMG20A | SOX5 | 18227 | 26787 | 41113 | 0.549 | 58.151 | HepG2 |
| CEBPD | CEBPG | 15790 | 19702 | 42425 | 0.546 | 80.301 | HepG2 |
| PPARG | RXRB | 17984 | 26097 | 41739 | 0.545 | 57.834 | HepG2 |
| FOXP1 | SOX13 | 20539 | 28473 | 50089 | 0.544 | 51.167 | HepG2 |
| TCF7 | TCF7L2 | 14501 | 19413 | 36713 | 0.543 | 76.552 | HepG2 |
| KLF16 | THAP11 | 18357 | 30628 | 37306 | 0.543 | 51.087 | HepG2 |
| FOXP1 | HMG20A | 14762 | 28473 | 26787 | 0.535 | 70.238 | HepG2 |
| IRF2 | THAP11 | 18195 | 31142 | 37306 | 0.534 | 50.361 | HepG2 |
| ERF | IRF2 | 16957 | 32451 | 31142 | 0.533 | 63.155 | HepG2 |
| FOXK1 | FOXO1 | 13880 | 26575 | 25752 | 0.531 | 65.333 | HepG2 |
| E2F4 | TFDP2 | 11757 | 21936 | 22516 | 0.529 | 87.141 | HepG2 |
| FOXP4 | ISL2 | 17005 | 35653 | 29175 | 0.527 | 51.977 | HepG2 |
| RXRB | ZNF331 | 19432 | 41739 | 32566 | 0.527 | 52.458 | HepG2 |
| IRF2 | RXRB | 18984 | 31142 | 41739 | 0.527 | 50.367 | HepG2 |
| FOXK1 | THAP11 | 16574 | 26575 | 37306 | 0.526 | 65.008 | HepG2 |
| FOXO1 | SMAD4 | 17992 | 25752 | 45641 | 0.525 | 52.198 | HepG2 |
| FOXO1 | FOXP4 | 15889 | 25752 | 35653 | 0.524 | 60.604 | HepG2 |
| FOXO1 | ZNF331 | 15181 | 25752 | 32566 | 0.524 | 73.343 | HepG2 |
| NFYA | NFYC | 8865 | 17604 | 16430 | 0.521 | 82.759 | HepG2 |
| FOXO1 | RXRB | 17049 | 25752 | 41739 | 0.520 | 53.886 | HepG2 |
| FOXO1 | LCOR | 15984 | 25752 | 36711 | 0.520 | 60.288 | HepG2 |
| IRF2 | ZNF331 | 16484 | 31142 | 32566 | 0.518 | 59.380 | HepG2 |
| NFYA | NFYB | 8512 | 17604 | 15448 | 0.516 | 95.749 | HepG2 |
| FOXA3 | FOXO1 | 18097 | 47916 | 25752 | 0.515 | 50.026 | HepG2 |

| | | | | | | | |
|---|---|---|---|---|---|---|---|
| TEAD3 | TEAD4 | 16038 | 45689 | 21380 | 0.513 | 56.079 | HepG2 |
| IKZF5 | KLF16 | 15806 | 31009 | 30628 | 0.513 | 55.947 | HepG2 |
| HMG20A | TEAD1 | 15874 | 26787 | 35808 | 0.513 | 53.894 | HepG2 |
| IKZF5 | IRF2 | 15826 | 31009 | 31142 | 0.509 | 57.340 | HepG2 |
| ATF2 | THAP11 | 15110 | 23795 | 37306 | 0.507 | 52.831 | HepG2 |
| NR2F6 | THRA | 16960 | 39697 | 28190 | 0.507 | 51.437 | HepG2 |
| ELF1 | ETV5 | 13716 | 22892 | 32099 | 0.506 | 64.266 | HepG2 |
| ERF | RBPJ | 15648 | 32451 | 29519 | 0.506 | 63.368 | HepG2 |
| THRA | ZNF331 | 15296 | 28190 | 32566 | 0.505 | 61.122 | HepG2 |
| ATF2 | ZNF331 | 14004 | 23795 | 32566 | 0.503 | 60.583 | HepG2 |
| IRF2 | RBPJ | 15223 | 31142 | 29519 | 0.502 | 52.537 | HepG2 |
| FOXK1 | IRF2 | 14436 | 26575 | 31142 | 0.502 | 59.147 | HepG2 |
| IRF2 | ZNF614 | 13969 | 31142 | 25059 | 0.500 | 57.225 | HepG2 |
| ERF | KLF16 | 15660 | 32451 | 30628 | 0.497 | 58.645 | HepG2 |
| FOXO1 | THRA | 13382 | 25752 | 28190 | 0.497 | 62.695 | HepG2 |
| CREB1 | CREM | 14010 | 26848 | 29846 | 0.495 | 50.271 | HepG2 |
| SIX1 | SIX4 | 7024 | 14572 | 13829 | 0.495 | 106.576 | HepG2 |
| ATF2 | IRF2 | 13469 | 23795 | 31142 | 0.495 | 64.286 | HepG2 |
| ZNF217 | ZNF609 | 13915 | 31645 | 25025 | 0.494 | 63.644 | HepG2 |
| FOXO1 | FOXP1 | 13389 | 25752 | 28473 | 0.494 | 55.301 | HepG2 |
| IRF2 | KLF16 | 15250 | 31142 | 30628 | 0.494 | 63.354 | HepG2 |
| FOXK1 | FOXP4 | 15159 | 26575 | 35653 | 0.492 | 55.047 | HepG2 |
| ATF2 | FOXK1 | 12347 | 23795 | 26575 | 0.491 | 88.658 | HepG2 |
| ATF2 | KLF16 | 13255 | 23795 | 30628 | 0.491 | 63.838 | HepG2 |
| ISL2 | ZNF503 | 11383 | 29175 | 18529 | 0.490 | 73.894 | HepG2 |
| ERF | ZNF792 | 13486 | 32451 | 23423 | 0.489 | 59.180 | HepG2 |
| FOXO1 | IRF2 | 13844 | 25752 | 31142 | 0.489 | 57.851 | HepG2 |
| IKZF5 | RBPJ | 14776 | 31009 | 29519 | 0.488 | 51.777 | HepG2 |
| FOXO1 | ZNF217 | 13899 | 25752 | 31645 | 0.487 | 50.301 | HepG2 |
| SPEN | ZNF574 | 15266 | 27137 | 36345 | 0.486 | 53.363 | HepG2 |
| ELF3 | FOXK1 | 15793 | 39910 | 26575 | 0.485 | 50.945 | HepG2 |
| IRF2 | ZNF792 | 13079 | 31142 | 23423 | 0.484 | 71.602 | HepG2 |
| ATF2 | LCOR | 14309 | 23795 | 36711 | 0.484 | 52.564 | HepG2 |
| FOXJ3 | ISL2 | 11994 | 21114 | 29175 | 0.483 | 65.753 | HepG2 |
| ATF2 | IKZF5 | 13090 | 23795 | 31009 | 0.482 | 71.210 | HepG2 |
| ATF2 | FOXO1 | 11925 | 23795 | 25752 | 0.482 | 67.167 | HepG2 |
| NR2F6 | PPARG | 15480 | 39697 | 26097 | 0.481 | 53.544 | HepG2 |
| ATF2 | THRA | 12441 | 23795 | 28190 | 0.480 | 59.543 | HepG2 |
| THRA | THRB | 10424 | 28190 | 16710 | 0.480 | 74.858 | HepG2 |
| KLF16 | SMAD3 | 12109 | 30628 | 20760 | 0.480 | 60.981 | HepG2 |
| ELF3 | HOMEZ | 15875 | 39910 | 27410 | 0.480 | 50.607 | HepG2 |
| FOXO1 | IKZF5 | 13558 | 25752 | 31009 | 0.480 | 50.793 | HepG2 |
| E2F4 | TFDP1 | 10871 | 21936 | 23515 | 0.479 | 59.392 | HepG2 |
| TEAD1 | TEAD4 | 13223 | 35808 | 21380 | 0.478 | 56.042 | HepG2 |
| DRAP1 | IRF2 | 13630 | 26223 | 31142 | 0.477 | 53.065 | HepG2 |
| ETV5 | FOXO1 | 13710 | 32099 | 25752 | 0.477 | 51.291 | HepG2 |
| FOXP4 | ZNF609 | 14237 | 35653 | 25025 | 0.477 | 52.430 | HepG2 |
| MLX | TFE3 | 9343 | 16042 | 23975 | 0.476 | 83.128 | HepG2 |
| RBPJ | ZNF614 | 12957 | 29519 | 25059 | 0.476 | 62.345 | HepG2 |
| RFX1 | RFX3 | 3160 | 6690 | 6577 | 0.476 | 110.611 | HepG2 |
| ERF | ZNF580 | 13060 | 32451 | 23174 | 0.476 | 62.539 | HepG2 |
| MIXL1 | PPARG | 12868 | 28115 | 26097 | 0.475 | 52.942 | HepG2 |

| | | | | | | | |
|---|---|---|---|---|---|---|---|
| ELF1 | THAP11 | 13870 | 22892 | 37306 | 0.475 | 56.524 | HepG2 |
| IRF2 | THRA | 14059 | 31142 | 28190 | 0.474 | 52.499 | HepG2 |
| KLF16 | RBPJ | 14156 | 30628 | 29519 | 0.471 | 54.654 | HepG2 |
| ATF2 | HOXA3 | 14561 | 23795 | 40277 | 0.470 | 58.445 | HepG2 |
| SMAD3 | THAP11 | 13059 | 20760 | 37306 | 0.469 | 53.789 | HepG2 |
| ZNF614 | ZNF792 | 11368 | 25059 | 23423 | 0.469 | 59.934 | HepG2 |
| MIXL1 | TEAD1 | 14879 | 28115 | 35808 | 0.469 | 55.955 | HepG2 |
| ERF | ZSCAN9 | 12011 | 32451 | 20246 | 0.469 | 67.172 | HepG2 |
| ZNF331 | ZNF614 | 13384 | 32566 | 25059 | 0.469 | 53.538 | HepG2 |
| ATF2 | SMAD3 | 10399 | 23795 | 20760 | 0.468 | 68.046 | HepG2 |
| FOXK1 | FOXP1 | 12860 | 26575 | 28473 | 0.468 | 64.739 | HepG2 |
| HNF1A | ZNF609 | 11711 | 25102 | 25025 | 0.467 | 50.833 | HepG2 |
| ATF2 | MYBL2 | 11250 | 23795 | 24417 | 0.467 | 66.398 | HepG2 |
| ERF | ZBTB25 | 12017 | 32451 | 20508 | 0.466 | 57.707 | HepG2 |
| FOXJ3 | FOXP4 | 12779 | 21114 | 35653 | 0.466 | 54.890 | HepG2 |
| IRF2 | TFE3 | 12718 | 31142 | 23975 | 0.465 | 54.780 | HepG2 |
| ATF2 | ERF | 12920 | 23795 | 32451 | 0.465 | 52.600 | HepG2 |
| ZNF331 | ZNF792 | 12826 | 32566 | 23423 | 0.464 | 60.230 | HepG2 |
| HOMEZ | MIXL1 | 12883 | 27410 | 28115 | 0.464 | 58.009 | HepG2 |
| FOXP1 | TCF7 | 10887 | 28473 | 19413 | 0.463 | 82.778 | HepG2 |
| RBPJ | ZNF792 | 12166 | 29519 | 23423 | 0.463 | 56.000 | HepG2 |
| FOXK1 | THRA | 12661 | 26575 | 28190 | 0.463 | 55.492 | HepG2 |
| ATF2 | ISL2 | 12177 | 23795 | 29175 | 0.462 | 55.392 | HepG2 |
| ISL2 | ZNF331 | 14213 | 29175 | 32566 | 0.461 | 53.067 | HepG2 |
| FOXK1 | SMAD3 | 10824 | 26575 | 20760 | 0.461 | 64.814 | HepG2 |
| IKZF5 | SMAD3 | 11686 | 31009 | 20760 | 0.461 | 51.406 | HepG2 |
| IRF2 | SMAD3 | 11705 | 31142 | 20760 | 0.460 | 64.898 | HepG2 |
| NR2F2 | NR2F6 | 12349 | 18140 | 39697 | 0.460 | 51.729 | HepG2 |
| SMAD3 | ZNF331 | 11918 | 20760 | 32566 | 0.458 | 53.056 | HepG2 |
| ERF | TFE3 | 12773 | 32451 | 23975 | 0.458 | 54.426 | HepG2 |
| IRF2 | ZBTB25 | 11537 | 31142 | 20508 | 0.457 | 64.028 | HepG2 |
| ATF2 | RBPJ | 12094 | 23795 | 29519 | 0.456 | 58.775 | HepG2 |
| IRF2 | ZSCAN9 | 11443 | 31142 | 20246 | 0.456 | 50.741 | HepG2 |
| ELF1 | ETV4 | 10174 | 22892 | 21788 | 0.456 | 58.677 | HepG2 |
| ISL2 | THRA | 13053 | 29175 | 28190 | 0.455 | 55.355 | HepG2 |
| HMG20A | TCF7 | 10378 | 26787 | 19413 | 0.455 | 63.700 | HepG2 |
| SOX5 | TFE3 | 14273 | 41113 | 23975 | 0.455 | 51.312 | HepG2 |
| KLF16 | ZBTB25 | 11389 | 30628 | 20508 | 0.454 | 64.961 | HepG2 |
| FOXJ3 | ZNF503 | 8976 | 21114 | 18529 | 0.454 | 81.682 | HepG2 |
| TFE3 | ZNF331 | 12675 | 23975 | 32566 | 0.454 | 57.864 | HepG2 |
| ERF | ZNF511 | 10867 | 32451 | 17697 | 0.453 | 68.702 | HepG2 |
| NR2F1 | NR2F2 | 8305 | 18530 | 18140 | 0.453 | 78.258 | HepG2 |
| ZNF792 | ZSCAN9 | 9854 | 23423 | 20246 | 0.453 | 64.898 | HepG2 |
| KMT2A | KMT2B | 10105 | 23007 | 21702 | 0.452 | 50.703 | HepG2 |
| MIXL1 | ZNF614 | 11986 | 28115 | 25059 | 0.452 | 55.322 | HepG2 |
| IRF2 | PPARG | 12858 | 31142 | 26097 | 0.451 | 51.277 | HepG2 |
| THAP9 | ZNF350 | 10731 | 21105 | 26991 | 0.450 | 55.816 | HepG2 |
| ATF2 | ZNF614 | 10975 | 23795 | 25059 | 0.449 | 55.708 | HepG2 |
| DLX6 | ISL2 | 10756 | 19647 | 29175 | 0.449 | 58.916 | HepG2 |
| FOXO1 | MLX | 9092 | 25752 | 16042 | 0.447 | 64.698 | HepG2 |
| TFDP2 | THAP11 | 12964 | 22516 | 37306 | 0.447 | 50.361 | HepG2 |
| IRF2 | MYBL2 | 12330 | 31142 | 24417 | 0.447 | 56.949 | HepG2 |

| | | | | | | | |
|---|---|---|---|---|---|---|---|
| THAP11 | ZBTB25 | 12357 | 37306 | 20508 | 0.447 | 50.568 | HepG2 |
| FOXK1 | MIXL1 | 12208 | 26575 | 28115 | 0.447 | 51.547 | HepG2 |
| IKZF5 | MYBL2 | 12281 | 31009 | 24417 | 0.446 | 50.226 | HepG2 |
| IRF2 | ZNF511 | 10475 | 31142 | 17697 | 0.446 | 55.354 | HepG2 |
| E2F4 | NFYA | 8767 | 21936 | 17604 | 0.446 | 76.636 | HepG2 |
| ZKSCAN8 | ZNF614 | 9725 | 18988 | 25059 | 0.446 | 61.977 | HepG2 |
| KLF16 | ZSCAN9 | 11101 | 30628 | 20246 | 0.446 | 56.262 | HepG2 |
| TFE3 | ZNF792 | 10561 | 23975 | 23423 | 0.446 | 53.797 | HepG2 |
| ERF | KMT2B | 11817 | 32451 | 21702 | 0.445 | 62.679 | HepG2 |
| FOXO1 | ZNF614 | 11309 | 25752 | 25059 | 0.445 | 51.064 | HepG2 |
| PPARG | THRA | 12066 | 26097 | 28190 | 0.445 | 53.019 | HepG2 |
| FOXK1 | TFE3 | 11222 | 26575 | 23975 | 0.445 | 51.732 | HepG2 |
| ZNF362 | ZNF384 | 9614 | 25385 | 18424 | 0.445 | 72.882 | HepG2 |
| DRAP1 | ZNF792 | 11012 | 26223 | 23423 | 0.444 | 62.869 | HepG2 |
| ZNF511 | ZNF792 | 9043 | 17697 | 23423 | 0.444 | 81.740 | HepG2 |
| DRAP1 | TBP | 9716 | 26223 | 18258 | 0.444 | 70.325 | HepG2 |
| ZNF511 | ZNF614 | 9333 | 17697 | 25059 | 0.443 | 67.530 | HepG2 |
| ISL2 | MYBL2 | 11814 | 29175 | 24417 | 0.443 | 54.728 | HepG2 |
| KLF16 | KMT2B | 11407 | 30628 | 21702 | 0.442 | 52.557 | HepG2 |
| ZNF580 | ZNF792 | 10308 | 23174 | 23423 | 0.442 | 56.258 | HepG2 |
| IRF2 | KMT2B | 11497 | 31142 | 21702 | 0.442 | 68.430 | HepG2 |
| FOXO1 | TFE3 | 10970 | 25752 | 23975 | 0.441 | 57.707 | HepG2 |
| ATF2 | ZNF792 | 10419 | 23795 | 23423 | 0.441 | 51.022 | HepG2 |
| FOXK1 | HOMEZ | 11902 | 26575 | 27410 | 0.441 | 58.042 | HepG2 |
| IRF2 | ZKSCAN8 | 10721 | 31142 | 18988 | 0.441 | 60.086 | HepG2 |
| ZBTB25 | ZNF614 | 9994 | 20508 | 25059 | 0.441 | 54.330 | HepG2 |
| RBPJ | ZBTB25 | 10844 | 29519 | 20508 | 0.441 | 51.122 | HepG2 |
| CREM | TCF7 | 10605 | 29846 | 19413 | 0.441 | 64.831 | HepG2 |
| ZNF614 | ZSCAN9 | 9908 | 25059 | 20246 | 0.440 | 51.320 | HepG2 |
| ATF2 | TFE3 | 10499 | 23795 | 23975 | 0.440 | 61.232 | HepG2 |
| ISL2 | PPARG | 12108 | 29175 | 26097 | 0.439 | 58.201 | HepG2 |
| TFE3 | ZSCAN9 | 9663 | 23975 | 20246 | 0.439 | 59.258 | HepG2 |
| IRF2 | MLX | 9803 | 31142 | 16042 | 0.439 | 51.178 | HepG2 |
| AHDC1 | FOXP1 | 12124 | 26848 | 28473 | 0.439 | 55.028 | HepG2 |
| HOMEZ | PPARG | 11727 | 27410 | 26097 | 0.438 | 50.769 | HepG2 |
| ATF2 | PPARG | 10911 | 23795 | 26097 | 0.438 | 57.197 | HepG2 |
| ZBTB25 | ZSCAN9 | 8920 | 20508 | 20246 | 0.438 | 62.545 | HepG2 |
| KLF16 | TFE3 | 11855 | 30628 | 23975 | 0.437 | 52.764 | HepG2 |
| FOXO1 | HOMEZ | 11616 | 25752 | 27410 | 0.437 | 55.453 | HepG2 |
| E2F1 | E2F4 | 7042 | 11827 | 21936 | 0.437 | 71.965 | HepG2 |
| FOXO1 | SMAD3 | 10107 | 25752 | 20760 | 0.437 | 52.930 | HepG2 |
| ZNF580 | ZNF614 | 10523 | 23174 | 25059 | 0.437 | 53.409 | HepG2 |
| SMAD3 | THRA | 10561 | 20760 | 28190 | 0.437 | 54.131 | HepG2 |
| ZBTB25 | ZNF792 | 9558 | 20508 | 23423 | 0.436 | 53.705 | HepG2 |
| KLF16 | KLF6 | 10376 | 30628 | 18502 | 0.436 | 57.019 | HepG2 |
| ERF | ZKSCAN8 | 10809 | 32451 | 18988 | 0.435 | 54.536 | HepG2 |
| TFE3 | ZNF614 | 10672 | 23975 | 25059 | 0.435 | 60.768 | HepG2 |
| ERF | ZNF205 | 11085 | 32451 | 20027 | 0.435 | 58.382 | HepG2 |
| FOXP4 | ZNF503 | 11158 | 35653 | 18529 | 0.434 | 50.278 | HepG2 |
| ZNF511 | ZSCAN9 | 8217 | 17697 | 20246 | 0.434 | 68.249 | HepG2 |
| NRL | ZNF891 | 8196 | 19698 | 18098 | 0.434 | 67.893 | HepG2 |
| MLX | SOX5 | 11147 | 16042 | 41113 | 0.434 | 58.398 | HepG2 |

| | | | | | | | |
|---|---|---|---|---|---|---|---|
| MXI1 | TFDP2 | 10373 | 25399 | 22516 | 0.434 | 63.113 | HepG2 |
| FOXO1 | ZNF792 | 10628 | 25752 | 23423 | 0.433 | 53.004 | HepG2 |
| GMEB1 | SPEN | 10477 | 21627 | 27137 | 0.432 | 57.392 | HepG2 |
| DLX6 | ZNF503 | 8251 | 19647 | 18529 | 0.432 | 66.675 | HepG2 |
| ATF2 | ZBTB25 | 9546 | 23795 | 20508 | 0.432 | 76.004 | HepG2 |
| IKZF5 | ZSCAN9 | 10826 | 31009 | 20246 | 0.432 | 51.297 | HepG2 |
| RBPJ | ZNF511 | 9849 | 29519 | 17697 | 0.431 | 54.342 | HepG2 |
| SPEN | TFDP2 | 10646 | 27137 | 22516 | 0.431 | 53.874 | HepG2 |
| MLX | PPARG | 8806 | 16042 | 26097 | 0.430 | 62.762 | HepG2 |
| MAZ | ZSCAN31 | 10465 | 27510 | 21564 | 0.430 | 52.184 | HepG2 |
| MYBL2 | SMAD3 | 9673 | 24417 | 20760 | 0.430 | 57.389 | HepG2 |
| TFE3 | THRA | 11161 | 23975 | 28190 | 0.429 | 50.135 | HepG2 |
| FOXO1 | RREB1 | 10347 | 25752 | 22566 | 0.429 | 53.865 | HepG2 |
| LCOR | SMAD3 | 11839 | 36711 | 20760 | 0.429 | 52.107 | HepG2 |
| DRAP1 | FOXK1 | 11311 | 26223 | 26575 | 0.428 | 51.557 | HepG2 |
| THRA | ZNF614 | 11382 | 28190 | 25059 | 0.428 | 54.580 | HepG2 |
| RBPJ | ZKSCAN8 | 10134 | 29519 | 18988 | 0.428 | 63.247 | HepG2 |
| MIXL1 | TFE3 | 11108 | 28115 | 23975 | 0.428 | 56.109 | HepG2 |
| ELF1 | FOXK1 | 10551 | 22892 | 26575 | 0.428 | 63.828 | HepG2 |
| FOXK1 | MLX | 8832 | 26575 | 16042 | 0.428 | 64.956 | HepG2 |
| ZNF205 | ZNF792 | 9264 | 20027 | 23423 | 0.428 | 58.641 | HepG2 |
| ATF2 | MLX | 8349 | 23795 | 16042 | 0.427 | 69.153 | HepG2 |
| FOXJ3 | FOXP1 | 10476 | 21114 | 28473 | 0.427 | 52.774 | HepG2 |
| MLX | ZNF331 | 9761 | 16042 | 32566 | 0.427 | 68.970 | HepG2 |
| FOXP4 | TCF7 | 11234 | 35653 | 19413 | 0.427 | 54.705 | HepG2 |
| ZKSCAN8 | ZNF792 | 9004 | 18988 | 23423 | 0.427 | 54.521 | HepG2 |
| GMEB1 | TFDP2 | 9419 | 21627 | 22516 | 0.427 | 53.629 | HepG2 |
| ZNF205 | ZNF614 | 9562 | 20027 | 25059 | 0.427 | 53.575 | HepG2 |
| KMT2B | MXD4 | 9361 | 21702 | 22169 | 0.427 | 54.906 | HepG2 |
| ZBTB25 | ZNF511 | 8126 | 20508 | 17697 | 0.427 | 62.676 | HepG2 |
| NRL | ZNF280B | 7494 | 19698 | 15695 | 0.426 | 73.671 | HepG2 |
| ATF2 | ZSCAN9 | 9353 | 23795 | 20246 | 0.426 | 60.518 | HepG2 |
| ERF | KLF6 | 10439 | 32451 | 18502 | 0.426 | 59.693 | HepG2 |
| MIXL1 | MLX | 9041 | 28115 | 16042 | 0.426 | 57.712 | HepG2 |
| SP1 | SP5 | 8544 | 16039 | 25217 | 0.425 | 51.180 | HepG2 |
| IKZF5 | ZBTB25 | 10711 | 31009 | 20508 | 0.425 | 56.403 | HepG2 |
| SMAD3 | ZNF614 | 9672 | 20760 | 25059 | 0.424 | 54.859 | HepG2 |
| KMT2B | ZBTB25 | 8946 | 21702 | 20508 | 0.424 | 58.876 | HepG2 |
| ATF2 | DRAP1 | 10586 | 23795 | 26223 | 0.424 | 53.390 | HepG2 |
| MYBL2 | TFDP2 | 9925 | 24417 | 22516 | 0.423 | 55.320 | HepG2 |
| MLX | ZNF614 | 8483 | 16042 | 25059 | 0.423 | 64.194 | HepG2 |
| SMAD3 | ZBTB25 | 8722 | 20760 | 20508 | 0.423 | 63.101 | HepG2 |
| GMEB1 | KMT2B | 9145 | 21627 | 21702 | 0.422 | 57.748 | HepG2 |
| HOMEZ | ZNF644 | 10858 | 27410 | 24199 | 0.422 | 52.256 | HepG2 |
| RBPJ | SMAD3 | 10435 | 29519 | 20760 | 0.422 | 53.166 | HepG2 |
| MLX | ZNF792 | 8170 | 16042 | 23423 | 0.421 | 65.996 | HepG2 |
| ATF1 | CREB1 | 8166 | 13987 | 26848 | 0.421 | 72.151 | HepG2 |
| SMAD3 | ZSCAN9 | 8628 | 20760 | 20246 | 0.421 | 59.294 | HepG2 |
| HMG20A | MLX | 8724 | 26787 | 16042 | 0.421 | 66.337 | HepG2 |
| ZKSCAN8 | ZSCAN9 | 8245 | 18988 | 20246 | 0.421 | 64.090 | HepG2 |
| FOXO1 | MYBL2 | 10540 | 25752 | 24417 | 0.420 | 50.336 | HepG2 |
| MLX | RBPJ | 9141 | 16042 | 29519 | 0.420 | 52.820 | HepG2 |

| | | | | | | | |
|---|---|---|---|---|---|---|---|
| ATF2 | ZNF511 | 8617 | 23795 | 17697 | 0.420 | 59.850 | HepG2 |
| KMT2B | ZNF580 | 9407 | 21702 | 23174 | 0.419 | 68.524 | HepG2 |
| KDM5B | MXD1 | 9550 | 28623 | 18125 | 0.419 | 59.947 | HepG2 |
| ATF1 | CREM | 8563 | 13987 | 29846 | 0.419 | 66.726 | HepG2 |
| MIXL1 | ZKSCAN8 | 9683 | 28115 | 18988 | 0.419 | 60.733 | HepG2 |
| MIXL1 | SMAD3 | 10118 | 28115 | 20760 | 0.419 | 51.088 | HepG2 |
| GMEB1 | PHF20 | 8174 | 21627 | 17625 | 0.419 | 58.746 | HepG2 |
| ZKSCAN8 | ZNF511 | 7673 | 18988 | 17697 | 0.419 | 72.727 | HepG2 |
| ZNF503 | ZNF609 | 9011 | 18529 | 25025 | 0.418 | 52.818 | HepG2 |
| KMT2B | ZNF792 | 9433 | 21702 | 23423 | 0.418 | 56.347 | HepG2 |
| TFE3 | ZNF511 | 8609 | 23975 | 17697 | 0.418 | 65.636 | HepG2 |
| ZBTB25 | ZNF580 | 9107 | 20508 | 23174 | 0.418 | 56.679 | HepG2 |
| DLX6 | FOXJ3 | 8506 | 19647 | 21114 | 0.418 | 54.908 | HepG2 |
| KLF16 | ZNF511 | 9718 | 30628 | 17697 | 0.417 | 51.706 | HepG2 |
| KLF6 | KMT2B | 8364 | 18502 | 21702 | 0.417 | 54.768 | HepG2 |
| DRAP1 | ELF1 | 10222 | 26223 | 22892 | 0.417 | 56.176 | HepG2 |
| ZNF580 | ZSCAN9 | 9027 | 23174 | 20246 | 0.417 | 50.599 | HepG2 |
| ATF2 | ZNF609 | 10163 | 23795 | 25025 | 0.416 | 57.501 | HepG2 |
| ATF2 | HOMEZ | 10620 | 23795 | 27410 | 0.416 | 50.278 | HepG2 |
| MIXL1 | ZSCAN9 | 9920 | 28115 | 20246 | 0.416 | 51.313 | HepG2 |
| KLF6 | ZBTB25 | 8099 | 18502 | 20508 | 0.416 | 61.230 | HepG2 |
| NFYA | SP1 | 6978 | 17604 | 16039 | 0.415 | 86.287 | HepG2 |
| ATF2 | ZKSCAN8 | 8822 | 23795 | 18988 | 0.415 | 66.871 | HepG2 |
| IKZF5 | MLX | 9251 | 31009 | 16042 | 0.415 | 66.055 | HepG2 |
| ISL2 | TCF7 | 9870 | 29175 | 19413 | 0.415 | 58.441 | HepG2 |
| ARNTL | ZNF609 | 9359 | 20368 | 25025 | 0.415 | 64.607 | HepG2 |
| ATF2 | ZNF124 | 8091 | 23795 | 16017 | 0.414 | 60.149 | HepG2 |
| ZKSCAN8 | ZNF331 | 10297 | 18988 | 32566 | 0.414 | 62.671 | HepG2 |
| FOXJ3 | FOXO1 | 9655 | 21114 | 25752 | 0.414 | 55.397 | HepG2 |
| ELF1 | KLF16 | 10962 | 22892 | 30628 | 0.414 | 55.395 | HepG2 |
| HMG20B | MLX | 6126 | 13676 | 16042 | 0.414 | 69.174 | HepG2 |
| SMAD3 | ZNF792 | 9120 | 20760 | 23423 | 0.414 | 52.544 | HepG2 |
| DRAP1 | SP1 | 8478 | 26223 | 16039 | 0.413 | 54.681 | HepG2 |
| IKZF5 | ZKSCAN8 | 10031 | 31009 | 18988 | 0.413 | 51.828 | HepG2 |
| KLF16 | ZNF124 | 9155 | 30628 | 16017 | 0.413 | 77.988 | HepG2 |
| FOXO1 | ZKSCAN8 | 9137 | 25752 | 18988 | 0.413 | 53.229 | HepG2 |
| CREM | NFIC | 9365 | 29846 | 17240 | 0.413 | 51.513 | HepG2 |
| KMT2B | TFDP2 | 9126 | 21702 | 22516 | 0.413 | 61.632 | HepG2 |
| HOXA5 | ISL2 | 10193 | 20899 | 29175 | 0.413 | 55.082 | HepG2 |
| DRAP1 | ZNF511 | 8891 | 26223 | 17697 | 0.413 | 53.639 | HepG2 |
| ZBTB25 | ZNF639 | 6846 | 20508 | 13418 | 0.413 | 76.872 | HepG2 |
| ATF2 | SIX4 | 7483 | 23795 | 13829 | 0.413 | 70.846 | HepG2 |
| SMAD3 | TFE3 | 9201 | 20760 | 23975 | 0.412 | 52.266 | HepG2 |
| FOXJ3 | HMG20A | 9798 | 21114 | 26787 | 0.412 | 52.003 | HepG2 |
| SP1 | TFDP2 | 7824 | 16039 | 22516 | 0.412 | 51.813 | HepG2 |
| MXI1 | SP1 | 8308 | 25399 | 16039 | 0.412 | 72.874 | HepG2 |
| MLX | THRA | 8753 | 16042 | 28190 | 0.412 | 50.802 | HepG2 |
| E2F4 | GMEB1 | 8959 | 21936 | 21627 | 0.411 | 63.681 | HepG2 |
| MXD4 | ZNF580 | 9318 | 22169 | 23174 | 0.411 | 52.585 | HepG2 |
| PPARG | ZKSCAN8 | 9140 | 26097 | 18988 | 0.411 | 57.055 | HepG2 |
| DLX6 | MIXL1 | 9649 | 19647 | 28115 | 0.411 | 52.448 | HepG2 |
| HMG20A | HMG20B | 7857 | 26787 | 13676 | 0.411 | 62.196 | HepG2 |

| | | | | | | | |
|---|---|---|---|---|---|---|---|
| ZNF205 | ZSCAN9 | 8264 | 20027 | 20246 | 0.410 | 52.568 | HepG2 |
| KLF6 | ZSCAN9 | 7942 | 18502 | 20246 | 0.410 | 53.834 | HepG2 |
| ATF2 | HNF1A | 10026 | 23795 | 25102 | 0.410 | 51.791 | HepG2 |
| IKZF5 | ZNF124 | 9123 | 31009 | 16017 | 0.409 | 54.512 | HepG2 |
| MLX | THAP11 | 10004 | 16042 | 37306 | 0.409 | 56.230 | HepG2 |
| ZBTB25 | ZKSCAN8 | 8056 | 20508 | 18988 | 0.408 | 56.149 | HepG2 |
| HOMEZ | MLX | 8560 | 27410 | 16042 | 0.408 | 67.367 | HepG2 |
| E2F4 | SP1 | 7655 | 21936 | 16039 | 0.408 | 57.090 | HepG2 |
| MLX | ZKSCAN8 | 7120 | 16042 | 18988 | 0.408 | 70.488 | HepG2 |
| ZNF205 | ZNF580 | 8781 | 20027 | 23174 | 0.408 | 62.816 | HepG2 |
| FOXO1 | HMG20B | 7647 | 25752 | 13676 | 0.407 | 52.382 | HepG2 |
| ZNF511 | ZNF580 | 8240 | 17697 | 23174 | 0.407 | 58.517 | HepG2 |
| ETV5 | MLX | 9229 | 32099 | 16042 | 0.407 | 54.950 | HepG2 |
| ZBTB25 | ZNF205 | 8238 | 20508 | 20027 | 0.406 | 58.972 | HepG2 |
| DRAP1 | TFDP2 | 9874 | 26223 | 22516 | 0.406 | 53.926 | HepG2 |
| KMT2B | MYBL2 | 9354 | 21702 | 24417 | 0.406 | 55.156 | HepG2 |
| ATF2 | ZNF205 | 8870 | 23795 | 20027 | 0.406 | 66.553 | HepG2 |
| NR2F2 | ZNF609 | 8649 | 18140 | 25025 | 0.406 | 50.277 | HepG2 |
| ZNF205 | ZNF511 | 7627 | 20027 | 17697 | 0.405 | 57.302 | HepG2 |
| FOXO1 | SIX4 | 7644 | 25752 | 13829 | 0.405 | 68.347 | HepG2 |
| PHF20 | SPEN | 8854 | 17625 | 27137 | 0.405 | 65.034 | HepG2 |
| MLX | ZNF511 | 6820 | 16042 | 17697 | 0.405 | 68.836 | HepG2 |
| MLX | ZNF609 | 8107 | 16042 | 25025 | 0.405 | 59.324 | HepG2 |
| MLX | ZNF217 | 9114 | 16042 | 31645 | 0.405 | 51.223 | HepG2 |
| FOXK1 | ZKSCAN8 | 9081 | 26575 | 18988 | 0.404 | 50.400 | HepG2 |
| MLX | ZSCAN9 | 7285 | 16042 | 20246 | 0.404 | 71.621 | HepG2 |
| NFYA | TFDP2 | 8044 | 17604 | 22516 | 0.404 | 56.310 | HepG2 |
| IKZF5 | ZNF205 | 10067 | 31009 | 20027 | 0.404 | 50.325 | HepG2 |
| DRAP1 | MLX | 8281 | 26223 | 16042 | 0.404 | 58.241 | HepG2 |
| FOXP1 | MLX | 8626 | 28473 | 16042 | 0.404 | 50.126 | HepG2 |
| TFE3 | ZKSCAN8 | 8608 | 23975 | 18988 | 0.403 | 53.712 | HepG2 |
| KMT2B | ZNF511 | 7906 | 21702 | 17697 | 0.403 | 59.908 | HepG2 |
| DLX6 | ZNF264 | 7717 | 19647 | 18666 | 0.403 | 62.584 | HepG2 |
| ZNF264 | ZNF503 | 7486 | 18666 | 18529 | 0.403 | 56.463 | HepG2 |
| HOXD1 | MEIS2 | 6853 | 13879 | 20907 | 0.402 | 57.617 | HepG2 |
| ATF2 | ELF1 | 9389 | 23795 | 22892 | 0.402 | 53.650 | HepG2 |
| SMAD3 | ZNF511 | 7709 | 20760 | 17697 | 0.402 | 57.999 | HepG2 |
| DRAP1 | ZBTB25 | 9324 | 26223 | 20508 | 0.402 | 53.456 | HepG2 |
| SMAD3 | ZKSCAN8 | 7982 | 20760 | 18988 | 0.402 | 58.574 | HepG2 |
| KLF16 | MLX | 8905 | 30628 | 16042 | 0.402 | 51.827 | HepG2 |
| NFIC | TCF7 | 7336 | 17240 | 19413 | 0.401 | 62.053 | HepG2 |
| DRAP1 | ZNF580 | 9877 | 26223 | 23174 | 0.401 | 55.314 | HepG2 |
| PPARG | ZNF511 | 8607 | 26097 | 17697 | 0.401 | 56.922 | HepG2 |
| DRAP1 | NFYC | 8311 | 26223 | 16430 | 0.400 | 53.978 | HepG2 |
| FOXK1 | SIX4 | 7672 | 26575 | 13829 | 0.400 | 65.602 | HepG2 |
| ATF3 | FOSL1 | 4655 | 15603 | 8676 | 0.400 | 77.954 | HepG2 |
| NR2F2 | TCF7 | 7502 | 18140 | 19413 | 0.400 | 73.219 | HepG2 |
| KMT2B | PHF20 | 7810 | 21702 | 17625 | 0.399 | 56.290 | HepG2 |
| FOXP1 | NR2F2 | 9075 | 28473 | 18140 | 0.399 | 53.629 | HepG2 |
| MLX | SP5 | 8030 | 16042 | 25217 | 0.399 | 55.770 | HepG2 |
| KLF6 | ZNF792 | 8310 | 18502 | 23423 | 0.399 | 53.133 | HepG2 |
| ATF2 | ATF3 | 7691 | 23795 | 15603 | 0.399 | 60.456 | HepG2 |

| | | | | | | | |
|---|---|---|---|---|---|---|---|
| KLF6 | ZNF580 | 8261 | 18502 | 23174 | 0.399 | 52.425 | HepG2 |
| PPARG | ZSCAN9 | 9169 | 26097 | 20246 | 0.399 | 53.010 | HepG2 |
| GMEB1 | NR2C2 | 10331 | 21627 | 31046 | 0.399 | 52.297 | HepG2 |
| MXD4 | ZNF792 | 9080 | 22169 | 23423 | 0.398 | 59.076 | HepG2 |
| TFE3 | ZNF580 | 9391 | 23975 | 23174 | 0.398 | 55.512 | HepG2 |
| MXD1 | ZNF776 | 6191 | 18125 | 13339 | 0.398 | 71.679 | HepG2 |
| KLF6 | MXD4 | 8063 | 18502 | 22169 | 0.398 | 53.980 | HepG2 |
| ELF1 | KMT2B | 8873 | 22892 | 21702 | 0.398 | 53.586 | HepG2 |
| FOXJ3 | ZNF609 | 9145 | 21114 | 25025 | 0.398 | 57.206 | HepG2 |
| E2F4 | SPEN | 9702 | 21936 | 27137 | 0.398 | 57.619 | HepG2 |
| ATF2 | HMG20B | 7173 | 23795 | 13676 | 0.398 | 54.066 | HepG2 |
| MXD3 | NFKB2 | 7836 | 17726 | 21911 | 0.398 | 66.199 | HepG2 |
| E2F1 | TFDP2 | 6488 | 11827 | 22516 | 0.398 | 61.172 | HepG2 |
| GMEB1 | NFYA | 7756 | 21627 | 17604 | 0.397 | 54.041 | HepG2 |
| SP1 | TFDP1 | 7719 | 16039 | 23515 | 0.397 | 58.453 | HepG2 |
| NFYA | SP2 | 5602 | 17604 | 11297 | 0.397 | 56.791 | HepG2 |
| SMAD3 | ZNF124 | 7243 | 20760 | 16017 | 0.397 | 63.454 | HepG2 |
| HMG20B | PPARG | 7501 | 13676 | 26097 | 0.397 | 66.929 | HepG2 |
| SIX4 | THRA | 7831 | 13829 | 28190 | 0.397 | 63.620 | HepG2 |
| ATF2 | SP5 | 9709 | 23795 | 25217 | 0.396 | 50.107 | HepG2 |
| ZNF280B | ZNF891 | 6674 | 15695 | 18098 | 0.396 | 64.366 | HepG2 |
| GMEB1 | SP1 | 7353 | 21627 | 16039 | 0.395 | 71.598 | HepG2 |
| MXD4 | ZBTB25 | 8415 | 22169 | 20508 | 0.395 | 57.635 | HepG2 |
| HMG20A | SIX1 | 7793 | 26787 | 14572 | 0.394 | 57.897 | HepG2 |
| THRA | ZKSCAN8 | 9123 | 28190 | 18988 | 0.394 | 53.177 | HepG2 |
| KMT2A | TFDP2 | 8969 | 23007 | 22516 | 0.394 | 51.770 | HepG2 |
| ATF2 | ZNF503 | 8268 | 23795 | 18529 | 0.394 | 51.293 | HepG2 |
| ISL2 | MLX | 8517 | 29175 | 16042 | 0.394 | 51.009 | HepG2 |
| ZKSCAN8 | ZNF205 | 7674 | 18988 | 20027 | 0.394 | 63.499 | HepG2 |
| MLX | SMAD3 | 7177 | 16042 | 20760 | 0.393 | 69.900 | HepG2 |
| ISL2 | SIX1 | 8107 | 29175 | 14572 | 0.393 | 57.196 | HepG2 |
| HMG20B | THRA | 7693 | 13676 | 28190 | 0.392 | 57.111 | HepG2 |
| HMG20B | MIXL1 | 7678 | 13676 | 28115 | 0.392 | 62.553 | HepG2 |
| HMG20B | TFE3 | 7089 | 13676 | 23975 | 0.391 | 68.908 | HepG2 |
| FOXP1 | NFIC | 8669 | 28473 | 17240 | 0.391 | 61.469 | HepG2 |
| HMG20B | LCOR | 8765 | 13676 | 36711 | 0.391 | 51.329 | HepG2 |
| ZFP1 | ZNF614 | 7313 | 13958 | 25059 | 0.391 | 61.833 | HepG2 |
| HMG20B | ZNF614 | 7235 | 13676 | 25059 | 0.391 | 60.796 | HepG2 |
| HMG20B | ZNF792 | 6991 | 13676 | 23423 | 0.391 | 50.846 | HepG2 |
| MLX | MYBL2 | 7723 | 16042 | 24417 | 0.390 | 50.215 | HepG2 |
| ISL2 | ZSCAN9 | 9474 | 29175 | 20246 | 0.390 | 53.232 | HepG2 |
| MXI1 | NFYA | 8242 | 25399 | 17604 | 0.390 | 50.296 | HepG2 |
| HMG20B | IRF2 | 8035 | 13676 | 31142 | 0.389 | 52.522 | HepG2 |
| ZKSCAN8 | ZNF580 | 8164 | 18988 | 23174 | 0.389 | 60.816 | HepG2 |
| GATA4 | ISL2 | 7834 | 13900 | 29175 | 0.389 | 53.714 | HepG2 |
| ISL2 | SIX4 | 7813 | 29175 | 13829 | 0.389 | 50.685 | HepG2 |
| ZNF124 | ZNF792 | 7531 | 16017 | 23423 | 0.389 | 56.701 | HepG2 |
| MXD4 | ZSCAN9 | 8213 | 22169 | 20246 | 0.388 | 50.069 | HepG2 |
| ATF2 | ZBED4 | 7693 | 23795 | 16556 | 0.388 | 51.736 | HepG2 |
| ZFP1 | ZKSCAN8 | 6309 | 13958 | 18988 | 0.388 | 64.147 | HepG2 |
| ARNTL | ZNF556 | 7820 | 20368 | 20009 | 0.387 | 54.332 | HepG2 |
| HOMEZ | ZSCAN9 | 9124 | 27410 | 20246 | 0.387 | 51.550 | HepG2 |

| | | | | | | | |
|---|---|---|---|---|---|---|---|
| THRA | ZNF511 | 8649 | 28190 | 17697 | 0.387 | 53.213 | HepG2 |
| KLF16 | ZNF335 | 8360 | 30628 | 15231 | 0.387 | 51.482 | HepG2 |
| HOXA3 | ZHX2 | 8026 | 40277 | 10676 | 0.387 | 53.797 | HepG2 |
| MLX | ZBTB25 | 7017 | 16042 | 20508 | 0.387 | 54.952 | HepG2 |
| ZNF274 | ZNF547 | 7148 | 17918 | 19097 | 0.386 | 55.822 | HepG2 |
| RBPJ | ZNF124 | 8391 | 29519 | 16017 | 0.386 | 51.075 | HepG2 |
| FOXJ3 | NFIC | 7355 | 21114 | 17240 | 0.386 | 53.270 | HepG2 |
| ZFP1 | ZNF511 | 6058 | 13958 | 17697 | 0.385 | 70.627 | HepG2 |
| IRF2 | ZFP1 | 8034 | 31142 | 13958 | 0.385 | 56.628 | HepG2 |
| SIX4 | SMAD3 | 6528 | 13829 | 20760 | 0.385 | 57.746 | HepG2 |
| HOXA5 | MEIS2 | 8046 | 20899 | 20907 | 0.385 | 54.908 | HepG2 |
| PPARG | ZBTB25 | 8904 | 26097 | 20508 | 0.385 | 51.090 | HepG2 |
| ZFP1 | ZSCAN9 | 6470 | 13958 | 20246 | 0.385 | 66.338 | HepG2 |
| KLF6 | SMAD3 | 7543 | 18502 | 20760 | 0.385 | 53.580 | HepG2 |
| MLX | ZNF205 | 6896 | 16042 | 20027 | 0.385 | 62.751 | HepG2 |
| ZFP1 | ZNF792 | 6951 | 13958 | 23423 | 0.384 | 52.672 | HepG2 |
| KMT2B | ZNF639 | 6559 | 21702 | 13418 | 0.384 | 63.480 | HepG2 |
| FOXP1 | TBX3 | 8011 | 28473 | 15274 | 0.384 | 54.110 | HepG2 |
| ARNTL | MLX | 6942 | 20368 | 16042 | 0.384 | 60.502 | HepG2 |
| HHEX | TBX3 | 4287 | 8174 | 15274 | 0.384 | 76.463 | HepG2 |
| HOXA5 | HOXD1 | 6534 | 20899 | 13879 | 0.384 | 58.060 | HepG2 |
| SP1 | SPEN | 7996 | 16039 | 27137 | 0.383 | 56.962 | HepG2 |
| ZNF556 | ZNF691 | 7508 | 20009 | 19190 | 0.383 | 55.494 | HepG2 |
| MXD4 | ZNF511 | 7588 | 22169 | 17697 | 0.383 | 50.292 | HepG2 |
| HOXD1 | ZNF503 | 6135 | 13879 | 18529 | 0.383 | 61.640 | HepG2 |
| HBP1 | TGIF2 | 6837 | 15317 | 20855 | 0.383 | 51.777 | HepG2 |
| KLF6 | ZNF511 | 6921 | 18502 | 17697 | 0.382 | 79.297 | HepG2 |
| HOXD1 | ISL2 | 7695 | 13879 | 29175 | 0.382 | 52.413 | HepG2 |
| MLX | SIX4 | 5691 | 16042 | 13829 | 0.382 | 70.910 | HepG2 |
| ZNF124 | ZNF580 | 7360 | 16017 | 23174 | 0.382 | 53.056 | HepG2 |
| GMEB1 | GMEB2 | 4104 | 21627 | 5338 | 0.382 | 88.806 | HepG2 |
| GMEB1 | TFDP1 | 8608 | 21627 | 23515 | 0.382 | 50.006 | HepG2 |
| FOXK1 | HMG20B | 7276 | 26575 | 13676 | 0.382 | 59.864 | HepG2 |
| KLF6 | ZNF614 | 8217 | 18502 | 25059 | 0.382 | 55.970 | HepG2 |
| KMT2B | ZNF205 | 7955 | 21702 | 20027 | 0.382 | 50.773 | HepG2 |
| HMG20B | ZKSCAN8 | 6147 | 13676 | 18988 | 0.381 | 69.303 | HepG2 |
| FOXO1 | ZNF511 | 8139 | 25752 | 17697 | 0.381 | 51.154 | HepG2 |
| MLX | ZFP1 | 5704 | 16042 | 13958 | 0.381 | 65.036 | HepG2 |
| ATF2 | KLF6 | 7998 | 23795 | 18502 | 0.381 | 58.333 | HepG2 |
| KMT2A | ZNF580 | 8800 | 23007 | 23174 | 0.381 | 50.381 | HepG2 |
| NR2F2 | THRA | 8617 | 18140 | 28190 | 0.381 | 50.362 | HepG2 |
| FOXJ3 | MYBL2 | 8652 | 21114 | 24417 | 0.381 | 53.258 | HepG2 |
| ZBTB26 | ZNF580 | 7508 | 16775 | 23174 | 0.381 | 50.777 | HepG2 |
| FOXP1 | SIX1 | 7750 | 28473 | 14572 | 0.380 | 51.059 | HepG2 |
| KLF6 | TFE3 | 8011 | 18502 | 23975 | 0.380 | 51.754 | HepG2 |
| SMAD3 | ZNF205 | 7739 | 20760 | 20027 | 0.380 | 55.034 | HepG2 |
| MYBL2 | ZKSCAN8 | 8169 | 24417 | 18988 | 0.379 | 50.787 | HepG2 |
| NFYB | SP2 | 5007 | 15448 | 11297 | 0.379 | 74.848 | HepG2 |
| ATF2 | ZFP1 | 6906 | 23795 | 13958 | 0.379 | 52.593 | HepG2 |
| MIXL1 | SIX1 | 7670 | 28115 | 14572 | 0.379 | 50.279 | HepG2 |
| MEIS2 | ZNF503 | 7457 | 20907 | 18529 | 0.379 | 55.449 | HepG2 |
| HMG20B | SIX4 | 5208 | 13676 | 13829 | 0.379 | 75.582 | HepG2 |

| | | | | | | | |
|---|---|---|---|---|---|---|---|
| PPARG | ZNF503 | 8327 | 26097 | 18529 | 0.379 | 50.617 | HepG2 |
| MLX | PITX1 | 8206 | 16042 | 29314 | 0.378 | 63.082 | HepG2 |
| HMG20B | ZSCAN9 | 6293 | 13676 | 20246 | 0.378 | 59.035 | HepG2 |
| PPARG | SIX1 | 7373 | 26097 | 14572 | 0.378 | 58.279 | HepG2 |
| HMG20B | SMAD3 | 6370 | 13676 | 20760 | 0.378 | 58.976 | HepG2 |
| MXD1 | MXD3 | 6773 | 18125 | 17726 | 0.378 | 63.235 | HepG2 |
| ERF | ZBTB26 | 8802 | 32451 | 16775 | 0.377 | 58.575 | HepG2 |
| ZNF124 | ZSCAN9 | 6791 | 16017 | 20246 | 0.377 | 55.171 | HepG2 |
| ZNF639 | ZNF792 | 6685 | 13418 | 23423 | 0.377 | 50.820 | HepG2 |
| FOXO1 | SIX1 | 7296 | 25752 | 14572 | 0.377 | 55.123 | HepG2 |
| HNF4G | NR2F2 | 8005 | 24914 | 18140 | 0.377 | 52.178 | HepG2 |
| MYBL2 | SIX4 | 6907 | 24417 | 13829 | 0.376 | 52.955 | HepG2 |
| KLF6 | ZNF639 | 5921 | 18502 | 13418 | 0.376 | 76.981 | HepG2 |
| NFIC | NR2F2 | 6643 | 17240 | 18140 | 0.376 | 60.235 | HepG2 |
| HMG20B | ZNF511 | 5839 | 13676 | 17697 | 0.375 | 64.807 | HepG2 |
| ZBTB25 | ZFP1 | 6349 | 20508 | 13958 | 0.375 | 57.284 | HepG2 |
| TBX3 | ZNF609 | 7334 | 15274 | 25025 | 0.375 | 54.707 | HepG2 |
| ERF | HMG20B | 7900 | 32451 | 13676 | 0.375 | 50.968 | HepG2 |
| ELF1 | SP1 | 7177 | 22892 | 16039 | 0.375 | 55.916 | HepG2 |
| SIX4 | TFE3 | 6806 | 13829 | 23975 | 0.374 | 61.831 | HepG2 |
| HMG20B | IKZF5 | 7692 | 13676 | 31009 | 0.374 | 50.792 | HepG2 |
| ZNF547 | ZNF784 | 6256 | 19097 | 14697 | 0.373 | 64.976 | HepG2 |
| FOXJ3 | TCF7 | 7559 | 21114 | 19413 | 0.373 | 50.434 | HepG2 |
| HMG20B | ZNF503 | 5942 | 13676 | 18529 | 0.373 | 59.895 | HepG2 |
| FOXK1 | ZNF124 | 7698 | 26575 | 16017 | 0.373 | 54.422 | HepG2 |
| ZIK1 | ZNF451 | 4430 | 10981 | 12838 | 0.373 | 79.976 | HepG2 |
| KLF6 | ZNF205 | 7182 | 18502 | 20027 | 0.373 | 51.238 | HepG2 |
| HHEX | ZNF609 | 5331 | 8174 | 25025 | 0.373 | 75.948 | HepG2 |
| KLF6 | MXD3 | 6742 | 18502 | 17726 | 0.372 | 58.806 | HepG2 |
| SMAD3 | ZBED4 | 6896 | 20760 | 16556 | 0.372 | 56.178 | HepG2 |
| IRF2 | ZNF335 | 8099 | 31142 | 15231 | 0.372 | 56.283 | HepG2 |
| GMEB1 | ZHX2 | 5646 | 21627 | 10676 | 0.372 | 62.009 | HepG2 |
| ZBTB25 | ZNF124 | 6732 | 20508 | 16017 | 0.371 | 66.326 | HepG2 |
| NFYC | SP2 | 5060 | 16430 | 11297 | 0.371 | 77.358 | HepG2 |
| MLX | ZNF503 | 6395 | 16042 | 18529 | 0.371 | 64.863 | HepG2 |
| ZNF124 | ZNF511 | 6244 | 16017 | 17697 | 0.371 | 50.880 | HepG2 |
| ATF2 | ETV4 | 8440 | 23795 | 21788 | 0.371 | 50.243 | HepG2 |
| SMAD3 | ZNF503 | 7269 | 20760 | 18529 | 0.371 | 59.266 | HepG2 |
| RBPJ | ZNF639 | 7375 | 29519 | 13418 | 0.371 | 54.949 | HepG2 |
| DLX6 | HOXD1 | 6117 | 19647 | 13879 | 0.370 | 57.273 | HepG2 |
| CREM | SIX1 | 7719 | 29846 | 14572 | 0.370 | 50.321 | HepG2 |
| SMAD3 | ZFP1 | 6297 | 20760 | 13958 | 0.370 | 50.146 | HepG2 |
| HMG20B | ZBTB25 | 6192 | 13676 | 20508 | 0.370 | 61.650 | HepG2 |
| HOXA5 | NKX3-1 | 6125 | 20899 | 13144 | 0.370 | 61.945 | HepG2 |
| FOXP1 | HHEX | 5632 | 28473 | 8174 | 0.369 | 56.322 | HepG2 |
| NFYC | SP1 | 5992 | 16430 | 16039 | 0.369 | 60.867 | HepG2 |
| ZBED4 | ZNF580 | 7222 | 16556 | 23174 | 0.369 | 54.317 | HepG2 |
| ATF2 | ZNF335 | 7019 | 23795 | 15231 | 0.369 | 55.918 | HepG2 |
| FOXJ3 | NR2F2 | 7214 | 21114 | 18140 | 0.369 | 52.154 | HepG2 |
| TFE3 | ZFP1 | 6740 | 23975 | 13958 | 0.368 | 59.338 | HepG2 |
| HHEX | TCF7 | 4640 | 8174 | 19413 | 0.368 | 69.093 | HepG2 |
| MYBL2 | NFYA | 7634 | 24417 | 17604 | 0.368 | 50.015 | HepG2 |

| | | | | | | | |
|---|---|---|---|---|---|---|---|
| ETV4 | GATA4 | 6400 | 21788 | 13900 | 0.368 | 51.132 | HepG2 |
| NKX3-1 | ZNF232 | 5412 | 13144 | 16478 | 0.368 | 63.769 | HepG2 |
| HMG20B | ZFP1 | 5077 | 13676 | 13958 | 0.367 | 65.248 | HepG2 |
| MIXL1 | ZFP1 | 7279 | 28115 | 13958 | 0.367 | 53.347 | HepG2 |
| FOXO1 | TCF7 | 8215 | 25752 | 19413 | 0.367 | 50.987 | HepG2 |
| ZBED4 | ZNF792 | 7235 | 16556 | 23423 | 0.367 | 52.886 | HepG2 |
| KMT2B | ZNF335 | 6678 | 21702 | 15231 | 0.367 | 50.412 | HepG2 |
| MEIS2 | ZNF264 | 7254 | 20907 | 18666 | 0.367 | 52.842 | HepG2 |
| NKX3-1 | ZNF503 | 5727 | 13144 | 18529 | 0.367 | 57.391 | HepG2 |
| GATA4 | ZNF503 | 5882 | 13900 | 18529 | 0.367 | 53.795 | HepG2 |
| MXD4 | ZNF639 | 6321 | 22169 | 13418 | 0.366 | 50.307 | HepG2 |
| AHR | MXD3 | 5368 | 12116 | 17726 | 0.366 | 59.745 | HepG2 |
| FOXJ3 | GATA2 | 7928 | 21114 | 22221 | 0.366 | 50.441 | HepG2 |
| MLX | ZNF580 | 7048 | 16042 | 23174 | 0.366 | 50.245 | HepG2 |
| ZNF511 | ZNF639 | 5624 | 17697 | 13418 | 0.365 | 61.622 | HepG2 |
| KMT2B | MXD3 | 7153 | 21702 | 17726 | 0.365 | 60.522 | HepG2 |
| GATA4 | ZNF609 | 6801 | 13900 | 25025 | 0.365 | 54.923 | HepG2 |
| ZNF580 | ZNF639 | 6421 | 23174 | 13418 | 0.364 | 55.069 | HepG2 |
| HOXD1 | NKX3-1 | 4911 | 13879 | 13144 | 0.364 | 67.414 | HepG2 |
| MEF2D | MLX | 6969 | 22926 | 16042 | 0.363 | 68.213 | HepG2 |
| SIX4 | ZNF503 | 5815 | 13829 | 18529 | 0.363 | 55.770 | HepG2 |
| SIX1 | ZNF503 | 5968 | 14572 | 18529 | 0.363 | 64.697 | HepG2 |
| ZNF124 | ZNF205 | 6500 | 16017 | 20027 | 0.363 | 54.499 | HepG2 |
| ZNF205 | ZNF639 | 5940 | 20027 | 13418 | 0.362 | 60.925 | HepG2 |
| FOXK1 | ZNF335 | 7271 | 26575 | 15231 | 0.361 | 50.726 | HepG2 |
| ZBED4 | ZNF124 | 5884 | 16556 | 16017 | 0.361 | 62.284 | HepG2 |
| HIVEP1 | MXD1 | 6466 | 17669 | 18125 | 0.361 | 59.276 | HepG2 |
| TFDP2 | ZNF124 | 6856 | 22516 | 16017 | 0.361 | 61.439 | HepG2 |
| ETV4 | MLX | 6747 | 21788 | 16042 | 0.361 | 51.953 | HepG2 |
| MXD1 | NFKB2 | 7188 | 18125 | 21911 | 0.361 | 59.410 | HepG2 |
| HHEX | NR2F2 | 4384 | 8174 | 18140 | 0.360 | 62.319 | HepG2 |
| SIX4 | ZNF614 | 6702 | 13829 | 25059 | 0.360 | 54.985 | HepG2 |
| HNF1A | SIX4 | 6706 | 25102 | 13829 | 0.360 | 53.828 | HepG2 |
| ZBED4 | ZBTB25 | 6630 | 16556 | 20508 | 0.360 | 50.722 | HepG2 |
| RBPJ | SIX4 | 7250 | 29519 | 13829 | 0.359 | 53.609 | HepG2 |
| KLF11 | KMT2B | 5835 | 12213 | 21702 | 0.358 | 57.595 | HepG2 |
| HMG20B | SIX1 | 5058 | 13676 | 14572 | 0.358 | 57.560 | HepG2 |
| ZBED4 | ZSCAN9 | 6559 | 16556 | 20246 | 0.358 | 55.112 | HepG2 |
| ZFP1 | ZNF205 | 5983 | 13958 | 20027 | 0.358 | 60.310 | HepG2 |
| MYBL2 | ZBED4 | 7194 | 24417 | 16556 | 0.358 | 55.064 | HepG2 |
| ZNF232 | ZNF766 | 5241 | 16478 | 13024 | 0.358 | 64.243 | HepG2 |
| ELF1 | GABPA | 4897 | 22892 | 8201 | 0.357 | 86.064 | HepG2 |
| ARNTL | ESRRA | 7087 | 20368 | 19314 | 0.357 | 66.153 | HepG2 |
| HMG20B | HOMEZ | 6918 | 13676 | 27410 | 0.357 | 54.648 | HepG2 |
| HES4 | ZNF446 | 4055 | 13098 | 9839 | 0.357 | 68.445 | HepG2 |
| TFE3 | ZNF503 | 7526 | 23975 | 18529 | 0.357 | 54.736 | HepG2 |
| ZBTB26 | ZSCAN9 | 6577 | 16775 | 20246 | 0.357 | 52.423 | HepG2 |
| HMG20B | MYBL2 | 6521 | 13676 | 24417 | 0.357 | 51.169 | HepG2 |
| TBX3 | TCF7 | 6143 | 15274 | 19413 | 0.357 | 56.953 | HepG2 |
| FOXO1 | ZFP1 | 6760 | 25752 | 13958 | 0.357 | 50.736 | HepG2 |
| KLF16 | ZNF639 | 7227 | 30628 | 13418 | 0.356 | 52.451 | HepG2 |
| ATF3 | FOXO1 | 7144 | 15603 | 25752 | 0.356 | 51.515 | HepG2 |

| | | | | | | | |
|---|---|---|---|---|---|---|---|
| KLF11 | ZNF511 | 5239 | 12213 | 17697 | 0.356 | 57.169 | HepG2 |
| MLX | SIX1 | 5448 | 16042 | 14572 | 0.356 | 61.178 | HepG2 |
| MYBL2 | ZFP1 | 6578 | 24417 | 13958 | 0.356 | 60.931 | HepG2 |
| ZNF639 | ZSCAN9 | 5872 | 13418 | 20246 | 0.356 | 61.375 | HepG2 |
| HHEX | HMG20A | 5271 | 8174 | 26787 | 0.356 | 52.311 | HepG2 |
| ZNF343 | ZNF543 | 5526 | 13429 | 17952 | 0.356 | 51.313 | HepG2 |
| SP1 | SP2 | 4790 | 16039 | 11297 | 0.356 | 77.423 | HepG2 |
| ZNF483 | ZNF556 | 6547 | 16926 | 20009 | 0.356 | 52.446 | HepG2 |
| HINFP | ZNF556 | 6007 | 14261 | 20009 | 0.356 | 53.211 | HepG2 |
| TFE3 | ZNF124 | 6966 | 23975 | 16017 | 0.355 | 50.187 | HepG2 |
| TFDP1 | ZNF639 | 6314 | 23515 | 13418 | 0.355 | 56.484 | HepG2 |
| FOXK1 | FOXK2 | 6014 | 26575 | 10775 | 0.355 | 50.715 | HepG2 |
| MBD1 | TGIF2 | 6316 | 15149 | 20855 | 0.355 | 61.691 | HepG2 |
| FOXK1 | ZBED4 | 7442 | 26575 | 16556 | 0.355 | 53.155 | HepG2 |
| HINFP | ZNF691 | 5866 | 14261 | 19190 | 0.355 | 54.118 | HepG2 |
| KLF6 | ZNF124 | 6103 | 18502 | 16017 | 0.355 | 62.298 | HepG2 |
| IRF5 | TIGD6 | 4390 | 11111 | 13801 | 0.355 | 60.832 | HepG2 |
| KMT2B | ZKSCAN8 | 7191 | 21702 | 18988 | 0.354 | 54.302 | HepG2 |
| SIX4 | ZKSCAN8 | 5734 | 13829 | 18988 | 0.354 | 53.710 | HepG2 |
| ZNF548 | ZNF556 | 5974 | 14256 | 20009 | 0.354 | 59.924 | HepG2 |
| HHEX | HMG20B | 3734 | 8174 | 13676 | 0.353 | 81.188 | HepG2 |
| HOXD1 | ZNF232 | 5339 | 13879 | 16478 | 0.353 | 65.678 | HepG2 |
| ATF2 | ETS1 | 5249 | 23795 | 9307 | 0.353 | 56.237 | HepG2 |
| GMEB1 | ZNF143 | 6153 | 21627 | 14071 | 0.353 | 52.460 | HepG2 |
| SIX4 | ZSCAN9 | 5889 | 13829 | 20246 | 0.352 | 61.770 | HepG2 |
| KMT2B | ZHX2 | 5357 | 21702 | 10676 | 0.352 | 63.627 | HepG2 |
| ZFP37 | ZNF232 | 4443 | 9678 | 16478 | 0.352 | 68.041 | HepG2 |
| E2F4 | KLF12 | 6392 | 21936 | 15074 | 0.352 | 58.051 | HepG2 |
| ZNF232 | ZNF264 | 6161 | 16478 | 18666 | 0.351 | 50.632 | HepG2 |
| TFDP2 | ZNF639 | 6104 | 22516 | 13418 | 0.351 | 58.831 | HepG2 |
| ATF2 | GATA4 | 6382 | 23795 | 13900 | 0.351 | 50.430 | HepG2 |
| SIX1 | TCF7 | 5900 | 14572 | 19413 | 0.351 | 61.204 | HepG2 |
| KLF11 | ZSCAN9 | 5514 | 12213 | 20246 | 0.351 | 62.043 | HepG2 |
| ZNF335 | ZSCAN9 | 6155 | 15231 | 20246 | 0.351 | 53.916 | HepG2 |
| SPEN | TIGD6 | 6783 | 27137 | 13801 | 0.350 | 56.820 | HepG2 |
| ATF2 | ZHX2 | 5583 | 23795 | 10676 | 0.350 | 54.193 | HepG2 |
| SP1 | YY1 | 6759 | 16039 | 23215 | 0.350 | 52.201 | HepG2 |
| SAFB2 | TIGD6 | 4492 | 11917 | 13801 | 0.350 | 61.752 | HepG2 |
| SP1 | ZHX2 | 4582 | 16039 | 10676 | 0.350 | 68.439 | HepG2 |
| SIX4 | ZNF511 | 5475 | 13829 | 17697 | 0.350 | 68.410 | HepG2 |
| HES4 | ZNF709 | 5290 | 13098 | 17451 | 0.350 | 59.391 | HepG2 |
| MXD3 | ZNF639 | 5393 | 17726 | 13418 | 0.350 | 60.462 | HepG2 |
| NFYB | SP1 | 5501 | 15448 | 16039 | 0.349 | 53.609 | HepG2 |
| FOXJ3 | MLX | 6425 | 21114 | 16042 | 0.349 | 51.211 | HepG2 |
| ZNF230 | ZNF556 | 6140 | 15509 | 20009 | 0.349 | 65.118 | HepG2 |
| FOXK1 | ZHX2 | 5866 | 26575 | 10676 | 0.348 | 58.153 | HepG2 |
| ZBTB26 | ZNF205 | 6376 | 16775 | 20027 | 0.348 | 52.635 | HepG2 |
| DLX6 | ZKSCAN8 | 6705 | 19647 | 18988 | 0.347 | 52.061 | HepG2 |
| E2F1 | GMEB1 | 5544 | 11827 | 21627 | 0.347 | 54.805 | HepG2 |
| KLF11 | ZBTB25 | 5485 | 12213 | 20508 | 0.347 | 59.797 | HepG2 |
| HMG20B | NR2F2 | 5458 | 13676 | 18140 | 0.347 | 67.137 | HepG2 |
| NFE2 | NFE2L2 | 2094 | 3508 | 10434 | 0.346 | 114.129 | HepG2 |

| | | | | | | | |
|---|---|---|---|---|---|---|---|
| GMEB1 | ZNF639 | 5894 | 21627 | 13418 | 0.346 | 55.852 | HepG2 |
| HOXA5 | ZNF766 | 5706 | 20899 | 13024 | 0.346 | 60.663 | HepG2 |
| FOXJ3 | HMG20B | 5871 | 21114 | 13676 | 0.345 | 62.429 | HepG2 |
| KLF6 | ZBED4 | 6037 | 18502 | 16556 | 0.345 | 60.837 | HepG2 |
| ZIK1 | ZNF616 | 3422 | 10981 | 8986 | 0.344 | 79.525 | HepG2 |
| YY1 | ZHX2 | 5418 | 23215 | 10676 | 0.344 | 64.458 | HepG2 |
| ELK1 | HOXD1 | 3997 | 9720 | 13879 | 0.344 | 69.976 | HepG2 |
| KLF11 | ZNF792 | 5819 | 12213 | 23423 | 0.344 | 51.478 | HepG2 |
| NFYC | TFDP2 | 6611 | 16430 | 22516 | 0.344 | 51.016 | HepG2 |
| ZNF333 | ZNF776 | 4160 | 10988 | 13339 | 0.344 | 73.891 | HepG2 |
| DLX6 | NKX3-1 | 5521 | 19647 | 13144 | 0.344 | 52.524 | HepG2 |
| HMG20B | RREB1 | 6032 | 13676 | 22566 | 0.343 | 51.868 | HepG2 |
| DRAP1 | ZFP1 | 6568 | 26223 | 13958 | 0.343 | 52.932 | HepG2 |
| GABPA | PHF20 | 4127 | 8201 | 17625 | 0.343 | 59.940 | HepG2 |
| NKX3-1 | SATB2 | 3514 | 13144 | 8020 | 0.342 | 74.703 | HepG2 |
| MXI1 | ZHX2 | 5633 | 25399 | 10676 | 0.342 | 51.694 | HepG2 |
| ZNF483 | ZNF543 | 5957 | 16926 | 17952 | 0.342 | 50.040 | HepG2 |
| KLF6 | ZBTB26 | 6020 | 18502 | 16775 | 0.342 | 52.220 | HepG2 |
| HOXA5 | KLF12 | 6065 | 20899 | 15074 | 0.342 | 51.149 | HepG2 |
| MXD1 | ZNF333 | 4821 | 18125 | 10988 | 0.342 | 57.167 | HepG2 |
| DLX6 | SIX1 | 5780 | 19647 | 14572 | 0.342 | 62.352 | HepG2 |
| KLF11 | KLF6 | 5133 | 12213 | 18502 | 0.341 | 56.759 | HepG2 |
| ZNF335 | ZNF511 | 5604 | 15231 | 17697 | 0.341 | 63.850 | HepG2 |
| PHF20 | ZHX2 | 4681 | 17625 | 10676 | 0.341 | 68.149 | HepG2 |
| MLX | ZNF124 | 5470 | 16042 | 16017 | 0.341 | 67.386 | HepG2 |
| ZBTB26 | ZNF511 | 5879 | 16775 | 17697 | 0.341 | 51.759 | HepG2 |
| E2F1 | KLF12 | 4555 | 11827 | 15074 | 0.341 | 58.617 | HepG2 |
| AHR | ZBTB26 | 4862 | 12116 | 16775 | 0.341 | 55.177 | HepG2 |
| ZNF124 | ZNF335 | 5326 | 16017 | 15231 | 0.341 | 54.524 | HepG2 |
| ZNF230 | ZNF543 | 5684 | 15509 | 17952 | 0.341 | 52.965 | HepG2 |
| ZNF451 | ZNF616 | 3656 | 12838 | 8986 | 0.340 | 76.847 | HepG2 |
| ZNF274 | ZNF33B | 4714 | 17918 | 10715 | 0.340 | 57.754 | HepG2 |
| ETS1 | FOXK1 | 5350 | 9307 | 26575 | 0.340 | 50.155 | HepG2 |
| DR1 | GMEB1 | 5132 | 10534 | 21627 | 0.340 | 61.969 | HepG2 |
| ARNTL | HINFP | 5794 | 20368 | 14261 | 0.340 | 55.540 | HepG2 |
| KLF11 | MXD4 | 5588 | 12213 | 22169 | 0.340 | 56.472 | HepG2 |
| KMT2B | MLX | 6327 | 21702 | 16042 | 0.339 | 51.783 | HepG2 |
| NKX3-1 | ZNF766 | 4435 | 13144 | 13024 | 0.339 | 61.792 | HepG2 |
| DR1 | TBP | 4697 | 10534 | 18258 | 0.339 | 68.891 | HepG2 |
| CEBPZ | IRF3 | 200 | 491 | 712 | 0.338 | 124.587 | HepG2 |
| ELK1 | ZFP37 | 3280 | 9720 | 9678 | 0.338 | 90.422 | HepG2 |
| ZFP37 | ZNF557 | 3077 | 9678 | 8561 | 0.338 | 87.314 | HepG2 |
| ZNF446 | ZNF709 | 4426 | 9839 | 17451 | 0.338 | 73.649 | HepG2 |
| ZNF142 | ZNF766 | 4683 | 14794 | 13024 | 0.337 | 61.582 | HepG2 |
| KLF11 | ZNF580 | 5675 | 12213 | 23174 | 0.337 | 51.383 | HepG2 |
| ELK1 | ZNF232 | 4269 | 9720 | 16478 | 0.337 | 57.342 | HepG2 |
| ZNF230 | ZNF548 | 5015 | 15509 | 14256 | 0.337 | 61.541 | HepG2 |
| KLF6 | ZNF335 | 5659 | 18502 | 15231 | 0.337 | 52.259 | HepG2 |
| MEIS2 | NKX3-1 | 5586 | 20907 | 13144 | 0.337 | 54.519 | HepG2 |
| ZNF230 | ZNF343 | 4861 | 15509 | 13429 | 0.337 | 60.179 | HepG2 |
| HBP1 | KMT2B | 6141 | 15317 | 21702 | 0.337 | 52.341 | HepG2 |
| GMEB1 | SP2 | 5262 | 21627 | 11297 | 0.337 | 51.895 | HepG2 |

| | | | | | | | |
|---|---|---|---|---|---|---|---|
| TFDP2 | ZHX2 | 5219 | 22516 | 10676 | 0.337 | 51.826 | HepG2 |
| E2F4 | SP2 | 5295 | 21936 | 11297 | 0.336 | 52.203 | HepG2 |
| FOXJ3 | TBX3 | 6039 | 21114 | 15274 | 0.336 | 50.474 | HepG2 |
| E2F4 | ZNF143 | 5903 | 21936 | 14071 | 0.336 | 50.634 | HepG2 |
| MEF2A | SMAD3 | 4501 | 8647 | 20760 | 0.336 | 62.370 | HepG2 |
| ATF2 | MEF2A | 4817 | 23795 | 8647 | 0.336 | 54.831 | HepG2 |
| NFAT5 | ZNF773 | 2628 | 9150 | 6695 | 0.336 | 90.991 | HepG2 |
| TBX3 | ZNF503 | 5648 | 15274 | 18529 | 0.336 | 50.820 | HepG2 |
| NFIC | SIX1 | 5321 | 17240 | 14572 | 0.336 | 51.216 | HepG2 |
| GATA4 | SIX4 | 4650 | 13900 | 13829 | 0.335 | 51.055 | HepG2 |
| MEIS2 | ZNF766 | 5531 | 20907 | 13024 | 0.335 | 52.997 | HepG2 |
| ZNF33B | ZNF547 | 4794 | 10715 | 19097 | 0.335 | 55.113 | HepG2 |
| ZFP1 | ZNF124 | 5010 | 13958 | 16017 | 0.335 | 57.047 | HepG2 |
| GATA4 | MLX | 5002 | 13900 | 16042 | 0.335 | 53.578 | HepG2 |
| MXD3 | ZNF451 | 5052 | 17726 | 12838 | 0.335 | 63.928 | HepG2 |
| IKZF4 | ZNF274 | 4433 | 9788 | 17918 | 0.335 | 64.526 | HepG2 |
| PHF20 | SP1 | 5627 | 17625 | 16039 | 0.335 | 61.171 | HepG2 |
| EEA1 | IRF5 | 3399 | 9299 | 11111 | 0.334 | 68.825 | HepG2 |
| NFAT5 | ZNF232 | 4106 | 9150 | 16478 | 0.334 | 58.193 | HepG2 |
| IKZF4 | ZNF547 | 4571 | 9788 | 19097 | 0.334 | 73.208 | HepG2 |
| ZNF232 | ZNF557 | 3970 | 16478 | 8561 | 0.334 | 71.709 | HepG2 |
| HHEX | ZNF503 | 4105 | 8174 | 18529 | 0.334 | 75.158 | HepG2 |
| GMEB2 | ZHX2 | 2517 | 5338 | 10676 | 0.333 | 73.268 | HepG2 |
| GMEB2 | SP1 | 3084 | 5338 | 16039 | 0.333 | 58.882 | HepG2 |
| IRF5 | SAFB2 | 3833 | 11111 | 11917 | 0.333 | 58.399 | HepG2 |
| SIX4 | ZNF124 | 4956 | 13829 | 16017 | 0.333 | 56.689 | HepG2 |
| GATA4 | HMG20B | 4586 | 13900 | 13676 | 0.333 | 53.875 | HepG2 |
| FOXO1 | HHEX | 4825 | 25752 | 8174 | 0.333 | 54.223 | HepG2 |
| NR2F2 | TBX3 | 5529 | 18140 | 15274 | 0.332 | 50.963 | HepG2 |
| FOXP4 | HHEX | 5664 | 35653 | 8174 | 0.332 | 50.538 | HepG2 |
| MLX | TBX3 | 5190 | 16042 | 15274 | 0.332 | 50.736 | HepG2 |
| HMG20B | TBX3 | 4790 | 13676 | 15274 | 0.331 | 54.695 | HepG2 |
| E2F1 | NFYA | 4774 | 11827 | 17604 | 0.331 | 66.214 | HepG2 |
| DR1 | E2F1 | 3689 | 10534 | 11827 | 0.331 | 61.673 | HepG2 |
| DR1 | DRAP1 | 5491 | 10534 | 26223 | 0.330 | 50.614 | HepG2 |
| KLF12 | ZNF766 | 4625 | 15074 | 13024 | 0.330 | 57.222 | HepG2 |
| ZNF230 | ZNF483 | 5345 | 15509 | 16926 | 0.330 | 52.581 | HepG2 |
| ZNF543 | ZNF548 | 5273 | 17952 | 14256 | 0.330 | 51.312 | HepG2 |
| HHEX | TCF7L2 | 5709 | 8174 | 36713 | 0.330 | 53.042 | HepG2 |
| DZIP1 | NRL | 4446 | 9278 | 19698 | 0.329 | 64.804 | HepG2 |
| CAMTA2 | ZNF776 | 3447 | 8249 | 13339 | 0.329 | 86.500 | HepG2 |
| HHEX | SIX1 | 3583 | 8174 | 14572 | 0.328 | 60.881 | HepG2 |
| KLF9 | ZNF511 | 4017 | 8472 | 17697 | 0.328 | 74.373 | HepG2 |
| NFYA | ZNF143 | 5163 | 17604 | 14071 | 0.328 | 57.911 | HepG2 |
| ATF3 | ETS1 | 3950 | 15603 | 9307 | 0.328 | 58.597 | HepG2 |
| ATF2 | HBP1 | 6252 | 23795 | 15317 | 0.327 | 50.801 | HepG2 |
| PHF20 | ZNF143 | 5157 | 17625 | 14071 | 0.327 | 53.246 | HepG2 |
| ZBED4 | ZNF639 | 4870 | 16556 | 13418 | 0.327 | 61.843 | HepG2 |
| HOXD1 | ZNF766 | 4388 | 13879 | 13024 | 0.326 | 50.984 | HepG2 |
| ATF2 | ZBTB21 | 4935 | 23795 | 9611 | 0.326 | 63.248 | HepG2 |
| AHR | MXD4 | 5344 | 12116 | 22169 | 0.326 | 50.628 | HepG2 |
| GATA4 | TBP | 5193 | 13900 | 18258 | 0.326 | 53.405 | HepG2 |

| | | | | | | | |
|---|---|---|---|---|---|---|---|
| MXD3 | ZIK1 | 4541 | 17726 | 10981 | 0.325 | 55.013 | HepG2 |
| MEF2A | ZNF503 | 4119 | 8647 | 18529 | 0.325 | 51.585 | HepG2 |
| HMG20B | ZNF124 | 4813 | 13676 | 16017 | 0.325 | 64.412 | HepG2 |
| HOXD1 | ZFP37 | 3767 | 13879 | 9678 | 0.325 | 70.139 | HepG2 |
| NKX3-1 | ZNF557 | 3447 | 13144 | 8561 | 0.325 | 66.713 | HepG2 |
| AHR | ZNF639 | 4143 | 12116 | 13418 | 0.325 | 69.576 | HepG2 |
| NFYA | ZHX2 | 4451 | 17604 | 10676 | 0.325 | 59.488 | HepG2 |
| GLI4 | ZNF547 | 4584 | 10442 | 19097 | 0.325 | 61.148 | HepG2 |
| ETS1 | MLX | 3965 | 9307 | 16042 | 0.324 | 51.837 | HepG2 |
| DRAP1 | ZHX2 | 5428 | 26223 | 10676 | 0.324 | 56.600 | HepG2 |
| HBP1 | MBD1 | 4940 | 15317 | 15149 | 0.324 | 50.920 | HepG2 |
| ELK1 | ZNF557 | 2956 | 9720 | 8561 | 0.324 | 54.308 | HepG2 |
| ARNTL | ATF7 | 4813 | 20368 | 10839 | 0.324 | 55.718 | HepG2 |
| ZNF343 | ZNF483 | 4882 | 13429 | 16926 | 0.324 | 53.230 | HepG2 |
| E2F1 | HOXA5 | 5089 | 11827 | 20899 | 0.324 | 50.694 | HepG2 |
| MEIS2 | ZFP37 | 4602 | 20907 | 9678 | 0.324 | 50.643 | HepG2 |
| SIX1 | ZFP1 | 4612 | 14572 | 13958 | 0.323 | 51.200 | HepG2 |
| ZBTB26 | ZNF639 | 4851 | 16775 | 13418 | 0.323 | 56.839 | HepG2 |
| AHR | KLF6 | 4840 | 12116 | 18502 | 0.323 | 56.970 | HepG2 |
| HINFP | ZNF548 | 4609 | 14261 | 14256 | 0.323 | 55.938 | HepG2 |
| ZNF343 | ZNF548 | 4470 | 13429 | 14256 | 0.323 | 55.345 | HepG2 |
| HHEX | NFIC | 3834 | 8174 | 17240 | 0.323 | 58.818 | HepG2 |
| ZNF25 | ZNF781 | 3066 | 7096 | 12710 | 0.323 | 63.016 | HepG2 |
| ZNF230 | ZNF782 | 4346 | 15509 | 11685 | 0.323 | 53.659 | HepG2 |
| ZNF230 | ZNF276 | 4159 | 15509 | 10710 | 0.323 | 56.849 | HepG2 |
| HOXD1 | ZNF557 | 3516 | 13879 | 8561 | 0.323 | 73.392 | HepG2 |
| GATA4 | SIX1 | 4585 | 13900 | 14572 | 0.322 | 52.067 | HepG2 |
| ZNF230 | ZNF510 | 4179 | 15509 | 10854 | 0.322 | 53.685 | HepG2 |
| MEF2A | SIX1 | 3615 | 8647 | 14572 | 0.322 | 76.635 | HepG2 |
| LBX2 | ZNF616 | 2558 | 7038 | 8986 | 0.322 | 65.977 | HepG2 |
| ZNF124 | ZNF639 | 4708 | 16017 | 13418 | 0.321 | 55.166 | HepG2 |
| ZKSCAN8 | ZNF639 | 5122 | 18988 | 13418 | 0.321 | 50.129 | HepG2 |
| NFAT5 | ZNF766 | 3500 | 9150 | 13024 | 0.321 | 61.135 | HepG2 |
| ZNF343 | ZNF556 | 5253 | 13429 | 20009 | 0.320 | 51.987 | HepG2 |
| HBP1 | ZNF580 | 6031 | 15317 | 23174 | 0.320 | 50.457 | HepG2 |
| ZNF142 | ZNF232 | 4998 | 14794 | 16478 | 0.320 | 53.405 | HepG2 |
| MEF2A | SIX4 | 3500 | 8647 | 13829 | 0.320 | 64.477 | HepG2 |
| KLF11 | TFE3 | 5476 | 12213 | 23975 | 0.320 | 57.753 | HepG2 |
| HMG20B | ZNF264 | 5111 | 13676 | 18666 | 0.320 | 50.046 | HepG2 |
| ETS1 | HNF4G | 4868 | 9307 | 24914 | 0.320 | 51.936 | HepG2 |
| ELK1 | HOXA5 | 4556 | 9720 | 20899 | 0.320 | 58.113 | HepG2 |
| ELK1 | ZNF264 | 4304 | 9720 | 18666 | 0.320 | 53.514 | HepG2 |
| SP1 | ZNF639 | 4687 | 16039 | 13418 | 0.319 | 62.165 | HepG2 |
| HMG20B | KLF6 | 5080 | 13676 | 18502 | 0.319 | 53.640 | HepG2 |
| PRDM10 | ZKSCAN8 | 5883 | 17873 | 18988 | 0.319 | 56.383 | HepG2 |
| DR1 | PHF20 | 4346 | 10534 | 17625 | 0.319 | 53.933 | HepG2 |
| HMG20B | THRB | 4820 | 13676 | 16710 | 0.319 | 51.433 | HepG2 |
| ZNF691 | ZNF782 | 4774 | 19190 | 11685 | 0.319 | 53.505 | HepG2 |
| EEA1 | TIGD6 | 3608 | 9299 | 13801 | 0.318 | 57.739 | HepG2 |
| E2F1 | ZNF766 | 3950 | 11827 | 13024 | 0.318 | 62.518 | HepG2 |
| ZHX2 | ZHX3 | 2701 | 10676 | 6761 | 0.318 | 70.702 | HepG2 |
| MXD1 | ZNF451 | 4849 | 18125 | 12838 | 0.318 | 53.473 | HepG2 |

| | | | | | | | |
|---|---|---|---|---|---|---|---|
| PHF20 | ZNF639 | 4885 | 17625 | 13418 | 0.318 | 50.680 | HepG2 |
| MEF2A | MLX | 3739 | 8647 | 16042 | 0.317 | 67.286 | HepG2 |
| E2F1 | PHF20 | 4583 | 11827 | 17625 | 0.317 | 57.700 | HepG2 |
| ETS1 | TBX3 | 3781 | 9307 | 15274 | 0.317 | 54.382 | HepG2 |
| MLX | TCF12 | 2722 | 16042 | 4594 | 0.317 | 73.993 | HepG2 |
| DR1 | NFYA | 4314 | 10534 | 17604 | 0.317 | 56.524 | HepG2 |
| PHF20 | SP2 | 4470 | 17625 | 11297 | 0.317 | 53.902 | HepG2 |
| KLF11 | ZNF639 | 4055 | 12213 | 13418 | 0.317 | 61.443 | HepG2 |
| ELK1 | NKX3-1 | 3580 | 9720 | 13144 | 0.317 | 67.220 | HepG2 |
| KLF12 | ZNF142 | 4727 | 15074 | 14794 | 0.317 | 50.745 | HepG2 |
| NFAT5 | NKX3-1 | 3471 | 9150 | 13144 | 0.317 | 56.497 | HepG2 |
| CAMTA2 | ZNF333 | 3012 | 8249 | 10988 | 0.316 | 59.008 | HepG2 |
| MLX | ZNF335 | 4944 | 16042 | 15231 | 0.316 | 58.082 | HepG2 |
| HHEX | MLX | 3621 | 8174 | 16042 | 0.316 | 53.919 | HepG2 |
| E2F1 | NKX3-1 | 3939 | 11827 | 13144 | 0.316 | 51.279 | HepG2 |
| MEF2A | MEF2D | 4445 | 8647 | 22926 | 0.316 | 54.367 | HepG2 |
| NCOA1 | ZNF414 | 4333 | 11085 | 16996 | 0.316 | 51.705 | HepG2 |
| ZNF483 | ZNF548 | 4900 | 16926 | 14256 | 0.315 | 50.730 | HepG2 |
| GMEB2 | KMT2B | 3392 | 5338 | 21702 | 0.315 | 62.145 | HepG2 |
| ZNF548 | ZNF782 | 4067 | 14256 | 11685 | 0.315 | 67.457 | HepG2 |
| ZFP1 | ZNF639 | 4311 | 13958 | 13418 | 0.315 | 58.678 | HepG2 |
| NR2F2 | SIX4 | 4989 | 18140 | 13829 | 0.315 | 55.530 | HepG2 |
| ETS1 | SMAD3 | 4377 | 9307 | 20760 | 0.315 | 52.076 | HepG2 |
| DZIP1 | ZNF280B | 3786 | 9278 | 15695 | 0.314 | 50.154 | HepG2 |
| ZNF335 | ZNF639 | 4482 | 15231 | 13418 | 0.314 | 56.668 | HepG2 |
| ZBTB44 | ZHX3 | 1872 | 5276 | 6761 | 0.313 | 72.010 | HepG2 |
| HOXD1 | KLF12 | 4532 | 13879 | 15074 | 0.313 | 56.190 | HepG2 |
| ZNF510 | ZNF543 | 4370 | 10854 | 17952 | 0.313 | 51.707 | HepG2 |
| MXD1 | ZIK1 | 4414 | 18125 | 10981 | 0.313 | 59.418 | HepG2 |
| SMAD1 | ZNF619 | 3622 | 11315 | 11856 | 0.313 | 62.668 | HepG2 |
| E2F1 | KMT2B | 5010 | 11827 | 21702 | 0.313 | 55.596 | HepG2 |
| HINFP | ZNF276 | 3864 | 14261 | 10710 | 0.313 | 59.538 | HepG2 |
| KMT2A | ZHX2 | 4894 | 23007 | 10676 | 0.312 | 51.546 | HepG2 |
| E2F4 | GMEB2 | 3378 | 21936 | 5338 | 0.312 | 68.351 | HepG2 |
| FOXJ3 | MEF2A | 4217 | 21114 | 8647 | 0.312 | 57.707 | HepG2 |
| MEF2A | ZKSCAN8 | 3994 | 8647 | 18988 | 0.312 | 56.088 | HepG2 |
| HINFP | ZNF782 | 4022 | 14261 | 11685 | 0.312 | 67.328 | HepG2 |
| MLXIP | NKX3-1 | 2649 | 5502 | 13144 | 0.311 | 65.247 | HepG2 |
| ZNF280B | ZNF485 | 3762 | 15695 | 9308 | 0.311 | 65.411 | HepG2 |
| ZNF343 | ZNF510 | 3757 | 13429 | 10854 | 0.311 | 54.975 | HepG2 |
| ZC3H8 | ZNF556 | 4326 | 9667 | 20009 | 0.311 | 55.078 | HepG2 |
| GMEB2 | NFYA | 3015 | 5338 | 17604 | 0.311 | 55.936 | HepG2 |
| ETS1 | ZNF609 | 4746 | 9307 | 25025 | 0.311 | 50.709 | HepG2 |
| ZNF264 | ZNF766 | 4845 | 18666 | 13024 | 0.311 | 58.453 | HepG2 |
| NKX3-1 | ZFP37 | 3504 | 13144 | 9678 | 0.311 | 51.554 | HepG2 |
| HIVEP1 | ZNF333 | 4328 | 17669 | 10988 | 0.311 | 54.299 | HepG2 |
| JRK | WIZ | 3933 | 15666 | 10254 | 0.310 | 50.916 | HepG2 |
| DNMT1 | EEA1 | 2816 | 8862 | 9299 | 0.310 | 74.623 | HepG2 |
| ZNF483 | ZNF510 | 4203 | 16926 | 10854 | 0.310 | 61.719 | HepG2 |
| ETS1 | HMG20B | 3494 | 9307 | 13676 | 0.310 | 57.818 | HepG2 |
| FOXO1 | MEF2A | 4621 | 25752 | 8647 | 0.310 | 51.969 | HepG2 |
| HMG20B | MEF2A | 3365 | 13676 | 8647 | 0.309 | 66.587 | HepG2 |

| | | | | | | | |
|---|---|---|---|---|---|---|---|
| ZNF276 | ZNF548 | 3823 | 10710 | 14256 | 0.309 | 56.714 | HepG2 |
| CAMTA2 | MXD1 | 3782 | 8249 | 18125 | 0.309 | 51.599 | HepG2 |
| ELK1 | ZNF766 | 3480 | 9720 | 13024 | 0.309 | 56.150 | HepG2 |
| GABPA | SP2 | 2977 | 8201 | 11297 | 0.309 | 56.178 | HepG2 |
| ZHX3 | ZNF792 | 3891 | 6761 | 23423 | 0.309 | 55.775 | HepG2 |
| ZBED4 | ZFP1 | 4700 | 16556 | 13958 | 0.309 | 51.022 | HepG2 |
| E2F1 | E2F2 | 2762 | 11827 | 6781 | 0.308 | 58.387 | HepG2 |
| KLF9 | ZBTB25 | 4064 | 8472 | 20508 | 0.308 | 50.458 | HepG2 |
| KLF9 | ZNF639 | 3287 | 8472 | 13418 | 0.308 | 63.789 | HepG2 |
| KLF6 | ZHX3 | 3447 | 18502 | 6761 | 0.308 | 61.645 | HepG2 |
| ZFP37 | ZNF766 | 3460 | 9678 | 13024 | 0.308 | 53.258 | HepG2 |
| ZBTB21 | ZHX2 | 3121 | 9611 | 10676 | 0.308 | 66.986 | HepG2 |
| DZIP1 | ZNF485 | 2863 | 9278 | 9308 | 0.308 | 56.280 | HepG2 |
| ZNF343 | ZNF782 | 3859 | 13429 | 11685 | 0.308 | 54.669 | HepG2 |
| KLF11 | ZFP1 | 4022 | 12213 | 13958 | 0.308 | 67.250 | HepG2 |
| MEF2A | MIXL1 | 4803 | 8647 | 28115 | 0.308 | 52.671 | HepG2 |
| ZNF138 | ZSCAN31 | 4263 | 8885 | 21564 | 0.308 | 54.002 | HepG2 |
| ZNF225 | ZNF274 | 4583 | 12359 | 17918 | 0.308 | 52.018 | HepG2 |
| CREB3 | NKX3-1 | 3051 | 7475 | 13144 | 0.308 | 69.781 | HepG2 |
| DNMT1 | IRF5 | 3054 | 8862 | 11111 | 0.308 | 54.501 | HepG2 |
| ZC3H8 | ZNF343 | 3506 | 9667 | 13429 | 0.308 | 61.713 | HepG2 |
| STAT5B | ZBTB44 | 1525 | 4656 | 5276 | 0.308 | 86.870 | HepG2 |
| GMEB2 | TFDP2 | 3372 | 5338 | 22516 | 0.308 | 53.001 | HepG2 |
| TIGD6 | ZNF451 | 4094 | 13801 | 12838 | 0.308 | 50.047 | HepG2 |
| ETS1 | TCF12 | 2009 | 9307 | 4594 | 0.307 | 82.521 | HepG2 |
| NFAT5 | ZNF142 | 3574 | 9150 | 14794 | 0.307 | 53.980 | HepG2 |
| SNAI1 | SNAPC4 | 3484 | 10844 | 11863 | 0.307 | 65.214 | HepG2 |
| MXD3 | NFAT5 | 3909 | 17726 | 9150 | 0.307 | 53.006 | HepG2 |
| ZHX2 | ZNF143 | 3760 | 10676 | 14071 | 0.307 | 61.014 | HepG2 |
| ZNF782 | ZZZ3 | 3062 | 11685 | 8530 | 0.307 | 57.567 | HepG2 |
| MLXIP | ZSCAN12 | 1858 | 5502 | 6684 | 0.306 | 78.276 | HepG2 |
| ZC3H8 | ZNF543 | 4036 | 9667 | 17952 | 0.306 | 54.350 | HepG2 |
| FOXO1 | TCF12 | 3331 | 25752 | 4594 | 0.306 | 55.971 | HepG2 |
| ZNF276 | ZNF691 | 4390 | 10710 | 19190 | 0.306 | 57.493 | HepG2 |
| MEF2A | ZFP1 | 3364 | 8647 | 13958 | 0.306 | 58.105 | HepG2 |
| MEF2A | ZNF511 | 3787 | 8647 | 17697 | 0.306 | 52.665 | HepG2 |
| ZNF276 | ZNF782 | 3421 | 10710 | 11685 | 0.306 | 53.360 | HepG2 |
| ZNF33B | ZNF44 | 3137 | 10715 | 9831 | 0.306 | 54.376 | HepG2 |
| AHR | MYRF | 2781 | 12116 | 6833 | 0.306 | 64.454 | HepG2 |
| ZFP1 | ZNF335 | 4455 | 13958 | 15231 | 0.306 | 53.768 | HepG2 |
| ZNF678 | ZNF782 | 2901 | 7739 | 11685 | 0.305 | 67.944 | HepG2 |
| ZNF503 | ZNF766 | 4737 | 18529 | 13024 | 0.305 | 50.293 | HepG2 |
| TEAD2 | ZNF773 | 1430 | 3288 | 6695 | 0.305 | 90.105 | HepG2 |
| THAP9 | ZNF225 | 4922 | 21105 | 12359 | 0.305 | 50.618 | HepG2 |
| ZHX2 | ZNF639 | 3646 | 10676 | 13418 | 0.305 | 57.464 | HepG2 |
| SATB2 | ZNF232 | 3501 | 8020 | 16478 | 0.305 | 67.430 | HepG2 |
| KMT2B | ZHX3 | 3687 | 21702 | 6761 | 0.304 | 64.118 | HepG2 |
| ELK1 | KMT2B | 4420 | 9720 | 21702 | 0.304 | 53.035 | HepG2 |
| ZNF556 | ZZZ3 | 3975 | 20009 | 8530 | 0.304 | 56.366 | HepG2 |
| HOXD1 | ZNF142 | 4359 | 13879 | 14794 | 0.304 | 54.568 | HepG2 |
| STAT5B | ZHX3 | 1705 | 4656 | 6761 | 0.304 | 81.246 | HepG2 |
| E2F2 | NFAT5 | 2392 | 6781 | 9150 | 0.304 | 61.648 | HepG2 |

| | | | | | | | |
|---|---|---|---|---|---|---|---|
| HINFP | ZZZ3 | 3346 | 14261 | 8530 | 0.303 | 56.472 | HepG2 |
| DLX6 | MEF2A | 3951 | 19647 | 8647 | 0.303 | 54.927 | HepG2 |
| SIX4 | ZHX3 | 2929 | 13829 | 6761 | 0.303 | 59.652 | HepG2 |
| LBX2 | ZIK1 | 2661 | 7038 | 10981 | 0.303 | 62.497 | HepG2 |
| ZNF274 | ZNF451 | 4589 | 17918 | 12838 | 0.303 | 52.647 | HepG2 |
| HHEX | SIX4 | 3214 | 8174 | 13829 | 0.302 | 57.908 | HepG2 |
| KLF9 | ZNF205 | 3935 | 8472 | 20027 | 0.302 | 62.391 | HepG2 |
| DNMT1 | TIGD6 | 3340 | 8862 | 13801 | 0.302 | 60.546 | HepG2 |
| ETS1 | SIX4 | 3426 | 9307 | 13829 | 0.302 | 55.519 | HepG2 |
| AKNA | MTF1 | 1218 | 3772 | 4316 | 0.302 | 96.871 | HepG2 |
| ETS1 | ZKSCAN8 | 4010 | 9307 | 18988 | 0.302 | 51.055 | HepG2 |
| NKX3-1 | ZSCAN12 | 2824 | 13144 | 6684 | 0.301 | 63.685 | HepG2 |
| HOXD1 | ZNF639 | 4108 | 13879 | 13418 | 0.301 | 51.385 | HepG2 |
| PHF20 | ZZZ3 | 3691 | 17625 | 8530 | 0.301 | 53.681 | HepG2 |
| ATF7 | ZNF691 | 4340 | 10839 | 19190 | 0.301 | 51.771 | HepG2 |
| HMG20B | REST | 4568 | 13676 | 16852 | 0.301 | 51.951 | HepG2 |
| ZNF343 | ZNF691 | 4830 | 13429 | 19190 | 0.301 | 51.979 | HepG2 |
| TIGD6 | ZIK1 | 3703 | 13801 | 10981 | 0.301 | 50.359 | HepG2 |
| LBX2 | ZNF451 | 2859 | 7038 | 12838 | 0.301 | 60.722 | HepG2 |
| TIGD6 | WIZ | 3577 | 13801 | 10254 | 0.301 | 55.318 | HepG2 |
| ZNF557 | ZNF766 | 3175 | 8561 | 13024 | 0.301 | 59.521 | HepG2 |
| ZHX3 | ZNF331 | 4460 | 6761 | 32566 | 0.301 | 50.934 | HepG2 |
| GMEB2 | PHF20 | 2912 | 5338 | 17625 | 0.300 | 61.491 | HepG2 |
| ATF7 | HINFP | 3731 | 10839 | 14261 | 0.300 | 55.738 | HepG2 |
| MAFF | MAFK | 19903 | 27875 | 27212 | 0.723 | 177.028 | K562 |
| TAL1 | TCF12 | 19712 | 29475 | 28573 | 0.679 | 185.816 | K562 |
| TCF12 | TCF3 | 19889 | 28573 | 33040 | 0.647 | 159.101 | K562 |
| MAFF | MAFG | 22716 | 27875 | 45814 | 0.636 | 127.799 | K562 |
| TAL1 | TCF3 | 19737 | 29475 | 33040 | 0.632 | 158.178 | K562 |
| USF1 | USF2 | 12325 | 22381 | 17154 | 0.629 | 182.878 | K562 |
| MAFG | MAFK | 20990 | 45814 | 27212 | 0.594 | 124.103 | K562 |
| MITF | TFE3 | 16056 | 34701 | 22683 | 0.572 | 140.066 | K562 |
| MEIS2 | PBX2 | 24894 | 53049 | 36550 | 0.565 | 97.906 | K562 |
| TCF12 | TEAD4 | 17744 | 28573 | 36089 | 0.553 | 128.129 | K562 |
| NFYA | NFYB | 3703 | 4570 | 9997 | 0.548 | 185.207 | K562 |
| TCF3 | TEAD4 | 18911 | 33040 | 36089 | 0.548 | 132.300 | K562 |
| TAL1 | TEAD4 | 17840 | 29475 | 36089 | 0.547 | 141.793 | K562 |
| NR2F1 | NR2F6 | 16517 | 38727 | 23922 | 0.543 | 122.715 | K562 |
| JUN | JUNB | 6849 | 11072 | 15517 | 0.523 | 196.062 | K562 |
| CEBPB | CEBPG | 25038 | 50240 | 46899 | 0.516 | 64.370 | K562 |
| E2F6 | MAX | 23180 | 33733 | 59913 | 0.516 | 77.399 | K562 |
| CREB1 | CREM | 13505 | 17620 | 39050 | 0.515 | 121.823 | K562 |
| MEIS2 | PKNOX1 | 26324 | 53049 | 52253 | 0.500 | 60.246 | K562 |
| FOSL1 | JUN | 5935 | 12759 | 11072 | 0.499 | 170.572 | K562 |
| FOS | NFYA | 2953 | 7848 | 4570 | 0.493 | 173.805 | K562 |
| FOSL1 | JUNB | 6901 | 12759 | 15517 | 0.490 | 128.638 | K562 |
| JUN | JUND | 9432 | 11072 | 35378 | 0.477 | 134.815 | K562 |
| CREM | MAZ | 18884 | 39050 | 40397 | 0.475 | 77.403 | K562 |
| NR2F2 | NR2F6 | 10595 | 20900 | 23922 | 0.474 | 129.048 | K562 |
| E4F1 | ELF4 | 12802 | 35525 | 20589 | 0.473 | 102.417 | K562 |
| ATF3 | JUN | 6775 | 18779 | 11072 | 0.470 | 135.989 | K562 |
| PBX2 | PKNOX1 | 20488 | 36550 | 52253 | 0.469 | 68.802 | K562 |

| | | | | | | | |
|---|---|---|---|---|---|---|---|
| GATA1 | TAL1 | 9746 | 14675 | 29475 | 0.469 | 107.118 | K562 |
| JUNB | JUND | 10940 | 15517 | 35378 | 0.467 | 106.468 | K562 |
| MAFG | NFE2 | 21204 | 45814 | 45103 | 0.466 | 63.492 | K562 |
| MAZ | VEZF1 | 19982 | 40397 | 46847 | 0.459 | 71.832 | K562 |
| SOX6 | TEAD4 | 16813 | 37291 | 36089 | 0.458 | 81.656 | K562 |
| CTCF | DEAF1 | 16004 | 59802 | 20812 | 0.454 | 93.310 | K562 |
| DEAF1 | ZNF143 | 11279 | 20812 | 29808 | 0.453 | 121.332 | K562 |
| STAT5A | TEAD4 | 9974 | 13551 | 36089 | 0.451 | 120.491 | K562 |
| GATA1 | TCF12 | 9216 | 14675 | 28573 | 0.450 | 125.301 | K562 |
| SOX6 | TCF3 | 15717 | 37291 | 33040 | 0.448 | 71.717 | K562 |
| GATA1 | GATA2 | 7933 | 14675 | 21627 | 0.445 | 119.388 | K562 |
| KDM5B | SMAD5 | 9930 | 22367 | 22343 | 0.444 | 96.951 | K562 |
| CREM | E2F1 | 13163 | 39050 | 22524 | 0.444 | 75.310 | K562 |
| GATA1 | TEAD4 | 10170 | 14675 | 36089 | 0.442 | 99.880 | K562 |
| CTCF | ZNF143 | 18645 | 59802 | 29808 | 0.442 | 68.537 | K562 |
| STAT5A | TAL1 | 8823 | 13551 | 29475 | 0.441 | 117.873 | K562 |
| STAT5A | TCF12 | 8665 | 13551 | 28573 | 0.440 | 117.188 | K562 |
| STAT5A | TCF3 | 9310 | 13551 | 33040 | 0.440 | 129.113 | K562 |
| ATF7 | CREM | 18795 | 47196 | 39050 | 0.438 | 50.527 | K562 |
| ATF1 | CREM | 11199 | 16924 | 39050 | 0.436 | 84.582 | K562 |
| CREB1 | E2F1 | 8651 | 17620 | 22524 | 0.434 | 124.724 | K562 |
| NFYA | SP2 | 1719 | 4570 | 3435 | 0.434 | 177.218 | K562 |
| GATA1 | STAT5A | 6106 | 14675 | 13551 | 0.433 | 134.982 | K562 |
| MBD2 | NEUROD1 | 6379 | 15550 | 14154 | 0.430 | 119.029 | K562 |
| GATA1 | TCF3 | 9457 | 14675 | 33040 | 0.429 | 100.593 | K562 |
| NR2F2 | STAT5A | 7183 | 20900 | 13551 | 0.427 | 127.621 | K562 |
| KLF16 | STAT5A | 6629 | 17849 | 13551 | 0.426 | 144.240 | K562 |
| CC2D1A | NCOA1 | 10553 | 26436 | 23304 | 0.425 | 102.469 | K562 |
| NR2F1 | NR2F2 | 11946 | 38727 | 20900 | 0.420 | 62.146 | K562 |
| SMAD5 | TBP | 9300 | 22343 | 22020 | 0.419 | 126.705 | K562 |
| ZNF215 | ZNF311 | 3047 | 7306 | 7247 | 0.419 | 156.430 | K562 |
| ELF1 | ELF4 | 8626 | 20717 | 20589 | 0.418 | 102.836 | K562 |
| KLF16 | TEAD4 | 10589 | 17849 | 36089 | 0.417 | 72.962 | K562 |
| FOXM1 | STAT5A | 6532 | 18104 | 13551 | 0.417 | 133.079 | K562 |
| NR2F2 | TEAD4 | 11399 | 20900 | 36089 | 0.415 | 75.342 | K562 |
| GMEB1 | MNT | 13793 | 29364 | 37785 | 0.414 | 68.458 | K562 |
| ATF1 | E2F1 | 8082 | 16924 | 22524 | 0.414 | 100.281 | K562 |
| ATF7 | JUND | 16804 | 47196 | 35378 | 0.411 | 58.538 | K562 |
| ATF3 | FOSL1 | 6362 | 18779 | 12759 | 0.411 | 112.732 | K562 |
| FOS | JUN | 3831 | 7848 | 11072 | 0.411 | 123.408 | K562 |
| FOSL1 | JUND | 8693 | 12759 | 35378 | 0.409 | 101.227 | K562 |
| CREB1 | ZBTB12 | 5066 | 17620 | 8719 | 0.409 | 142.655 | K562 |
| GATA1 | SOX6 | 9552 | 14675 | 37291 | 0.408 | 98.211 | K562 |
| LEF1 | SOX6 | 11678 | 21938 | 37291 | 0.408 | 90.602 | K562 |
| ATF7 | E2F1 | 13258 | 47196 | 22524 | 0.407 | 76.342 | K562 |
| CREB1 | MBD2 | 6710 | 17620 | 15550 | 0.405 | 103.566 | K562 |
| GATA1 | KLF16 | 6553 | 14675 | 17849 | 0.405 | 100.356 | K562 |
| SOX6 | TCF12 | 13210 | 37291 | 28573 | 0.405 | 55.205 | K562 |
| NR2C1 | NR2C2 | 13492 | 24799 | 45215 | 0.403 | 60.960 | K562 |
| ATF1 | CREB1 | 6949 | 16924 | 17620 | 0.402 | 112.731 | K562 |
| KLF16 | TCF3 | 9761 | 17849 | 33040 | 0.402 | 86.371 | K562 |
| KLF6 | SHOX2 | 2833 | 7751 | 6459 | 0.400 | 163.315 | K562 |

| | | | | | | | |
|---|---|---|---|---|---|---|---|
| KLF16 | TCF12 | 9024 | 17849 | 28573 | 0.400 | 107.057 | K562 |
| ARID2 | DEAF1 | 6296 | 11951 | 20812 | 0.399 | 98.513 | K562 |
| MAZ | ZNF143 | 13832 | 40397 | 29808 | 0.399 | 66.851 | K562 |
| FOS | NFYB | 3509 | 7848 | 9997 | 0.396 | 135.318 | K562 |
| MITF | USF1 | 11032 | 34701 | 22381 | 0.396 | 78.438 | K562 |
| ZNF148 | ZNF281 | 12143 | 23453 | 40285 | 0.395 | 72.645 | K562 |
| CTCF | CTCFL | 11220 | 59802 | 13502 | 0.395 | 81.595 | K562 |
| FOXM1 | GATA1 | 6435 | 18104 | 14675 | 0.395 | 114.763 | K562 |
| MAX | MYC | 11729 | 59913 | 14737 | 0.395 | 63.206 | K562 |
| BHLHE40 | MAX | 16097 | 27807 | 59913 | 0.394 | 52.704 | K562 |
| SNAPC5 | STAT5B | 246 | 642 | 607 | 0.394 | 166.934 | K562 |
| LEF1 | NCOA1 | 8870 | 21938 | 23304 | 0.392 | 89.200 | K562 |
| FOXM1 | NCOA1 | 8046 | 18104 | 23304 | 0.392 | 99.172 | K562 |
| ATF7 | SKIL | 13240 | 47196 | 24237 | 0.391 | 63.574 | K562 |
| SOX6 | TAL1 | 12946 | 37291 | 29475 | 0.390 | 58.230 | K562 |
| NR2F2 | TCF3 | 10218 | 20900 | 33040 | 0.389 | 90.250 | K562 |
| ATF3 | CREM | 10528 | 18779 | 39050 | 0.389 | 76.080 | K562 |
| ATF3 | JUND | 10009 | 18779 | 35378 | 0.388 | 75.396 | K562 |
| E4F1 | ELF1 | 10508 | 35525 | 20717 | 0.387 | 95.184 | K562 |
| NR2F2 | TCF12 | 9457 | 20900 | 28573 | 0.387 | 85.278 | K562 |
| FOXM1 | TCF12 | 8754 | 18104 | 28573 | 0.385 | 90.810 | K562 |
| FOXM1 | SOX6 | 9999 | 18104 | 37291 | 0.385 | 80.043 | K562 |
| CREM | NEUROD1 | 9047 | 39050 | 14154 | 0.385 | 78.438 | K562 |
| ATF1 | JUND | 9416 | 16924 | 35378 | 0.385 | 94.832 | K562 |
| KLF16 | TAL1 | 8810 | 17849 | 29475 | 0.384 | 78.865 | K562 |
| ATF3 | JUNB | 6542 | 18779 | 15517 | 0.383 | 108.628 | K562 |
| CREM | MBD2 | 9425 | 39050 | 15550 | 0.382 | 94.694 | K562 |
| FOXM1 | TEAD4 | 9735 | 18104 | 36089 | 0.381 | 77.123 | K562 |
| NEUROD1 | SP1 | 4020 | 14154 | 7876 | 0.381 | 124.082 | K562 |
| E2F6 | MGA | 12802 | 33733 | 33582 | 0.380 | 55.349 | K562 |
| SOX6 | STAT5A | 8534 | 37291 | 13551 | 0.380 | 105.386 | K562 |
| GABPA | MBD2 | 6063 | 16492 | 15550 | 0.379 | 102.887 | K562 |
| KLF16 | SOX6 | 9757 | 17849 | 37291 | 0.378 | 89.646 | K562 |
| CREB1 | GABPA | 6446 | 17620 | 16492 | 0.378 | 108.884 | K562 |
| FOXM1 | LEF1 | 7529 | 18104 | 21938 | 0.378 | 92.686 | K562 |
| GMEB1 | ZBTB2 | 8754 | 29364 | 18463 | 0.376 | 84.997 | K562 |
| ARNT | NCOA1 | 7334 | 16351 | 23304 | 0.376 | 92.700 | K562 |
| NR2F2 | TAL1 | 9323 | 20900 | 29475 | 0.376 | 77.028 | K562 |
| FOXM1 | KLF16 | 6747 | 18104 | 17849 | 0.375 | 96.659 | K562 |
| KLF6 | ZNF239 | 2645 | 7751 | 6412 | 0.375 | 131.698 | K562 |
| NR2C1 | NR2F1 | 11612 | 24799 | 38727 | 0.375 | 73.864 | K562 |
| SOX6 | ZEB2 | 12003 | 37291 | 27537 | 0.375 | 59.756 | K562 |
| MAZ | NEUROD1 | 8955 | 40397 | 14154 | 0.374 | 82.829 | K562 |
| ATF1 | JUN | 5112 | 16924 | 11072 | 0.373 | 110.674 | K562 |
| MBD2 | SP1 | 4129 | 15550 | 7876 | 0.373 | 117.493 | K562 |
| ETV1 | ETV5 | 7175 | 21785 | 17004 | 0.373 | 94.643 | K562 |
| ARNT | LEF1 | 7055 | 16351 | 21938 | 0.373 | 102.384 | K562 |
| CREM | GABPA | 9444 | 39050 | 16492 | 0.372 | 80.593 | K562 |
| MNT | MYC | 8772 | 37785 | 14737 | 0.372 | 86.514 | K562 |
| CREM | SOX6 | 14149 | 39050 | 37291 | 0.371 | 53.014 | K562 |
| E4F1 | MNT | 13539 | 35525 | 37785 | 0.370 | 52.516 | K562 |
| NCOA1 | SOX6 | 10892 | 23304 | 37291 | 0.369 | 70.775 | K562 |

| | | | | | | | |
|---|---|---|---|---|---|---|---|
| E2F6 | MYC | 8237 | 33733 | 14737 | 0.369 | 119.267 | K562 |
| ZBTB17 | ZBTB9 | 5527 | 15161 | 14782 | 0.369 | 116.561 | K562 |
| CREB1 | NEUROD1 | 5830 | 17620 | 14154 | 0.369 | 101.579 | K562 |
| ATF1 | ATF7 | 10432 | 16924 | 47196 | 0.369 | 67.155 | K562 |
| KLF16 | NCOA1 | 7523 | 17849 | 23304 | 0.369 | 88.212 | K562 |
| ATF7 | CREB1 | 10576 | 47196 | 17620 | 0.367 | 78.536 | K562 |
| BHLHE40 | CREM | 12083 | 27807 | 39050 | 0.367 | 55.873 | K562 |
| ERF | ZBTB9 | 4436 | 9999 | 14782 | 0.365 | 129.323 | K562 |
| MITF | USF2 | 8890 | 34701 | 17154 | 0.364 | 83.447 | K562 |
| SHOX2 | ZNF215 | 2503 | 6459 | 7306 | 0.364 | 133.227 | K562 |
| FOXM1 | TCF3 | 8907 | 18104 | 33040 | 0.364 | 61.908 | K562 |
| FOXM1 | TAL1 | 8362 | 18104 | 29475 | 0.362 | 75.712 | K562 |
| BACH1 | NFE2L1 | 2143 | 4739 | 7399 | 0.362 | 157.676 | K562 |
| NCOA1 | STAT5A | 6423 | 23304 | 13551 | 0.361 | 129.609 | K562 |
| LEF1 | TEAD4 | 10138 | 21938 | 36089 | 0.360 | 64.445 | K562 |
| LEF1 | STAT5A | 6196 | 21938 | 13551 | 0.359 | 103.767 | K562 |
| KLF16 | NR2F2 | 6939 | 17849 | 20900 | 0.359 | 83.509 | K562 |
| KDM5B | TBP | 7941 | 22367 | 22020 | 0.358 | 74.137 | K562 |
| E4F1 | ETV1 | 9945 | 35525 | 21785 | 0.357 | 66.326 | K562 |
| TBX18 | ZNF76 | 2153 | 6175 | 5886 | 0.357 | 130.638 | K562 |
| TSHZ1 | ZNF79 | 1060 | 2648 | 3330 | 0.357 | 120.357 | K562 |
| SHOX2 | ZNF583 | 2792 | 6459 | 9537 | 0.356 | 97.935 | K562 |
| SHOX2 | ZNF239 | 2289 | 6459 | 6412 | 0.356 | 127.512 | K562 |
| CC2D1A | FOXM1 | 7774 | 26436 | 18104 | 0.355 | 74.358 | K562 |
| HIVEP1 | TSHZ1 | 1220 | 4459 | 2648 | 0.355 | 160.544 | K562 |
| ETV1 | GABPA | 6726 | 21785 | 16492 | 0.355 | 91.838 | K562 |
| CREB3L1 | MNT | 10270 | 22214 | 37785 | 0.354 | 69.947 | K562 |
| BCL6 | ZNF257 | 1220 | 2832 | 4190 | 0.354 | 164.736 | K562 |
| MBD2 | TBP | 6538 | 15550 | 22020 | 0.353 | 89.333 | K562 |
| CREB3L1 | SKIL | 8185 | 22214 | 24237 | 0.353 | 75.698 | K562 |
| ETV5 | GABPA | 5894 | 17004 | 16492 | 0.352 | 101.828 | K562 |
| E2F1 | ZBTB12 | 4924 | 22524 | 8719 | 0.351 | 107.956 | K562 |
| HES1 | LEF1 | 5499 | 11186 | 21938 | 0.351 | 106.442 | K562 |
| BHLHE40 | MAZ | 11762 | 27807 | 40397 | 0.351 | 54.988 | K562 |
| ZBTB9 | ZNF583 | 4161 | 14782 | 9537 | 0.350 | 111.362 | K562 |
| SHOX2 | TSHZ1 | 1449 | 6459 | 2648 | 0.350 | 141.976 | K562 |
| ZNF583 | ZNF76 | 2623 | 9537 | 5886 | 0.350 | 103.885 | K562 |
| KLF6 | ZNF583 | 3010 | 7751 | 9537 | 0.350 | 115.876 | K562 |
| BCL6 | ZNF79 | 1075 | 2832 | 3330 | 0.350 | 149.303 | K562 |
| GABPA | ZBTB11 | 3879 | 16492 | 7469 | 0.350 | 99.314 | K562 |
| ATF3 | ATF7 | 10397 | 18779 | 47196 | 0.349 | 62.444 | K562 |
| SHOX2 | ZNF79 | 1617 | 6459 | 3330 | 0.349 | 158.817 | K562 |
| KLF6 | TEAD1 | 3498 | 7751 | 12990 | 0.349 | 105.186 | K562 |
| TBX18 | ZNF583 | 2673 | 6175 | 9537 | 0.348 | 113.812 | K562 |
| LEF1 | TCF12 | 8703 | 21938 | 28573 | 0.348 | 61.344 | K562 |
| ELF4 | ETV1 | 7360 | 20589 | 21785 | 0.348 | 78.601 | K562 |
| BCL6 | SHOX2 | 1484 | 2832 | 6459 | 0.347 | 126.217 | K562 |
| E4F1 | MBD2 | 8154 | 35525 | 15550 | 0.347 | 69.303 | K562 |
| CREB1 | MAZ | 9230 | 17620 | 40397 | 0.346 | 76.084 | K562 |
| NR2C1 | NR2F6 | 8426 | 24799 | 23922 | 0.346 | 76.612 | K562 |
| FOS | FOSL1 | 3459 | 7848 | 12759 | 0.346 | 108.446 | K562 |
| LEF1 | ZNF281 | 10263 | 21938 | 40285 | 0.345 | 57.542 | K562 |

| | | | | | | | |
|---|---|---|---|---|---|---|---|
| NFYA | SP1 | 2069 | 4570 | 7876 | 0.345 | 127.247 | K562 |
| FOS | SP2 | 1790 | 7848 | 3435 | 0.345 | 109.248 | K562 |
| BCL6 | TSHZ1 | 943 | 2832 | 2648 | 0.344 | 125.251 | K562 |
| GABPA | NEUROD1 | 5261 | 16492 | 14154 | 0.344 | 93.207 | K562 |
| ATF1 | ZBTB12 | 4177 | 16924 | 8719 | 0.344 | 112.147 | K562 |
| CTCFL | ZNF143 | 6898 | 13502 | 29808 | 0.344 | 76.169 | K562 |
| ATF3 | CREB1 | 6250 | 18779 | 17620 | 0.344 | 92.748 | K562 |
| SHOX2 | ZNF311 | 2350 | 6459 | 7247 | 0.343 | 108.824 | K562 |
| HIVEP1 | TBX18 | 1799 | 4459 | 6175 | 0.343 | 134.715 | K562 |
| LEF1 | TCF3 | 9227 | 21938 | 33040 | 0.343 | 59.375 | K562 |
| E2F4 | HMGN3 | 4709 | 10379 | 18201 | 0.343 | 91.565 | K562 |
| NR2F2 | SOX6 | 9555 | 20900 | 37291 | 0.342 | 58.615 | K562 |
| CC2D1A | KLF16 | 7431 | 26436 | 17849 | 0.342 | 109.936 | K562 |
| ATF7 | CREB3L1 | 11057 | 47196 | 22214 | 0.341 | 54.045 | K562 |
| BCL6 | ZNF239 | 1455 | 2832 | 6412 | 0.341 | 131.331 | K562 |
| ATF1 | ATF3 | 6067 | 16924 | 18779 | 0.340 | 93.189 | K562 |
| HIVEP1 | ZNF76 | 1743 | 4459 | 5886 | 0.340 | 125.545 | K562 |
| E4F1 | GABPA | 8235 | 35525 | 16492 | 0.340 | 69.801 | K562 |
| HIVEP1 | KLF6 | 2000 | 4459 | 7751 | 0.340 | 120.016 | K562 |
| E4F1 | NCOA1 | 9780 | 35525 | 23304 | 0.340 | 59.148 | K562 |
| MBD2 | MNT | 8236 | 15550 | 37785 | 0.340 | 73.387 | K562 |
| MAZ | TBP | 10124 | 40397 | 22020 | 0.339 | 62.035 | K562 |
| NCOA1 | TEAD4 | 9843 | 23304 | 36089 | 0.339 | 59.622 | K562 |
| E4F1 | LEF1 | 9475 | 35525 | 21938 | 0.339 | 61.837 | K562 |
| KLF6 | ZNF79 | 1722 | 7751 | 3330 | 0.339 | 123.784 | K562 |
| GATA1 | NR2F2 | 5934 | 14675 | 20900 | 0.339 | 89.208 | K562 |
| HES1 | SOX6 | 6916 | 11186 | 37291 | 0.339 | 72.575 | K562 |
| SKIL | SOX6 | 10180 | 24237 | 37291 | 0.339 | 59.376 | K562 |
| ARNT | FOXM1 | 5825 | 16351 | 18104 | 0.339 | 115.278 | K562 |
| ATF7 | NEUROD1 | 8749 | 47196 | 14154 | 0.339 | 64.832 | K562 |
| E4F1 | GMEB1 | 10932 | 35525 | 29364 | 0.338 | 53.625 | K562 |
| TEAD1 | ZNF239 | 3083 | 12990 | 6412 | 0.338 | 100.726 | K562 |
| NFE2 | NFE2L1 | 6166 | 45103 | 7399 | 0.338 | 87.119 | K562 |
| MAZ | MYC | 8230 | 40397 | 14737 | 0.337 | 57.173 | K562 |
| GABPA | MAZ | 8706 | 16492 | 40397 | 0.337 | 70.986 | K562 |
| E2F4 | E2F6 | 6309 | 10379 | 33733 | 0.337 | 98.126 | K562 |
| ARID2 | CC2D1A | 5986 | 11951 | 26436 | 0.337 | 68.273 | K562 |
| E2F3 | ERF | 3123 | 8607 | 9999 | 0.337 | 106.431 | K562 |
| CREM | MYC | 8073 | 39050 | 14737 | 0.337 | 69.885 | K562 |
| CREB1 | MNT | 8667 | 17620 | 37785 | 0.336 | 63.280 | K562 |
| ATF3 | FOS | 4074 | 18779 | 7848 | 0.336 | 104.143 | K562 |
| CREB1 | GMEB1 | 7628 | 17620 | 29364 | 0.335 | 73.881 | K562 |
| ESRRA | SKIL | 9572 | 33637 | 24237 | 0.335 | 54.823 | K562 |
| JUND | TEAD4 | 11957 | 35378 | 36089 | 0.335 | 51.358 | K562 |
| FOXM1 | NR2F2 | 6508 | 18104 | 20900 | 0.335 | 78.333 | K562 |
| GMEB1 | MBD2 | 7145 | 29364 | 15550 | 0.334 | 85.081 | K562 |
| HIVEP1 | ZNF79 | 1288 | 4459 | 3330 | 0.334 | 117.355 | K562 |
| SHOX2 | TBX18 | 2108 | 6459 | 6175 | 0.334 | 111.300 | K562 |
| CREB1 | E2F4 | 4512 | 17620 | 10379 | 0.334 | 112.541 | K562 |
| KLF6 | TSHZ1 | 1510 | 7751 | 2648 | 0.333 | 116.006 | K562 |
| E4F1 | NEUROD1 | 7471 | 35525 | 14154 | 0.333 | 73.127 | K562 |
| GATA2 | TEAD4 | 9304 | 21627 | 36089 | 0.333 | 67.305 | K562 |

| | | | | | | | |
|---|---|---|---|---|---|---|---|
| HIVEP1 | ZNF583 | 2171 | 4459 | 9537 | 0.333 | 114.945 | K562 |
| MAZ | MBD2 | 8344 | 40397 | 15550 | 0.333 | 65.281 | K562 |
| CREM | KLF16 | 8785 | 39050 | 17849 | 0.333 | 53.225 | K562 |
| ATF7 | MBD2 | 8998 | 47196 | 15550 | 0.332 | 77.488 | K562 |
| KLF6 | TBX18 | 2297 | 7751 | 6175 | 0.332 | 110.842 | K562 |
| CREB5 | PATZ1 | 729 | 1990 | 2428 | 0.332 | 130.988 | K562 |
| E4F1 | FOXK2 | 9516 | 35525 | 23181 | 0.332 | 53.895 | K562 |
| CC2D1A | CHAMP1 | 6534 | 26436 | 14730 | 0.331 | 65.850 | K562 |
| FOXM1 | HES1 | 4710 | 18104 | 11186 | 0.331 | 104.445 | K562 |
| MNT | NCOA1 | 9820 | 37785 | 23304 | 0.331 | 63.845 | K562 |
| LEF1 | NR2F2 | 7071 | 21938 | 20900 | 0.330 | 88.788 | K562 |
| E2F4 | TBP | 4988 | 10379 | 22020 | 0.330 | 111.848 | K562 |
| HIVEP1 | SHOX2 | 1770 | 4459 | 6459 | 0.330 | 113.747 | K562 |
| ELF1 | GABPA | 6096 | 20717 | 16492 | 0.330 | 96.149 | K562 |
| ZNF175 | ZNF589 | 6573 | 18986 | 20981 | 0.329 | 63.197 | K562 |
| GATA2 | TCF12 | 8178 | 21627 | 28573 | 0.329 | 74.736 | K562 |
| ETS1 | GABPA | 4493 | 11321 | 16492 | 0.329 | 89.265 | K562 |
| KLF6 | ZNF215 | 2472 | 7751 | 7306 | 0.328 | 113.483 | K562 |
| NCOA1 | NR2F2 | 7248 | 23304 | 20900 | 0.328 | 70.108 | K562 |
| TEAD4 | ZEB2 | 10337 | 36089 | 27537 | 0.328 | 50.758 | K562 |
| GATA2 | SOX6 | 9303 | 21627 | 37291 | 0.328 | 52.487 | K562 |
| BCL6 | ZNF354B | 914 | 2832 | 2758 | 0.327 | 151.426 | K562 |
| ZNF239 | ZNF79 | 1510 | 6412 | 3330 | 0.327 | 111.166 | K562 |
| MXI1 | MYC | 3777 | 9080 | 14737 | 0.327 | 104.868 | K562 |
| SP1 | SP2 | 1698 | 7876 | 3435 | 0.326 | 123.072 | K562 |
| BCL6 | KLF6 | 1529 | 2832 | 7751 | 0.326 | 121.882 | K562 |
| FOS | JUNB | 3598 | 7848 | 15517 | 0.326 | 132.523 | K562 |
| BHLHE40 | MYC | 6592 | 27807 | 14737 | 0.326 | 66.968 | K562 |
| SHOX2 | ZNF257 | 1694 | 6459 | 4190 | 0.326 | 119.828 | K562 |
| BHLHE40 | USF1 | 8122 | 27807 | 22381 | 0.326 | 67.509 | K562 |
| LEF1 | ZNF318 | 4791 | 21938 | 9879 | 0.325 | 91.662 | K562 |
| ELF1 | ETV1 | 6911 | 20717 | 21785 | 0.325 | 70.900 | K562 |
| ARNT | NFATC3 | 4811 | 16351 | 13378 | 0.325 | 89.539 | K562 |
| HMGN3 | MAZ | 8819 | 18201 | 40397 | 0.325 | 68.094 | K562 |
| TSHZ1 | ZNF257 | 1081 | 2648 | 4190 | 0.325 | 138.452 | K562 |
| ASH1L | NFYA | 1958 | 7970 | 4570 | 0.324 | 142.323 | K562 |
| LEF1 | ZEB2 | 7970 | 21938 | 27537 | 0.324 | 54.382 | K562 |
| BCL6 | ZNF507 | 1275 | 2832 | 5465 | 0.324 | 109.294 | K562 |
| GATA1 | HES1 | 4147 | 14675 | 11186 | 0.324 | 90.281 | K562 |
| NEUROD1 | SOX6 | 7434 | 14154 | 37291 | 0.324 | 75.169 | K562 |
| GATA1 | LEF1 | 5803 | 14675 | 21938 | 0.323 | 79.355 | K562 |
| E2F4 | MAZ | 6618 | 10379 | 40397 | 0.323 | 61.865 | K562 |
| GATA2 | STAT5A | 5530 | 21627 | 13551 | 0.323 | 91.618 | K562 |
| KLF16 | MAZ | 8674 | 17849 | 40397 | 0.323 | 50.569 | K562 |
| MNT | NEUROD1 | 7443 | 37785 | 14154 | 0.322 | 71.267 | K562 |
| NFYB | SP2 | 1886 | 9997 | 3435 | 0.322 | 131.906 | K562 |
| ERF | TFCP2 | 2995 | 9999 | 8671 | 0.322 | 103.542 | K562 |
| TBX18 | ZBTB9 | 3064 | 6175 | 14782 | 0.321 | 101.113 | K562 |
| KLF6 | ZNF257 | 1827 | 7751 | 4190 | 0.321 | 108.955 | K562 |
| ZNF239 | ZNF583 | 2507 | 6412 | 9537 | 0.321 | 99.444 | K562 |
| HES1 | STAT5A | 3947 | 11186 | 13551 | 0.321 | 79.135 | K562 |
| NCOA1 | TCF12 | 8264 | 23304 | 28573 | 0.320 | 54.147 | K562 |

| | | | | | | | |
|---|---|---|---|---|---|---|---|
| NCOA1 | TCF3 | 8877 | 23304 | 33040 | 0.320 | 62.267 | K562 |
| MAFG | NFE2L1 | 5888 | 45814 | 7399 | 0.320 | 74.105 | K562 |
| E2F5 | ZNF639 | 7331 | 19514 | 26973 | 0.320 | 56.605 | K562 |
| E2F4 | MYC | 3948 | 10379 | 14737 | 0.319 | 97.039 | K562 |
| CC2D1A | DEAF1 | 7487 | 26436 | 20812 | 0.319 | 63.475 | K562 |
| E2F5 | ZNF766 | 7244 | 19514 | 26421 | 0.319 | 71.831 | K562 |
| SHOX2 | TEAD1 | 2922 | 6459 | 12990 | 0.319 | 104.498 | K562 |
| TEAD1 | ZNF583 | 3550 | 12990 | 9537 | 0.319 | 77.599 | K562 |
| CREB1 | TBP | 6263 | 17620 | 22020 | 0.318 | 77.886 | K562 |
| HIVEP1 | ZNF239 | 1700 | 4459 | 6412 | 0.318 | 98.231 | K562 |
| ARNT | CC2D1A | 6605 | 16351 | 26436 | 0.318 | 95.934 | K562 |
| E2F8 | E4F1 | 6763 | 12762 | 35525 | 0.318 | 72.917 | K562 |
| ZNF215 | ZNF583 | 2651 | 7306 | 9537 | 0.318 | 119.422 | K562 |
| ZNF395 | ZNF639 | 8293 | 25294 | 26973 | 0.317 | 64.836 | K562 |
| ZBTB9 | ZNF76 | 2960 | 14782 | 5886 | 0.317 | 89.240 | K562 |
| CREB5 | RBPJ | 661 | 1990 | 2181 | 0.317 | 144.570 | K562 |
| CHAMP1 | FOXM1 | 5179 | 14730 | 18104 | 0.317 | 81.526 | K562 |
| CREM | ZBTB12 | 5849 | 39050 | 8719 | 0.317 | 75.073 | K562 |
| ZNF257 | ZNF79 | 1184 | 4190 | 3330 | 0.317 | 122.505 | K562 |
| TBX18 | ZNF239 | 1994 | 6175 | 6412 | 0.317 | 105.014 | K562 |
| BCL6 | ZNF215 | 1441 | 2832 | 7306 | 0.317 | 109.700 | K562 |
| MAFF | NFE2L1 | 4547 | 27875 | 7399 | 0.317 | 82.732 | K562 |
| BCL6 | HIVEP1 | 1125 | 2832 | 4459 | 0.317 | 105.840 | K562 |
| GATA2 | TAL1 | 7990 | 21627 | 29475 | 0.316 | 57.446 | K562 |
| ZNF250 | ZNF397 | 851 | 2468 | 2931 | 0.316 | 118.846 | K562 |
| E2F3 | ZBTB9 | 3568 | 8607 | 14782 | 0.316 | 77.943 | K562 |
| ATF1 | SOX6 | 7942 | 16924 | 37291 | 0.316 | 53.347 | K562 |
| TCF3 | ZEB2 | 9522 | 33040 | 27537 | 0.316 | 56.966 | K562 |
| HES1 | NCOA1 | 5092 | 11186 | 23304 | 0.315 | 85.206 | K562 |
| SHOX2 | ZNF76 | 1942 | 6459 | 5886 | 0.315 | 96.779 | K562 |
| KDM5B | THAP12 | 4742 | 22367 | 10136 | 0.315 | 99.343 | K562 |
| ZNF175 | ZNF584 | 5303 | 18986 | 14944 | 0.315 | 76.748 | K562 |
| MYC | TBP | 5666 | 14737 | 22020 | 0.315 | 73.713 | K562 |
| BCL6 | TBX18 | 1315 | 2832 | 6175 | 0.314 | 127.612 | K562 |
| ERF | ZNF583 | 3070 | 9999 | 9537 | 0.314 | 108.341 | K562 |
| CREB1 | E4F1 | 7862 | 17620 | 35525 | 0.314 | 62.357 | K562 |
| TSHZ1 | ZNF239 | 1294 | 2648 | 6412 | 0.314 | 102.233 | K562 |
| E2F3 | NR4A1 | 2016 | 8607 | 4794 | 0.314 | 115.584 | K562 |
| MYC | NEUROD1 | 4527 | 14737 | 14154 | 0.313 | 72.998 | K562 |
| CREB1 | MYC | 5049 | 17620 | 14737 | 0.313 | 79.819 | K562 |
| ZNF239 | ZNF257 | 1624 | 6412 | 4190 | 0.313 | 130.013 | K562 |
| TCF12 | ZEB2 | 8783 | 28573 | 27537 | 0.313 | 54.708 | K562 |
| GMEB1 | NEUROD1 | 6379 | 29364 | 14154 | 0.313 | 85.516 | K562 |
| ATF7 | JUN | 7146 | 47196 | 11072 | 0.313 | 53.433 | K562 |
| E2F4 | MBD2 | 3967 | 10379 | 15550 | 0.312 | 96.055 | K562 |
| E2F5 | ZNF644 | 4963 | 19514 | 12949 | 0.312 | 82.940 | K562 |
| KLF16 | LEF1 | 6176 | 17849 | 21938 | 0.312 | 81.278 | K562 |
| GATA2 | TCF3 | 8336 | 21627 | 33040 | 0.312 | 57.195 | K562 |
| E2F3 | ZNF76 | 2218 | 8607 | 5886 | 0.312 | 102.552 | K562 |
| FOXM1 | GATA2 | 6166 | 18104 | 21627 | 0.312 | 87.902 | K562 |
| E2F8 | ELF4 | 5051 | 12762 | 20589 | 0.312 | 84.323 | K562 |
| ERF | NR4A1 | 2155 | 9999 | 4794 | 0.311 | 116.094 | K562 |

| | | | | | | | |
|---|---|---|---|---|---|---|---|
| NCOA1 | NEUROD1 | 5651 | 23304 | 14154 | 0.311 | 81.824 | K562 |
| MBD2 | SMAD5 | 5793 | 15550 | 22343 | 0.311 | 76.276 | K562 |
| ARNT | SOX6 | 7670 | 16351 | 37291 | 0.311 | 61.887 | K562 |
| HES1 | TEAD4 | 6238 | 11186 | 36089 | 0.310 | 68.411 | K562 |
| FOS | NEUROD1 | 3268 | 7848 | 14154 | 0.310 | 84.596 | K562 |
| ETS1 | MYC | 4005 | 11321 | 14737 | 0.310 | 88.376 | K562 |
| MYNN | NEUROD1 | 4581 | 15431 | 14154 | 0.310 | 77.182 | K562 |
| KLF6 | ZNF311 | 2323 | 7751 | 7247 | 0.310 | 103.736 | K562 |
| CREB3L1 | GMEB1 | 7914 | 22214 | 29364 | 0.310 | 59.176 | K562 |
| MBD2 | SMAD1 | 3627 | 15550 | 8819 | 0.310 | 86.177 | K562 |
| TBX18 | TSHZ1 | 1252 | 6175 | 2648 | 0.310 | 103.812 | K562 |
| CREB1 | HMGN3 | 5544 | 17620 | 18201 | 0.310 | 74.229 | K562 |
| E4F1 | FOXM1 | 7851 | 35525 | 18104 | 0.310 | 57.125 | K562 |
| FOXK2 | NCOA1 | 7183 | 23181 | 23304 | 0.309 | 68.382 | K562 |
| GATA1 | NCOA1 | 5711 | 14675 | 23304 | 0.309 | 62.934 | K562 |
| ZBTB17 | ZNF583 | 3713 | 15161 | 9537 | 0.309 | 80.625 | K562 |
| TEAD1 | TEAD4 | 6685 | 12990 | 36089 | 0.309 | 57.202 | K562 |
| FOS | SP1 | 2426 | 7848 | 7876 | 0.309 | 100.453 | K562 |
| DDIT3 | ZSCAN32 | 1328 | 4129 | 4494 | 0.308 | 128.417 | K562 |
| E4F1 | ETV5 | 7568 | 35525 | 17004 | 0.308 | 73.510 | K562 |
| CREM | JUN | 6400 | 39050 | 11072 | 0.308 | 68.077 | K562 |
| KLF6 | ZNF76 | 2075 | 7751 | 5886 | 0.307 | 109.510 | K562 |
| SMAD5 | THAP12 | 4622 | 22343 | 10136 | 0.307 | 87.006 | K562 |
| E2F3 | ZNF583 | 2782 | 8607 | 9537 | 0.307 | 100.648 | K562 |
| TSHZ1 | ZNF76 | 1212 | 2648 | 5886 | 0.307 | 111.038 | K562 |
| ERF | ZBTB17 | 3779 | 9999 | 15161 | 0.307 | 84.672 | K562 |
| DLX4 | KLF10 | 231 | 529 | 1071 | 0.307 | 153.313 | K562 |
| TSHZ1 | ZNF583 | 1542 | 2648 | 9537 | 0.307 | 105.107 | K562 |
| TBX18 | ZNF79 | 1391 | 6175 | 3330 | 0.307 | 123.521 | K562 |
| ELF4 | MBD2 | 5485 | 20589 | 15550 | 0.307 | 70.393 | K562 |
| TSHZ1 | ZNF215 | 1348 | 2648 | 7306 | 0.306 | 107.151 | K562 |
| TSHZ1 | ZNF354B | 828 | 2648 | 2758 | 0.306 | 119.007 | K562 |
| HIVEP1 | ZNF165 | 1205 | 4459 | 3470 | 0.306 | 110.400 | K562 |
| E2F5 | ZNF197 | 4428 | 19514 | 10708 | 0.306 | 83.040 | K562 |
| STAT5A | ZNF281 | 7157 | 13551 | 40285 | 0.306 | 64.785 | K562 |
| LEF1 | TAL1 | 7782 | 21938 | 29475 | 0.306 | 53.615 | K562 |
| FOXK2 | LEF1 | 6900 | 23181 | 21938 | 0.306 | 60.070 | K562 |
| ZNF197 | ZNF644 | 3601 | 10708 | 12949 | 0.306 | 93.804 | K562 |
| E2F4 | ETS1 | 3314 | 10379 | 11321 | 0.306 | 106.401 | K562 |
| ETS1 | MBD2 | 4056 | 11321 | 15550 | 0.306 | 78.038 | K562 |
| ERF | ZNF76 | 2345 | 9999 | 5886 | 0.306 | 98.253 | K562 |
| ERF | RREB1 | 2827 | 9999 | 8561 | 0.306 | 77.783 | K562 |
| CREM | MYNN | 7500 | 39050 | 15431 | 0.306 | 58.519 | K562 |
| NCOA1 | ZEB2 | 7733 | 23304 | 27537 | 0.305 | 55.162 | K562 |
| HMGN3 | MBD2 | 5134 | 18201 | 15550 | 0.305 | 90.306 | K562 |
| PATZ1 | RBPJ | 702 | 2428 | 2181 | 0.305 | 113.269 | K562 |
| CREM | E2F4 | 6140 | 39050 | 10379 | 0.305 | 61.104 | K562 |
| ELF4 | GABPA | 5616 | 20589 | 16492 | 0.305 | 72.206 | K562 |
| NR2C1 | NR2F2 | 6938 | 24799 | 20900 | 0.305 | 52.819 | K562 |
| JUND | LEF1 | 8490 | 35378 | 21938 | 0.305 | 51.087 | K562 |
| BHLHE40 | NEUROD1 | 6045 | 27807 | 14154 | 0.305 | 57.738 | K562 |
| MNT | MXI1 | 5633 | 37785 | 9080 | 0.304 | 73.413 | K562 |

| | | | | | | | |
|---|---|---|---|---|---|---|---|
| FOXO4 | ZNF57 | 734 | 2724 | 2140 | 0.304 | 119.593 | K562 |
| FOS | MBD2 | 3354 | 7848 | 15550 | 0.304 | 91.629 | K562 |
| NFXL1 | TSHZ1 | 864 | 3067 | 2648 | 0.303 | 134.769 | K562 |
| TEAD1 | ZBTB9 | 4200 | 12990 | 14782 | 0.303 | 98.440 | K562 |
| GATA2 | KLF16 | 5948 | 21627 | 17849 | 0.303 | 72.606 | K562 |
| NEUROD1 | VEZF1 | 7791 | 14154 | 46847 | 0.303 | 50.086 | K562 |
| ATF3 | MAZ | 8332 | 18779 | 40397 | 0.303 | 55.626 | K562 |
| ETS1 | HMGN3 | 4342 | 11321 | 18201 | 0.302 | 83.474 | K562 |
| RREB1 | ZNF76 | 2146 | 8561 | 5886 | 0.302 | 112.446 | K562 |
| BACH1 | MAFK | 3432 | 4739 | 27212 | 0.302 | 83.207 | K562 |
| ZNF239 | ZNF507 | 1788 | 6412 | 5465 | 0.302 | 121.439 | K562 |
| NEUROD1 | TBP | 5332 | 14154 | 22020 | 0.302 | 66.951 | K562 |
| BCL6 | ZNF583 | 1569 | 2832 | 9537 | 0.302 | 89.167 | K562 |
| ZNF215 | ZNF239 | 2066 | 7306 | 6412 | 0.302 | 97.543 | K562 |
| RREB1 | ZBTB9 | 3395 | 8561 | 14782 | 0.302 | 78.942 | K562 |
| HIVEP1 | NR4A1 | 1395 | 4459 | 4794 | 0.302 | 122.645 | K562 |
| ATF3 | NEUROD1 | 4913 | 18779 | 14154 | 0.301 | 88.160 | K562 |
| ATF1 | JUNB | 4883 | 16924 | 15517 | 0.301 | 70.137 | K562 |
| HIVEP1 | ZNF257 | 1302 | 4459 | 4190 | 0.301 | 107.485 | K562 |
| GABPA | MNT | 7517 | 16492 | 37785 | 0.301 | 52.879 | K562 |
| RREB1 | ZNF583 | 2720 | 8561 | 9537 | 0.301 | 98.597 | K562 |
| HES1 | KLF16 | 4253 | 11186 | 17849 | 0.301 | 98.749 | K562 |
| NR4A1 | RREB1 | 1928 | 4794 | 8561 | 0.301 | 107.550 | K562 |
| ARNT | E4F1 | 7249 | 16351 | 35525 | 0.301 | 68.277 | K562 |
| SKIL | ZNF282 | 5389 | 24237 | 13248 | 0.301 | 79.033 | K562 |
| HOMEZ | ZNF133 | 585 | 2447 | 1548 | 0.301 | 131.007 | K562 |
| TEAD1 | ZNF507 | 2531 | 12990 | 5465 | 0.300 | 97.573 | K562 |
| NR4A1 | ZNF76 | 1595 | 4794 | 5886 | 0.300 | 113.060 | K562 |
| ARID3A | STAT5A | 3728 | 11377 | 13551 | 0.300 | 90.409 | K562 |
| MAX | MYC | 18866 | 38331 | 33008 | 0.530 | 116.512 | MCF-7 |
| GATA3 | NR2F2 | 20118 | 47474 | 34427 | 0.498 | 98.989 | MCF-7 |
| FOS | FOSL2 | 17907 | 63152 | 21640 | 0.484 | 90.741 | MCF-7 |
| FOSL2 | JUND | 8584 | 21640 | 14651 | 0.482 | 140.291 | MCF-7 |
| ELF1 | GABPA | 8325 | 21038 | 14816 | 0.472 | 131.683 | MCF-7 |
| MAX | MNT | 17014 | 38331 | 35594 | 0.461 | 105.996 | MCF-7 |
| FOXM1 | TCF12 | 4366 | 9332 | 11601 | 0.420 | 154.692 | MCF-7 |
| MNT | MYC | 14381 | 35594 | 33008 | 0.420 | 101.032 | MCF-7 |
| ELK1 | GABPA | 4013 | 8180 | 14816 | 0.365 | 130.458 | MCF-7 |
| MBD2 | TCF12 | 4228 | 11612 | 11601 | 0.364 | 116.726 | MCF-7 |
| ZNF592 | ZNF687 | 4711 | 8037 | 21097 | 0.362 | 110.615 | MCF-7 |
| NR2F2 | TCF12 | 7228 | 34427 | 11601 | 0.362 | 90.671 | MCF-7 |
| GATA3 | TCF12 | 8474 | 47474 | 11601 | 0.361 | 76.286 | MCF-7 |
| MAZ | MNT | 9655 | 20397 | 35594 | 0.358 | 82.951 | MCF-7 |
| TCF12 | ZNF217 | 4425 | 11601 | 13180 | 0.358 | 122.703 | MCF-7 |
| FOXM1 | MBD2 | 3655 | 9332 | 11612 | 0.351 | 129.410 | MCF-7 |
| OVOL1 | SPDEF | 7980 | 22812 | 22779 | 0.350 | 99.253 | MCF-7 |
| NCOA3 | NEUROD1 | 2773 | 8114 | 7850 | 0.347 | 120.982 | MCF-7 |
| ELF1 | MAX | 9841 | 21038 | 38331 | 0.347 | 74.541 | MCF-7 |
| FOXM1 | JUND | 4025 | 9332 | 14651 | 0.344 | 110.771 | MCF-7 |
| CEBPB | CEBPG | 8259 | 37725 | 15351 | 0.343 | 96.323 | MCF-7 |
| MAX | ZNF687 | 9726 | 38331 | 21097 | 0.342 | 84.113 | MCF-7 |
| FOXM1 | NR2F2 | 6115 | 9332 | 34427 | 0.341 | 94.842 | MCF-7 |

| | | | | | | | |
|---|---|---|---|---|---|---|---|
| ELF1 | ELK1 | 4379 | 21038 | 8180 | 0.334 | 104.482 | MCF-7 |
| JUND | TCF12 | 4336 | 14651 | 11601 | 0.333 | 109.296 | MCF-7 |
| ELF1 | MYC | 8743 | 21038 | 33008 | 0.332 | 69.580 | MCF-7 |
| GATAD2B | NEUROD1 | 3015 | 10644 | 7850 | 0.330 | 96.385 | MCF-7 |
| MAZ | MYC | 8551 | 20397 | 33008 | 0.330 | 60.072 | MCF-7 |
| FOXM1 | NCOA3 | 2859 | 9332 | 8114 | 0.329 | 148.073 | MCF-7 |
| NR2F2 | SPDEF | 9156 | 34427 | 22779 | 0.327 | 64.925 | MCF-7 |
| FOXM1 | ZNF217 | 3537 | 9332 | 13180 | 0.319 | 118.709 | MCF-7 |
| JUND | NR2F2 | 7126 | 14651 | 34427 | 0.317 | 86.661 | MCF-7 |
| GATA3 | ZNF217 | 7885 | 47474 | 13180 | 0.315 | 66.847 | MCF-7 |
| FOXM1 | GATA3 | 6629 | 9332 | 47474 | 0.315 | 81.074 | MCF-7 |
| NCOA3 | TCF12 | 3027 | 8114 | 11601 | 0.312 | 115.095 | MCF-7 |
| FOSL2 | JUN | 3686 | 21640 | 6469 | 0.312 | 95.804 | MCF-7 |
| MAX | MAZ | 8644 | 38331 | 20397 | 0.309 | 59.142 | MCF-7 |
| FOXM1 | SPDEF | 4475 | 9332 | 22779 | 0.307 | 93.257 | MCF-7 |
| GATA3 | JUND | 8056 | 47474 | 14651 | 0.305 | 51.605 | MCF-7 |
| NR2F2 | ZNF217 | 6497 | 34427 | 13180 | 0.305 | 75.785 | MCF-7 |
| SPDEF | TCF12 | 4947 | 22779 | 11601 | 0.304 | 100.617 | MCF-7 |
| JUN | JUND | 2949 | 6469 | 14651 | 0.303 | 93.697 | MCF-7 |
| ELF1 | MNT | 8233 | 21038 | 35594 | 0.301 | 77.550 | MCF-7 |
